# SWATH-MS co-expression profiles reveal paralogue interference in protein complex evolution

**DOI:** 10.1101/2020.09.08.287334

**Authors:** Luzia Stalder, Amir Banaei-Esfahani, Rodolfo Ciuffa, Joshua L Payne, Ruedi Aebersold

**Affiliations:** Department of Biology, Institute of Molecular Systems Biology, ETH Zurich, 8093 Zurich, Switzerland; Institute of Integrative Biology, ETH Zurich, 8092 Zurich, Switzerland; Swiss Institute of Bioinformatics, 1015 Lausanne, Switzerland; Faculty of Science, University of Zurich, 8057 Zurich, Switzerland

## Abstract

Understanding the conservation and evolution of protein complexes is of critical value to decode their function in physiological and pathological processes. One prominent proposal posits gene duplication as a potential mechanism for protein complex evolution. In this study we take advantage of large-scale proteome expression datasets to systematically investigate the role of paralogues, and specifically self-interacting paralogues, in shaping the evolutionary trajectories of protein complexes. First, we show that protein co-expression derived from quantitative proteomic matrices is a good indicator for complex membership and is conserved across species. Second, we suggest that paralogues are commonly strongly co-expressed and that for the subset of paralogues that show diverging co-expression patterns, the divergent co-expression patterns reflect both sequence and functional divergence. Finally, on this basis, we show that homomeric paralogues known to be part of protein complexes display a unique co-expression pattern distribution, with a subset of them being highly diverging. These findings support the idea that homomeric paralogues can avoid cross-interference by diversifying their expression patterns, and corroborates the role of this mechanism as a force shaping protein complex evolution and specialization.

## Introduction

Protein complexes – described as stable protein assemblies that can be isolated by biochemical means – are one of the main modes of proteome organization and fundamental functional entities of the cell. The dramatic increase in structural knowledge of these complexes, as well as the advent of proteome-wide profiling methods (most prominently via mass spectrometry) have allowed the interrogation of general principles governing protein complex formation, function and evolution. For instance, bioinformatics approaches of the Protein Data Bank (PDB) and of large-scale proteomics datasets, respectively, have identified core symmetries that complexes obey, and the extent to which the co-regulation of their subunits is constrained^1,2^. An especially fertile area of investigation relates to the evolution of protein complexes.

It has been demonstrated that a possible mechanism for protein complex core formation starts with the duplication of self-interacting proteins (homomeric paralogues; Figure 1A)^3,4^. Importantly, recent research on the fate of paralogues pointed out that duplication of genes for obligate homomeric proteins can lead to interference between the resulting paralogues: mutations in only one of the paralogues can “poison” the oligomer and affect the ancestral function^5^. It has been suggested that this constrains the evolution of paralogues on the one hand, but also promotes additional regulatory complexity on the other^5^. Specifically, if sequence divergence of the homomeric paralogues impairs the ancestral function, the mutated paralogue acts as a highly specific competitive inhibitor for the ancestral protein. This is referred to as paralogue interference^5^. To prevent this type of interference, the paralogues will be under selection pressure to develop mechanisms that prevent their cross-interaction, for example by mutations that drive expressional separation. Importantly, if paralogues that are separated by expression give rise to protein complexes, they will contain either the ancestral or the duplicated paralogous pair, but not both at the same time. Expressional separation after duplication is a common pattern on the RNA level^5^. However, expressional separation has not been analyzed on the protein level in a general way and its role in protein complex evolution remains elusive^5^.

**Figure 1.**
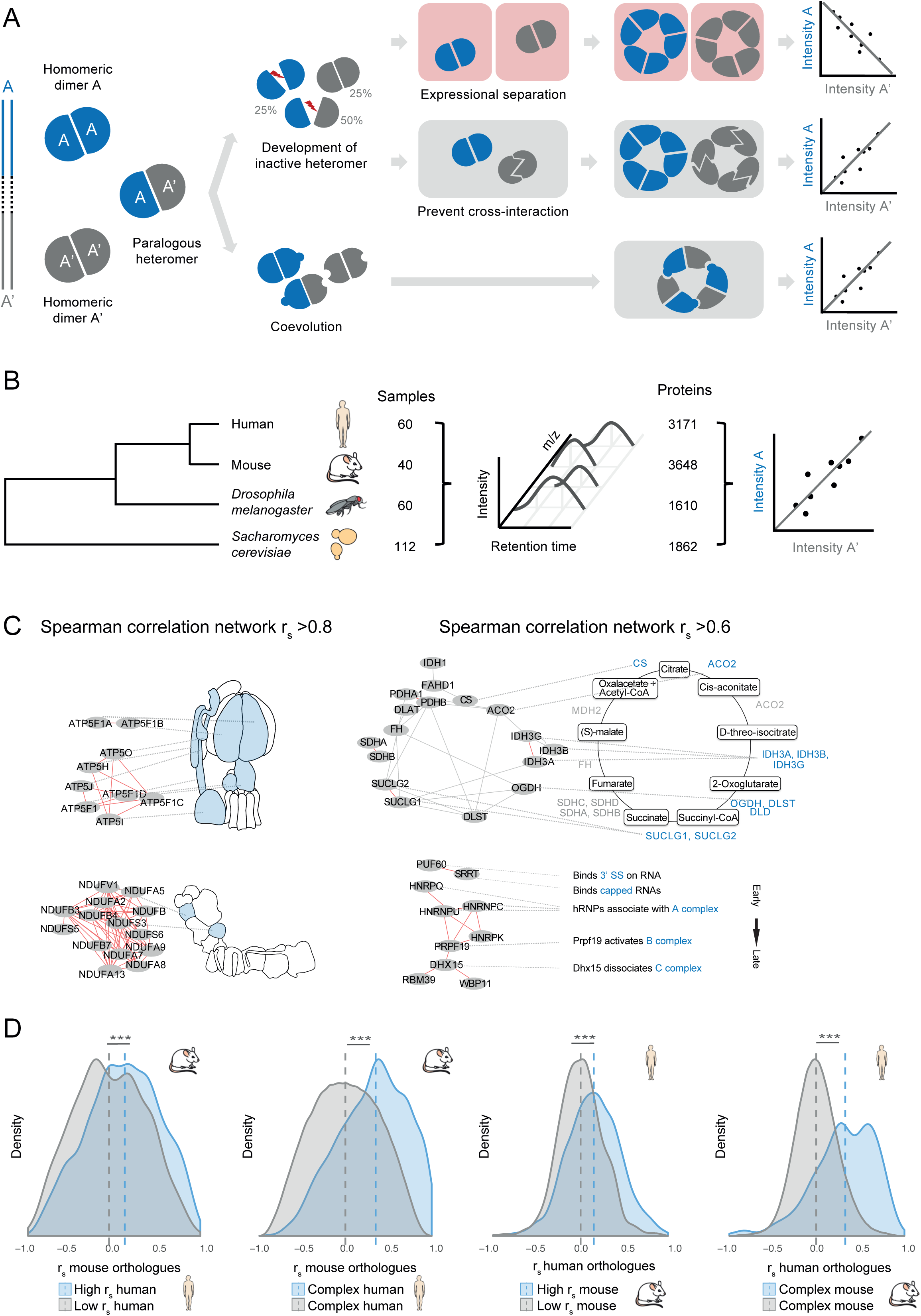
SWATH-MS co-expression profiles of protein complex modules are conserved across species. **(A)** Schematic drawing illustrating possible evolutionary routes from homomer-derived paralogues into protein complexes. **(B)** Study design. Protein co-expression profiles were acquired in SWATH-MS from 272 samples covering the proteome of four species. Spearman’s rank correlation (r_s_) was used to measure profile similarity. **(C)** Conserved correlation networks of human and mouse. Nodes represent proteins, edges indicate high correlation in both human and mouse datasets. Red edges are annotated in CORUM. In panels on the left, a stringent correlation cutoff of r_s_ > 0.8 was chosen, on the right a more relaxed cutoff of r_s_ > 0.6 was used. For comparison, correlation values of human and mouse were quantile normalized. ***Left:*** Proteins of the ATP-synthase complex (top) and proteins of the NADH dehydrogenase complex (bottom). ***Right***: Proteins involved in the tricarboxylic acid cycle (top) and proteins involved in the spliceosome (bottom). **(D) *(I)*** Mouse Spearman correlation for mouse orthologues that are highly correlated in human (top 0.5 percentile, r_s_ > 0.8, n_pairs_ = 1429) and for mouse orthologues that not correlate in human (bottom 0.5 percentile, n_pair_ = 1429). Pairs that highly correlate in human correlate also significantly higher in mouse (md 0.2 vs 0.0, Wilcoxon signed-rank test p-value < 0.001). ***(II)*** Mouse Spearman correlation for pairs that are annotated as CORUM complex pairs in human (n_pairs_ = 1807) and for pairs that are not annotated in CORUM (n_pairs_ = 712803). Human CORUM pairs correlate significantly higher in mouse (md 0.35 vs 0.0, Wilcoxon signed-rank test p-value < 0.001). ***(III)*** Human Spearman correlation for human orthologues that are highly correlated in mouse (top 0.5 percentile, r_s >_ 0.8, n_pairs_ = 1429) and for human orthologues that not correlate in mouse (bottom 0.5 percentile, n_pairs_ = 1429). Pairs that highly correlate in mouse correlate also significantly higher in human (md 0.2 vs 0.0, Wilcoxon signed-rank test p-value < 0.001). ***(IV)*** Human Spearman correlation for pairs that are annotated as CORUM complex pairs in mouse (n_pairs_ = 331) and for pairs that are not annotated in CORUM (n_pairs_ = 714279). Mouse CORUM pairs correlate significantly higher in human (md 0.3 vs 0.0, Wilcoxon signed-rank test p-value < 0.001).

A main limitation in studying protein complex evolution has been the difficulty of measuring protein complex states in a high-throughput manner. Here we address this by using data-independent acquisition and analysis of mass spectrometry data, particularly by the sequential windowed acquisition of all theoretical mass spectra (SWATH-MS)^6^. We study complex states at the proteome level in human, mouse, *Drosophila melanogaster* and *Saccharomyces cerevisiae*^7–9^. We demonstrate that protein co-expression profiles determined by SWATH-MS are substantially conserved across species and that they can be used to map the diversification of protein paralogues, as measured for instance by divergence in sequence or interactors. Critically, we find that protein paralogues that are known to be part of a protein complex are enriched in homomer-derived paralogues that separated in their expression levels. These are indeed the proteins shown to be affected by interference, and our finding therefore supports the notion that this mechanism can propel protein complex diversification and expression divergence. In conclusion, our study indicates that quantitative proteomic data can be used to infer protein complex relationships and identifies paralogue interference as a constraint of their evolution.

## Results

### SWATH-MS co-expression profiles recover protein complexes and are conserved across species

To analyze the role of paralogue diversification in the evolution of protein complexes, we first set out to define a suitable methodological framework. In recent years, a number of studies have taken advantage of the rapidly accumulating body of data on protein-protein interactions and protein expression to analyze constraints on, and evolution of, protein complexes^1–4,10–13^. Some of these studies have shown that the expression levels of protein complex subunits are generally covarying, and that, conversely, co-expression patterns can be used as a proxy for functional, interaction and complex relatedness. Here we used abundance profiles derived from SWATH-MS data across four species: human, mouse, *Drosophila melanogaster* and *Saccharomyces cerevisiae*^7–9^ (Figure 1B). The proteome dataset of each species contains 40 to 112 samples. Overall, between 1610 and 3171 proteins were consistently quantified per species dataset across samples (Table S1 and Figure S1). To examine covariance of protein pairs, we used Spearman‘s rank coefficient (r_s_). First, we aimed at showing that covariation patterns in our data can indeed preferentially recall known protein complexes and detail their conservation across species. We used manually curated catalogues of protein complexes as a benchmark, specifically CORUM^14,15^ for human and mouse complexes, DroID^16^ for *Drosophila melanogaster* complexes and CYC2008^17^ for *Saccharomyces cerevisiae* complexes. The results showed that in the datasets of all four species complex members have significantly higher r_s_ compared to those not annotated as members of the same complex (Figure S2, Wilcoxon sign-rank test p-values < 0.001, number of pairs human = 17453, mouse = 1739, *Drosophila melanogaster* = 2261, *Saccharomyces cerevisiae* = 1807). To further characterize conserved covariance profiles, we performed functional enrichment by DAVID^18,19^. We found that protein pairs with highly covarying abundance profiles in human and mouse (r_s_ > 0.8) are enriched in mitochondrial processes. Interestingly, when we relaxed the cutoff to r_s_ > 0.6 we noted that protein pairs functionally annotated with “splicing” and “signaling pathways regulating metabolism”, “proliferation”, “cell-cell adhesion” and “immune responses” were additionally enriched (Table S2, EASE score, a modified Fisher exact test p-value < 0.05). Figure 1C illustrates this point by showing that the correlation network with edges r_s_ > 0.8 recovers the ATP synthase complex and the NADH dehydrogenase complex, whereas the correlation network with edges r_s_ > 0.6 additionally recovers complexes in the tricarboxylic acid cycle and the spliceosome. Next, we asked whether protein pairs with highly correlated abundance in one species are also highly correlated in another species. For this, we selected the most and least correlating orthologues in one species (top and bottom 0.5 percentile) and tested whether these most correlating pairs also showed higher covariance than the least correlating pairs in a second species. We found that this was the case for all species combinations, indicating that covariance profiles are conserved across species (Figure 1D and Figure S3, Wilcoxon signed-rank test p-values < 0.001, number of mouse orthologue pairs that are highly correlated in human/ number of human orthologue pairs that are highly correlated in mouse = 1429). Consequently, proteins that belong to a protein complex in one species also correlate higher in a second species (Figure 1D and Figure S3, Wilcoxon signed-rank test p-values < 0.001, number of pairs in mouse that are annotated as CORUM complex pairs in human = 1807, number of pairs in human that are annotated as CORUM complex pairs in mouse = 331). Taken together, our analyses indicate that protein complex members exhibit coordinated expression, and that such coordinated expression is conserved across species.

### SWATH-MS correlation profiles reflect evolutionary trajectories of paralogues

We next asked whether our framework is able to capture important principles driving the evolution of protein complexes. To this end, we focused on protein paralogues, because paralogue diversification has been proposed as a significant factor of complex evolution^4,5^. First, we wanted to verify that our protein abundance matrices recapitulated paralogue divergence over time, as well as diversification of protein interactions. We identified paralogues using Ensembl 92 (ref ^27^) and we classified them into paralogue families, whereby a family was defined as the genes emerged from a single ancestral gene by duplication (Figure S4 and Table S3, number of paralogue families human = 73, mouse = 114, *Drosophila melanogaster* = 3, *Saccharomyces cerevisiae* = 9, mean size of paralogue families human = 9, mouse = 9, *Drosophila melanogaster* = 11, *Saccharomyces cerevisiae* = 9). On a general scale, we found that paralogous proteins exhibit a stronger degree of covariance than non-paralogous proteins in all species examined (Figure 2A, Wilcoxon signed-rank test p-values < 0.001, number of paralogous pairs human = 1302, mouse = 1964, *Drosophila melanogaster =* 119 and *Saccharomyces cerevisiae* = 191). To assess whether paralogue covariance patterns recapitulated sequence diversification, we tested whether a higher frequency of differentiating mutations between paralogous pairs corresponded to a decrease in protein covariance. To do so, we quantified all pairwise correlations among paralogue family members and determined for each pair the rate of synonymous and non-synonymous amino acid sequence changes, i.e. the number of nucleotide changes among the two paralogues that affects, respectively not affects, the resulting codon sequence, relative to the paralogue length. As expected, the co-expression of paralogous pairs within a paralogue family was negatively associated with the rate of synonymous and non-synonymous nucleotide changes (Figure 2B left and center, respectively; due to limited numbers of observations we could not examine the *Drosophila melanogaster* and *Saccharomyces cerevisiae* dataset in this and subsequent analysis). Furthermore, the association of covariance with non-synonymous changes was stronger than with synonymous changes, in line with intuition that non-synonymous mutations have generally a greater phenotypic effect (Paired sample t-test p-values non-synonymous changes: human = 0.005 and mouse = 0.009, synonymous changes: human = 0.04 and mouse = 0.05; number of paralogue groups: human = 23 and mouse = 64). Finally, we reasoned that, if covariance is a good proxy for sequence diversification, paralogues with more strongly diverging interactomes – i.e. a smaller fraction of shared interactors – should also exhibit more strongly diverging expression patterns, as interactome rewiring is likely a consequence of sequence change. To test whether the paralogue covariance correlates to the diversity of protein interactions, we calculated for each paralogous pair the Jaccard index of interaction partners, defined as the intersection of the interaction partners divided by the union of the interaction partners of each pair. By these means, we found that lower covariance within a paralogue family was associated with more strongly diverging interactomes (Figure 2B right, Paired sample t-test p-values human = 0.008 and mouse = 0.09; number of paralogue groups in human = 35 and mouse = 7). Taken together, the data show that protein covariance recapitulates paralogue divergence over time as well as diversification of protein interactions that drives the evolution of new protein complexes.

**Figure 2:**
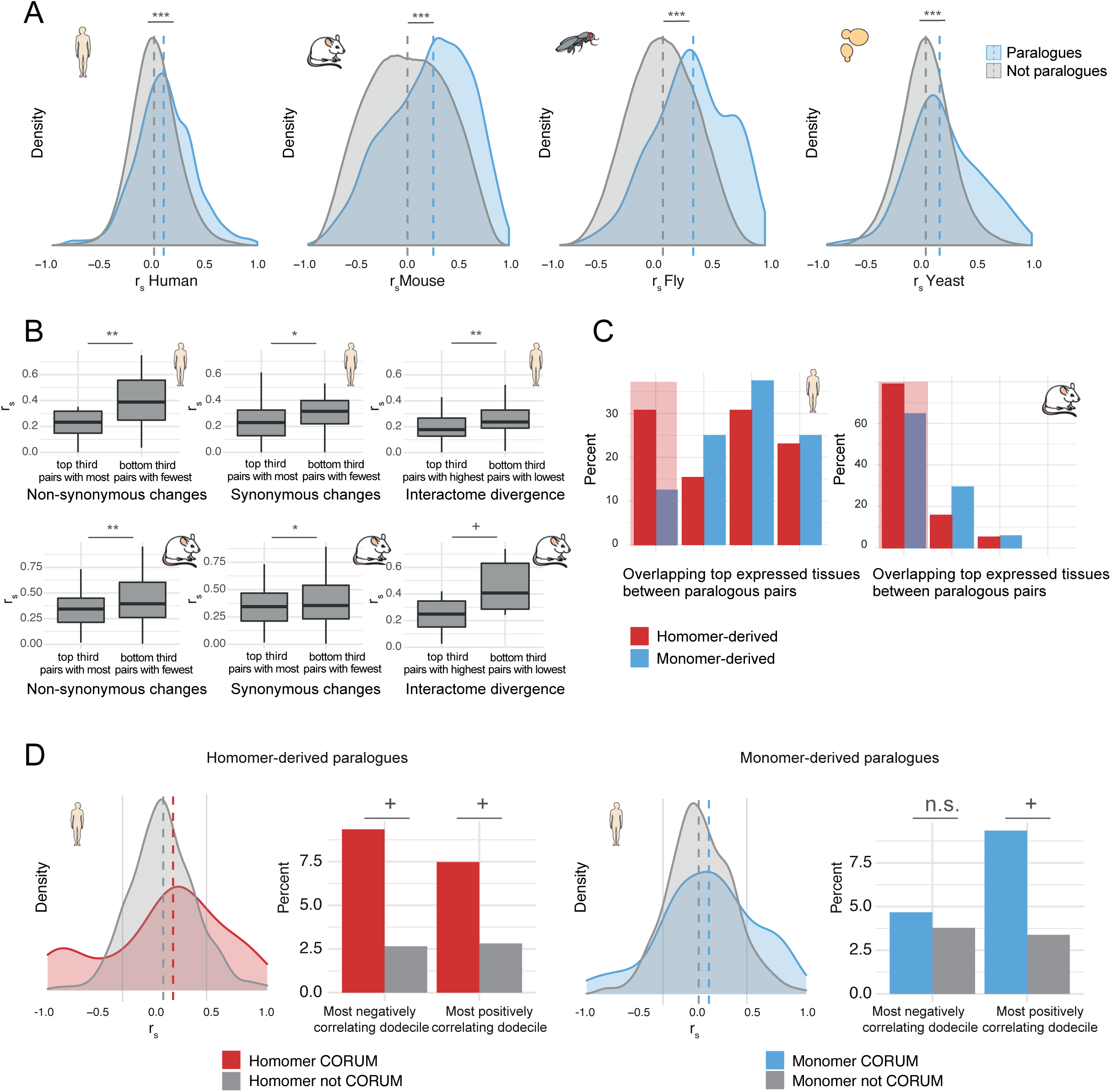
Co-expression profiles of paralogues are conserved across species and reflect sequence divergence within the paralogue family. Negatively correlating homomer-derived paralogues are functional divergent and prominent in complexes. **(A)** Spearman correlation of paralogous pairs in yeast, fly, mouse and human compared to all measured non-paralogous pairs (number of paralogous pairs human = 1302, mouse = 1964, fly = 119 and yeast = 191). Paralogues correlate significantly higher in all species (Wilcoxon signed-rank test p-values < 0.001, md 0.12 vs 0.03, md 0.25 vs 0.01, md 0.34 vs 0.08 and md 0.17 vs 0.04, respectively). **(B) *Left:*** Spearman correlation coefficients of paralogous pairs within the top, respectively bottom third of non-synonymous sequence changes within their paralogue family. Families that have a range in Spearman correlation of < 0.4 and a range in non-synonymous changes of < 0.2 were excluded. Paralogous pairs with fewer non-synonymous changes tend to correlate higher (Paired sample t-test p-values human = 0.005 and mouse = 0.009; number of paralogue families in human = 23 and mouse = 64). ***Middle:*** Spearman correlation coefficients of paralogous pairs within the top, respectively bottom third of synonymous sequence changes within their paralogue family. Families that have a range in Spearman correlation of < 0.4 and a range in synonymous changes of < 0.2 were excluded. Paralogous pairs with fewer synonymous changes tend to correlate higher (Paired sample t-test p-values human = 0.04 and mouse = 0.05, number of paralogue families in human = 28 and mouse = 78). ***Right:*** Spearman correlation coefficients of paralogous pairs within the top, respectively bottom third of interactome divergence within their paralogue family. To quantify interactome divergence, the Jaccard index, defined as the intersection of the interaction partners divided by the union of the interaction partners of each pair was calculated. All interaction partners from Biogrid were considered. Families that have a range of the Jaccard index of < 0.2 and range in Spearman correlation of < 0.2 were excluded. Paralogous pairs with more similar interaction partners tend to correlate higher (Paired sample t-test p-values human = 0.008 and mouse = 0.09; number of paralogue families in human = 35 and mouse = 7). **(C)** The overlap of the four most expressed tissues between paralogous pairs (as defined by HPM) is compared between strongly negative correlating homomer-derived and monomer-derived pairs paralogous pairs (r_s <_ -0.5, human n_pairs =_ 21, mouse n_pairs_ = 36). Negatively correlating homomer-derived paralogues tend to be expressed more diversely across different tissues compared to monomer-derived paralogues. **(D) *Left***: Spearman correlation of homomer-derived paralogues that are annotated as CORUM complex pairs (n_pairs_ = 42) and for pairs that are not annotated in CORUM (n_pairs_ = 474). Homomer-derived paralogues are enriched in the bottom and top dodecile of the correlation distribution (Fisher’s exact test p-values 0.06 and 0.1). ***Right***: Spearman correlation of monomer-derived paralogues that are annotated as CORUM complex pairs (n_pairs_ = 51) and for pairs that are not annotated in CORUM (n_pairs_ = 569). Monomer-derived paralogues are enriched not in the bottom but in the top dodecile of the correlation distribution (Fisher’s exact test p-values 0.5 and 0.06). *** indicate p-values ≤ 0.001, ** ≤ 0.01, * ≤ 0.01, + ≤ 0.1.

### Negatively correlating SWATH-MS profiles from homomer-derived paralogues are functionally divergent and prominent in complex members

Since correlation of quantitative proteomics data can inform us about the divergence of evolutionary trajectories, we used it to assess, on a proteome wide scale, the notion of paralogue interference (Figure 2C), first on a whole proteome level and then focusing specifically on known protein complexes. We first reasoned that, if the divergence in protein abundance can serve as a mechanism to escape interference, then homomeric paralogues should be differentially abundant to a greater extent than monomer-derived paralogues. To test that, we retrieved homo-, hetero-and monomer annotations from InterEvols^32^ and classified paralogous pairs either as homomer-derived if at least one member was annotated as homomer, or as monomer-derived when none of the members was annotated as either homo-or heteromer. In line with our expectations, we found that among negatively correlating pairs, homomer-derived paralogues showed an enrichment factor of 6.9 in human and 1.8 in mouse, respectively, over monomer-derived paralogues (Figure S5; negative correlation r_s_ < -0.7; number of human homomeric pairs = 540, human monomeric pairs = 620, mouse homomeric pairs = 615, mouse monomeric pairs = 1121, Fisher’s exact test p-values = 0.04 and 0.2, respectively). Of note, we also found that among all negatively correlating paralogous pairs, homomer-derived pairs were more likely to be of different abundance in tissues than monomer-derived paralogues, as defined by the tissue specific expression analysis of the Human Proteome Map^20^ (HPM) (Figure 2D and Figure S6, number of pairs human = 21 and mouse = 36). This indicates that homomeric paralogues are more prone to be affected by spatial separation of expression, and gives credence to the notion that this separation has evolved in response to protein interference.

Finally, we asked what impact the mechanism of paralogue interference has had on the diversification of protein complexes. If interference from homomeric paralogues has played any role in the organization of complexome diversity, then there must be a subset of complexes whose homomer-derived paralogue members exhibit highly divergent expression patterns; and, as a corollary, such divergence should not be observed in the case of monomer-derived paralogues. We therefore plotted the distribution of correlations of protein abundance for all paralogues present in CORUM complexes across the two classes listed above. Strikingly, we found that protein complexes containing homomer-derived paralogues contained two discrete subsets of highly positively and highly negatively correlating paralogue members (Figure 2E and Figure S7). In support of the homomer-derived paralogue specificity of such a pattern, we found no evidence for a similar distribution for the monomer-derived paralogues. Furthermore, by calculating the covariance correlation for all CORUM complex members, irrespective of them being paralogous or non-paralogous proteins, we found that homomer-derived paralogues are strongly enriched among the negatively correlating complex pairs (Figure S8; negative correlation r_s_ < -0.7; number of human homomeric CORUM pairs = 5645, human CORUM pairs = 11820, mouse homomeric CORUM pairs = 457, mouse CORUM pairs = 1572, Fisher’s exact test p-values < 0.001 and 0.05, respectively). This leads us to suggest a classification of protein complexes containing homomer-derived paralogues in two distinct groups: those where paralogues are not interfering with each other’s function and therefore the need to minimize their spatiotemporal co-existence is alleviated; and those that have diverged in their abundance levels under the pressure of negative interference. To further corroborate our findings and conceptualization, we manually curated the whole set of protein complexes in the ‘escaped’, i.e. negatively correlating class. Consistent with our proposal, we found for 84% of the human and 67% of the mouse complexes in this category, respectively, literature evidence supporting mutual exclusivity/complementarity (Number of complexes with negatively correlating pairs human = 13 and mouse = 6, for the complete list of paralogue correlations see Table S4 and Figure S9-10). In the human dataset, for example, negatively correlating, homomer-derived CORUM paralogous pairs included two subcomplexes of the emerin complex, one with lamin A and the other with lamin B1. Lamin A has been shown to regulate nuclear mechanisms and is also associated with several diseases, including Emery–Dreifuss muscular dystrophy. In contrast, Lamin B1 is involved in intermediate filaments from the cytoskeleton, but not in nuclear mechanisms^21^. Another example was the homomer-derived paralogues Hspa5 and Hspa8, which are part of the HCF-1 complex involved in cell cycle and transcriptional regulation^21^. Whereas Hspa5 localizes in the ER lumen, Hspa8 resides in the nucleolus and the cell membrane^21^. In both the human and the mouse dataset, the paralogues were among the most negatively correlated pairs. Further examples from the mouse dataset include negatively correlating homomeric paralogous pairs of the ubiquitin E3 ligase complex, that is Cul1 and Cul2, as well as Cul2 and Cul3. This is consistent with studies that established that the paralogues Cul1, Cul2 and Cul3 are involved in three distinct subcomplexes^22^. Additionally, the mouse dataset showed two negatively correlating members of the Ubiquitin-proteasome complex, UBQLN1 and UBQLN2. Only UBQLN2 was shown to be able to translocate to the nucleus, and it has been shown that after heat stress, the two proteins are in distinct subcellular locations^23^. Taken together, our data indicate that the class of proteins postulated to be more prone to protein interference, that is, homomer-derived paralogues, and especially those that are part of protein complexes, exhibit a stronger divergence in protein abundance than other paralogues, as well as subcellular specialization. By this means, our correlation studies pinpoint interference escape as an important mechanism of protein complex evolution.

## Discussion

In this study we show that rigorous statistical analysis of sets of protein abundance maps across species and tissues can inform us about the evolution of protein complexes. We used this framework to address the role of paralogue interference and diversification in protein complex evolution and specialization. Besides demonstrating that protein complex members tend to display highly correlated expression profiles, and that these profiles are conserved across species, we also indicate that protein abundance matrices recapitulate paralogue divergence over time, as well as diversification of protein interactions. Our study culminates with the observation that homomeric paralogues that are part of protein complexes show highly divergent expression patterns. This supports the notion that this is a mechanism by which protein interference among homomeric paralogues is avoided and complexes are diversified.

While many general aspects of protein complex architectural principles and evolution have been addressed in previous studies, to the best of our knowledge this is the first time that the escape of paralogue interference, specifically by separation of expression, has been analyzed on the proteome level and across species. Typically, positive co-variance of expression has been used to study conservation of protein complexes. Here we show that, since divergence of co-expression patterns seem to scale to some extent with functional and structural diversification, negative correlation may pinpoint specific evolutionary constraints in maintaining separation of homomer-derived paralogues. In fact, our data suggest that anti-correlating complex pairs are enriched in homomer-derived paralogues (Figure 2E). This is in agreement with the suggestion that homomer-derived paralogues are the most likely to be affected by protein-protein interference which can be resolved by separating expression. We find only a fraction of homomer-derived paralogues to exhibit such anticorrelation, while others strongly correlate. We therefore suggest that complex paralogues can be divided into two main groups. First, highly correlating co-evolved subunits and second, negatively correlating paralogues which, by being expressed in complementary fashions minimize the risk of interference. We indicate several examples of paralogues belonging to the latter class, for example lamin A and B1, which could represent suitable targets for follow-up studies.

At present, the scope and generalizability of our conclusions is limited by several factors. First, protein complex formation and evolution are likely to have many determinants, which can mask the effect of paralogue interference or compensate for it in ways other than expression divergence. Second, our analyses rests on resources curating protein complexes and distinguishing homomer-from monomer-derived paralogues. Broadening and improvement in such curations and annotation will allow more extensive and statistically more robust analyses and conclusions. This has a clear impact on comparative studies, where the extent of annotation between species varies greatly. Finally, the correlational nature of this, and many other methodologically related studies, must be stressed. We show that proteome-wide, cross-tissue and cross-species analyses are capable of capturing patterns that would otherwise be indiscernible. In this respect, we identified specific trends behind complex evolution which give support to specific proposals, such as homomer-derived paralogues-driven complex formation and interference escape. However, targeted studies should decode the mechanisms underlying these trends. Such an investigation, together with correlation analyses covering larger sets of conditions, are what in our view holds the greatest promise to refine our understanding of the forces shaping protein complex evolution.

## Material and methods

### SWATH-MS datasets and co-expression measures

We obtained all protein abundance data from publicly available SWATH-MS datasets. For human we used the data of Guo et al. 2019^7^, for mouse the data from Williams at al. 2018^9^, for fly the data of Okada et al. 2016^8^ and for yeast the SWATH-MS dataset of the yeast strains described Zhu et al., 2008^24^ and Brem et al., 2002^25^ (manuscript in preparation). For further description of the datasets see Table S1. All analyses were based on the available protein matrix with relative protein intensities.

For each protein pair, we calculated the Spearman correlation of raw protein abundances across all samples. We used the *cor* function of the R package stats v. 3.4.4.^26^ with the option *pairwise*.*complete*.*obs* to compute the correlation between each pair of proteins using all complete pairs of observations on those proteins.

### Orthologue identification

Orthologue mapping was conducted with BioMart Ensembl 92^27^ (release April 2018). We considered only genes with a “one2one” mapping, i.e. when the gene in one species has only one defined ortholog in another species. To translate protein and gene identifiers, we used the R package biomaRt v2.34.2^28,29^.

### Paralogue identification and analysis

We identified paralogues and paralogue families with Ensembl 92 (release April 2018) ^27^. As paralogue family we defined all paralogues connected via direct pairs. We determined the rate of non-synonymous changes and synonymous changes between the paralogous pairs using Ensembl 92 (ref^27^). We next assessed the similarity of interaction partners among paralogous pairs. We obtained the interaction partners of each paralogue from BioGrid v3.5^30,31^ and we calculated the Jaccard similarity index, that is the intersection divided by the union of distinct interaction partners.

To determine whether a paralogous pair is derived from an ancestral protein that either formed homomers or was only present as a monomer, we used the InterEvol database (release 2010), designed for the analysis of co-evolution events at the structural interfaces of hetero-and homo-oligomers^32^. We considered a paralogous pair as homomer-derived if at least one member was annotated as homomer, and a pair as monomer-derived when none of the members was annotated as either homo-or heteromer. For manual paralogue annotation, we additionally considered the UniProt database (release November 2018)^21^.

### Functional annotation

For functional enrichment analysis we used DAVID v6.8^18^ with the following parameters: Annotation categories: GOTERM_BP_DIRECT; GO_Kappa similarity: Similarity term overlap = 3, similarity threshold = 0.5; Classification: Initial group membership = 2, final group membership = 2, multiple linkage threshold = 0.5; Enrichment thresholds: EASE = 0.05. We retrieved lists of protein complexes from CORUM v2.0^14,15^. Tissue specific expression data was retrieved from the Human Proteome Map portal^20^ (HPM). To compare expression between tissue, the data was normalized using the *normalizeBetweenArrays* function from the R package limma v. 3.40.6^33^.

### Statistics and visualization

We conducted all statistical analysis with R v3.4.4^26^. For the Fisher’s exact tests and the Wilcoxon tests we used a one-sided alternative hypothesis if applicable. We drew density graphs, bar-and boxplots with the R package ggplot2 v3.1.0^34^. Boxplots were drawn in default settings (lower and upper hinges correspond to the first and third quartiles, whiskers extend up to 1.5 x the inter-quartile range or the distance between the first and the third quartile). In addition, we used the Van de Peer’s webtool to draw the Venn diagrams^35^. For network representations, we used Cytoscape v3.6.1^36^.

## Acknowledgments

We thank Marija Buljan for help with the project supervision. This project was supported by the Swiss National Science Foundation through grant # SNSF 31003A_166435 to R.A. and by the European Research Council (ERC) through grant (ERC-20140AdG 670821 to R.A. R.C. was supported in part by the IMI project ULTRA-DD (FP07/2007-2013, grant no. 115766). J.L.P. acknowledges support from Swiss National Science Foundation Grant PP00P3_170604.

## Author contributions

L.S. conceived the study, performed the analysis and wrote the manuscript. R.C. helped with manuscript writing. R.A. and J.L.P supervised the project and provided feedback on the manuscript. A. B.E. helped with project supervision.

