## Supplementary information for "SWATH-MS co-expression profiles reveal paralogue interference in protein complex evolution"

Supplementary Figure 1

A

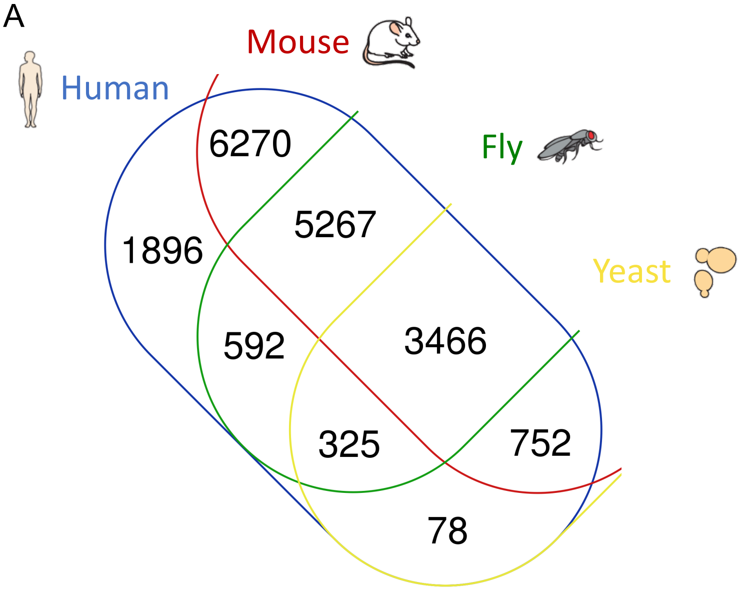

| Overlap | Overlap proteins/ unique Human proteins | Percent from unique Human proteins |
|---|---|---|
| Human-Mouse | 15755/18646 | 84.5 |
| Human-Fly | 9650/18646 | 51.8 |
| Human-Yeast | 4621/18646 | 24.7 |
| All four species | 3466/18646 | 18.6 |

B

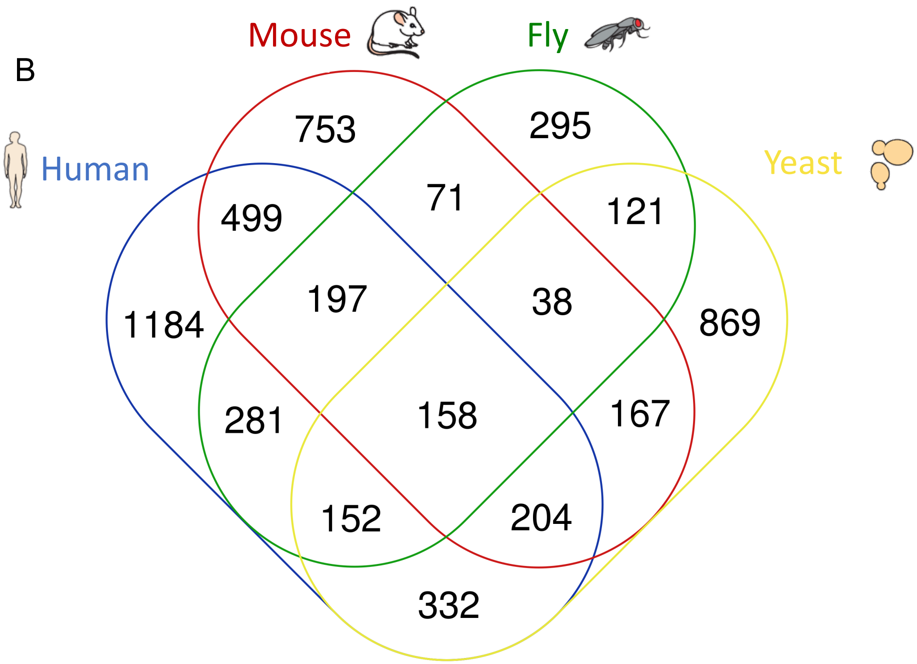

| Overlap | Overlap proteins/ unique Human proteins | Percent from unique Human proteins |
|---|---|---|
| Human-Mouse | 1058/3007 | 35.2 |
| Human-Fly | 788/3007 | 26.2 |
| Human-Yeast | 846/3007 | 28.1 |
| All four species | 158/3007 | 5.3 |

Supplementary Figure 2

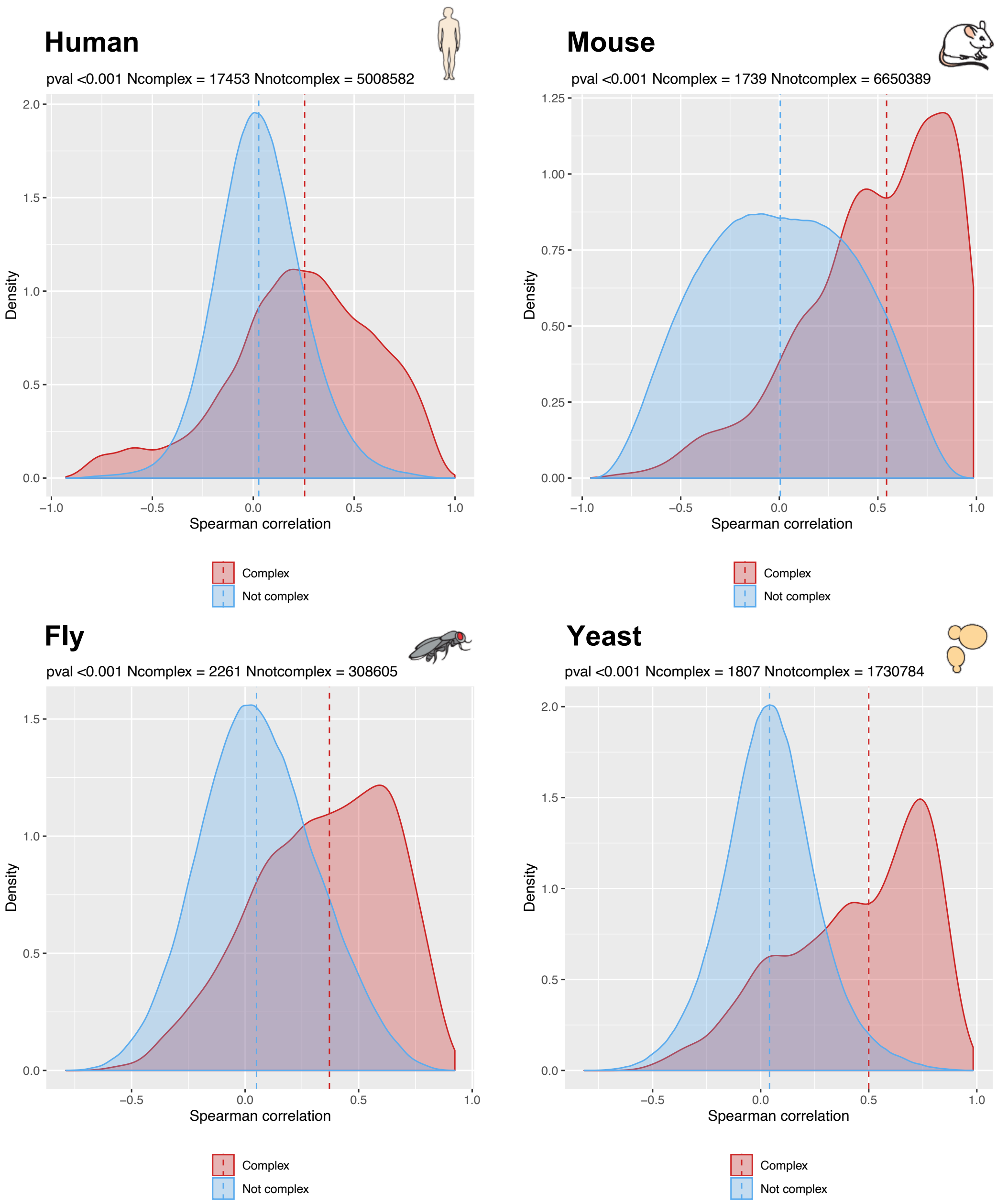

Supplementary Figure 3

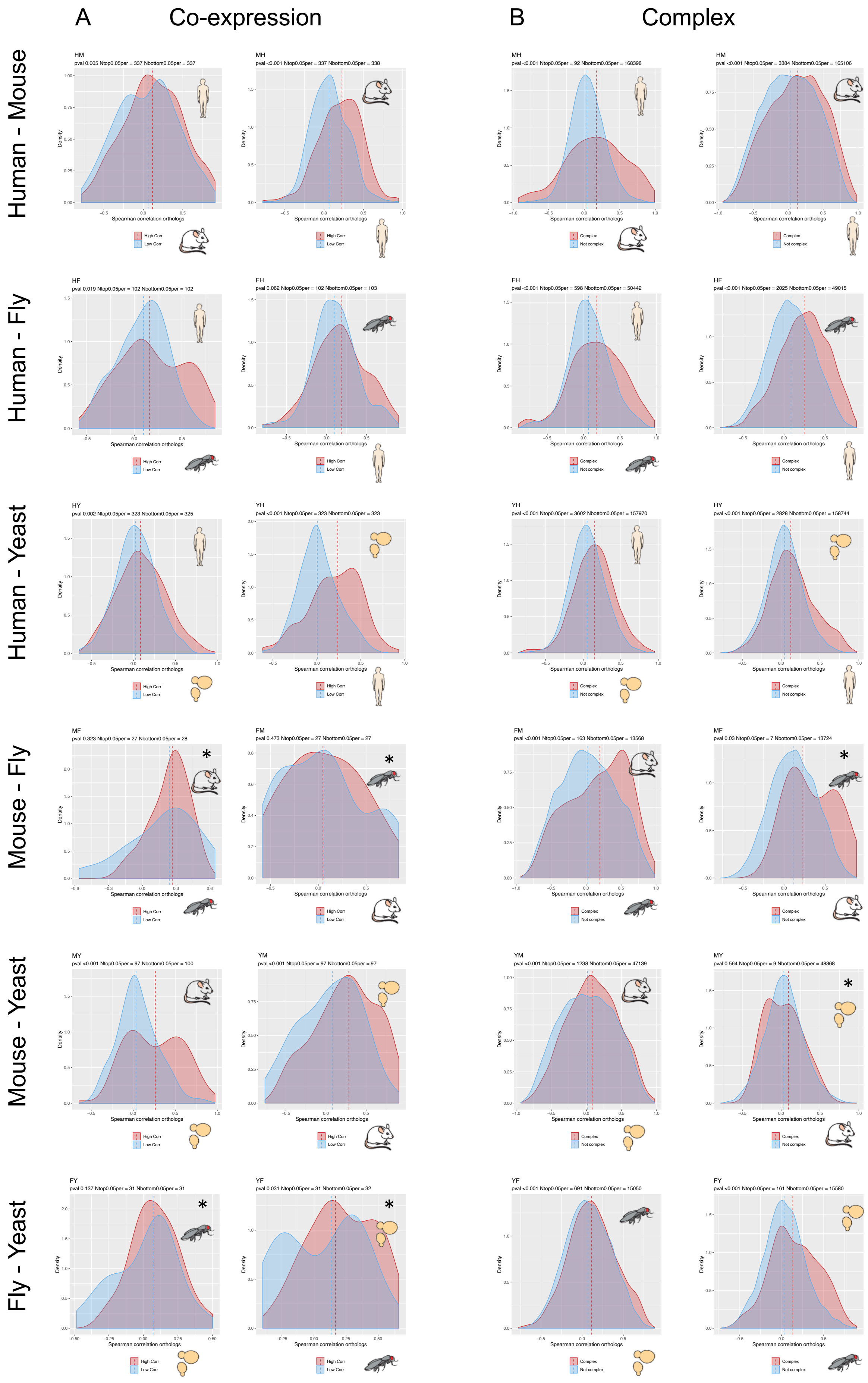

Supplementary Figure 4

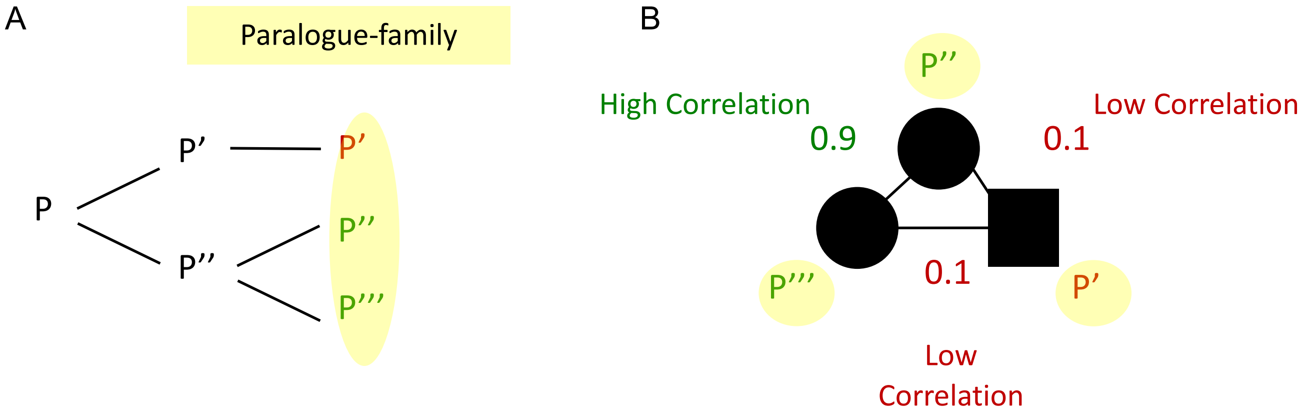

Supplementary Figure 5

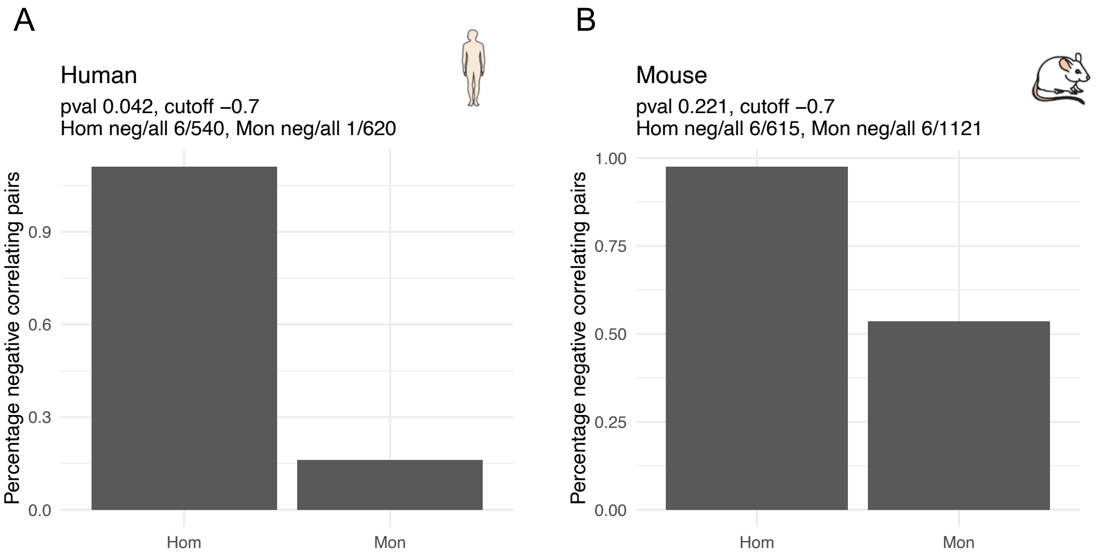

Supplementary Figure 6

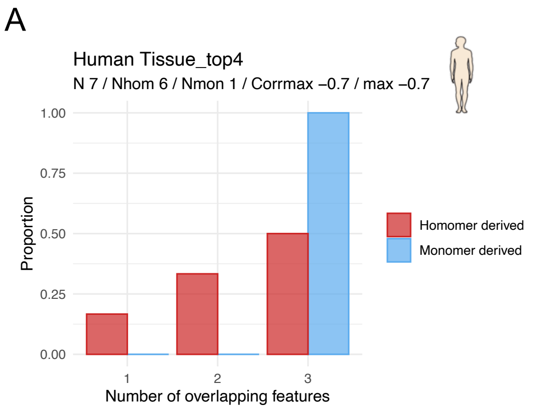

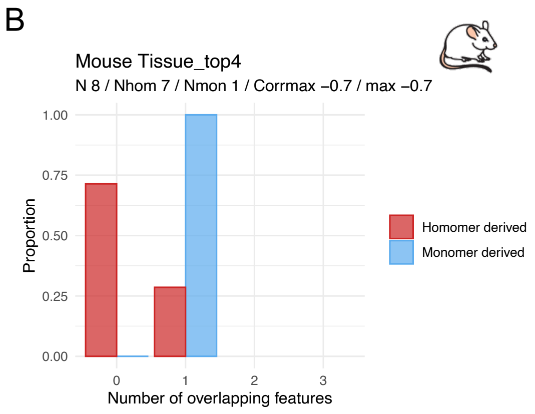

### Supplementary Figure 7

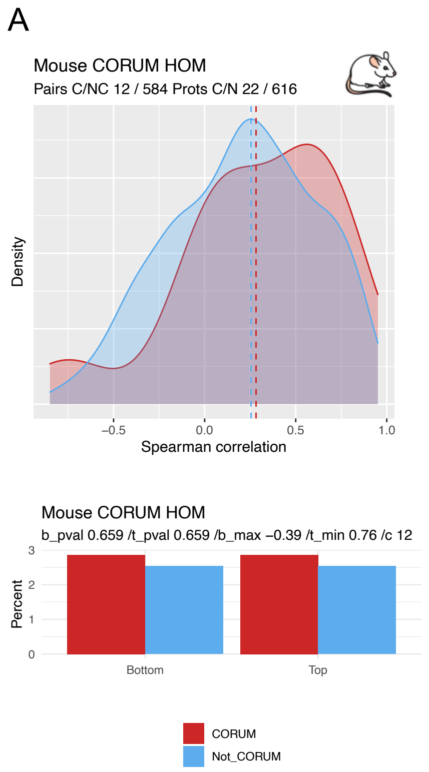

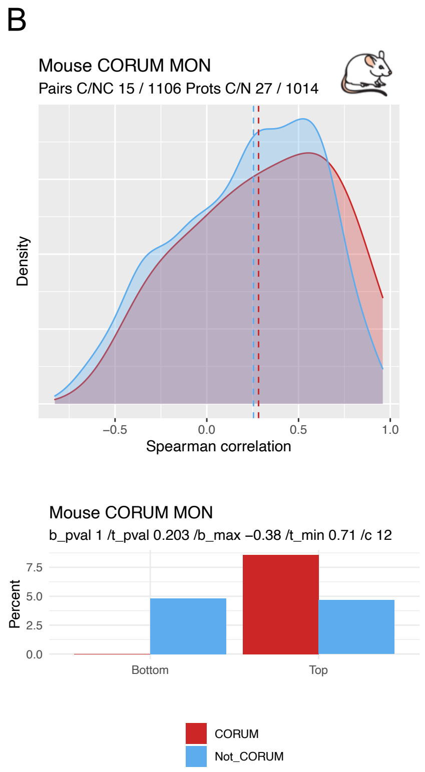

### Supplementary Figure 8

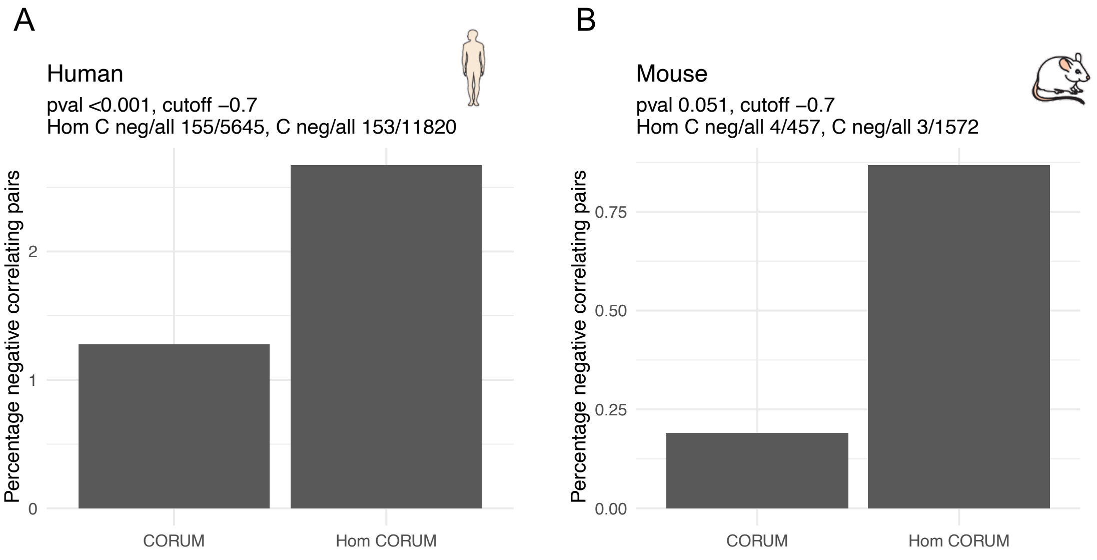

Supplementary Figure 9

Paralogue pairs correlating >0.5 in both Human and Mouse

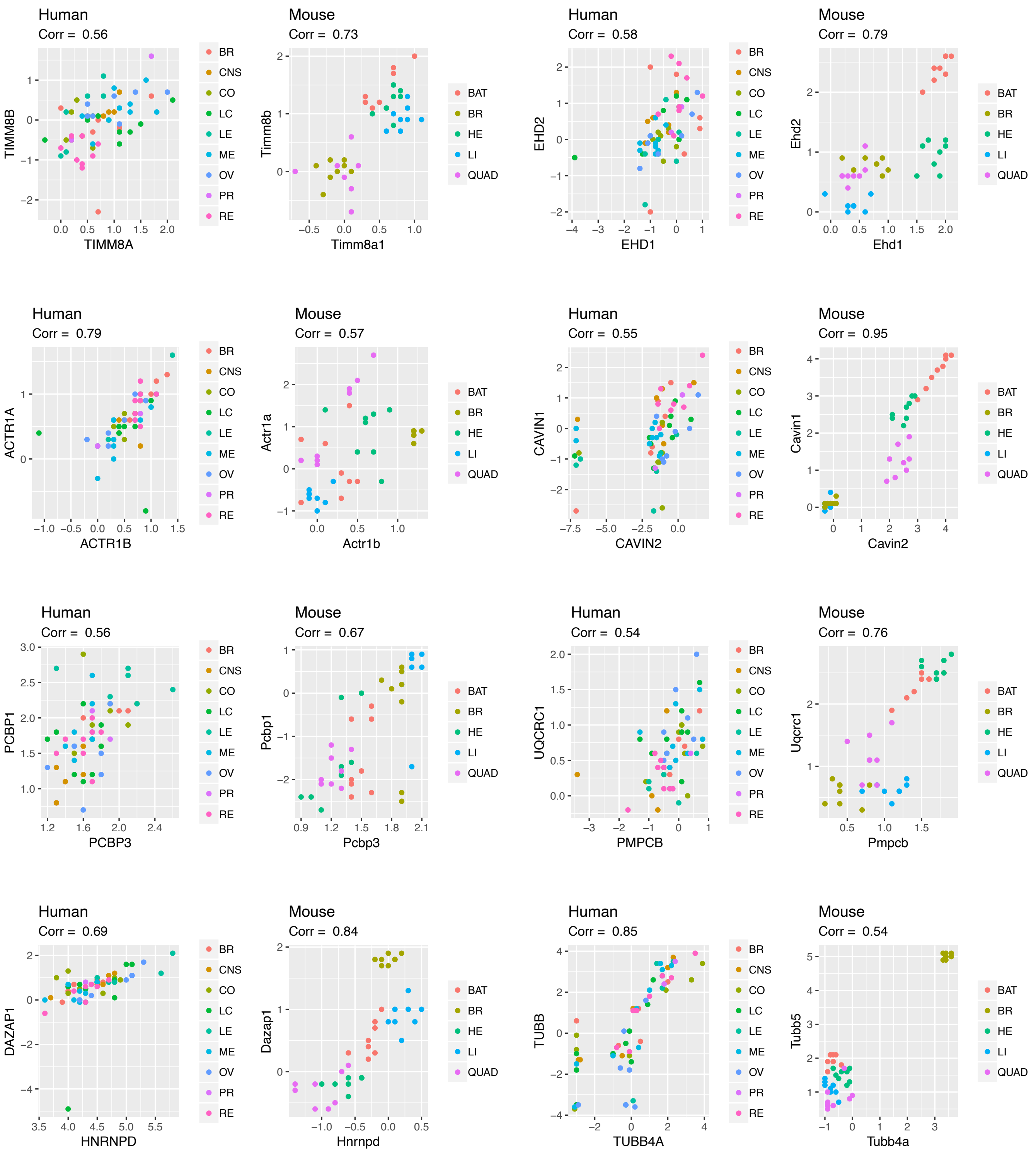

Supplementary Figure 10

Paralouge pairs correlating <(-0.2) in both Human and Mouse

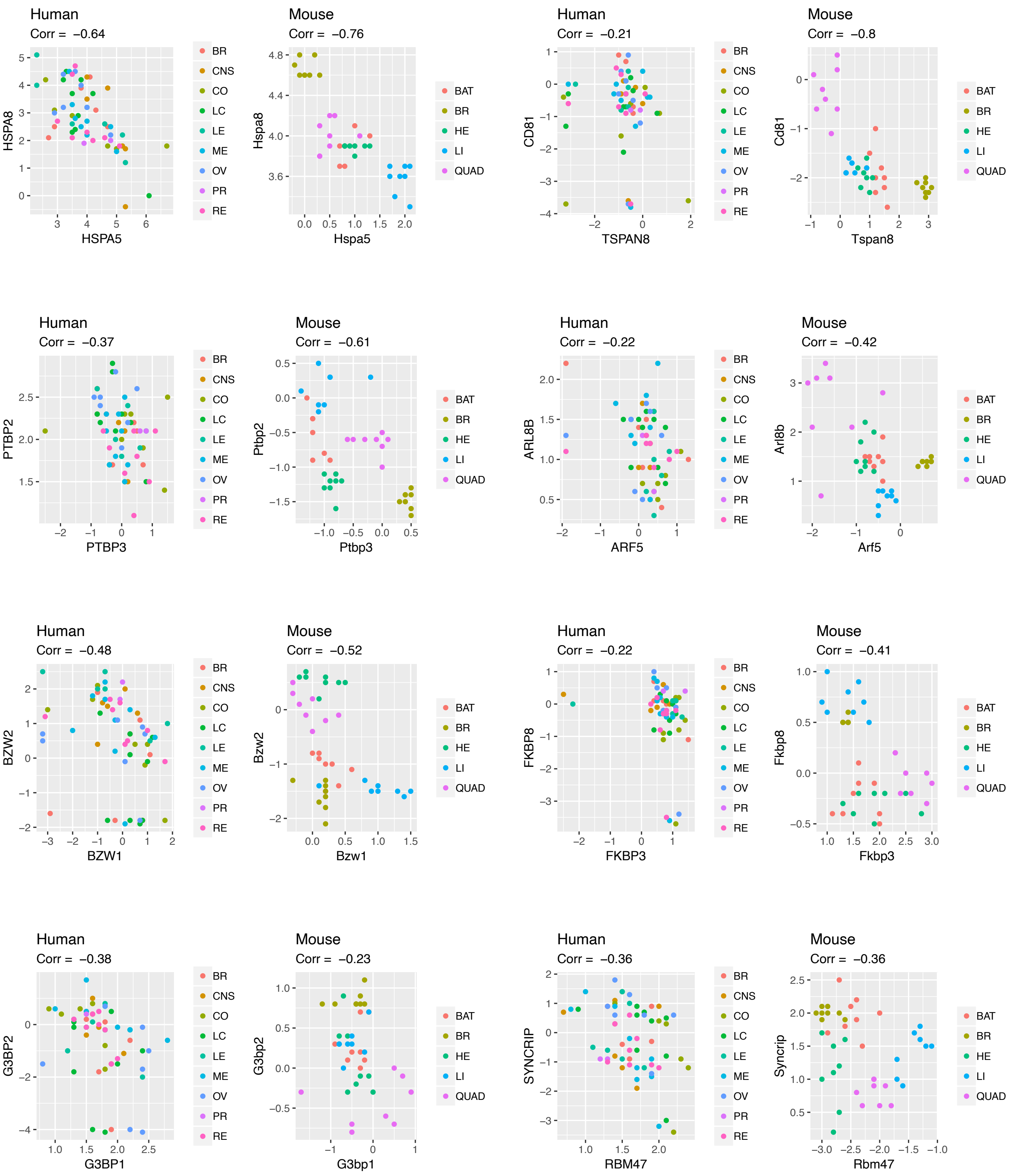

**Figure S1: Overlap of the datasets.**

(**A**) Theoretically possible overlap based on the genomic data for human orthologues in mouse, fly and yeast as reported in Ensembl. 84.5% of the human genes have at least one reported orthologue in mouse, 18.6% have at least one reported orthologue in mouse, fly and yeast.

(**B**) Human orthologues measured in the human, mouse, fly and yeast SWATH-MS datasets. From 3007 measured human proteins, 35.2% have a measured orthologue in mouse, 5.3% have a measured orthologue in all four species.

**Figure S2: Protein complexes show high SWATH-MS co-expression profiles.**

Density plot of the Spearman correlation of the expression profiles of protein pairs that are annotated as members of protein complexes (red) and for protein pairs that are not annotated as members of protein complexes (blue). Expression profiles of protein pairs part of protein complexes correlate significantly more than non-complex pairs in all species (Wilcoxon signed-rank test p-values < 0.001, number of complex pairs human = 17453, mouse = 1739, fly = 2261, yeast = 1807). The definition of protein complexes is based on CORUM (human and mouse), DroID (fly) and CYC2008 (yeast). Dashed lines indicate sample medians.

**Figure S3: SWATH-MS co-expression profiles of protein complex modules are conserved across species.**

For each species combination we tested

(**A**) whether protein pairs that are co-expressed in one species are co-expressed in the other species and

(**B**) whether known complex members described in one species are co-expressed as well in the other species. For complex member prediction we used BIOGRID protein interaction data with physical evidence in the respective species.

The number of observations and the Wilcoxon signed-rank test p-values are indicated above the density plots. Dashed lines indicate sample medians. Note that * marked density plots have less than 40 observations.

**Figure S4: Paralogue family analysis.**

(**A**) A paralogue family is defined as all genes emerged from the same ancestral gene by duplication.

(**B**) Hypothetical pairwise correlation values from the paralogue family depicted in (A).

**Figure S5: Negative correlations are enriched in homomer derived paralogues compared to monomer derived paralogues.**

Percentage of negative correlating pairs (correlation < -0.7) from homomer and monomer derived paralogues in human (**A**) and mouse (**B**) (number of human homomeric pairs = 540, human monomeric pairs = 620, mouse homomeric pairs = 615, mouse monomeric pairs = 1121, Fisher's exact test p-values = 0.04 and 0.2, respectively).

**Figure S6: Negative correlating paralogues show more divergent tissue expression.**

The overlap of the four most expressed tissues between paralogous pairs (as defined by HPM) is compared between strongly negative correlating homomer-derived and monomer-derived pairs paralogous pairs in human (**A**) and mouse (**B**) ($r_{s\,<}$ -0.7, human $n_{pairs\,=}$ 7, mouse $n_{pairs}$ = 8). Negatively correlating homomer-derived paralogues tend to be expressed more diversely across different tissues compared to monomer-derived paralogues.

**Figure S7: Among negatively correlating paralogues in mouse, homomer-derived pairs are enriched in complex members compared to non-complex members.**

(**A**) Spearman correlation of homomer-derived paralogues that are annotated as CORUM complex pairs ($n_{pairs}$ = 12) and for pairs that are not annotated in CORUM ($n_{pairs}$ = 584).

**(B)** Spearman correlation of monomer-derived paralogues that are annotated as CORUM complex pairs ($n_{pairs}$ = 15) and for pairs that are not annotated in CORUM ($n_{pairs}$ = 1106).

**Figure S8: Negative correlating CORUM pairs are enriched in homomer derived paralogues.**

Percentage of negative correlating pairs (correlation < -0.7) from homomer derived and non-homomer derived CORUM pairs in human (**A**) and mouse (**B**) (number of human homomeric CORUM pairs = 5645, human CORUM pairs = 11820, mouse homomeric CORUM pairs = 457, mouse CORUM pairs = 1572, Fisher's exact test p-values < 0.001 and 0.05, respectively).

**Figure S9: Positively correlating paralogous pairs in human and mouse.**

Scatter plots for all paralogous pairs that correlate > 0.5 in both human and mouse. Each dot indicates a sample according to its expression in the two paralogous proteins indicated on the axis. Above each plot the Spearman correlation is indicated. Samples are colored according to their tissue of origin. Human tissues are breast (BR), central nervous system (CNS), colon (CO), lung (LC), blood (LE), skin (ME), ovary (OV), prostate (PR) and kidney (RE); mouse tissues are brown adipose (BAT), brain (BR), heart (HE), liver (LI), quadriceps (QUAD).

**Figure S10: Negatively correlating paralogous pairs in human and mouse.**

Scatter plots for all paralogous pairs that correlate < (-0.2) in both human and mouse. Each dot indicates a sample according to its expression in the two paralogous proteins indicated on the axis. Above each plot the Spearman correlation is indicated. Samples are colored according to their tissue of origin. Human tissues are breast (BR), central nervous system (CNS), colon (CO), lung (LC), blood (LE), skin (ME), ovary (OV), prostate (PR) and kidney (RE); mouse tissues are brown adipose (BAT), brain (BR), heart (HE), liver (LI), quadriceps (QUAD).

**Table S1: Overview of used datasets.**

For each species dataset, the dataset source, sample number, sample source, number of biological and technical replicates as well as the used mass spectrometer instrument are indicated.

**Table S2: DAVID functional annotation of highly correlated pairs conserved in human and mouse.**

(**A**) Proteins correlated $\rho > 0.8$ in both human and mouse 2018 are compared to all proteins measured in both human and mouse. "Term" indicates the associated GO biological process, "Count" shows the total number of genes associated with the term, "P-value" indicates the EASE score (modified fisher exact test p-value) for the respective term and "Genes" shows the human uniprot identifier of the associated genes.

(**B**) Proteins correlated $\rho > 0.6$ in both human and mouse 2018 are compared to all proteins measured in both human and mouse. "Term" indicates the associated GO biological process, "Count" shows the total number of genes associated with the term, "P-value" indicates the EASE score (modified fisher exact test p-value) for the respective term and "Genes" shows the human uniprot identifier of the associated genes.

**Table S3: Number of paralogue families in the different species.**

Number of paralogue families with the respective mean and median sizes.

**Table S4: Paralogue correlation in human and mouse.**

Specifications of paralogous pairs. Paralogous pairs are denoted with Uniprot identifiers (Prot_1 and Prot_2). Corr indicates the respective Spearman correlation. Prot_1_Hom and Prot_2_Hom indicates whether the respective protein is annotated as homomer or heteromer in the InterEvol database. Hom and Mon indicates whether we defined this pair as homomer-, respectively monomer-derived paralogous pair. CORUM indicates whether the pair is annotated in the CORUM database, where cluster_id and CORUM_organism and CORUM_name indicate the complex specifications indicated in CORUM. Biogrid indicates whether the pair is annotated in the Biogrid database.

Table S1

| | Human | Mouse | Fly | Yeast |
|---|---|---|---|---|
| **Source** | Guo, T. et al. (2019). Quantitative Proteome Landscape of the NCI-60 Cancer Cell Lines. iScience 21, 664–680. | Williams, E. G. et al. (2018). Quantifying and Localizing the Mitochondrial Proteome Across Five Tissues in A Mouse Population. Mol Cell Proteomics, 17(9), 1766-1777. | Okada, H. et al. (2016). Proteome-wide association studies identify biochemical modules associated with a wing-size phenotype in Drosophila melanogaster. Nat Commun, 7, 12649. | Proteomics data unpublished, Yeast strain references: Brem, R. B. et al. (2002). Genetic dissection of transcriptional regulation in budding yeast. Science, 296(5568), 752-755.; Zhu, J. et al. (2008). Integrating large-scale functional genomic data to dissect the complexity of yeast regulatory networks. Nat Genet, 40(7), 854-861. |
| **Sample number** | 60 | 40 | 60 | 112 |
| **Sample source** | NCI-60 cancer cell line panel (leukemia, melanoma, non-small-cell lung carcinoma, and cancers of the brain, ovary, breast, colon, kidney and prostate) | Mouse tissue samples (liver, quadriceps, heart, brain and brown adipose) | Drosophila genetic reference panel (DGRP) lines, wing tissue | 170 yeast strains from cross of haploid derivatives of a standard laboratory strain (BY) and a wild isolate from a California vineyard (RM) |
| **Biological replicates** | 5 biological replicates per cell line | 8 different mouse BXD lines per tissue | 30 bit wing size and 30 small wing size replicates | 1-8 replicates per strain |
| **Technical replicates** | 2 technical replicates | None | 3 | None |
| **Instrument** | SCIEX TripleTOF 5600 | SCIEX TripleTOF 5600 | SCIEX TripleTOF 5600 | SCIEX TripleTOF 5600 |
| **Analysis** | OpenSWATH | OpenSWATH | OpenSWATH | OpenSWATH |

**Table S1 DAVID functional annotation of highly correlated pairs conserved in human and mouse.**

(a) Proteins correlated ρ>0.8 in both human and mouse 2018 are compared to all proteins measured in both human and mouse. "Term" indicates the associated GO biological process, "Count" shows the total number of genes associated with the term, "P-value" indicates the EASE score (modified fisher exact test p-value) for the respective term and "Genes" shows the human uniprot identifier of the associated genes.

| **Annotation Cluster 1: Enrichment Score: 3.6** | | | |
|---|---|---|---|
| **Term** | **Count** | **PValue** | **Genes** |
| GO:0042776~mitochondrial ATP synthesis coupled proton transport | 6 | 0.000 | P06576, P08574, P25705, P56385, O75947, P36542 |
| GO:0006754~ATP biosynthetic process | 5 | 0.000 | P06576, P25705, P56385, O75947, P36542 |
| GO:0015986~ATP synthesis coupled proton transport | 4 | 0.002 | P06576, P25705, O75947, P36542 |
| **Annotation Cluster 2: Enrichment Score: 1.4** | | | |
| **Term** | **Count** | **PValue** | **Genes** |
| GO:0006120~mitochondrial electron transport, NADH to ubiquinone | 4 | 0.035 | O14561, O95299, P49821, O43678 |
| GO:0032981~mitochondrial respiratory chain complex I assembly | 4 | 0.044 | O14561, O95299, P49821, O43678 |

(b) Proteins correlated ρ>0.6 in both human and mouse 2018 are compared to all proteins measured in both human and mouse. "Term" indicates the associated GO biological process, "Count" shows the total number of genes associated with the term, "P-value" indicates the EASE score (modified fisher exact test p-value) for the respective term and "Genes" shows the human uniprot identifier of the associated genes.

| **Annotation Cluster 1: Enrichment Score: 4.4** |
|---|

| Term | Count | PValue | Genes |
|---|---|---|---|
| GO:0006120~mitochondrial electron transport, NADH to ubiquinone | 17 | 0.000 | O14561, O43676, P49821, O75489, Q9P0J0, O96000, P51970, P56556, O43678, P09622, P17568, O95168, O95299, P28331, Q16718, Q9Y6M9, O75380 |
| GO:0032981~mitochondrial respiratory chain complex I assembly | 18 | 0.000 | O14561, O43676, P49821, O75489, Q9P0J0, O96000, P51970, P56556, O43678, Q9H845, O95831, P17568, O95168, O95299, P28331, Q16718, Q9Y6M9, O75380 |
| **Annotation Cluster 2: Enrichment Score: 2.7** | | | |
| **Term** | **Count** | **PValue** | **Genes** |
| GO:0000398~mRNA splicing, via spliceosome | 22 | 0.000 | Q13151, O75643, Q13247, P14866, Q9UMS4, Q8WYA6, Q9UKF6, P31943, Q13838, Q15717, P11142, P55795, P07910, Q9BXP5, P35637, O43143, Q14103, Q00839, P61978, O43809, Q15365, O60506 |
| GO:0010467~gene expression | 9 | 0.014 | Q13151, P55795, P07910, P14866, Q14103, P31943, P61978, Q00839, Q15365 |
| **Annotation Cluster 3: Enrichment Score: 2.6** | | | |
| **Term** | **Count** | **PValue** | **Genes** |
| GO:0042776~mitochondrial ATP synthesis coupled proton transport | 10 | 0.001 | P24539, P06576, P08574, P56134, P30049, P18859, P25705, P56385, O75947, P36542 |
| GO:0006754~ATP biosynthetic process | 10 | 0.001 | P24539, P06576, P56134, P30049, P18859, P25705, P56385, P14618, O75947, P36542 |

| | | | |
|---|---|---|---|
| GO:0015986~ATP synthesis coupled proton transport | 7 | 0.016 | P24539, P06576, P30049, P18859, P25705, O75947, P36542 |
| **Annotation Cluster 4: Enrichment Score: 2.3** | | | |
| **Term** | **Count** | **PValue** | **Genes** |
| GO:0070125~mitochondrial translational elongation | 18 | 0.004 | Q9NYK5, Q9H2W6, P82914, Q96RP9, Q9Y2R9, Q9HD33, P51398, Q92552, Q8N5N7, Q96DV4, P52815, Q7Z7H8, Q6P1L8, P82933, Q9NRX2, P49406, Q8IXM3, Q13405 |
| GO:0070126~mitochondrial translational termination | 17 | 0.005 | Q9NYK5, Q9H2W6, P82914, Q9Y2R9, Q9HD33, P51398, Q92552, Q8N5N7, Q96DV4, P52815, Q7Z7H8, Q6P1L8, P82933, Q9NRX2, P49406, Q8IXM3, Q13405 |
| **Annotation Cluster 5: Enrichment Score: 2.2** | | | |
| **Term** | **Count** | **PValue** | **Genes** |
| GO:0006413~translational initiation | 21 | 0.002 | P60228, P46776, P05387, P49207, P39019, Q92905, P62910, Q14152, P55884, Q9Y262, P60842, Q15056, Q13347, O60841, O15371, P62266, P62829, P05198, P50914, O00303, P63220 |
| GO:0000184~nuclear-transcribed mRNA catabolic process, nonsense-mediated decay | 14 | 0.002 | P60228, Q9BRP8, P46776, P05387, P49207, P39019, P62495, P30153, P62910, P62266, P62829, P50914, P63220, Q92900 |
| GO:0006412~translation | 23 | 0.003 | P82914, P46776, P05387, Q9Y2R9, P49207, P39019, Q92905, P26639, P62910, P05141, Q9NSD9, P62266, P62829, Q02978, Q6P1L8, P50914, |

| | | | |
|---|---|---|---|
| | | | Q7Z7H8, P82933, Q9NRX2, P63220, P49406, Q13405, Q8IXM3 |
| GO:0006364~rRNA processing | 11 | 0.011 | P46776, P05387, P49207, P39019, Q9Y2W2, P62266, P62829, P62910, P50914, P63220, P22087 |
| GO:0019083~viral transcription | 9 | 0.014 | P46776, P05387, P49207, P39019, P62266, P62829, P62910, P50914, P63220 |
| GO:0006446~regulation of translational initiation | 10 | 0.038 | P60228, Q15056, Q13347, O60841, O15371, Q14152, O00303, P55884, Q9Y262, P60842 |
| **Annotation Cluster 6: Enrichment Score: 1.6** | | | |
| **Term** | **Count** | **PValue** | **Genes** |
| GO:0033209~tumor necrosis factor-mediated signaling pathway | 12 | 0.010 | P28074, O43242, P28072, P25789, Q9BRA2, P28070, P17980, P05787, P55036, P60900, P49721, O00487 |
| GO:0038061~NIK/NF-kappaB signaling | 10 | 0.023 | P28074, O43242, P28072, P25789, P28070, P17980, P55036, P60900, P49721, O00487 |
| GO:0006521~regulation of cellular amino acid metabolic process | 10 | 0.023 | P28074, O43242, P28072, P25789, P28070, P17980, P55036, P60900, P49721, O00487 |
| GO:0090090~negative regulation of canonical Wnt signaling pathway | 12 | 0.027 | P28074, O43242, P28072, P25789, P28070, Q13283, P17980, P55036, P60900, P49721, Q03135, O00487 |
| GO:0090263~positive regulation of canonical Wnt signaling pathway | 12 | 0.027 | P28074, O43242, P28072, P25789, P28070, P17980, P55036, P14923, P60900, P49721, Q03135, O00487 |
| GO:0002223~stimulatory C-type lectin receptor signaling pathway | 11 | 0.032 | P28074, O43242, P28072, P25789, P28070, P17980, P55036, P17612, P60900, P49721, O00487 |

| | | | |
|---|---|---|---|
| GO:0038095~Fc-epsilon receptor signaling pathway | 11 | 0.032 | P28074, O43242, P28072, P25789, P62993, P28070, P17980, P55036, P60900, P49721, O00487 |
| GO:0031145~anaphase-promoting complex-dependent catabolic process | 10 | 0.038 | P28074, O43242, P28072, P25789, P28070, P17980, P55036, P60900, P49721, O00487 |
| GO:0051436~negative regulation of ubiquitin-protein ligase activity involved in mitotic cell cycle | 10 | 0.038 | P28074, O43242, P28072, P25789, P28070, P17980, P55036, P60900, P49721, O00487 |
| GO:0051437~positive regulation of ubiquitin-protein ligase activity involved in regulation of mitotic cell cycle transition | 10 | 0.038 | P28074, O43242, P28072, P25789, P28070, P17980, P55036, P60900, P49721, O00487 |

**Table S3 Number of paralogue families in the different species.**

| | Human | Mouse | Fly | Yeast |
|---|---|---|---|---|
| Number of paralogue families with >2 measured paralogues | 73 | 114 | 3 | 9 |
| Mean paralogue family size | 9 | 9 | 11 | 9 |
| Median paralogue family size | 8 | 9 | 7 | 6 |
| Number of paralogue family with available interaction Jaccard-index and >2 measured paralogues | 73 | 92 | 0 | 7 |
| Mean paralogue family size | 9 | 9 | NA | 7 |
| Median paralogue family size | 8 | 9 | NA | 5 |

Table S4 Human

| Prot_1 | Prot_2 | Corr | Prot_1_Hom | Prot_2_Hom | Hom | Mon | CORUM | cluster_id | CORUM_organism | CORUM_name | Biogrid |
|---|---|---|---|---|---|---|---|---|---|---|---|
| Q07955 | Q13247 | -0.81 | HOM | HOM | TRUE | FALSE | TRUE | 351 | Human | Spliceosome | TRUE |
| P08238 | P14625 | -0.766 | HOM | HOM | TRUE | FALSE | FALSE | | | | FALSE |
| P27348 | P62258 | -0.765 | | HOM | TRUE | FALSE | TRUE | 5199,5615 | Human | Kinase maturation complex 1,Emerin complex 52 | TRUE |
| Q00839 | Q9BUJ2 | -0.755 | | | FALSE | TRUE | TRUE | 1332 | Human | Large Drosha complex | FALSE |
| P09651 | P51991 | -0.752 | HOM | | TRUE | FALSE | TRUE | 1181 | Human | C complex spliceosome | TRUE |
| P07900 | P08238 | -0.735 | HOM | HOM | TRUE | FALSE | TRUE | 5199,5212,5234,5266,5268,5269,5286,25 | Human,Mouse | Kinase maturation complex 1,Kinase maturation complex 2,IKBKB-CDC37-KIAA1967-HSP90AB1-HSP90AA1 complex,TNF-alpha/NF-kappa B signaling complex 6,TNF-alpha/NF-kappa B signaling complex 7,TNF-alpha/NF-kappa B signaling complex 8,TNF-alpha/NF-kappa B signaling complex 10,9S-cytosolic aryl hydrocarbon receptor non-ligand activated complex | TRUE |
| P49792 | P62937 | -0.709 | HET | HOM | TRUE | FALSE | FALSE | | | | FALSE |
| Q07955 | Q13242 | -0.7 | HOM | HOM | TRUE | FALSE | TRUE | 351 | Human | Spliceosome | TRUE |
| P27348 | P63104 | -0.654 | | | FALSE | TRUE | TRUE | 5199 | Human | Kinase maturation complex 1 | TRUE |
| O95758 | P26599 | -0.644 | | | FALSE | TRUE | FALSE | | | | FALSE |
| P11021 | P11142 | -0.64 | HOM | HOM | TRUE | FALSE | TRUE | 929,2721 | Human | CEN complex,HCF-1 complex | TRUE |
| P28062 | P28074 | -0.635 | HOM | HOM | TRUE | FALSE | FALSE | | | | TRUE |
| Q8IWZ3 | Q8TC84 | -0.622 | | | FALSE | TRUE | FALSE | | | | FALSE |
| O95758 | P14866 | -0.606 | | | FALSE | TRUE | FALSE | | | | FALSE |

Table S4 Human

| | | | | | | | | | | | |
|---|---|---|---|---|---|---|---|---|---|---|---|
| P02545 | P20700 | -0.596 | HOM | | TRUE | FALSE | TRUE | 5608,5611 | Human | Emerin architectural complex,Emerin complex 24 | TRUE |
| P11021 | P34931 | -0.593 | HOM | | TRUE | FALSE | FALSE | | | | TRUE |
| Q04323 | Q92575 | -0.57 | | | FALSE | TRUE | FALSE | | | | FALSE |
| Q1KMD3 | Q9BUJ2 | -0.544 | | | FALSE | TRUE | FALSE | | | | TRUE |
| P08670 | P20700 | -0.523 | HOM | | TRUE | FALSE | FALSE | | | | TRUE |
| P62820 | Q15286 | -0.514 | HET | HET | FALSE | FALSE | FALSE | | | | FALSE |
| Q01844 | Q92804 | -0.512 | | | FALSE | TRUE | TRUE | 1332 | Human | Large Drosha complex | FALSE |
| Q7L1Q6 | Q9Y6E2 | -0.484 | | | FALSE | TRUE | FALSE | | | | TRUE |
| O43447 | Q9H2H8 | -0.48 | HET | | FALSE | FALSE | TRUE | 351 | Human | Spliceosome | FALSE |
| P28065 | P28072 | -0.473 | HOM | HOM | TRUE | FALSE | FALSE | | | | TRUE |
| Q9BXS5 | Q9Y6Q5 | -0.46 | HET | HET | FALSE | FALSE | TRUE | 36,144 | Human | AP1 adaptor complex,Gamma-BAR-AP1 complex | FALSE |
| P35637 | Q92804 | -0.459 | | | FALSE | TRUE | TRUE | 1332 | Human | Large Drosha complex | TRUE |
| P60953 | P84095 | -0.458 | HOM | | TRUE | FALSE | FALSE | | | | FALSE |
| P68371 | Q13885 | -0.446 | HET | HET | FALSE | FALSE | FALSE | | | | TRUE |
| Q07955 | Q13243 | -0.445 | HOM | HOM | TRUE | FALSE | TRUE | 351 | Human | Spliceosome | TRUE |
| P23284 | Q9Y3C6 | -0.439 | | | FALSE | TRUE | FALSE | | | | FALSE |
| P23284 | Q9H2H8 | -0.437 | | | FALSE | TRUE | FALSE | | | | FALSE |
| P09651 | Q13151 | -0.433 | HOM | | TRUE | FALSE | FALSE | | | | TRUE |
| P55084 | Q9BWD1 | -0.428 | HOM | HOM | TRUE | FALSE | FALSE | | | | FALSE |
| P62244 | Q15041 | -0.41 | | | FALSE | TRUE | FALSE | | | | FALSE |
| P13796 | P13797 | -0.409 | | | FALSE | TRUE | FALSE | | | | TRUE |
| P53675 | Q00610 | -0.398 | | | FALSE | TRUE | FALSE | | | | TRUE |
| P51858 | Q7Z4V5 | -0.393 | HOM | | TRUE | FALSE | FALSE | | | | FALSE |
| Q13283 | Q9UN86 | -0.383 | HOM | | TRUE | FALSE | FALSE | | | | TRUE |
| P35268 | Q6P5R6 | -0.378 | | | FALSE | TRUE | FALSE | | | | FALSE |

Table S4 Human

| | | | | | | | | | | | |
|---|---|---|---|---|---|---|---|---|---|---|---|
| O95758 | Q9UKA9 | -0.368 | | | FALSE | TRUE | FALSE | | | | FALSE |
| P04271 | P31949 | -0.366 | HOM | | TRUE | FALSE | FALSE | | | | TRUE |
| P04350 | Q13885 | -0.363 | HET | HET | FALSE | FALSE | FALSE | | | | FALSE |
| A0AV96 | O60506 | -0.357 | | | FALSE | TRUE | FALSE | | | | FALSE |
| B0I1T2 | O00159 | -0.356 | HOM | HOM | TRUE | FALSE | FALSE | | | | FALSE |
| P61978 | Q15366 | -0.355 | HOM | HOM | TRUE | FALSE | FALSE | | | | TRUE |
| O60610 | O95466 | -0.355 | HOM | | TRUE | FALSE | FALSE | | | | FALSE |
| P18085 | P84077 | -0.348 | | HOM | TRUE | FALSE | FALSE | | | | TRUE |
| Q15363 | Q9Y3B3 | -0.346 | HOM | HOM | TRUE | FALSE | FALSE | | | | TRUE |
| Q92499 | Q9BUJ2 | -0.345 | | | FALSE | TRUE | TRUE | 1332 | Human | Large Drosha complex | FALSE |
| Q14257 | Q9BRK5 | -0.342 | | | FALSE | TRUE | FALSE | | | | FALSE |
| P09936 | P15374 | -0.34 | HET | HET | FALSE | FALSE | FALSE | | | | FALSE |
| Q02790 | Q14318 | -0.338 | HOM | HET | TRUE | FALSE | FALSE | | | | FALSE |
| P53396 | P53597 | -0.328 | | | FALSE | TRUE | FALSE | | | | FALSE |
| P13639 | Q15029 | -0.328 | | | FALSE | TRUE | FALSE | | | | FALSE |
| O00629 | P52292 | -0.325 | HOM | HET,HOM | TRUE | FALSE | FALSE | | | | TRUE |
| O95319 | Q92879 | -0.322 | | HOM | TRUE | FALSE | FALSE | | | | FALSE |
| Q96CS3 | Q9UNN5 | -0.322 | | HET | FALSE | FALSE | FALSE | | | | FALSE |
| O60884 | Q9UBS4 | -0.321 | HOM | HOM | TRUE | FALSE | FALSE | | | | FALSE |
| Q15293 | Q9BRK5 | -0.321 | | | FALSE | TRUE | FALSE | | | | FALSE |
| Q99714 | Q9BUT1 | -0.314 | HOM | HOM | TRUE | FALSE | FALSE | | | | FALSE |
| Q03252 | Q16352 | -0.31 | | | FALSE | TRUE | FALSE | | | | FALSE |
| O75525 | Q07666 | -0.31 | HOM | HOM | TRUE | FALSE | FALSE | | | | TRUE |
| P09496 | P09497 | -0.31 | | | FALSE | TRUE | FALSE | | | | TRUE |
| Q969X5 | Q96RQ1 | -0.309 | | | FALSE | TRUE | FALSE | | | | FALSE |
| Q15393 | Q16531 | -0.309 | | HOM | TRUE | FALSE | FALSE | | | | FALSE |

Table S4 Human

| | | | | | | | | | | | |
|---|---|---|---|---|---|---|---|---|---|---|---|
| P50151 | P63218 | -0.308 | | | FALSE | TRUE | FALSE | | | | FALSE |
| P33176 | Q02241 | -0.306 | HOM | HET | TRUE | FALSE | FALSE | | | | FALSE |
| P20700 | Q16352 | -0.306 | | | FALSE | TRUE | FALSE | | | | FALSE |
| P62820 | Q9H0U4 | -0.303 | HET | HET | FALSE | FALSE | FALSE | | | | TRUE |
| Q5M775 | Q8TDZ2 | -0.3 | | | FALSE | TRUE | FALSE | | | | FALSE |
| P60842 | Q14240 | -0.298 | HET | | FALSE | FALSE | FALSE | | | | TRUE |
| P61981 | P62258 | -0.296 | | HOM | TRUE | FALSE | TRUE | 5199 | Human | Kinase maturation complex 1 | TRUE |
| P55795 | Q9NTZ6 | -0.295 | | HOM | TRUE | FALSE | FALSE | | | | FALSE |
| Q92598 | Q9Y4L1 | -0.292 | HOM | HOM | TRUE | FALSE | FALSE | | | | FALSE |
| Q06265 | Q96B26 | -0.29 | HOM, HET | HOM, HET | TRUE, FALSE | FALSE | TRUE | 789 | Human | Exosome | TRUE |
| Q9BTT6 | Q9H9A6 | -0.288 | | | FALSE | TRUE | FALSE | | | | FALSE |
| P30153 | P30154 | -0.287 | HET | HET | FALSE | FALSE | TRUE | 5876 | Human | PPP2R1A-PPP2R1B-PPP2CA-PPME1-EIF4A1 complex | TRUE |
| Q7Z4W1 | Q9BUT1 | -0.286 | HOM | HOM | TRUE | FALSE | FALSE | | | | FALSE |
| O15020 | P12814 | -0.286 | | | FALSE | TRUE | FALSE | | | | FALSE |
| P68371 | Q9BVA1 | -0.282 | HET | HET | FALSE | FALSE | FALSE | | | | FALSE |
| Q15555 | Q15691 | -0.278 | | HOM | TRUE | FALSE | FALSE | | | | TRUE |
| P08133 | P12429 | -0.275 | | | FALSE | TRUE | FALSE | | | | FALSE |
| P31943 | Q6NXG1 | -0.274 | | | FALSE | TRUE | FALSE | | | | FALSE |
| P23297 | P25815 | -0.273 | HET | HOM | TRUE | FALSE | FALSE | | | | TRUE |
| Q01970 | Q15147 | -0.265 | | | FALSE | TRUE | FALSE | | | | FALSE |
| P15311 | P26038 | -0.264 | | | FALSE | TRUE | FALSE | | | | TRUE |
| P20337 | P61006 | -0.264 | HOM | HET,HOM | TRUE | FALSE | FALSE | | | | TRUE |
| Q92817 | Q9UPN3 | -0.263 | HOM | | TRUE | FALSE | FALSE | | | | FALSE |
| P51159 | P61006 | -0.261 | HET,HOM | HET,HOM | FALSE,TRUE | FALSE | FALSE | | | | FALSE |

Table S4 Human

| | | | | | | | | | | | |
|---|---|---|---|---|---|---|---|---|---|---|---|
| Q96MG7 | Q9UNF1 | -0.26 | HET | | FALSE | FALSE | FALSE | | | | FALSE |
| Q9BWM7 | Q9H9B4 | -0.259 | | | FALSE | TRUE | FALSE | | | | TRUE |
| O95071 | Q7Z6Z7 | -0.258 | | | FALSE | TRUE | FALSE | | | | FALSE |
| P34931 | Q0VDF9 | -0.257 | | HET | FALSE | FALSE | FALSE | | | | FALSE |
| Q01085 | Q14498 | -0.257 | | | FALSE | TRUE | FALSE | | | | FALSE |
| P45877 | Q6UX04 | -0.255 | | | FALSE | TRUE | FALSE | | | | FALSE |
| P45877 | Q9Y3C6 | -0.254 | | | FALSE | TRUE | FALSE | | | | FALSE |
| P07196 | Q16352 | -0.252 | | | FALSE | TRUE | FALSE | | | | FALSE |
| P43121 | P50895 | -0.25 | | | FALSE | TRUE | FALSE | | | | FALSE |
| Q07955 | Q08170 | -0.249 | HOM | HOM | TRUE | FALSE | TRUE | 351 | Human | Spliceosome | TRUE |
| O15143 | Q92747 | -0.247 | | | FALSE | TRUE | FALSE | | | | FALSE |
| Q14203 | Q99426 | -0.246 | HOM | HET | TRUE | FALSE | FALSE | | | | FALSE |
| P43034 | P61964 | -0.245 | HOM | | TRUE | FALSE | FALSE | | | | TRUE |
| P23297 | P26447 | -0.245 | HET | | FALSE | FALSE | FALSE | | | | TRUE |
| P11137 | P27816 | -0.244 | | | FALSE | TRUE | FALSE | | | | FALSE |
| P08134 | P62745 | -0.243 | | | FALSE | TRUE | FALSE | | | | FALSE |
| P07437 | Q13885 | -0.24 | HET | HET | FALSE | FALSE | FALSE | | | | TRUE |
| Q96AB3 | Q96CN7 | -0.235 | | | FALSE | TRUE | FALSE | | | | FALSE |
| P63218 | Q9UBI6 | -0.234 | | | FALSE | TRUE | FALSE | | | | FALSE |
| O00159 | Q12965 | -0.233 | HOM | HOM | TRUE | FALSE | FALSE | | | | FALSE |
| P15311 | Q9Y4F1 | -0.232 | | | FALSE | TRUE | FALSE | | | | FALSE |
| O14976 | Q68CZ2 | -0.231 | | | FALSE | TRUE | FALSE | | | | FALSE |
| Q14677 | Q9Y6I3 | -0.231 | HET | | FALSE | FALSE | FALSE | | | | FALSE |
| P11142 | Q0VDF9 | -0.231 | HOM | HET | TRUE | FALSE | FALSE | | | | FALSE |
| P84077 | Q9NVJ2 | -0.229 | HOM | | TRUE | FALSE | FALSE | | | | FALSE |
| P17655 | Q9UMQ6 | -0.227 | HOM | | TRUE | FALSE | FALSE | | | | FALSE |

Table S4 Human

| | | | | | | | | | | | |
|---|---|---|---|---|---|---|---|---|---|---|---|
| P31146 | Q9ULV4 | -0.226 | | HOM | TRUE | FALSE | FALSE | | | | TRUE |
| P04271 | P60903 | -0.226 | HOM | | TRUE | FALSE | FALSE | | | | FALSE |
| P05026 | P54709 | -0.225 | | | FALSE | TRUE | FALSE | | | | FALSE |
| P00533 | P43403 | -0.225 | HOM | HOM | TRUE | FALSE | FALSE | | | | TRUE |
| A0AV96 | O43390 | -0.225 | | | FALSE | TRUE | FALSE | | | | FALSE |
| O15479 | P43358 | -0.225 | | | FALSE | TRUE | FALSE | | | | FALSE |
| O43865 | P23526 | -0.223 | | | FALSE | TRUE | FALSE | | | | TRUE |
| P36404 | P84077 | -0.223 | HET | HOM | TRUE | FALSE | FALSE | | | | FALSE |
| Q06830 | Q13162 | -0.222 | HOM | HOM | TRUE | FALSE | FALSE | | | | TRUE |
| O60732 | Q96MG7 | -0.222 | | HET | FALSE | FALSE | FALSE | | | | FALSE |
| Q00688 | Q14318 | -0.22 | | HET | FALSE | FALSE | FALSE | | | | FALSE |
| P84085 | Q9NVJ2 | -0.219 | | | FALSE | TRUE | FALSE | | | | FALSE |
| P12830 | P19022 | -0.219 | HOM | HET,HOM | TRUE | FALSE | FALSE | | | | FALSE |
| O43491 | P11171 | -0.219 | | | FALSE | TRUE | FALSE | | | | FALSE |
| O15382 | P54687 | -0.218 | HOM | HOM | TRUE | FALSE | FALSE | | | | FALSE |
| Q92945 | Q96AE4 | -0.216 | | HET | FALSE | FALSE | FALSE | | | | FALSE |
| P35908 | Q7Z794 | -0.215 | | | FALSE | TRUE | FALSE | | | | TRUE |
| P04271 | P06703 | -0.213 | HOM | HOM | TRUE | FALSE | FALSE | | | | TRUE |
| P61978 | Q15365 | -0.212 | HOM | | TRUE | FALSE | FALSE | | | | TRUE |
| B0I1T2 | O43795 | -0.212 | HOM | HOM | TRUE | FALSE | FALSE | | | | FALSE |
| Q14315 | Q8NF91 | -0.212 | HOM | HOM | TRUE | FALSE | FALSE | | | | FALSE |
| P08579 | P09012 | -0.211 | HET | HOM | TRUE | FALSE | TRUE | 351 | Human | Spliceosome | FALSE |
| O95817 | Q9UL15 | -0.211 | | HET | FALSE | FALSE | FALSE | | | | FALSE |
| P08758 | P50995 | -0.21 | | | FALSE | TRUE | FALSE | | | | FALSE |
| O95425 | P09327 | -0.21 | | HOM | TRUE | FALSE | FALSE | | | | FALSE |
| P19075 | P60033 | -0.206 | | HOM | TRUE | FALSE | FALSE | | | | FALSE |

Table S4 Human

| | | | | | | | | | | | |
|---|---|---|---|---|---|---|---|---|---|---|---|
| P07737 | P35080 | -0.203 | | HOM | TRUE | FALSE | FALSE | | | | FALSE |
| O00291 | Q9Y490 | -0.203 | HOM | HOM | TRUE | FALSE | FALSE | | | | FALSE |
| P10644 | P13861 | -0.202 | | HOM | TRUE | FALSE | FALSE | | | | FALSE |
| Q15436 | Q15437 | -0.201 | HET,HOM | HOM | TRUE | FALSE | FALSE | | | | TRUE |
| P36952 | P50454 | -0.199 | | | FALSE | TRUE | FALSE | | | | FALSE |
| P57723 | P61978 | -0.193 | | HOM | TRUE | FALSE | FALSE | | | | FALSE |
| P07384 | Q9UMQ6 | -0.193 | HET | | FALSE | FALSE | FALSE | | | | FALSE |
| P55795 | Q8IXT5 | -0.193 | | | FALSE | TRUE | FALSE | | | | TRUE |
| P11310 | P49748 | -0.191 | HOM | HOM | TRUE | FALSE | FALSE | | | | FALSE |
| O15020 | Q01082 | -0.189 | | HOM | TRUE | FALSE | FALSE | | | | FALSE |
| P21333 | Q8NF91 | -0.188 | HOM | HOM | TRUE | FALSE | FALSE | | | | FALSE |
| Q08211 | Q7L2E3 | -0.188 | HOM | | TRUE | FALSE | FALSE | | | | FALSE |
| O00505 | O00629 | -0.185 | HOM | HOM | TRUE | FALSE | FALSE | | | | TRUE |
| P62993 | Q13588 | -0.184 | HOM | | TRUE | FALSE | FALSE | | | | FALSE |
| P55011 | Q9UP95 | -0.183 | | | FALSE | TRUE | FALSE | | | | FALSE |
| P32119 | Q13162 | -0.183 | HOM | HOM | TRUE | FALSE | FALSE | | | | TRUE |
| P52888 | Q9BYT8 | -0.181 | | | FALSE | TRUE | FALSE | | | | FALSE |
| P09211 | P21266 | -0.181 | HOM | | TRUE | FALSE | FALSE | | | | FALSE |
| P31689 | Q9UBS4 | -0.18 | HOM | HOM | TRUE | FALSE | FALSE | | | | FALSE |
| P04271 | P26447 | -0.18 | HOM | | TRUE | FALSE | FALSE | | | | TRUE |
| Q92597 | Q9UGV2 | -0.179 | | | FALSE | TRUE | FALSE | | | | FALSE |
| P15924 | Q9UPN3 | -0.178 | HOM | | TRUE | FALSE | FALSE | | | | FALSE |
| O00231 | P61201 | -0.174 | | | FALSE | TRUE | FALSE | | | | FALSE |
| P27348 | P31946 | -0.173 | | | FALSE | TRUE | TRUE | 5199 | Human | Kinase maturation complex 1 | TRUE |
| O00429 | P20591 | -0.168 | | HOM | TRUE | FALSE | FALSE | | | | FALSE |
| P55786 | Q9UIQ6 | -0.168 | | | FALSE | TRUE | FALSE | | | | FALSE |

Table S4 Human

| | | | | | | | | | | | |
|---|---|---|---|---|---|---|---|---|---|---|---|
| Q13330 | Q9BTC8 | -0.168 | | | FALSE | TRUE | TRUE | 587 | Human | NuRD.1 complex | TRUE |
| O43765 | Q9H6T3 | -0.165 | | HOM | TRUE | FALSE | FALSE | | | | FALSE |
| P62136 | P62140 | -0.163 | HOM | | TRUE | FALSE | FALSE | | | | TRUE |
| Q9BXB4 | Q9BZF1 | -0.163 | | | FALSE | TRUE | FALSE | | | | FALSE |
| P61981 | P63104 | -0.161 | | | FALSE | TRUE | TRUE | 5199 | Human | Kinase maturation complex 1 | TRUE |
| P38159 | P98179 | -0.161 | | | FALSE | TRUE | FALSE | | | | TRUE |
| P62942 | Q14318 | -0.161 | HOM | HET | TRUE | FALSE | FALSE | | | | FALSE |
| P68371 | Q9BUF5 | -0.16 | HET | HET | FALSE | FALSE | FALSE | | | | TRUE |
| O43491 | Q9H4G0 | -0.16 | | | FALSE | TRUE | FALSE | | | | TRUE |
| P02545 | Q03252 | -0.159 | HOM | | TRUE | FALSE | FALSE | | | | TRUE |
| P07196 | P20700 | -0.159 | | | FALSE | TRUE | FALSE | | | | FALSE |
| O15479 | Q9UNF1 | -0.158 | | | FALSE | TRUE | FALSE | | | | FALSE |
| Q16643 | Q9UJU6 | -0.157 | | | FALSE | TRUE | FALSE | | | | FALSE |
| O95487 | P53992 | -0.155 | HET | HET | FALSE | FALSE | FALSE | | | | FALSE |
| P38919 | P60842 | -0.155 | HET | HET | FALSE | FALSE | FALSE | | | | TRUE |
| P08133 | Q5VT79 | -0.155 | | | FALSE | TRUE | FALSE | | | | FALSE |
| P43034 | Q96DI7 | -0.154 | HOM | | TRUE | FALSE | FALSE | | | | FALSE |
| Q00013 | Q9UDY2 | -0.154 | HOM | HOM | TRUE | FALSE | FALSE | | | | FALSE |
| P43357 | Q96MG7 | -0.15 | | HET | FALSE | FALSE | FALSE | | | | FALSE |
| Q15637 | Q96PU8 | -0.147 | | HOM | TRUE | FALSE | FALSE | | | | FALSE |
| Q969X5 | Q9Y282 | -0.147 | | | FALSE | TRUE | FALSE | | | | TRUE |
| P35908 | Q5XKE5 | -0.146 | | | FALSE | TRUE | FALSE | | | | FALSE |
| O15479 | P43363 | -0.146 | | | FALSE | TRUE | FALSE | | | | FALSE |
| P51911 | Q15417 | -0.145 | | | FALSE | TRUE | FALSE | | | | FALSE |
| P31944 | P55212 | -0.145 | | HOM | TRUE | FALSE | FALSE | | | | FALSE |
| Q96NB2 | Q9H9B4 | -0.144 | | | FALSE | TRUE | FALSE | | | | FALSE |

Table S4 Human

| | | | | | | | | | | | |
|---|---|---|---|---|---|---|---|---|---|---|---|
| P22626 | P51991 | -0.141 | | | FALSE | TRUE | TRUE | 1181 | Human | C complex spliceosome | TRUE |
| O60732 | P43358 | -0.14 | | | FALSE | TRUE | FALSE | | | | FALSE |
| P26885 | Q9Y680 | -0.14 | | | FALSE | TRUE | FALSE | | | | FALSE |
| Q13356 | Q9H2H8 | -0.14 | | | FALSE | TRUE | TRUE | 351 | Human | Spliceosome | FALSE |
| P15311 | P35240 | -0.139 | | HOM | TRUE | FALSE | FALSE | | | | TRUE |
| P00352 | P47895 | -0.139 | | | FALSE | TRUE | FALSE | | | | FALSE |
| P61289 | Q06323 | -0.138 | | HOM | TRUE | FALSE | FALSE | | | | FALSE |
| P12268 | P36959 | -0.138 | | | FALSE | TRUE | FALSE | | | | FALSE |
| P13647 | Q5XKE5 | -0.137 | | | FALSE | TRUE | FALSE | | | | FALSE |
| P53814 | Q8TDZ2 | -0.135 | | | FALSE | TRUE | FALSE | | | | FALSE |
| P36873 | P62140 | -0.135 | | | FALSE | TRUE | FALSE | | | | TRUE |
| P06396 | Q13045 | -0.135 | | | FALSE | TRUE | FALSE | | | | FALSE |
| P04350 | Q9BVA1 | -0.134 | HET | HET | FALSE | FALSE | FALSE | | | | FALSE |
| P07947 | P12931 | -0.134 | | HOM | TRUE | FALSE | FALSE | | | | FALSE |
| P07437 | Q9BVA1 | -0.133 | HET | HET | FALSE | FALSE | FALSE | | | | FALSE |
| P04350 | Q9BUF5 | -0.133 | HET | HET | FALSE | FALSE | FALSE | | | | FALSE |
| P35240 | P35241 | -0.132 | HOM | | TRUE | FALSE | FALSE | | | | FALSE |
| P50238 | Q16527 | -0.131 | | | FALSE | TRUE | FALSE | | | | FALSE |
| O00159 | O94832 | -0.129 | HOM | HOM | TRUE | FALSE | FALSE | | | | FALSE |
| P21399 | Q99798 | -0.126 | | | FALSE | TRUE | FALSE | | | | FALSE |
| P12829 | P60660 | -0.125 | | | FALSE | TRUE | FALSE | | | | FALSE |
| O94826 | Q9H3U1 | -0.125 | HOM | HOM | TRUE | FALSE | FALSE | | | | FALSE |
| Q13268 | Q99714 | -0.125 | | HOM | TRUE | FALSE | FALSE | | | | FALSE |
| P11310 | Q9H845 | -0.123 | HOM | | TRUE | FALSE | FALSE | | | | FALSE |
| O60711 | Q96HC4 | -0.122 | | | FALSE | TRUE | FALSE | | | | FALSE |
| Q9H223 | Q9UBC2 | -0.122 | HET | HET | FALSE | FALSE | FALSE | | | | FALSE |

Table S4 Human

| | | | | | | | | |
|---|---|---|---|---|---|---|---|---|
| Q8IVL5 | Q8IVL6 | -0.119 | | | FALSE | TRUE | FALSE | FALSE |
| P31483 | Q01085 | -0.119 | | | FALSE | TRUE | FALSE | FALSE |
| P43357 | P43363 | -0.119 | | | FALSE | TRUE | FALSE | FALSE |
| P08133 | P50995 | -0.119 | | | FALSE | TRUE | FALSE | FALSE |
| P19367 | Q2TB90 | -0.118 | HOM | | TRUE | FALSE | FALSE | TRUE |
| O75369 | Q8NF91 | -0.116 | | HOM | TRUE | FALSE | FALSE | FALSE |
| P68371 | Q13509 | -0.116 | HET | HET | FALSE | FALSE | FALSE | FALSE |
| P02545 | P14136 | -0.115 | HOM | | TRUE | FALSE | FALSE | FALSE |
| P31947 | P63104 | -0.114 | HOM | | TRUE | FALSE | FALSE | TRUE |
| O95881 | O95994 | -0.112 | | HOM | TRUE | FALSE | FALSE | FALSE |
| P34932 | Q9Y4L1 | -0.112 | HOM | HOM | TRUE | FALSE | FALSE | FALSE |
| P43007 | Q15758 | -0.111 | | | FALSE | TRUE | FALSE | FALSE |
| P09455 | Q01469 | -0.111 | | | FALSE | TRUE | FALSE | FALSE |
| P61106 | Q15907 | -0.109 | HET | HET | FALSE | FALSE | FALSE | FALSE |
| Q9GZM7 | Q9UBR2 | -0.108 | | | FALSE | TRUE | FALSE | FALSE |
| P08670 | P14136 | -0.107 | HOM | | TRUE | FALSE | FALSE | TRUE |
| O60437 | Q9UPN3 | -0.107 | HET | | FALSE | FALSE | FALSE | FALSE |
| O43447 | P45877 | -0.106 | HET | | FALSE | FALSE | FALSE | FALSE |
| Q92945 | Q96I24 | -0.106 | | | FALSE | TRUE | FALSE | TRUE |
| P12429 | Q5VT79 | -0.106 | | | FALSE | TRUE | FALSE | FALSE |
| P06753 | P07951 | -0.103 | | | FALSE | TRUE | FALSE | TRUE |
| Q00013 | Q07157 | -0.103 | HOM | HOM | TRUE | FALSE | FALSE | FALSE |
| P15144 | Q9NZ08 | -0.102 | | HET | FALSE | FALSE | FALSE | FALSE |
| O95425 | Q13045 | -0.1 | | | FALSE | TRUE | FALSE | FALSE |
| P23284 | Q6UX04 | -0.1 | | | FALSE | TRUE | FALSE | FALSE |
| P04083 | P27216 | -0.1 | | | FALSE | TRUE | FALSE | FALSE |

Table S4 Human

| | | | | | | | | |
|---|---|---|---|---|---|---|---|---|
| P05121 | P36952 | -0.099 | HOM | | TRUE | FALSE | FALSE | FALSE |
| Q7L2E3 | Q9H2U1 | -0.099 | | | FALSE | TRUE | FALSE | FALSE |
| P38159 | Q14011 | -0.098 | | | FALSE | TRUE | FALSE | TRUE |
| Q15233 | Q8WXF1 | -0.095 | HET | HET | FALSE | FALSE | FALSE | TRUE |
| P53814 | Q8NDI1 | -0.095 | | | FALSE | TRUE | FALSE | FALSE |
| P26038 | P35240 | -0.094 | | HOM | TRUE | FALSE | FALSE | FALSE |
| P14136 | Q16352 | -0.094 | | | FALSE | TRUE | FALSE | TRUE |
| O60343 | Q9Y3P9 | -0.093 | | | FALSE | TRUE | FALSE | FALSE |
| P35580 | P35749 | -0.092 | | | FALSE | TRUE | FALSE | FALSE |
| Q9H4M9 | Q9UBC2 | -0.091 | HET | HET | FALSE | FALSE | FALSE | FALSE |
| P49419 | Q8IZ83 | -0.091 | HOM | | TRUE | FALSE | FALSE | FALSE |
| P40121 | Q13045 | -0.091 | | | FALSE | TRUE | FALSE | FALSE |
| P42765 | Q9BWD1 | -0.09 | | HOM | TRUE | FALSE | FALSE | FALSE |
| Q01105 | Q86VY4 | -0.09 | HOM | | TRUE | FALSE | FALSE | FALSE |
| P04271 | P25815 | -0.09 | HOM | HOM | TRUE | FALSE | FALSE | TRUE |
| P42226 | P51692 | -0.089 | HOM | HOM | TRUE | FALSE | FALSE | FALSE |
| P04626 | P43403 | -0.088 | HOM | HOM | TRUE | FALSE | FALSE | FALSE |
| O00186 | P61764 | -0.088 | | | FALSE | TRUE | FALSE | FALSE |
| P08758 | Q5VT79 | -0.088 | | | FALSE | TRUE | FALSE | FALSE |
| P50995 | Q5VT79 | -0.087 | | | FALSE | TRUE | FALSE | FALSE |
| Q9NZN4 | Q9UBC2 | -0.087 | HOM, HET | HET | TRUE, FALSE | FALSE | FALSE | FALSE |
| P42224 | P51692 | -0.087 | HOM | HOM | TRUE | FALSE | FALSE | FALSE |
| O75475 | P51858 | -0.087 | | HOM | TRUE | FALSE | FALSE | TRUE |
| Q15555 | Q9UPY8 | -0.086 | | | FALSE | TRUE | FALSE | TRUE |
| P49773 | Q9BX68 | -0.086 | | | FALSE | TRUE | FALSE | FALSE |

Table S4 Human

| | | | | | | | | |
|---|---|---|---|---|---|---|---|---|
| P08575 | P10586 | -0.085 | | | FALSE | TRUE | FALSE | FALSE |
| P04259 | P13647 | -0.084 | | | FALSE | TRUE | FALSE | FALSE |
| Q13347 | Q9Y3F4 | -0.084 | HET | HET | FALSE | FALSE | FALSE | FALSE |
| P61006 | Q6IQ22 | -0.084 | HET,HOM | HOM | TRUE | FALSE | FALSE | FALSE |
| P40763 | P42226 | -0.083 | HOM | HOM | TRUE | FALSE | FALSE | TRUE |
| Q5T8P6 | Q9P2N5 | -0.082 | | | FALSE | TRUE | FALSE | FALSE |
| P35237 | P50453 | -0.082 | | | FALSE | TRUE | FALSE | FALSE |
| P21926 | P60033 | -0.081 | HOM | HOM | TRUE | FALSE | FALSE | TRUE |
| P02751 | P24821 | -0.081 | HOM | HOM | TRUE | FALSE | FALSE | TRUE |
| P31943 | Q8IXT5 | -0.081 | | | FALSE | TRUE | FALSE | FALSE |
| Q15542 | Q8WWQ0 | -0.08 | | | FALSE | TRUE | FALSE | FALSE |
| P07355 | P20073 | -0.079 | HOM | | TRUE | FALSE | FALSE | FALSE |
| P30740 | P48594 | -0.076 | | | FALSE | TRUE | FALSE | FALSE |
| P31947 | P61981 | -0.076 | HOM | | TRUE | FALSE | FALSE | TRUE |
| O95235 | P33176 | -0.075 | | HOM | TRUE | FALSE | FALSE | FALSE |
| P58107 | Q9UPN3 | -0.073 | HET | | FALSE | FALSE | FALSE | FALSE |
| Q02388 | Q05707 | -0.072 | | | FALSE | TRUE | FALSE | FALSE |
| P43358 | Q96MG7 | -0.072 | | HET | FALSE | FALSE | FALSE | FALSE |
| Q07666 | Q96PU8 | -0.072 | HOM | HOM | TRUE | FALSE | FALSE | FALSE |
| O00151 | Q96JY6 | -0.071 | | | FALSE | TRUE | FALSE | FALSE |
| P08648 | P23229 | -0.071 | | | FALSE | TRUE | FALSE | FALSE |
| P30740 | P50453 | -0.07 | | | FALSE | TRUE | FALSE | FALSE |
| P30048 | P32119 | -0.069 | HOM | HOM | TRUE | FALSE | FALSE | TRUE |
| P13645 | P19012 | -0.069 | HET | | FALSE | FALSE | FALSE | FALSE |
| O15540 | Q01469 | -0.068 | | | FALSE | TRUE | FALSE | FALSE |
| P31947 | P62258 | -0.067 | HOM | HOM | TRUE | FALSE | FALSE | FALSE |

Table S4 Human

| | | | | | | | | | | | |
|---|---|---|---|---|---|---|---|---|---|---|---|
| P48059 | Q96HC4 | -0.067 | HET | | FALSE | FALSE | FALSE | | | | FALSE |
| P12429 | P27216 | -0.067 | | | FALSE | TRUE | FALSE | | | | FALSE |
| O00429 | P50570 | -0.066 | | | FALSE | TRUE | FALSE | | | | FALSE |
| P20337 | P51159 | -0.066 | HOM | HET,HOM | TRUE | FALSE | FALSE | | | | FALSE |
| Q13356 | Q9Y3C6 | -0.064 | | | FALSE | TRUE | TRUE | 351 | Human | Spliceosome | FALSE |
| O00629 | O60684 | -0.063 | HOM | HOM | TRUE | FALSE | FALSE | | | | FALSE |
| P13164 | Q01628 | -0.063 | | | FALSE | TRUE | FALSE | | | | FALSE |
| P62258 | Q04917 | -0.063 | HOM | | TRUE | FALSE | TRUE | 5199 | Human | Kinase maturation complex 1 | TRUE |
| P11940 | P29558 | -0.062 | HOM | | TRUE | FALSE | FALSE | | | | FALSE |
| Q13217 | Q99615 | -0.062 | | | FALSE | TRUE | FALSE | | | | FALSE |
| P26639 | Q9NYK5 | -0.062 | | | FALSE | TRUE | FALSE | | | | FALSE |
| B0I1T2 | O94832 | -0.06 | HOM | HOM | TRUE | FALSE | FALSE | | | | FALSE |
| P61006 | P61026 | -0.058 | HET,HOM | HOM | TRUE | FALSE | FALSE | | | | TRUE |
| O95486 | P53992 | -0.057 | HET | HET | FALSE | FALSE | FALSE | | | | TRUE |
| O43432 | Q04637 | -0.057 | HET | HOM, HET | TRUE, FALSE | FALSE | FALSE | | | | TRUE |
| P14317 | Q14247 | -0.057 | | HOM | TRUE | FALSE | FALSE | | | | FALSE |
| P13796 | Q14651 | -0.057 | | | FALSE | TRUE | FALSE | | | | TRUE |
| P16219 | Q9H845 | -0.056 | HOM | | TRUE | FALSE | FALSE | | | | FALSE |
| Q6NXG1 | Q9NTZ6 | -0.056 | | HOM | TRUE | FALSE | FALSE | | | | FALSE |
| O95235 | Q02241 | -0.056 | | HET | FALSE | FALSE | FALSE | | | | FALSE |
| P36405 | Q9NVJ2 | -0.055 | | | FALSE | TRUE | FALSE | | | | FALSE |
| Q96D15 | Q9BRK5 | -0.055 | | | FALSE | TRUE | FALSE | | | | FALSE |
| Q13451 | Q14318 | -0.054 | HET | HET | FALSE | FALSE | FALSE | | | | FALSE |
| Q9H4G0 | Q9Y2J2 | -0.054 | | | FALSE | TRUE | FALSE | | | | TRUE |
| Q13362 | Q16537 | -0.054 | HET | HET | FALSE | FALSE | FALSE | | | | TRUE |

Table S4 Human

| | | | | | | | | | | | |
|---|---|---|---|---|---|---|---|---|---|---|---|
| P49748 | Q92947 | -0.054 | HOM | HOM | TRUE | FALSE | FALSE | | | | FALSE |
| P47755 | P52907 | -0.053 | | | FALSE | TRUE | FALSE | | | | TRUE |
| P35237 | P48594 | -0.053 | | | FALSE | TRUE | FALSE | | | | FALSE |
| Q13098 | Q15008 | -0.052 | | | FALSE | TRUE | FALSE | | | | FALSE |
| P14550 | P15121 | -0.051 | | HOM | TRUE | FALSE | FALSE | | | | FALSE |
| Q14258 | Q96LD4 | -0.051 | | | FALSE | TRUE | FALSE | | | | FALSE |
| O14964 | Q8IWB7 | -0.051 | HET,HOM | | FALSE,TRUE | FALSE | FALSE | | | | FALSE |
| O75170 | Q5H9R7 | -0.051 | HET | HET | FALSE | FALSE | FALSE | | | | TRUE |
| P31946 | P61981 | -0.049 | | | FALSE | TRUE | TRUE | 5199 | Human | Kinase maturation complex 1 | TRUE |
| P19012 | P35527 | -0.049 | | | FALSE | TRUE | FALSE | | | | FALSE |
| P05787 | P13647 | -0.047 | | | FALSE | TRUE | FALSE | | | | FALSE |
| Q00013 | Q12959 | -0.047 | HOM | HOM | TRUE | FALSE | FALSE | | | | FALSE |
| Q15691 | Q9UPY8 | -0.046 | HOM | | TRUE | FALSE | FALSE | | | | TRUE |
| P24386 | P50395 | -0.046 | HET | HET | FALSE | FALSE | FALSE | | | | FALSE |
| P28288 | P33897 | -0.045 | HET | HOM | TRUE | FALSE | FALSE | | | | TRUE |
| P23246 | Q8WXF1 | -0.045 | HOM | HET | TRUE | FALSE | FALSE | | | | TRUE |
| P07355 | P08133 | -0.044 | HOM | | TRUE | FALSE | FALSE | | | | FALSE |
| Q9H2H8 | Q9Y3C6 | -0.044 | | | FALSE | TRUE | TRUE | 351,1181 | Human | Spliceosome,C complex spliceosome | FALSE |
| P07355 | P50995 | -0.042 | HOM | | TRUE | FALSE | FALSE | | | | TRUE |
| P13861 | P31323 | -0.041 | HOM | | TRUE | FALSE | TRUE | 878 | Human | PKA-AKAP5-ADRB1 complex | TRUE |
| P51911 | Q99439 | -0.041 | | | FALSE | TRUE | FALSE | | | | FALSE |
| Q96MU7 | Q9BYJ9 | -0.041 | | | FALSE | TRUE | FALSE | | | | FALSE |
| P55786 | Q9NZ08 | -0.041 | | HET | FALSE | FALSE | FALSE | | | | FALSE |
| O75525 | Q96PU8 | -0.04 | HOM | HOM | TRUE | FALSE | FALSE | | | | FALSE |
| Q13356 | Q6UX04 | -0.039 | | | FALSE | TRUE | FALSE | | | | FALSE |

Table S4 Human

| | | | | | | | | |
|---|---|---|---|---|---|---|---|---|
| P20337 | Q6IQ22 | -0.039 | HOM | HOM | TRUE | FALSE | FALSE | FALSE |
| P62820 | Q9NP72 | -0.038 | HET | HET | FALSE | FALSE | FALSE | FALSE |
| P61586 | P62745 | -0.038 | HOM | | TRUE | FALSE | FALSE | FALSE |
| P43357 | Q9UNF1 | -0.036 | | | FALSE | TRUE | FALSE | FALSE |
| P13647 | Q7Z794 | -0.036 | | | FALSE | TRUE | FALSE | FALSE |
| Q92841 | Q9NR30 | -0.036 | | HOM | TRUE | FALSE | FALSE | FALSE |
| O60711 | Q9NR12 | -0.034 | | | FALSE | TRUE | FALSE | TRUE |
| P04264 | Q5XKE5 | -0.034 | HET | | FALSE | FALSE | FALSE | FALSE |
| P04083 | Q5VT79 | -0.034 | | | FALSE | TRUE | FALSE | FALSE |
| P08670 | Q03252 | -0.033 | HOM | | TRUE | FALSE | FALSE | FALSE |
| P09327 | P40121 | -0.033 | HOM | | TRUE | FALSE | FALSE | FALSE |
| P52789 | Q2TB90 | -0.031 | | | FALSE | TRUE | FALSE | FALSE |
| P31146 | Q9BR76 | -0.031 | | HOM | TRUE | FALSE | FALSE | TRUE |
| P06756 | P23229 | -0.031 | HET | | FALSE | FALSE | FALSE | FALSE |
| P37802 | P51911 | -0.03 | | | FALSE | TRUE | FALSE | FALSE |
| O75525 | Q15637 | -0.03 | HOM | | TRUE | FALSE | FALSE | FALSE |
| P02511 | P04792 | -0.029 | HOM | | TRUE | FALSE | FALSE | TRUE |
| O15020 | O43707 | -0.028 | | | FALSE | TRUE | FALSE | FALSE |
| O95833 | Q9Y696 | -0.028 | | | FALSE | TRUE | FALSE | FALSE |
| P05787 | P35908 | -0.028 | | | FALSE | TRUE | FALSE | FALSE |
| O43795 | Q12965 | -0.028 | HOM | HOM | TRUE | FALSE | FALSE | FALSE |
| P30622 | Q99426 | -0.025 | | HET | FALSE | FALSE | FALSE | FALSE |
| P31944 | Q14790 | -0.024 | | HET | FALSE | FALSE | FALSE | TRUE |
| P12814 | Q13813 | -0.024 | | HOM | TRUE | FALSE | FALSE | FALSE |
| P07355 | P27216 | -0.023 | HOM | | TRUE | FALSE | FALSE | TRUE |
| P05937 | P22676 | -0.021 | | | FALSE | TRUE | FALSE | FALSE |

Table S4 Human

| | | | | | | | | | | | |
|---|---|---|---|---|---|---|---|---|---|---|---|
| P06753 | P67936 | -0.021 | | | FALSE | TRUE | FALSE | | | | TRUE |
| P29373 | Q01469 | -0.02 | HOM | | TRUE | FALSE | FALSE | | | | FALSE |
| P00352 | P49419 | -0.019 | | HOM | TRUE | FALSE | FALSE | | | | FALSE |
| P50151 | Q9UBI6 | -0.019 | | | FALSE | TRUE | FALSE | | | | FALSE |
| P02545 | P07196 | -0.018 | HOM | | TRUE | FALSE | FALSE | | | | FALSE |
| Q7Z2T5 | Q9NXH9 | -0.016 | | | FALSE | TRUE | FALSE | | | | FALSE |
| P06733 | P09104 | -0.016 | HOM | HOM | TRUE | FALSE | FALSE | | | | TRUE |
| P11171 | Q9H4G0 | -0.015 | | | FALSE | TRUE | FALSE | | | | FALSE |
| P46063 | P54132 | -0.015 | HOM | | TRUE | FALSE | FALSE | | | | FALSE |
| P05091 | P47895 | -0.015 | HOM | | TRUE | FALSE | FALSE | | | | FALSE |
| Q0D2K2 | Q2WGJ6 | -0.015 | | | FALSE | TRUE | FALSE | | | | FALSE |
| P02100 | P69892 | -0.014 | | | FALSE | TRUE | FALSE | | | | FALSE |
| O75340 | Q9UBV8 | -0.014 | HOM | HET | TRUE | FALSE | TRUE | 6375 | Human | PEF1-ALG2 complex | TRUE |
| Q6UX04 | Q9H2H8 | -0.014 | | | FALSE | TRUE | TRUE | 1181 | Human | C complex spliceosome | FALSE |
| Q9BXB4 | Q9BXB5 | -0.013 | | | FALSE | TRUE | FALSE | | | | FALSE |
| P35240 | Q9Y4F1 | -0.011 | HOM | | TRUE | FALSE | FALSE | | | | FALSE |
| P11940 | Q15717 | -0.011 | HOM | HOM | TRUE | FALSE | FALSE | | | | TRUE |
| P04264 | Q7Z794 | -0.011 | HET | | FALSE | FALSE | FALSE | | | | TRUE |
| P35527 | Q04695 | -0.01 | | | FALSE | TRUE | FALSE | | | | FALSE |
| P08729 | P35908 | -0.01 | | | FALSE | TRUE | FALSE | | | | FALSE |
| P16144 | P18084 | -0.009 | HET | | FALSE | FALSE | FALSE | | | | FALSE |
| P52597 | Q8IXT5 | -0.009 | | | FALSE | TRUE | FALSE | | | | FALSE |
| P21291 | P50238 | -0.009 | | | FALSE | TRUE | FALSE | | | | FALSE |
| P63104 | Q04917 | -0.009 | | | FALSE | TRUE | TRUE | 5199 | Human | Kinase maturation complex 1 | TRUE |
| O95757 | Q9Y4L1 | -0.008 | HOM | HOM | TRUE | FALSE | FALSE | | | | FALSE |
| P15153 | P60953 | -0.008 | HET | HOM | TRUE | FALSE | FALSE | | | | FALSE |

Table S4 Human

| | | | | | | | | |
|---|---|---|---|---|---|---|---|---|
| O60218 | P42330 | -0.007 | | | FALSE | TRUE | FALSE | FALSE |
| P26038 | Q9Y4F1 | -0.007 | | | FALSE | TRUE | FALSE | FALSE |
| P29558 | Q9UHX1 | -0.006 | | HOM | TRUE | FALSE | FALSE | FALSE |
| P04259 | Q7Z794 | -0.005 | | | FALSE | TRUE | FALSE | FALSE |
| P42025 | P63261 | -0.004 | | | FALSE | TRUE | FALSE | FALSE |
| P08758 | P27216 | -0.004 | | | FALSE | TRUE | FALSE | FALSE |
| P12277 | P12532 | -0.004 | | HOM | TRUE | FALSE | FALSE | FALSE |
| P48163 | Q16798 | -0.004 | | | FALSE | TRUE | FALSE | FALSE |
| P58107 | Q92817 | -0.004 | HET | HOM | TRUE | FALSE | FALSE | FALSE |
| P31942 | P55795 | -0.004 | | | FALSE | TRUE | FALSE | TRUE |
| O94804 | Q13188 | -0.003 | HOM | | TRUE | FALSE | FALSE | FALSE |
| P23368 | Q16798 | -0.003 | HOM | | TRUE | FALSE | FALSE | FALSE |
| Q7Z739 | Q9BYJ9 | -0.003 | | | FALSE | TRUE | FALSE | TRUE |
| Q6NXG1 | Q8IXT5 | -0.002 | | | FALSE | TRUE | FALSE | FALSE |
| P17931 | P56470 | -0.002 | HET | | FALSE | FALSE | FALSE | FALSE |
| Q8ND56 | Q9BX40 | -0.002 | | | FALSE | TRUE | FALSE | FALSE |
| Q86UX7 | Q96AC1 | -0.001 | | | FALSE | TRUE | FALSE | TRUE |
| P34931 | P54652 | -0.001 | | | FALSE | TRUE | FALSE | FALSE |

Table S4 Human

| | | | | | | | | | | | |
|---|---|---|---|---|---|---|---|---|---|---|---|
| Q13547 | Q92769 | 0.0004 | | | FALSE | TRUE | TRUE | 61,282,614,620,632,633,634,636,642,685,696,714,732,738,739,743,749,778,871,888,1257,1505,2721,2814,2851,3167,6373,6421 | Human | Mi2/NuRD complex,SNF2h-cohesin-NuRD complex,NRD complex,CoREST-HDAC complex,Anti-HDAC2 complex,anti-BHC110 complex,XFIM complex,BHC complex,CtBP complex,MeCP1 complex,BRMS1-SIN3-HDAC complex,BRM-SIN3A complex,SIN3 complex,SIN3-ING1b complex I,SIN3-ING1b complex II,SIN3-SAP25 complex,MeCP2-SIN3A-HDAC complex,LARC complex,BRAF53-BRCA2 complex,MTA2 complex,ALL-1 supercomplex,NCOR2 complex,HCF-1 complex,BRCA1- | TRUE |
| P08727 | Q04695 | 0.0005 | | | FALSE | TRUE | FALSE | | | | FALSE |
| Q96CW1 | Q9Y6Q5 | 0.0015 | HET | HET | FALSE | FALSE | FALSE | | | | FALSE |
| P53814 | Q5M775 | 0.0016 | | | FALSE | TRUE | FALSE | | | | FALSE |
| O15226 | Q9NXV6 | 0.0018 | | | FALSE | TRUE | FALSE | | | | FALSE |
| O14787 | Q92973 | 0.0029 | | HOM | TRUE | FALSE | FALSE | | | | FALSE |
| P27708 | P31327 | 0.0041 | | HOM | TRUE | FALSE | FALSE | | | | FALSE |

Table S4 Human

| | | | | | | | | | | | |
|---|---|---|---|---|---|---|---|---|---|---|---|
| Q8TAQ2 | Q92922 | 0.0045 | | | FALSE | TRUE | TRUE | 86,149,189,238,570,713,714,739,778,806,1230,1237,1239,1252,1257,1413,2829,3064,3065,3067,5614 | Human | NUMAC complex,PBAF complex,BAF complex,SWI-SNF chromatin remodeling-related-BRCA1 complex,p300-CBP-p270-SWI/SNF complex,BRG1-SIN3A complex,BRM-SIN3A complex,SIN3-ING1b complex II,LARC complex,BRM-SIN3A-HDAC complex,WINAC complex,EBAFb complex,EBAFa complex,ALL-1 supercomplex,NCOR1 complex,RSmad complex,RNA polymerase II complex, chromatin structure modifying,RNA polymerase II complex, incomplete, chromatin structure modifying,Emerin | TRUE |
| P30405 | P62937 | 0.0057 | | HOM | TRUE | FALSE | FALSE | | | | FALSE |
| P01040 | P04080 | 0.0058 | HOM | HOM | TRUE | FALSE | FALSE | | | | FALSE |
| P16219 | Q92947 | 0.0059 | HOM | HOM | TRUE | FALSE | FALSE | | | | FALSE |
| Q08257 | Q99536 | 0.0079 | HOM | | TRUE | FALSE | FALSE | | | | FALSE |
| P46734 | Q02750 | 0.0082 | | HOM | TRUE | FALSE | FALSE | | | | FALSE |
| P07196 | P14136 | 0.0092 | | | FALSE | TRUE | FALSE | | | | FALSE |
| P51114 | Q06787 | 0.0094 | | HET | FALSE | FALSE | FALSE | | | | TRUE |
| Q96RT1 | Q9H9A6 | 0.0095 | | | FALSE | TRUE | FALSE | | | | FALSE |
| P06396 | P09327 | 0.0101 | | HOM | TRUE | FALSE | FALSE | | | | FALSE |
| P17643 | P40126 | 0.0102 | | | FALSE | TRUE | FALSE | | | | FALSE |
| P24752 | Q9BWD1 | 0.0103 | HOM | HOM | TRUE | FALSE | FALSE | | | | TRUE |
| O94826 | Q99614 | 0.0104 | HOM | HOM | TRUE | FALSE | FALSE | | | | FALSE |

Table S4 Human

| | | | | | | | | | | | |
|---|---|---|---|---|---|---|---|---|---|---|---|
| P38159 | P62995 | 0.0106 | | | FALSE | TRUE | FALSE | | | | TRUE |
| P50479 | Q96JY6 | 0.0109 | HOM | | TRUE | FALSE | FALSE | | | | FALSE |
| O43294 | Q13642 | 0.0122 | | | FALSE | TRUE | FALSE | | | | FALSE |
| P04259 | P08729 | 0.0126 | | | FALSE | TRUE | FALSE | | | | FALSE |
| P13645 | Q04695 | 0.0128 | HET | | FALSE | FALSE | FALSE | | | | FALSE |
| O95466 | Q27J81 | 0.013 | | | FALSE | TRUE | FALSE | | | | FALSE |
| P04259 | P05787 | 0.0139 | | | FALSE | TRUE | FALSE | | | | FALSE |
| P14317 | Q16643 | 0.0146 | | | FALSE | TRUE | FALSE | | | | FALSE |
| P43355 | P43363 | 0.0146 | | | FALSE | TRUE | FALSE | | | | FALSE |
| P35579 | P35580 | 0.0151 | | | FALSE | TRUE | FALSE | | | | TRUE |
| P45877 | Q9H2H8 | 0.0152 | | | FALSE | TRUE | FALSE | | | | FALSE |
| O95757 | Q92598 | 0.0156 | HOM | HOM | TRUE | FALSE | FALSE | | | | FALSE |
| P04350 | Q13509 | 0.0156 | HET | HET | FALSE | FALSE | FALSE | | | | FALSE |
| O00299 | Q9Y696 | 0.0158 | HOM | | TRUE | FALSE | FALSE | | | | FALSE |
| O15230 | P55268 | 0.0173 | | | FALSE | TRUE | TRUE | 2319 | Human | ITGA6-ITGB4-Laminin10/12 complex | FALSE |
| P13667 | Q8IXB1 | 0.0174 | | | FALSE | TRUE | FALSE | | | | FALSE |
| O43143 | Q92620 | 0.0175 | | | FALSE | TRUE | TRUE | 351 | Human | Spliceosome | FALSE |
| P48960 | Q6QNK2 | 0.0175 | HET | | FALSE | FALSE | FALSE | | | | FALSE |
| P21291 | Q16527 | 0.0179 | | | FALSE | TRUE | FALSE | | | | FALSE |
| P20073 | Q5VT79 | 0.0191 | | | FALSE | TRUE | FALSE | | | | FALSE |
| P22626 | Q13151 | 0.02 | | | FALSE | TRUE | FALSE | | | | FALSE |
| O00151 | P50479 | 0.0201 | | HOM | TRUE | FALSE | FALSE | | | | FALSE |
| P05121 | P07093 | 0.0213 | HOM | | TRUE | FALSE | FALSE | | | | FALSE |
| P27105 | Q9UJZ1 | 0.0217 | HOM | HOM | TRUE | FALSE | FALSE | | | | TRUE |
| P43355 | P43357 | 0.0224 | | | FALSE | TRUE | FALSE | | | | FALSE |

Table S4 Human

| | | | | | | | | |
|---|---|---|---|---|---|---|---|---|
| P40306 | Q99436 | 0.0226 | HOM | HOM | TRUE | FALSE | FALSE | FALSE |
| P08133 | P27216 | 0.0228 | | | FALSE | TRUE | FALSE | FALSE |
| P04259 | P04264 | 0.0231 | | HET | FALSE | FALSE | FALSE | FALSE |
| P42566 | Q9NZN4 | 0.0236 | HOM, HET | HOM, HET | TRUE, FALSE | FALSE | FALSE | FALSE |
| O43175 | Q13363 | 0.0238 | HOM | HET | TRUE | FALSE | FALSE | FALSE |
| O43813 | Q9NS86 | 0.0241 | | | FALSE | TRUE | FALSE | FALSE |
| P07203 | Q8TED1 | 0.0257 | HOM | | TRUE | FALSE | FALSE | FALSE |
| P30405 | P49792 | 0.0257 | | HET | FALSE | FALSE | FALSE | FALSE |
| O43396 | P10599 | 0.0262 | HET | HOM, HET | TRUE, FALSE | FALSE | FALSE | FALSE |
| Q8IVL5 | Q92791 | 0.0264 | | | FALSE | TRUE | FALSE | FALSE |
| O75340 | P30626 | 0.0266 | HOM | | TRUE | FALSE | FALSE | FALSE |
| O43294 | P48059 | 0.0269 | | HET | FALSE | FALSE | FALSE | FALSE |
| P42025 | P60709 | 0.027 | | HET | FALSE | FALSE | FALSE | FALSE |
| P08727 | P35527 | 0.0273 | | | FALSE | TRUE | FALSE | FALSE |
| Q9NRV9 | Q9Y5Z4 | 0.0274 | | | FALSE | TRUE | FALSE | FALSE |
| P09327 | Q13045 | 0.0278 | HOM | | TRUE | FALSE | FALSE | FALSE |
| P27216 | Q5VT79 | 0.028 | | | FALSE | TRUE | FALSE | FALSE |
| P18085 | P84085 | 0.0281 | | | FALSE | TRUE | FALSE | FALSE |
| P11310 | P16219 | 0.0294 | HOM | HOM | TRUE | FALSE | FALSE | FALSE |
| P15121 | P52895 | 0.0295 | HOM | HOM | TRUE | FALSE | FALSE | FALSE |
| P07900 | Q12931 | 0.0295 | HOM | HOM | TRUE | FALSE | FALSE | FALSE |
| P45877 | Q13356 | 0.0299 | | | FALSE | TRUE | FALSE | FALSE |
| O75874 | P48735 | 0.0301 | HOM | HOM | TRUE | FALSE | FALSE | TRUE |
| O76070 | P37840 | 0.0302 | | | FALSE | TRUE | FALSE | FALSE |
| P61163 | P63261 | 0.0302 | | | FALSE | TRUE | FALSE | FALSE |

Table S4 Human

| | | | | | | | | | | | |
|---|---|---|---|---|---|---|---|---|---|---|---|
| O95810 | Q969G5 | 0.0307 | | | FALSE | TRUE | FALSE | | | | FALSE |
| O75131 | Q99829 | 0.0316 | | HOM | TRUE | FALSE | FALSE | | | | FALSE |
| P23297 | P60903 | 0.0317 | HET | | FALSE | FALSE | FALSE | | | | FALSE |
| P36405 | P84077 | 0.032 | | HOM | TRUE | FALSE | FALSE | | | | FALSE |
| P58107 | Q15149 | 0.0328 | HET | HOM | TRUE | FALSE | FALSE | | | | FALSE |
| Q9Y490 | Q9Y4G6 | 0.0331 | HOM | | TRUE | FALSE | FALSE | | | | FALSE |
| P43363 | Q96MG7 | 0.0334 | | HET | FALSE | FALSE | FALSE | | | | FALSE |
| P07437 | Q13509 | 0.0334 | HET | HET | FALSE | FALSE | FALSE | | | | TRUE |
| P06703 | P23297 | 0.0337 | HOM | HET | TRUE | FALSE | FALSE | | | | FALSE |
| Q86XP3 | Q92841 | 0.0345 | | | FALSE | TRUE | FALSE | | | | FALSE |
| P25815 | P60903 | 0.0346 | HOM | | TRUE | FALSE | FALSE | | | | FALSE |
| O95232 | Q9NQ29 | 0.0346 | HOM | | TRUE | FALSE | FALSE | | | | TRUE |
| P50851 | Q6ZNJ1 | 0.0351 | | | FALSE | TRUE | FALSE | | | | FALSE |
| Q02978 | Q9UBX3 | 0.0352 | | | FALSE | TRUE | FALSE | | | | FALSE |
| O15479 | P43357 | 0.0357 | | | FALSE | TRUE | FALSE | | | | FALSE |
| Q5T2T1 | Q9NZW5 | 0.0378 | HOM | HOM | TRUE | FALSE | FALSE | | | | FALSE |
| P40763 | P42224 | 0.0385 | HOM | HOM | TRUE | FALSE | FALSE | | | | TRUE |
| P57721 | P61978 | 0.0386 | | HOM | TRUE | FALSE | FALSE | | | | FALSE |
| P30530 | Q12866 | 0.0392 | HET | | FALSE | FALSE | FALSE | | | | FALSE |
| O60218 | P52895 | 0.0398 | | HOM | TRUE | FALSE | FALSE | | | | FALSE |
| P35241 | Q9Y4F1 | 0.0399 | | | FALSE | TRUE | FALSE | | | | FALSE |
| Q96RT1 | Q9BTT6 | 0.0407 | | | FALSE | TRUE | FALSE | | | | FALSE |
| Q15293 | Q96D15 | 0.041 | | | FALSE | TRUE | FALSE | | | | FALSE |
| O60231 | Q92620 | 0.0413 | | | FALSE | TRUE | TRUE | 351 | Human | Spliceosome | TRUE |
| Q9BXB5 | Q9BZF1 | 0.0418 | | | FALSE | TRUE | FALSE | | | | FALSE |
| P61764 | Q15833 | 0.0425 | | | FALSE | TRUE | FALSE | | | | FALSE |

Table S4 Human

| | | | | | | | | | | | |
|---|---|---|---|---|---|---|---|---|---|---|---|
| Q8IXT5 | Q9NTZ6 | 0.0431 | | HOM | TRUE | FALSE | FALSE | | | | FALSE |
| P16104 | Q8IUE6 | 0.0431 | HET | HET | FALSE | FALSE | FALSE | | | | TRUE |
| P13646 | Q04695 | 0.0437 | | | FALSE | TRUE | FALSE | | | | FALSE |
| P84077 | P84085 | 0.0443 | HOM | | TRUE | FALSE | FALSE | | | | TRUE |
| P43121 | Q13740 | 0.0456 | | | FALSE | TRUE | FALSE | | | | FALSE |
| P04259 | Q5XKE5 | 0.046 | | | FALSE | TRUE | FALSE | | | | FALSE |
| P04083 | P08758 | 0.046 | | | FALSE | TRUE | FALSE | | | | TRUE |
| O95758 | Q8WVV9 | 0.0465 | | | FALSE | TRUE | FALSE | | | | FALSE |
| P34896 | P34897 | 0.0466 | | | FALSE | TRUE | FALSE | | | | TRUE |
| Q13617 | Q93034 | 0.0469 | HET | HET | FALSE | FALSE | FALSE | | | | TRUE |
| Q02388 | Q99715 | 0.047 | | | FALSE | TRUE | FALSE | | | | FALSE |
| Q14739 | Q9UBM7 | 0.0481 | | | FALSE | TRUE | FALSE | | | | FALSE |
| P20073 | P27216 | 0.0489 | | | FALSE | TRUE | FALSE | | | | FALSE |
| O60231 | Q14562 | 0.0508 | | HOM | TRUE | FALSE | TRUE | 351 | Human | Spliceosome | FALSE |
| P42785 | Q9UHL4 | 0.0509 | HOM | HOM | TRUE | FALSE | FALSE | | | | FALSE |
| O00629 | P52294 | 0.051 | HOM | HOM | TRUE | FALSE | FALSE | | | | FALSE |
| Q9NZ08 | Q9UIQ6 | 0.051 | HET | | FALSE | FALSE | FALSE | | | | FALSE |
| O43294 | O60711 | 0.0514 | | | FALSE | TRUE | FALSE | | | | FALSE |
| P05091 | P49419 | 0.0514 | HOM | HOM | TRUE | FALSE | FALSE | | | | FALSE |
| P04259 | P35908 | 0.0515 | | | FALSE | TRUE | FALSE | | | | FALSE |
| P05164 | Q92626 | 0.0528 | | | FALSE | TRUE | FALSE | | | | FALSE |
| O43491 | Q9Y2J2 | 0.0528 | | | FALSE | TRUE | FALSE | | | | TRUE |
| P69891 | P69905 | 0.0533 | HOM | HOM | TRUE | FALSE | FALSE | | | | FALSE |
| Q8N129 | Q9BT09 | 0.0541 | | | FALSE | TRUE | FALSE | | | | FALSE |
| P23229 | P26006 | 0.0544 | | | FALSE | TRUE | FALSE | | | | FALSE |
| A0AVT1 | P22314 | 0.0544 | | | FALSE | TRUE | FALSE | | | | FALSE |

Table S4 Human

| | | | | | | | | |
|---|---|---|---|---|---|---|---|---|
| P06239 | P12931 | 0.0547 | HOM | HOM | TRUE | FALSE | FALSE | FALSE |
| P07437 | Q9BUF5 | 0.0573 | HET | HET | FALSE | FALSE | FALSE | FALSE |
| Q9ULC6 | Q9ULW8 | 0.0576 | | | FALSE | TRUE | FALSE | TRUE |
| P27216 | P50995 | 0.0583 | | | FALSE | TRUE | FALSE | FALSE |
| Q09666 | Q8IVF2 | 0.0589 | | | FALSE | TRUE | FALSE | FALSE |
| P11142 | P54652 | 0.0591 | HOM | | TRUE | FALSE | FALSE | TRUE |
| P18085 | P36405 | 0.0596 | | | FALSE | TRUE | FALSE | FALSE |
| P08133 | P20073 | 0.0597 | | | FALSE | TRUE | FALSE | FALSE |
| O14979 | Q96DH6 | 0.0598 | | | FALSE | TRUE | FALSE | FALSE |
| Q07157 | Q9NZW5 | 0.0601 | HOM | HOM | TRUE | FALSE | FALSE | FALSE |
| Q96A00 | Q96C90 | 0.0609 | | | FALSE | TRUE | FALSE | FALSE |
| P43355 | P43358 | 0.062 | | | FALSE | TRUE | FALSE | FALSE |
| O94973 | O95782 | 0.0622 | HET | HET | FALSE | FALSE | FALSE | TRUE |
| P05091 | Q8IZ83 | 0.0645 | HOM | | TRUE | FALSE | FALSE | FALSE |
| P15153 | P63000 | 0.0647 | HET | HOM | TRUE | FALSE | FALSE | FALSE |
| P02008 | P69891 | 0.0649 | | HOM | TRUE | FALSE | FALSE | FALSE |
| P04083 | P08133 | 0.0653 | | | FALSE | TRUE | FALSE | FALSE |
| Q8IVD9 | Q9Y266 | 0.0659 | | | FALSE | TRUE | FALSE | FALSE |
| O15400 | Q86Y82 | 0.0663 | | | FALSE | TRUE | FALSE | TRUE |
| P07355 | Q5VT79 | 0.0672 | HOM | | TRUE | FALSE | FALSE | FALSE |
| P21291 | P52943 | 0.0677 | | | FALSE | TRUE | FALSE | FALSE |
| P51452 | Q5VZP5 | 0.0689 | HOM | | TRUE | FALSE | FALSE | FALSE |
| Q12959 | Q9NZW5 | 0.069 | HOM | HOM | TRUE | FALSE | FALSE | FALSE |
| O95486 | O95487 | 0.0696 | HET | HET | FALSE | FALSE | FALSE | TRUE |
| O60711 | Q14192 | 0.0697 | | | FALSE | TRUE | FALSE | FALSE |
| P29558 | Q13310 | 0.0705 | | | FALSE | TRUE | FALSE | FALSE |

Table S4 Human

| | | | | | | | | |
|---|---|---|---|---|---|---|---|---|
| P52597 | P55795 | 0.0708 | | | FALSE | TRUE | FALSE | TRUE |
| O60610 | Q27J81 | 0.0715 | HOM | | TRUE | FALSE | FALSE | FALSE |
| P52597 | Q9NTZ6 | 0.072 | | HOM | TRUE | FALSE | FALSE | FALSE |
| O60732 | P43363 | 0.0723 | | | FALSE | TRUE | FALSE | FALSE |
| P38919 | Q14240 | 0.0728 | HET | | FALSE | FALSE | FALSE | TRUE |
| P12111 | Q99715 | 0.0731 | | | FALSE | TRUE | FALSE | FALSE |
| P11940 | Q13310 | 0.0732 | HOM | | TRUE | FALSE | FALSE | FALSE |
| Q9NRR5 | Q9UMX0 | 0.0745 | HET | HOM, HET | TRUE, FALSE | FALSE | FALSE | TRUE |
| P51784 | Q9Y4E8 | 0.075 | | HOM | TRUE | FALSE | FALSE | TRUE |
| P15170 | Q9Y450 | 0.0763 | HET | | FALSE | FALSE | FALSE | FALSE |
| P04264 | P08729 | 0.0766 | HET | | FALSE | FALSE | FALSE | FALSE |
| P13797 | Q14651 | 0.0778 | | | FALSE | TRUE | FALSE | TRUE |
| P04083 | P50995 | 0.0778 | | | FALSE | TRUE | FALSE | FALSE |
| Q13177 | Q9UEW8 | 0.078 | | | FALSE | TRUE | FALSE | FALSE |
| O75323 | Q9BPW8 | 0.0782 | | | FALSE | TRUE | FALSE | FALSE |
| P53634 | Q9GZM7 | 0.0785 | HET | | FALSE | FALSE | FALSE | FALSE |
| P08758 | P12429 | 0.0788 | | | FALSE | TRUE | FALSE | FALSE |
| Q86V48 | Q9P2B4 | 0.079 | | HOM | TRUE | FALSE | FALSE | FALSE |
| P11021 | P54652 | 0.0802 | HOM | | TRUE | FALSE | FALSE | FALSE |
| P51610 | Q7Z6M1 | 0.0803 | | | FALSE | TRUE | FALSE | FALSE |
| P02461 | P20908 | 0.0808 | | | FALSE | TRUE | FALSE | FALSE |
| P05783 | P35900 | 0.0813 | | | FALSE | TRUE | FALSE | FALSE |
| O75607 | P06748 | 0.0813 | | | FALSE | TRUE | FALSE | TRUE |
| P68104 | Q05639 | 0.082 | HET | HET | FALSE | FALSE | FALSE | TRUE |
| Q5M775 | Q8NDI1 | 0.0836 | | | FALSE | TRUE | FALSE | FALSE |

Table S4 Human

| | | | | | | | | | |
|---|---|---|---|---|---|---|---|---|---|
| Q9NZW5 | Q9UDY2 | 0.0845 | HOM | HOM | TRUE | FALSE | FALSE | | FALSE |
| Q53H96 | Q96C36 | 0.0846 | HOM | HOM | TRUE | FALSE | FALSE | | FALSE |
| O60437 | Q15149 | 0.0851 | HET | HOM | TRUE | FALSE | FALSE | | FALSE |
| P04083 | P12429 | 0.0851 | | | FALSE | TRUE | FALSE | | FALSE |
| P08727 | P13645 | 0.0853 | | HET | FALSE | FALSE | FALSE | | FALSE |
| Q9BR76 | Q9ULV4 | 0.0857 | HOM | HOM | TRUE | FALSE | FALSE | | TRUE |
| Q9UHR4 | Q9UQB8 | 0.0858 | | HOM | TRUE | FALSE | FALSE | | TRUE |
| P08134 | P61586 | 0.0858 | | HOM | TRUE | FALSE | FALSE | | TRUE |
| P69891 | P69892 | 0.0859 | HOM | | TRUE | FALSE | FALSE | | FALSE |
| Q05707 | Q99715 | 0.0861 | | | FALSE | TRUE | FALSE | | FALSE |
| O60711 | P49023 | 0.087 | | | FALSE | TRUE | FALSE | | FALSE |
| P06239 | P07947 | 0.0876 | HOM | | TRUE | FALSE | FALSE | | FALSE |
| O00291 | Q9Y4G6 | 0.089 | HOM | | TRUE | FALSE | FALSE | | FALSE |
| P61225 | P62834 | 0.0897 | | HET | FALSE | FALSE | FALSE | | FALSE |
| P36404 | Q9NVJ2 | 0.0899 | HET | | FALSE | FALSE | FALSE | | FALSE |
| P40763 | P51692 | 0.0904 | HOM | HOM | TRUE | FALSE | FALSE | | TRUE |
| P36404 | P36405 | 0.0908 | HET | | FALSE | FALSE | FALSE | | TRUE |
| Q6P587 | Q96GK7 | 0.0915 | HOM | HOM | TRUE | FALSE | FALSE | | FALSE |
| Q00013 | Q5T2T1 | 0.0917 | HOM | HOM | TRUE | FALSE | FALSE | | FALSE |
| P31942 | Q6NXG1 | 0.0919 | | | FALSE | TRUE | FALSE | | FALSE |
| Q13617 | Q13620 | 0.092 | HET | HET | FALSE | FALSE | FALSE | | TRUE |
| P20591 | P50570 | 0.092 | HOM | | TRUE | FALSE | FALSE | | FALSE |
| P15924 | Q92817 | 0.093 | HOM | HOM | TRUE | FALSE | FALSE | | TRUE |
| Q96AE4 | Q96I24 | 0.0932 | HET | | FALSE | FALSE | FALSE | | TRUE |
| Q86X55 | Q99873 | 0.0934 | HOM | HOM | TRUE | FALSE | FALSE | | FALSE |
| P04083 | P20073 | 0.0939 | | | FALSE | TRUE | FALSE | | FALSE |

Table S4 Human

| | | | | | | | | | | | |
|---|---|---|---|---|---|---|---|---|---|---|---|
| P14550 | P52895 | 0.0941 | | HOM | TRUE | FALSE | FALSE | | | | FALSE |
| P00505 | P17174 | 0.0942 | HOM | HOM | TRUE | FALSE | FALSE | | | | FALSE |
| P15311 | P35241 | 0.0946 | | | FALSE | TRUE | FALSE | | | | TRUE |
| P35749 | Q7Z406 | 0.0947 | | | FALSE | TRUE | FALSE | | | | FALSE |
| O43432 | P78344 | 0.0948 | HET | HET | FALSE | FALSE | FALSE | | | | TRUE |
| P23528 | P60981 | 0.0952 | | | FALSE | TRUE | FALSE | | | | FALSE |
| P47895 | P49419 | 0.0959 | | HOM | TRUE | FALSE | FALSE | | | | FALSE |
| Q96B97 | Q9Y5K6 | 0.0973 | HOM | | TRUE | FALSE | TRUE | 846 | Human | RICH1/AMOT polarity complex, Flag-Rich1 precipitated | TRUE |
| Q14247 | Q16643 | 0.0983 | HOM | | TRUE | FALSE | FALSE | | | | TRUE |
| P62879 | Q9HAV0 | 0.0986 | | | FALSE | TRUE | FALSE | | | | FALSE |
| A1L0T0 | Q9UJ83 | 0.0987 | | | FALSE | TRUE | FALSE | | | | FALSE |
| P50238 | P52943 | 0.0987 | | | FALSE | TRUE | FALSE | | | | FALSE |
| P47895 | Q8IZ83 | 0.0988 | | | FALSE | TRUE | FALSE | | | | FALSE |
| Q6PCE3 | Q96G03 | 0.0989 | | | FALSE | TRUE | FALSE | | | | FALSE |
| P30048 | Q06830 | 0.0996 | HOM | HOM | TRUE | FALSE | FALSE | | | | TRUE |
| O60218 | P14550 | 0.1001 | | | FALSE | TRUE | FALSE | | | | FALSE |
| P15144 | Q9UIQ6 | 0.1008 | | | FALSE | TRUE | FALSE | | | | FALSE |
| P33240 | Q9H0L4 | 0.101 | HET | HET | FALSE | FALSE | FALSE | | | | TRUE |
| P13647 | P35908 | 0.101 | | | FALSE | TRUE | FALSE | | | | TRUE |
| P51911 | Q01995 | 0.1016 | | | FALSE | TRUE | FALSE | | | | FALSE |
| O95747 | Q9UEW8 | 0.1018 | HOM | | TRUE | FALSE | FALSE | | | | FALSE |
| P52292 | P52294 | 0.1021 | HET,HOM | HOM | TRUE | FALSE | FALSE | | | | TRUE |
| P55212 | Q14790 | 0.1023 | HOM | HET | TRUE | FALSE | FALSE | | | | TRUE |
| P43357 | P43358 | 0.1027 | | | FALSE | TRUE | FALSE | | | | FALSE |
| O75521 | Q13011 | 0.1032 | HOM | HOM | TRUE | FALSE | FALSE | | | | TRUE |

Table S4 Human

| | | | | | | | | |
|---|---|---|---|---|---|---|---|---|
| O00186 | Q15833 | 0.1037 | | | FALSE | TRUE | FALSE | FALSE |
| O95425 | P40121 | 0.1048 | | | FALSE | TRUE | FALSE | FALSE |
| O15479 | Q96MG7 | 0.1048 | | HET | FALSE | FALSE | FALSE | FALSE |
| Q9H5I1 | Q9H9B1 | 0.1053 | | HOM | TRUE | FALSE | FALSE | TRUE |
| O43795 | O94832 | 0.1057 | HOM | HOM | TRUE | FALSE | FALSE | FALSE |
| O60218 | P15121 | 0.1065 | | HOM | TRUE | FALSE | FALSE | FALSE |
| O43707 | Q13813 | 0.1067 | | HOM | TRUE | FALSE | FALSE | FALSE |
| P04264 | P35908 | 0.1069 | HET | | FALSE | FALSE | FALSE | TRUE |
| P11171 | Q9Y2J2 | 0.107 | | | FALSE | TRUE | FALSE | FALSE |
| Q32P28 | Q8IVL5 | 0.1086 | | | FALSE | TRUE | FALSE | FALSE |
| Q96DH6 | Q96EP5 | 0.1087 | | | FALSE | TRUE | FALSE | FALSE |
| Q8IXB1 | Q8NBS9 | 0.1087 | | | FALSE | TRUE | FALSE | FALSE |
| O94832 | Q12965 | 0.1087 | HOM | HOM | TRUE | FALSE | FALSE | FALSE |
| P00533 | Q05397 | 0.109 | HOM | HOM | TRUE | FALSE | FALSE | TRUE |
| P49588 | Q5JTZ9 | 0.1092 | | | FALSE | TRUE | FALSE | FALSE |
| P63000 | P84095 | 0.1092 | HOM | | TRUE | FALSE | FALSE | FALSE |
| P27144 | Q9UIJ7 | 0.111 | | | FALSE | TRUE | FALSE | FALSE |
| P43355 | Q9UNF1 | 0.1113 | | | FALSE | TRUE | FALSE | FALSE |
| P10644 | P31323 | 0.1118 | | | FALSE | TRUE | FALSE | TRUE |
| Q08257 | Q53FA7 | 0.1119 | HOM | HOM | TRUE | FALSE | FALSE | FALSE |
| P40939 | Q16836 | 0.1119 | | HOM | TRUE | FALSE | FALSE | FALSE |
| Q13268 | Q9BUT1 | 0.1137 | | HOM | TRUE | FALSE | FALSE | FALSE |
| P14625 | Q12931 | 0.1138 | HOM | HOM | TRUE | FALSE | FALSE | FALSE |
| P42765 | P55084 | 0.1139 | | HOM | TRUE | FALSE | FALSE | FALSE |
| P36507 | P46734 | 0.1146 | | | FALSE | TRUE | FALSE | FALSE |
| P20742 | Q6YHK3 | 0.1149 | | HET | FALSE | FALSE | FALSE | FALSE |

Table S4 Human

| | | | | | | | | | | | |
|---|---|---|---|---|---|---|---|---|---|---|---|
| O94880 | Q6IE81 | 0.1155 | | | FALSE | TRUE | FALSE | | | | FALSE |
| P07355 | P08758 | 0.1163 | HOM | | TRUE | FALSE | FALSE | | | | FALSE |
| Q13838 | Q9UHI6 | 0.1168 | HOM | | TRUE | FALSE | FALSE | | | | FALSE |
| Q13148 | Q96DH6 | 0.1172 | | | FALSE | TRUE | FALSE | | | | TRUE |
| P12111 | Q05707 | 0.1175 | | | FALSE | TRUE | FALSE | | | | FALSE |
| P20337 | P61026 | 0.118 | HOM | HOM | TRUE | FALSE | FALSE | | | | FALSE |
| P51159 | P61026 | 0.1185 | HET,HOM | HOM | TRUE | FALSE | FALSE | | | | FALSE |
| Q9H936 | Q9UJS0 | 0.119 | | | FALSE | TRUE | FALSE | | | | FALSE |
| P12814 | Q01082 | 0.1193 | | HOM | TRUE | FALSE | FALSE | | | | FALSE |
| P98179 | Q96E39 | 0.1194 | | | FALSE | TRUE | FALSE | | | | FALSE |
| O60437 | Q92817 | 0.1199 | HET | HOM | TRUE | FALSE | FALSE | | | | TRUE |
| Q96DI7 | Q9NVX2 | 0.1201 | | | FALSE | TRUE | FALSE | | | | FALSE |
| P52597 | Q12849 | 0.1206 | | | FALSE | TRUE | FALSE | | | | FALSE |
| Q9BUF5 | Q9BVA1 | 0.1214 | HET | HET | FALSE | FALSE | FALSE | | | | FALSE |
| P15144 | P55786 | 0.1217 | | | FALSE | TRUE | FALSE | | | | FALSE |
| Q9BY43 | Q9H444 | 0.1223 | | | FALSE | TRUE | TRUE | 1186 | Human | ESCRT-III complex | TRUE |
| P02749 | P15529 | 0.1223 | HET | | FALSE | FALSE | FALSE | | | | FALSE |
| P42566 | Q9UBC2 | 0.123 | HOM, HET | HET | TRUE, FALSE | FALSE | FALSE | | | | FALSE |
| Q16222 | Q3KQV9 | 0.1231 | HOM | | TRUE | FALSE | FALSE | | | | FALSE |
| P78362 | Q96SB4 | 0.1232 | HOM | HET,HOM | TRUE | FALSE | FALSE | | | | TRUE |
| Q07157 | Q5T2T1 | 0.1233 | HOM | HOM | TRUE | FALSE | FALSE | | | | FALSE |
| Q8NDI1 | Q8TDZ2 | 0.1234 | | | FALSE | TRUE | FALSE | | | | FALSE |
| O43447 | Q6UX04 | 0.1234 | HET | | FALSE | FALSE | FALSE | | | | FALSE |
| O00505 | O60684 | 0.1238 | HOM | HOM | TRUE | FALSE | FALSE | | | | TRUE |
| Q12792 | Q6IBS0 | 0.1244 | HOM | | TRUE | FALSE | FALSE | | | | FALSE |

Table S4 Human

| | | | | | | | | |
|---|---|---|---|---|---|---|---|---|
| O00505 | P52294 | 0.1244 | HOM | HOM | TRUE | FALSE | FALSE | FALSE |
| P14136 | Q03252 | 0.125 | | | FALSE | TRUE | FALSE | FALSE |
| O75718 | Q8IVL6 | 0.1251 | | | FALSE | TRUE | FALSE | FALSE |
| Q15063 | Q15582 | 0.1265 | | | FALSE | TRUE | FALSE | FALSE |
| P04264 | P05787 | 0.1266 | HET | | FALSE | FALSE | FALSE | FALSE |
| P31943 | Q12849 | 0.1271 | | | FALSE | TRUE | FALSE | FALSE |
| P07093 | P36952 | 0.1272 | | | FALSE | TRUE | FALSE | FALSE |
| Q99584 | Q9HCY8 | 0.1277 | HOM | | TRUE | FALSE | FALSE | FALSE |
| P61964 | Q96DI7 | 0.1287 | | | FALSE | TRUE | FALSE | TRUE |
| P11310 | Q92947 | 0.129 | HOM | HOM | TRUE | FALSE | FALSE | FALSE |
| Q9HC16 | Q9NRW3 | 0.1291 | HOM | | TRUE | FALSE | FALSE | FALSE |
| P26885 | Q96AY3 | 0.1292 | | | FALSE | TRUE | FALSE | FALSE |
| P07196 | P08670 | 0.1295 | | HOM | TRUE | FALSE | FALSE | TRUE |
| P02008 | P02100 | 0.1301 | | | FALSE | TRUE | FALSE | FALSE |
| O43815 | P53621 | 0.1305 | | HOM | TRUE | FALSE | FALSE | FALSE |
| P30048 | Q13162 | 0.1306 | HOM | HOM | TRUE | FALSE | FALSE | TRUE |
| Q9NWM8 | Q9Y680 | 0.1312 | | | FALSE | TRUE | FALSE | FALSE |
| P42574 | P55212 | 0.1318 | HOM | HOM | TRUE | FALSE | FALSE | TRUE |
| Q14195 | Q16555 | 0.1325 | HET | HOM | TRUE | FALSE | FALSE | TRUE |
| Q7Z739 | Q96MU7 | 0.1325 | | | FALSE | TRUE | FALSE | FALSE |
| O14936 | Q5T2T1 | 0.134 | HET,HOM | HOM | TRUE | FALSE | FALSE | FALSE |
| P19075 | P21926 | 0.1347 | | HOM | TRUE | FALSE | FALSE | FALSE |
| P55795 | Q6NXG1 | 0.1364 | | | FALSE | TRUE | FALSE | FALSE |
| P30101 | Q8IXB1 | 0.1366 | HET | | FALSE | FALSE | FALSE | FALSE |
| Q9NR46 | Q9Y371 | 0.1372 | HET | HET | FALSE | FALSE | FALSE | TRUE |
| P24752 | P55084 | 0.1373 | HOM | HOM | TRUE | FALSE | FALSE | FALSE |

Table S4 Human

| | | | | | | | | | | | |
|---|---|---|---|---|---|---|---|---|---|---|---|
| P61019 | Q15907 | 0.1383 | HET | HET | FALSE | FALSE | FALSE | | | | FALSE |
| P24386 | P31150 | 0.1387 | HET | HET | FALSE | FALSE | FALSE | | | | FALSE |
| O60711 | Q13642 | 0.1388 | | | FALSE | TRUE | FALSE | | | | FALSE |
| P07858 | Q9UBR2 | 0.1388 | HET | | FALSE | FALSE | FALSE | | | | FALSE |
| P02100 | P69891 | 0.1395 | | HOM | TRUE | FALSE | FALSE | | | | FALSE |
| O43143 | Q14562 | 0.1399 | | HOM | TRUE | FALSE | TRUE | 351 | Human | Spliceosome | FALSE |
| Q96NB2 | Q9BWM7 | 0.1408 | | | FALSE | TRUE | FALSE | | | | FALSE |
| P17858 | Q01813 | 0.1416 | HET | HET | FALSE | FALSE | FALSE | | | | TRUE |
| P14136 | P20700 | 0.1416 | | | FALSE | TRUE | FALSE | | | | FALSE |
| P84103 | Q16629 | 0.1423 | | | FALSE | TRUE | TRUE | 351 | Human | Spliceosome | TRUE |
| P08729 | P13647 | 0.1427 | | | FALSE | TRUE | FALSE | | | | FALSE |
| P19367 | P52789 | 0.1429 | HOM | | TRUE | FALSE | FALSE | | | | TRUE |
| P08238 | Q12931 | 0.143 | HOM | HOM | TRUE | FALSE | FALSE | | | | FALSE |
| P23368 | P48163 | 0.1438 | HOM | | TRUE | FALSE | FALSE | | | | FALSE |
| Q14562 | Q92620 | 0.1439 | HOM | | TRUE | FALSE | TRUE | 351,1181 | Human | Spliceosome,C complex spliceosome | FALSE |
| P49902 | Q5TFE4 | 0.1448 | HOM | | TRUE | FALSE | FALSE | | | | FALSE |
| Q12849 | Q8IXT5 | 0.145 | | | FALSE | TRUE | FALSE | | | | FALSE |
| P07332 | P41240 | 0.1454 | | | FALSE | TRUE | FALSE | | | | FALSE |
| Q5T2T1 | Q9UDY2 | 0.1457 | HOM | HOM | TRUE | FALSE | FALSE | | | | FALSE |
| O00299 | O95833 | 0.1463 | HOM | | TRUE | FALSE | FALSE | | | | FALSE |
| Q9NZI8 | Q9Y6M1 | 0.1469 | HOM | HOM | TRUE | FALSE | FALSE | | | | TRUE |
| P07093 | P50454 | 0.1469 | | | FALSE | TRUE | FALSE | | | | FALSE |
| P23297 | P31949 | 0.147 | HET | | FALSE | FALSE | FALSE | | | | FALSE |
| P60709 | Q562R1 | 0.1476 | HET | | FALSE | FALSE | FALSE | | | | FALSE |
| P13646 | P35527 | 0.1477 | | | FALSE | TRUE | FALSE | | | | FALSE |
| P07858 | Q9GZM7 | 0.1477 | HET | | FALSE | FALSE | FALSE | | | | FALSE |

Table S4 Human

| | | | | | | | | | | | |
|---|---|---|---|---|---|---|---|---|---|---|---|
| Q9H0U4 | Q9NP72 | 0.1483 | HET | HET | FALSE | FALSE | FALSE | | | | FALSE |
| P37802 | Q01995 | 0.1485 | | | FALSE | TRUE | FALSE | | | | FALSE |
| O15020 | Q13813 | 0.1487 | | HOM | TRUE | FALSE | FALSE | | | | TRUE |
| P13667 | P30101 | 0.1489 | | HET | FALSE | FALSE | FALSE | | | | TRUE |
| Q7L014 | Q9NR30 | 0.1493 | | HOM | TRUE | FALSE | FALSE | | | | FALSE |
| P49748 | Q9H845 | 0.1499 | HOM | | TRUE | FALSE | FALSE | | | | FALSE |
| P26599 | Q8WVV9 | 0.1503 | | | FALSE | TRUE | FALSE | | | | FALSE |
| P60709 | P61163 | 0.1509 | HET | | FALSE | FALSE | FALSE | | | | FALSE |
| P02008 | P69905 | 0.1519 | | HOM | TRUE | FALSE | FALSE | | | | FALSE |
| Q13617 | Q13619 | 0.1526 | HET | HOM, HET | TRUE, FALSE | FALSE | FALSE | | | | FALSE |
| Q96MU7 | Q9Y5A9 | 0.1528 | | | FALSE | TRUE | FALSE | | | | FALSE |
| P31146 | P57737 | 0.1533 | | | FALSE | TRUE | FALSE | | | | FALSE |
| O43852 | Q9BRK5 | 0.1534 | | | FALSE | TRUE | FALSE | | | | FALSE |
| Q96KQ7 | Q9H5I1 | 0.1539 | HET | | FALSE | FALSE | FALSE | | | | TRUE |
| P14866 | Q8WVV9 | 0.1543 | | | FALSE | TRUE | FALSE | | | | FALSE |
| P54136 | Q5T160 | 0.1545 | | | FALSE | TRUE | FALSE | | | | FALSE |
| Q14103 | Q96DH6 | 0.1548 | | | FALSE | TRUE | FALSE | | | | FALSE |
| Q07666 | Q15637 | 0.1556 | HOM | | TRUE | FALSE | FALSE | | | | FALSE |
| P61289 | Q9UL46 | 0.1577 | | | FALSE | TRUE | FALSE | | | | FALSE |
| P09651 | P22626 | 0.1582 | HOM | | TRUE | FALSE | TRUE | 1181 | Human | C complex spliceosome | TRUE |
| P06703 | P60903 | 0.1582 | HOM | | TRUE | FALSE | FALSE | | | | FALSE |
| P49023 | Q13642 | 0.1585 | | | FALSE | TRUE | FALSE | | | | FALSE |
| P42566 | Q9H4M9 | 0.1585 | HOM, HET | HET | TRUE, FALSE | FALSE | FALSE | | | | FALSE |
| Q13268 | Q7Z4W1 | 0.1593 | | HOM | TRUE | FALSE | FALSE | | | | FALSE |
| B0I1T2 | Q12965 | 0.1595 | HOM | HOM | TRUE | FALSE | FALSE | | | | FALSE |

Table S4 Human

| | | | | | | | | | | | |
|---|---|---|---|---|---|---|---|---|---|---|---|
| P19012 | Q04695 | 0.1597 | | | FALSE | TRUE | FALSE | | | | FALSE |
| O75475 | Q7Z4V5 | 0.1602 | | | FALSE | TRUE | FALSE | | | | FALSE |
| O15230 | P07942 | 0.1607 | | | FALSE | TRUE | TRUE | 2318 | Human | ITGA6-ITGB4-Laminin10/12 complex | FALSE |
| P18858 | P49916 | 0.1609 | | HOM | TRUE | FALSE | FALSE | | | | FALSE |
| Q02218 | Q96HY7 | 0.1612 | | | FALSE | TRUE | FALSE | | | | FALSE |
| Q00653 | Q04206 | 0.1614 | HOM | HET | TRUE | FALSE | TRUE | 5193,5194,5196,5230,5232,5233 | Human | TNF-alpha/NF-kappa B signaling complex,CHUK-NFKB2-REL-IKBKG-SPAG9-NFKB1-NFKBIE-COPB2-TNIP1-NFKBIA-RELA-TNIP2 complex,TNF-alpha/Nf-kappa B signaling complex,TNF-alpha/NF-kappa B signaling complex 5 | TRUE |
| Q7L014 | Q86XP3 | 0.1616 | | | FALSE | TRUE | FALSE | | | | FALSE |
| Q8IVL6 | Q92791 | 0.1618 | | | FALSE | TRUE | FALSE | | | | FALSE |
| Q86UP2 | Q9P2E9 | 0.1628 | HOM | | TRUE | FALSE | FALSE | | | | FALSE |
| P31947 | Q04917 | 0.1633 | HOM | | TRUE | FALSE | FALSE | | | | FALSE |
| P61163 | Q562R1 | 0.1635 | | | FALSE | TRUE | FALSE | | | | FALSE |
| O75367 | Q9P0M6 | 0.1637 | HOM | HET | TRUE | FALSE | FALSE | | | | TRUE |
| P08246 | P24158 | 0.1643 | HOM | HOM | TRUE | FALSE | FALSE | | | | FALSE |
| P39059 | P39060 | 0.1644 | HOM | HOM | TRUE | FALSE | FALSE | | | | FALSE |
| P05997 | P08123 | 0.1646 | | | FALSE | TRUE | FALSE | | | | FALSE |
| Q92900 | Q9HCE1 | 0.165 | HOM | | TRUE | FALSE | FALSE | | | | TRUE |
| P60981 | Q9Y281 | 0.1651 | | | FALSE | TRUE | FALSE | | | | TRUE |
| Q9BRQ6 | Q9NX63 | 0.1656 | | | FALSE | TRUE | TRUE | 6249,6255 | Human | MIB complex,MICOS complex | FALSE |
| P02100 | P69905 | 0.1659 | | HOM | TRUE | FALSE | FALSE | | | | FALSE |

Table S4 Human

| | | | | | | | | | | | |
|---|---|---|---|---|---|---|---|---|---|---|---|
| Q8IUE6 | Q9P0M6 | 0.1661 | HET | HET | FALSE | FALSE | FALSE | | | | TRUE |
| P54652 | Q0VDF9 | 0.1681 | | HET | FALSE | FALSE | FALSE | | | | FALSE |
| P04626 | Q05397 | 0.1689 | HOM | HOM | TRUE | FALSE | FALSE | | | | TRUE |
| Q00688 | Q13451 | 0.1692 | | HET | FALSE | FALSE | FALSE | | | | FALSE |
| Q12959 | Q5T2T1 | 0.171 | HOM | HOM | TRUE | FALSE | TRUE | 6260,6263 | Human | MPP7-DLG1-LIN7A complex,MPP7-DLG1-LIN7C complex | FALSE |
| O95425 | P06396 | 0.1722 | | | FALSE | TRUE | FALSE | | | | FALSE |
| O43447 | Q9Y3C6 | 0.1728 | HET | | FALSE | FALSE | TRUE | 351 | Human | Spliceosome | FALSE |
| P20339 | P61020 | 0.1729 | HOM | HOM | TRUE | FALSE | FALSE | | | | FALSE |
| P15153 | P84095 | 0.173 | HET | | FALSE | FALSE | FALSE | | | | FALSE |
| P48059 | Q14192 | 0.173 | HET | | FALSE | FALSE | FALSE | | | | FALSE |
| P31944 | P42574 | 0.1735 | | HOM | TRUE | FALSE | FALSE | | | | FALSE |
| O60784 | Q6ZVM7 | 0.1743 | | | FALSE | TRUE | FALSE | | | | FALSE |
| Q13188 | Q9P289 | 0.1744 | | | FALSE | TRUE | FALSE | | | | FALSE |
| Q7Z4W1 | Q99714 | 0.1744 | HOM | HOM | TRUE | FALSE | FALSE | | | | FALSE |
| P38159 | Q96E39 | 0.1748 | | | FALSE | TRUE | FALSE | | | | FALSE |
| P36507 | Q02750 | 0.1755 | | HOM | TRUE | FALSE | TRUE | 5872 | Human | BRAF-MAP2K1-MAP2K2-YWHAE complex | TRUE |
| O14936 | Q07157 | 0.1771 | HET,HOM | HOM | TRUE | FALSE | FALSE | | | | FALSE |
| P08133 | P08758 | 0.1778 | | | FALSE | TRUE | FALSE | | | | FALSE |
| Q32P28 | Q8IVL6 | 0.1781 | | | FALSE | TRUE | FALSE | | | | FALSE |
| P08123 | P20908 | 0.1786 | | | FALSE | TRUE | FALSE | | | | FALSE |
| Q53FA7 | Q99536 | 0.1791 | HOM | | TRUE | FALSE | FALSE | | | | FALSE |
| P48059 | Q9NR12 | 0.1791 | HET | | FALSE | FALSE | FALSE | | | | FALSE |
| P31943 | Q9NTZ6 | 0.1797 | | HOM | TRUE | FALSE | FALSE | | | | FALSE |
| P04899 | Q14344 | 0.1799 | | | FALSE | TRUE | FALSE | | | | FALSE |

Table S4 Human

| | | | | | | | | | | | |
|---|---|---|---|---|---|---|---|---|---|---|---|
| P57737 | Q9BR76 | 0.1801 | | HOM | TRUE | FALSE | FALSE | | | | FALSE |
| Q14247 | Q9UJU6 | 0.1806 | HOM | | TRUE | FALSE | FALSE | | | | FALSE |
| P30622 | Q14203 | 0.1807 | | HOM | TRUE | FALSE | FALSE | | | | TRUE |
| O43294 | P49023 | 0.181 | | | FALSE | TRUE | FALSE | | | | TRUE |
| P23284 | Q13356 | 0.1811 | | | FALSE | TRUE | FALSE | | | | FALSE |
| P24752 | P42765 | 0.1812 | HOM | | TRUE | FALSE | FALSE | | | | TRUE |
| O43854 | Q96PD2 | 0.1812 | | | FALSE | TRUE | FALSE | | | | FALSE |
| P04899 | P50148 | 0.1816 | | | FALSE | TRUE | FALSE | | | | FALSE |
| P35580 | Q7Z406 | 0.1817 | | | FALSE | TRUE | FALSE | | | | FALSE |
| O95302 | P26885 | 0.1831 | | | FALSE | TRUE | FALSE | | | | FALSE |
| Q13642 | Q96HC4 | 0.1834 | | | FALSE | TRUE | FALSE | | | | FALSE |
| O00148 | Q9UHI6 | 0.1839 | | | FALSE | TRUE | FALSE | | | | FALSE |
| Q13509 | Q9BVA1 | 0.1842 | HET | HET | FALSE | FALSE | FALSE | | | | TRUE |
| Q15084 | Q8IXB1 | 0.1845 | | | FALSE | TRUE | FALSE | | | | FALSE |
| Q12849 | Q9NTZ6 | 0.1848 | | HOM | TRUE | FALSE | FALSE | | | | FALSE |
| Q6UX04 | Q9Y3C6 | 0.1854 | | | FALSE | TRUE | TRUE | 1181 | Human | C complex spliceosome | FALSE |
| Q14257 | Q96D15 | 0.1855 | | | FALSE | TRUE | FALSE | | | | FALSE |
| P00352 | P05091 | 0.1868 | | HOM | TRUE | FALSE | FALSE | | | | TRUE |
| O75718 | Q8IVL5 | 0.1881 | | | FALSE | TRUE | FALSE | | | | FALSE |
| P08758 | P20073 | 0.1883 | | | FALSE | TRUE | FALSE | | | | FALSE |
| P23284 | P45877 | 0.19 | | | FALSE | TRUE | FALSE | | | | FALSE |
| P31942 | P52597 | 0.19 | | | FALSE | TRUE | FALSE | | | | TRUE |
| O15540 | P29373 | 0.19 | | HOM | TRUE | FALSE | FALSE | | | | FALSE |
| Q9UHD9 | Q9UMX0 | 0.1909 | HET | HOM, HET | TRUE, FALSE | FALSE | TRUE | 5209 | Human | Ubiquilin-proteasome complex | TRUE |
| P00352 | Q8IZ83 | 0.191 | | | FALSE | TRUE | FALSE | | | | FALSE |

Table S4 Human

| | | | | | | | | | | | |
|---|---|---|---|---|---|---|---|---|---|---|---|
| O43291 | P48307 | 0.1919 | | | FALSE | TRUE | FALSE | | | | FALSE |
| Q7Z739 | Q9Y5A9 | 0.1921 | | | FALSE | TRUE | FALSE | | | | FALSE |
| Q06323 | Q9UL46 | 0.1925 | HOM | | TRUE | FALSE | TRUE | 30,192,193 | Human | PA28 complex,PA28-20S proteasome,PA700-20S-PA28 complex | TRUE |
| P00533 | P04626 | 0.193 | HOM | HOM | TRUE | FALSE | TRUE | 1185 | Human | EGFR-containing signaling complex | TRUE |
| O60437 | P58107 | 0.1944 | HET | HET | FALSE | FALSE | FALSE | | | | FALSE |
| P55795 | Q12849 | 0.1948 | | | FALSE | TRUE | FALSE | | | | FALSE |
| O00505 | P52292 | 0.1951 | HOM | HET,HOM | TRUE | FALSE | FALSE | | | | TRUE |
| P26447 | P60903 | 0.1952 | | | FALSE | TRUE | FALSE | | | | FALSE |
| P07951 | P67936 | 0.1957 | | | FALSE | TRUE | FALSE | | | | TRUE |
| O95302 | Q9Y680 | 0.1961 | | | FALSE | TRUE | FALSE | | | | FALSE |
| P52597 | Q6NXG1 | 0.1967 | | | FALSE | TRUE | FALSE | | | | FALSE |
| Q8IX12 | Q8N163 | 0.1974 | | | FALSE | TRUE | FALSE | | | | FALSE |
| O75367 | Q8IUE6 | 0.1977 | HOM | HET | TRUE | FALSE | FALSE | | | | FALSE |
| P07910 | Q9UKM9 | 0.1978 | | | FALSE | TRUE | TRUE | 1181 | Human | C complex spliceosome | TRUE |
| O60732 | Q9UNF1 | 0.1983 | | | FALSE | TRUE | FALSE | | | | FALSE |
| P32322 | Q53H96 | 0.1986 | HOM | HOM | TRUE | FALSE | FALSE | | | | FALSE |
| P80303 | Q02818 | 0.1988 | | | FALSE | TRUE | FALSE | | | | FALSE |
| P14550 | P42330 | 0.2002 | | | FALSE | TRUE | FALSE | | | | FALSE |
| Q08211 | Q9H2U1 | 0.2013 | HOM | | TRUE | FALSE | FALSE | | | | FALSE |
| P06756 | P08648 | 0.2014 | HET | | FALSE | FALSE | FALSE | | | | FALSE |
| P30101 | Q8NBS9 | 0.2019 | HET | | FALSE | FALSE | FALSE | | | | FALSE |
| P57737 | Q9ULV4 | 0.2021 | | HOM | TRUE | FALSE | FALSE | | | | FALSE |
| P43363 | Q9UNF1 | 0.2021 | | | FALSE | TRUE | FALSE | | | | FALSE |
| O14936 | Q9NZW5 | 0.2024 | HET,HOM | HOM | TRUE | FALSE | FALSE | | | | FALSE |
| P48059 | Q13642 | 0.2033 | HET | | FALSE | FALSE | FALSE | | | | FALSE |

Table S4 Human

| | | | | | | | | | | | |
|---|---|---|---|---|---|---|---|---|---|---|---|
| P16104 | Q9P0M6 | 0.2042 | HET | HET | FALSE | FALSE | FALSE | | | | FALSE |
| Q12849 | Q6NXG1 | 0.2052 | | | FALSE | TRUE | FALSE | | | | FALSE |
| P13646 | P19012 | 0.2062 | | | FALSE | TRUE | FALSE | | | | FALSE |
| O00571 | Q9UJV9 | 0.2063 | | | FALSE | TRUE | TRUE | 351 | Human | Spliceosome | FALSE |
| O75340 | P04632 | 0.2064 | HOM | HET | TRUE | FALSE | FALSE | | | | FALSE |
| O15540 | P09455 | 0.2065 | | | FALSE | TRUE | FALSE | | | | FALSE |
| P36405 | P84085 | 0.2085 | | | FALSE | TRUE | FALSE | | | | FALSE |
| Q13620 | Q93034 | 0.2088 | HET | HET | FALSE | FALSE | FALSE | | | | TRUE |
| P31946 | P31947 | 0.2091 | | HOM | TRUE | FALSE | FALSE | | | | TRUE |
| P20339 | Q9UL25 | 0.2093 | HOM | HET | TRUE | FALSE | FALSE | | | | FALSE |
| P49023 | Q96HC4 | 0.21 | | | FALSE | TRUE | FALSE | | | | FALSE |
| Q9NQ29 | Q9Y383 | 0.2103 | | | FALSE | TRUE | FALSE | | | | TRUE |
| P15121 | P42330 | 0.2108 | HOM | | TRUE | FALSE | FALSE | | | | FALSE |
| P61981 | Q04917 | 0.211 | | | FALSE | TRUE | TRUE | 5199 | Human | Kinase maturation complex 1 | TRUE |

Table S4 Human

| | | | | | | | | | | | |
|---|---|---|---|---|---|---|---|---|---|---|---|
| Q09028 | Q16576 | 0.2112 | | | FALSE | TRUE | TRUE | 61,100,282,587,632,646,649,650,685,696,732,738,739,743,886,888,889,996,1257,2721,2851,2750 | Human,Mouse | Mi2/NuRD complex,hNURF complex,SNF2h-cohesin-NuRD complex,NuRD.1 complex,Anti-HDAC2 complex,HDAC1-associated protein complex,HDAC1-associated core complex cII,HDAC2-asscociated core complex,MeCP1 complex,BRMS1-SIN3-HDAC complex,SIN3 complex,SIN3-ING1b complex I,SIN3-ING1b complex II,SIN3-SAP25 complex,MTA1 complex,MTA2 complex,MTA1-HDAC core complex,Polycomb repressive complex 2,ALL-1 supercomplex,HCF-1 complex,ING2 complex,Gata1-Fog1- | TRUE |
| P52943 | Q16527 | 0.2118 | | | FALSE | TRUE | FALSE | | | | FALSE |
| P18206 | P35221 | 0.2121 | HET | HOM | TRUE | FALSE | FALSE | | | | TRUE |
| Q15149 | Q9UPN3 | 0.2135 | HOM | | TRUE | FALSE | FALSE | | | | FALSE |
| O60437 | P15924 | 0.2135 | HET | HOM | TRUE | FALSE | FALSE | | | | FALSE |
| Q96I99 | Q9P2R7 | 0.2137 | | | FALSE | TRUE | FALSE | | | | FALSE |
| P51159 | Q6IQ22 | 0.2141 | HET,HOM | HOM | TRUE | FALSE | FALSE | | | | FALSE |
| Q99470 | Q9HCN8 | 0.2142 | | | FALSE | TRUE | FALSE | | | | FALSE |
| P50552 | Q8N8S7 | 0.2148 | HOM | HET | TRUE | FALSE | FALSE | | | | TRUE |
| P42025 | Q562R1 | 0.2156 | | | FALSE | TRUE | FALSE | | | | FALSE |
| P15924 | Q15149 | 0.2161 | HOM | HOM | TRUE | FALSE | FALSE | | | | FALSE |
| O14936 | Q00013 | 0.2173 | HET,HOM | HOM | TRUE | FALSE | FALSE | | | | FALSE |
| P02008 | P69892 | 0.2174 | | | FALSE | TRUE | FALSE | | | | FALSE |

Table S4 Human

| | | | | | | | | | | | |
|---|---|---|---|---|---|---|---|---|---|---|---|
| Q15149 | Q92817 | 0.2185 | HOM | HOM | TRUE | FALSE | FALSE | | | | FALSE |
| P27797 | P27824 | 0.2194 | | | FALSE | TRUE | FALSE | | | | TRUE |
| P07711 | P09668 | 0.2196 | HOM | | TRUE | FALSE | FALSE | | | | TRUE |
| P08237 | P17858 | 0.2197 | HET | HET | FALSE | FALSE | FALSE | | | | TRUE |
| O00264 | O15173 | 0.2205 | HOM | | TRUE | FALSE | FALSE | | | | FALSE |
| O43707 | Q01082 | 0.2213 | | HOM | TRUE | FALSE | FALSE | | | | FALSE |
| O43237 | Q9Y6G 9 | 0.222 | HOM | HOM | TRUE | FALSE | FALSE | | | | TRUE |
| P21796 | P45880 | 0.2221 | | | FALSE | TRUE | FALSE | | | | TRUE |
| Q02790 | Q13451 | 0.2234 | HOM | HET | TRUE | FALSE | FALSE | | | | FALSE |
| P49023 | Q9NR1 2 | 0.2234 | | | FALSE | TRUE | FALSE | | | | FALSE |
| O75436 | Q4G0F 5 | 0.2235 | HET | HET | FALSE | FALSE | FALSE | | | | TRUE |
| O76041 | Q14847 | 0.2237 | | | FALSE | TRUE | FALSE | | | | FALSE |
| P05556 | P16144 | 0.225 | HOM | HET | TRUE | FALSE | FALSE | | | | FALSE |
| P43403 | Q05397 | 0.2251 | HOM | HOM | TRUE | FALSE | FALSE | | | | FALSE |
| P06493 | P24941 | 0.2256 | | HET | FALSE | FALSE | TRUE | 5559 | Human | CDC2-CCNA2-CDK2 complex | TRUE |
| Q15654 | Q93052 | 0.2288 | | | FALSE | TRUE | FALSE | | | | FALSE |
| P11940 | Q9UH X1 | 0.2311 | HOM | HOM | TRUE | FALSE | TRUE | 351 | Human | Spliceosome | TRUE |
| Q13310 | Q15717 | 0.2316 | | HOM | TRUE | FALSE | FALSE | | | | TRUE |
| O95292 | Q9P0L 0 | 0.2327 | HET | HET | FALSE | FALSE | FALSE | | | | TRUE |
| P60709 | P63261 | 0.233 | HET | | FALSE | FALSE | TRUE | 189,22 54 | Human | BAF complex,CTGF/Hcs24-actin complex | TRUE |
| P08727 | P13646 | 0.2341 | | | FALSE | TRUE | FALSE | | | | FALSE |
| P42566 | Q9H22 3 | 0.2348 | HOM, HET | HET | TRUE, FALSE | FALSE | FALSE | | | | FALSE |
| O60684 | P52294 | 0.2357 | HOM | HOM | TRUE | FALSE | FALSE | | | | FALSE |
| P50148 | Q14344 | 0.2361 | | | FALSE | TRUE | FALSE | | | | FALSE |
| O43447 | Q13356 | 0.2363 | HET | | FALSE | FALSE | TRUE | 351 | Human | Spliceosome | FALSE |

Table S4 Human

| | | | | | | | | |
|---|---|---|---|---|---|---|---|---|
| P31942 | Q9NTZ6 | 0.2365 | | HOM | TRUE | FALSE | FALSE | FALSE |
| P11021 | Q0VDF9 | 0.2372 | HOM | HET | TRUE | FALSE | FALSE | FALSE |
| P18085 | P36404 | 0.2382 | | HET | FALSE | FALSE | FALSE | FALSE |
| P24941 | Q00534 | 0.2393 | HET | HOM | TRUE | FALSE | FALSE | TRUE |
| O75494 | Q01130 | 0.2402 | | HOM | TRUE | FALSE | FALSE | FALSE |
| O15479 | O60732 | 0.2405 | | | FALSE | TRUE | FALSE | FALSE |
| Q9H3G5 | Q9HB40 | 0.2412 | | | FALSE | TRUE | FALSE | FALSE |
| P53634 | Q9UBR2 | 0.2414 | HET | | FALSE | FALSE | FALSE | FALSE |
| O14974 | Q8WUF5 | 0.2415 | | | FALSE | TRUE | FALSE | FALSE |
| O60831 | O75915 | 0.2426 | | | FALSE | TRUE | FALSE | FALSE |
| P13645 | P13646 | 0.2441 | HET | | FALSE | FALSE | FALSE | FALSE |
| P18206 | P26232 | 0.2442 | HET | | FALSE | FALSE | FALSE | FALSE |
| P04632 | Q9UBV8 | 0.2445 | HET | HET | FALSE | FALSE | FALSE | FALSE |
| P04899 | P63096 | 0.2456 | | HOM | TRUE | FALSE | FALSE | TRUE |
| O60256 | P60891 | 0.2461 | HOM | HOM | TRUE | FALSE | FALSE | TRUE |
| P06396 | P40121 | 0.2468 | | | FALSE | TRUE | FALSE | FALSE |
| O95757 | P34932 | 0.2474 | HOM | HOM | TRUE | FALSE | FALSE | FALSE |
| O75521 | P30084 | 0.2476 | HOM | HOM | TRUE | FALSE | FALSE | FALSE |
| O94804 | Q9Y6E0 | 0.248 | HOM | | TRUE | FALSE | FALSE | FALSE |
| Q00577 | Q96QR8 | 0.248 | HET | HET | FALSE | FALSE | FALSE | TRUE |
| P07942 | P55268 | 0.2482 | | | FALSE | TRUE | FALSE | FALSE |
| Q13642 | Q14192 | 0.2484 | | | FALSE | TRUE | FALSE | TRUE |
| P15170 | Q8IYD1 | 0.2494 | HET | | FALSE | FALSE | FALSE | TRUE |
| Q13595 | Q96E39 | 0.25 | | | FALSE | TRUE | FALSE | FALSE |
| O15479 | P43355 | 0.2501 | | | FALSE | TRUE | FALSE | FALSE |
| Q56VL3 | Q9NX40 | 0.251 | | | FALSE | TRUE | FALSE | FALSE |

Table S4 Human

| | | | | | | | | |
|---|---|---|---|---|---|---|---|---|
| O60732 | P43357 | 0.2517 | | | FALSE | TRUE | FALSE | FALSE |
| Q01995 | Q15417 | 0.2519 | | | FALSE | TRUE | FALSE | FALSE |
| P27348 | P31947 | 0.253 | | HOM | TRUE | FALSE | FALSE | TRUE |
| Q9BYJ9 | Q9Y5A9 | 0.2544 | | | FALSE | TRUE | FALSE | FALSE |
| P05121 | P50454 | 0.2546 | HOM | | TRUE | FALSE | FALSE | FALSE |
| P14317 | Q9UJU6 | 0.2547 | | | FALSE | TRUE | FALSE | FALSE |
| P08727 | P19012 | 0.2573 | | | FALSE | TRUE | FALSE | TRUE |
| P26885 | Q9NWM8 | 0.2581 | | | FALSE | TRUE | FALSE | FALSE |
| P36404 | P84085 | 0.2594 | HET | | FALSE | FALSE | FALSE | FALSE |
| Q86YP4 | Q8WXI9 | 0.2594 | HET | | FALSE | FALSE | FALSE | TRUE |
| P61964 | Q9NVX2 | 0.2608 | | | FALSE | TRUE | FALSE | TRUE |
| Q14011 | Q96E39 | 0.2611 | | | FALSE | TRUE | FALSE | FALSE |
| Q13425 | Q9NUP9 | 0.2623 | HOM | | TRUE | FALSE | FALSE | FALSE |
| Q05086 | Q5T447 | 0.2634 | HOM | | TRUE | FALSE | FALSE | FALSE |
| P48059 | P49023 | 0.2639 | HET | | FALSE | FALSE | FALSE | TRUE |
| P07196 | Q03252 | 0.265 | | | FALSE | TRUE | FALSE | FALSE |
| O94826 | Q15785 | 0.2651 | HOM | HOM | TRUE | FALSE | FALSE | FALSE |
| Q96CW1 | Q9BXS5 | 0.2657 | HET | HET | FALSE | FALSE | FALSE | FALSE |
| P07237 | Q8IXB1 | 0.2659 | HOM | | TRUE | FALSE | FALSE | FALSE |
| P51665 | Q7L5N1 | 0.2669 | HOM | | TRUE | FALSE | FALSE | FALSE |
| P29558 | Q15717 | 0.267 | | HOM | TRUE | FALSE | FALSE | FALSE |
| P20290 | Q96K17 | 0.2673 | HET | | FALSE | FALSE | FALSE | TRUE |
| P04899 | P08754 | 0.2674 | | HET | FALSE | FALSE | FALSE | TRUE |
| P43034 | Q9NVX2 | 0.2675 | HOM | | TRUE | FALSE | FALSE | FALSE |
| P00390 | Q16881 | 0.2702 | HOM | HOM | TRUE | FALSE | FALSE | FALSE |
| Q13642 | Q9NR12 | 0.2706 | | | FALSE | TRUE | FALSE | FALSE |

Table S4 Human

| | | | | | | | | | | | |
|---|---|---|---|---|---|---|---|---|---|---|---|
| Q15185 | Q9BTE6 | 0.2708 | | | FALSE | TRUE | FALSE | | | | FALSE |
| Q13595 | Q14011 | 0.271 | | | FALSE | TRUE | FALSE | | | | FALSE |
| O43852 | Q14257 | 0.2719 | | | FALSE | TRUE | FALSE | | | | FALSE |
| P36873 | P62136 | 0.2723 | | HOM | TRUE | FALSE | FALSE | | | | TRUE |
| O60488 | O95573 | 0.2725 | | | FALSE | TRUE | FALSE | | | | TRUE |
| P20700 | Q03252 | 0.2732 | | | FALSE | TRUE | FALSE | | | | TRUE |
| O00425 | Q9Y6M1 | 0.2734 | HOM | HOM | TRUE | FALSE | FALSE | | | | FALSE |
| P48729 | Q99986 | 0.2735 | | | FALSE | TRUE | FALSE | | | | FALSE |
| P67809 | Q9Y2T7 | 0.2746 | HOM | | TRUE | FALSE | FALSE | | | | FALSE |
| P62995 | Q14011 | 0.2764 | | | FALSE | TRUE | TRUE | 351 | Human | Spliceosome | FALSE |
| P56182 | Q14684 | 0.2774 | | | FALSE | TRUE | FALSE | | | | FALSE |
| Q13310 | Q9UHX1 | 0.2795 | | HOM | TRUE | FALSE | FALSE | | | | TRUE |
| P07951 | P09493 | 0.2811 | | | FALSE | TRUE | FALSE | | | | TRUE |
| P02461 | P05997 | 0.2833 | | | FALSE | TRUE | FALSE | | | | FALSE |
| P04271 | P23297 | 0.2844 | HOM | HET | TRUE | FALSE | FALSE | | | | TRUE |
| Q96RQ1 | Q9Y282 | 0.2844 | | | FALSE | TRUE | FALSE | | | | TRUE |
| P31942 | P31943 | 0.2845 | | | FALSE | TRUE | FALSE | | | | TRUE |
| O00425 | Q9NZI8 | 0.2846 | HOM | HOM | TRUE | FALSE | FALSE | | | | TRUE |
| P21796 | Q9Y277 | 0.2861 | | | FALSE | TRUE | FALSE | | | | TRUE |
| O43175 | P56545 | 0.2868 | HOM | HOM | TRUE | FALSE | FALSE | | | | FALSE |
| P30838 | P51648 | 0.2871 | | | FALSE | TRUE | FALSE | | | | TRUE |
| Q07866 | Q9H0B6 | 0.2885 | | HOM | TRUE | FALSE | FALSE | | | | TRUE |
| P26038 | P35241 | 0.2893 | | | FALSE | TRUE | FALSE | | | | FALSE |
| P69892 | P69905 | 0.291 | | HOM | TRUE | FALSE | FALSE | | | | FALSE |
| P49023 | Q14192 | 0.2919 | | | FALSE | TRUE | FALSE | | | | FALSE |
| O60732 | P43355 | 0.2931 | | | FALSE | TRUE | FALSE | | | | FALSE |

Table S4 Human

| | | | | | | | | | | | |
|---|---|---|---|---|---|---|---|---|---|---|---|
| Q8NBN7 | Q8TC12 | 0.2932 | | | FALSE | TRUE | FALSE | | | | FALSE |
| O15511 | Q9BPX5 | 0.2938 | | | FALSE | TRUE | FALSE | | | | TRUE |
| P37802 | Q15417 | 0.2941 | | | FALSE | TRUE | FALSE | | | | FALSE |
| P31942 | Q8IXT5 | 0.2964 | | | FALSE | TRUE | FALSE | | | | FALSE |
| Q14108 | Q8WTV0 | 0.2964 | | | FALSE | TRUE | FALSE | | | | FALSE |
| P60953 | P63000 | 0.2969 | HOM | HOM | TRUE | FALSE | FALSE | | | | FALSE |
| P21333 | Q14315 | 0.2972 | HOM | HOM | TRUE | FALSE | FALSE | | | | FALSE |
| P38159 | Q13595 | 0.2977 | | | FALSE | TRUE | FALSE | | | | FALSE |
| P27348 | Q04917 | 0.2978 | | | FALSE | TRUE | TRUE | 5199 | Human | Kinase maturation complex 1 | TRUE |
| P62942 | Q00688 | 0.2982 | HOM | | TRUE | FALSE | FALSE | | | | FALSE |
| Q00013 | Q9NZW5 | 0.3003 | HOM | HOM | TRUE | FALSE | FALSE | | | | FALSE |
| P31946 | Q04917 | 0.3009 | | | FALSE | TRUE | TRUE | 5199 | Human | Kinase maturation complex 1 | TRUE |
| P11047 | P55268 | 0.3009 | | | FALSE | TRUE | TRUE | 2319 | Human | ITGA6-ITGB4-Laminin10/12 complex | FALSE |
| O43294 | Q9NR12 | 0.3014 | | | FALSE | TRUE | FALSE | | | | FALSE |
| Q9H223 | Q9H4M9 | 0.3018 | HET | HET | FALSE | FALSE | FALSE | | | | TRUE |
| P42574 | Q14790 | 0.3026 | HOM | HET | TRUE | FALSE | FALSE | | | | TRUE |
| P08648 | P26006 | 0.3031 | | | FALSE | TRUE | FALSE | | | | FALSE |
| P05787 | P08729 | 0.3039 | | | FALSE | TRUE | FALSE | | | | FALSE |
| P05166 | Q9HCC0 | 0.3041 | | | FALSE | TRUE | FALSE | | | | FALSE |
| P06703 | P25815 | 0.3047 | HOM | HOM | TRUE | FALSE | FALSE | | | | FALSE |
| P30740 | P35237 | 0.305 | | | FALSE | TRUE | FALSE | | | | FALSE |
| P63244 | Q2TAY7 | 0.3051 | HOM | | TRUE | FALSE | FALSE | | | | FALSE |
| P55209 | Q99733 | 0.3055 | HOM | HOM | TRUE | FALSE | FALSE | | | | TRUE |
| P06732 | P12532 | 0.3056 | | HOM | TRUE | FALSE | FALSE | | | | FALSE |
| P25815 | P31949 | 0.3072 | HOM | | TRUE | FALSE | FALSE | | | | FALSE |

Table S4 Human

| | | | | | | | | | | | |
|---|---|---|---|---|---|---|---|---|---|---|---|
| P31943 | P52597 | 0.3091 | | | FALSE | TRUE | TRUE | 1181,5142 | Human,Mouse | C complex spliceosome,DCS complex | TRUE |
| P04632 | P30626 | 0.3095 | HET | | FALSE | FALSE | FALSE | | | | FALSE |
| P39023 | Q92901 | 0.3095 | | | FALSE | TRUE | FALSE | | | | FALSE |
| P24534 | P29692 | 0.3101 | HET | HET | FALSE | FALSE | FALSE | | | | TRUE |
| P05023 | P16615 | 0.3125 | | | FALSE | TRUE | FALSE | | | | FALSE |
| O43294 | Q14192 | 0.3133 | | | FALSE | TRUE | FALSE | | | | FALSE |
| P05997 | P20908 | 0.3134 | | | FALSE | TRUE | FALSE | | | | FALSE |
| Q12959 | Q9UDY2 | 0.3143 | HOM | HOM | TRUE | FALSE | FALSE | | | | FALSE |
| P62942 | Q02790 | 0.3144 | HOM | HOM | TRUE | FALSE | FALSE | | | | FALSE |
| P02461 | P08123 | 0.3146 | | | FALSE | TRUE | FALSE | | | | FALSE |
| P54577 | Q12904 | 0.3147 | HOM | HOM | TRUE | FALSE | FALSE | | | | FALSE |
| P50895 | Q13740 | 0.3153 | | | FALSE | TRUE | FALSE | | | | FALSE |
| P62873 | P62879 | 0.3159 | HET | | FALSE | FALSE | FALSE | | | | FALSE |
| O60716 | Q9Y446 | 0.3159 | | | FALSE | TRUE | FALSE | | | | FALSE |
| P05556 | P18084 | 0.3162 | HOM | | TRUE | FALSE | FALSE | | | | FALSE |
| P31946 | P63104 | 0.3165 | | | FALSE | TRUE | TRUE | 5199 | Human | Kinase maturation complex 1 | TRUE |
| P06703 | P26447 | 0.3166 | HOM | | TRUE | FALSE | FALSE | | | | TRUE |
| O00139 | Q99661 | 0.3169 | | | FALSE | TRUE | FALSE | | | | TRUE |
| P51148 | Q9UL25 | 0.3171 | | HET | FALSE | FALSE | FALSE | | | | FALSE |
| P57729 | Q13637 | 0.3185 | | | FALSE | TRUE | FALSE | | | | TRUE |
| P08670 | Q16352 | 0.3202 | HOM | | TRUE | FALSE | FALSE | | | | TRUE |
| P06756 | P26006 | 0.3207 | HET | | FALSE | FALSE | FALSE | | | | FALSE |
| P07355 | P12429 | 0.3209 | HOM | | TRUE | FALSE | FALSE | | | | FALSE |
| O43488 | Q13303 | 0.322 | | HET | FALSE | FALSE | FALSE | | | | FALSE |
| Q08170 | Q13247 | 0.3224 | HOM | HOM | TRUE | FALSE | TRUE | 351,2589 | Human | Spliceosome,PGC-1-SRp40-SRp55-SRp75 complex | TRUE |

Table S4 Human

| | | | | | | | | | | | |
|---|---|---|---|---|---|---|---|---|---|---|---|
| P18754 | Q9P258 | 0.3231 | HOM | | TRUE | FALSE | FALSE | | | | FALSE |
| P13667 | Q15084 | 0.3235 | | | FALSE | TRUE | FALSE | | | | FALSE |
| P07858 | P53634 | 0.3244 | HET | HET | FALSE | FALSE | FALSE | | | | FALSE |
| Q15286 | Q9NP72 | 0.3245 | HET | HET | FALSE | FALSE | FALSE | | | | FALSE |
| Q13619 | Q93034 | 0.3246 | HOM, HET | HET | TRUE, FALSE | FALSE | FALSE | | | | TRUE |
| P52565 | P52566 | 0.3261 | HET | HET | FALSE | FALSE | FALSE | | | | FALSE |
| O00469 | O60568 | 0.327 | | | FALSE | TRUE | FALSE | | | | FALSE |
| O43852 | Q96D15 | 0.327 | | | FALSE | TRUE | FALSE | | | | FALSE |
| P12429 | P20073 | 0.3287 | | | FALSE | TRUE | FALSE | | | | FALSE |
| Q96TA1 | Q9BZQ8 | 0.3289 | | | FALSE | TRUE | FALSE | | | | FALSE |
| P61019 | P61106 | 0.3289 | HET | HET | FALSE | FALSE | FALSE | | | | FALSE |
| P17844 | Q9NR30 | 0.3291 | | HOM | TRUE | FALSE | FALSE | | | | FALSE |
| Q13885 | Q9BVA1 | 0.3294 | HET | HET | FALSE | FALSE | FALSE | | | | FALSE |
| P63096 | Q14344 | 0.3296 | HOM | | TRUE | FALSE | FALSE | | | | TRUE |
| P06753 | P09493 | 0.3308 | | | FALSE | TRUE | FALSE | | | | TRUE |
| P45880 | Q9Y277 | 0.331 | | | FALSE | TRUE | FALSE | | | | TRUE |
| P31946 | P62258 | 0.3311 | | HOM | TRUE | FALSE | TRUE | 5199,5613 | Human | Kinase maturation complex 1,Emerin complex 25 | TRUE |
| P02749 | P08174 | 0.3314 | HET | HOM | TRUE | FALSE | FALSE | | | | FALSE |
| P41091 | Q2VIR3 | 0.3321 | HET | HET | FALSE | FALSE | FALSE | | | | FALSE |
| P20073 | P50995 | 0.3354 | | | FALSE | TRUE | FALSE | | | | FALSE |
| P30876 | Q9NW08 | 0.3358 | | | FALSE | TRUE | FALSE | | | | FALSE |
| P09455 | P29373 | 0.3359 | | HOM | TRUE | FALSE | FALSE | | | | FALSE |
| P11233 | P62070 | 0.336 | | | FALSE | TRUE | FALSE | | | | FALSE |
| P08754 | Q14344 | 0.3383 | HET | | FALSE | FALSE | FALSE | | | | FALSE |
| O43294 | Q96HC4 | 0.3388 | | | FALSE | TRUE | FALSE | | | | FALSE |

Table S4 Human

| | | | | | | | | |
|---|---|---|---|---|---|---|---|---|
| O14556 | P04406 | 0.3394 | HOM | HOM | TRUE | FALSE | FALSE | TRUE |
| P16989 | Q9Y2T7 | 0.3394 | | | FALSE | TRUE | FALSE | FALSE |
| P61026 | Q6IQ22 | 0.3408 | HOM | HOM | TRUE | FALSE | FALSE | FALSE |
| Q9H223 | Q9NZN4 | 0.341 | HET | HOM, HET | TRUE, FALSE | FALSE | FALSE | FALSE |
| P50579 | Q9UQ80 | 0.3413 | | | FALSE | TRUE | FALSE | FALSE |
| P08237 | Q01813 | 0.3414 | HET | HET | FALSE | FALSE | FALSE | TRUE |
| P68366 | Q9BQE3 | 0.3416 | HET | HET | FALSE | FALSE | FALSE | TRUE |
| Q8WVV9 | Q9UKA9 | 0.3425 | | | FALSE | TRUE | FALSE | FALSE |
| P04083 | P07355 | 0.3428 | | HOM | TRUE | FALSE | FALSE | FALSE |
| O14936 | Q9UDY2 | 0.343 | HET,HOM | HOM | TRUE | FALSE | FALSE | FALSE |
| P26447 | P31949 | 0.3435 | | | FALSE | TRUE | FALSE | FALSE |
| P43358 | Q9UNF1 | 0.3438 | | | FALSE | TRUE | FALSE | FALSE |
| P61020 | Q9UL25 | 0.3443 | HOM | HET | TRUE | FALSE | FALSE | FALSE |
| P31942 | Q12849 | 0.3449 | | | FALSE | TRUE | FALSE | FALSE |
| O00159 | O43795 | 0.3457 | HOM | HOM | TRUE | FALSE | FALSE | TRUE |
| P62995 | P98179 | 0.3464 | | | FALSE | TRUE | FALSE | FALSE |
| P42677 | Q71UM5 | 0.347 | | | FALSE | TRUE | FALSE | TRUE |
| O75368 | Q9H299 | 0.3472 | | HOM | TRUE | FALSE | FALSE | FALSE |
| P40967 | Q14956 | 0.3475 | HET | | FALSE | FALSE | FALSE | FALSE |
| P31483 | Q14498 | 0.3489 | | | FALSE | TRUE | FALSE | FALSE |
| P42330 | P52895 | 0.3491 | | HOM | TRUE | FALSE | FALSE | TRUE |
| P34932 | Q92598 | 0.3492 | HOM | HOM | TRUE | FALSE | FALSE | TRUE |
| P63261 | Q562R1 | 0.3506 | | | FALSE | TRUE | FALSE | TRUE |
| O95747 | Q13177 | 0.3506 | HOM | | TRUE | FALSE | FALSE | FALSE |
| O75367 | P16104 | 0.3515 | HOM | HET | TRUE | FALSE | FALSE | TRUE |

Table S4 Human

| | | | | | | | | | | | |
|---|---|---|---|---|---|---|---|---|---|---|---|
| Q9NZJ7 | Q9Y6C9 | 0.3518 | | | FALSE | TRUE | FALSE | | | | FALSE |
| P09960 | Q9H4A4 | 0.3523 | | | FALSE | TRUE | FALSE | | | | FALSE |
| P62942 | Q13451 | 0.3527 | HOM | HET | TRUE | FALSE | FALSE | | | | FALSE |
| Q9NRR5 | Q9UHD9 | 0.3527 | HET | HET | FALSE | FALSE | FALSE | | | | TRUE |
| P08754 | P50148 | 0.3531 | HET | | FALSE | FALSE | FALSE | | | | FALSE |
| O15230 | P11047 | 0.3534 | | | FALSE | TRUE | TRUE | 2318,2319 | Human | ITGA6-ITGB4-Laminin10/12 complex | TRUE |
| P29992 | P50148 | 0.3543 | | | FALSE | TRUE | TRUE | 117 | Human | GPR56-CD81-Galpha-Gbeta complex | FALSE |
| P23528 | Q9Y281 | 0.3547 | | | FALSE | TRUE | FALSE | | | | TRUE |
| P14923 | P35222 | 0.356 | | HET,HOM | FALSE,TRUE | FALSE | TRUE | 5177 | Human | Polycystin-1 multiprotein complex | TRUE |
| P06737 | P11216 | 0.3561 | HOM | HOM | TRUE | FALSE | FALSE | | | | TRUE |
| P20339 | P51148 | 0.3566 | HOM | | TRUE | FALSE | FALSE | | | | TRUE |
| P12109 | P12110 | 0.3574 | | | FALSE | TRUE | FALSE | | | | FALSE |
| P04264 | P13647 | 0.3575 | HET | | FALSE | FALSE | FALSE | | | | TRUE |
| O14618 | P00441 | 0.3576 | HOM | HOM | TRUE | FALSE | FALSE | | | | TRUE |
| P07384 | P17655 | 0.3588 | HET | HOM | TRUE | FALSE | FALSE | | | | TRUE |
| P09493 | P67936 | 0.359 | | | FALSE | TRUE | FALSE | | | | TRUE |
| P09758 | P16422 | 0.3596 | | | FALSE | TRUE | FALSE | | | | FALSE |
| P35579 | P35749 | 0.3599 | | | FALSE | TRUE | FALSE | | | | FALSE |
| Q16630 | Q8N684 | 0.3599 | HET | HET | FALSE | FALSE | TRUE | 1141 | Human | CF IIAm complex | TRUE |
| O75369 | P21333 | 0.3609 | | HOM | TRUE | FALSE | FALSE | | | | TRUE |
| Q15084 | Q8NBS9 | 0.3624 | | | FALSE | TRUE | FALSE | | | | FALSE |
| P06737 | P11217 | 0.3624 | HOM | HOM | TRUE | FALSE | FALSE | | | | FALSE |
| P51148 | P61020 | 0.3627 | | HOM | TRUE | FALSE | FALSE | | | | FALSE |
| P04899 | P29992 | 0.363 | | | FALSE | TRUE | FALSE | | | | FALSE |

Table S4 Human

| | | | | | | | | | | | |
|---|---|---|---|---|---|---|---|---|---|---|---|
| O60749 | Q13596 | 0.3639 | HOM | HOM | TRUE | FALSE | TRUE | 657,1060,1070,1091,1093,1095,1096,1104 | Human | Retromer complex,SNX complex | TRUE |
| Q01082 | Q13813 | 0.3641 | HOM | HOM | TRUE | FALSE | FALSE | | | | TRUE |
| P31943 | P55795 | 0.3658 | | | FALSE | TRUE | FALSE | | | | TRUE |
| P12429 | P50995 | 0.3684 | | | FALSE | TRUE | FALSE | | | | TRUE |
| Q9NYF8 | Q9Y2W1 | 0.3687 | | | FALSE | TRUE | FALSE | | | | TRUE |
| P16401 | P16403 | 0.3695 | | | FALSE | TRUE | FALSE | | | | FALSE |
| P30626 | Q9UBV8 | 0.3703 | | HET | FALSE | FALSE | FALSE | | | | FALSE |
| Q86XP3 | Q9NR30 | 0.3705 | | HOM | TRUE | FALSE | FALSE | | | | FALSE |
| Q01995 | Q99439 | 0.3709 | | | FALSE | TRUE | FALSE | | | | FALSE |
| P13667 | Q8NBS9 | 0.3717 | | | FALSE | TRUE | FALSE | | | | FALSE |
| Q12906 | Q96KR1 | 0.3717 | HET | | FALSE | FALSE | FALSE | | | | FALSE |
| O00469 | Q8NBJ5 | 0.372 | | | FALSE | TRUE | FALSE | | | | TRUE |
| Q14151 | Q15424 | 0.3722 | | HOM | TRUE | FALSE | FALSE | | | | TRUE |
| Q92643 | Q99538 | 0.373 | | | FALSE | TRUE | FALSE | | | | FALSE |
| O75369 | Q14315 | 0.3747 | | HOM | TRUE | FALSE | FALSE | | | | TRUE |
| Q14141 | Q9NVA2 | 0.3747 | HOM | HOM | TRUE | FALSE | FALSE | | | | FALSE |
| P63167 | Q96FJ2 | 0.375 | HOM | HOM | TRUE | FALSE | TRUE | 6236 | Human | Dynein-2 complex, cytoplasmic | TRUE |
| Q08170 | Q13243 | 0.3752 | HOM | HOM | TRUE | FALSE | TRUE | 351,2589 | Human | Spliceosome,PGC-1-SRp40-SRp55-SRp75 complex | FALSE |
| P22307 | P51659 | 0.3767 | HET | HOM | TRUE | FALSE | FALSE | | | | TRUE |
| P06732 | P12277 | 0.3789 | | | FALSE | TRUE | FALSE | | | | TRUE |
| Q6NZI2 | Q969G5 | 0.3797 | | | FALSE | TRUE | FALSE | | | | FALSE |
| Q96DB5 | Q96TC7 | 0.3802 | | | FALSE | TRUE | FALSE | | | | FALSE |
| Q13188 | Q9Y6E0 | 0.3812 | | | FALSE | TRUE | FALSE | | | | FALSE |

Table S4 Human

| | | | | | | | | | | | |
|---|---|---|---|---|---|---|---|---|---|---|---|
| P54725 | P54727 | 0.3818 | HOM, HET | HET | TRUE, FALSE | FALSE | FALSE | | | | TRUE |
| P56545 | Q13363 | 0.3831 | HOM | HET | TRUE | FALSE | TRUE | 642,2921 | Human | CtBP complex,SHARP-CtBP complex | TRUE |
| O00303 | Q7L5N1 | 0.3836 | | | FALSE | TRUE | FALSE | | | | FALSE |
| P43355 | Q96MG7 | 0.3837 | | HET | FALSE | FALSE | FALSE | | | | FALSE |
| P25325 | Q16762 | 0.3839 | | | FALSE | TRUE | FALSE | | | | FALSE |
| Q96AY3 | Q9Y680 | 0.3841 | | | FALSE | TRUE | FALSE | | | | FALSE |
| P29992 | P63096 | 0.3859 | | HOM | TRUE | FALSE | FALSE | | | | FALSE |
| O60716 | Q99959 | 0.3883 | | | FALSE | TRUE | FALSE | | | | FALSE |
| P48594 | P50453 | 0.3883 | | | FALSE | TRUE | FALSE | | | | FALSE |
| P18085 | Q9NVJ2 | 0.3884 | | | FALSE | TRUE | FALSE | | | | FALSE |
| P62995 | Q96E39 | 0.389 | | | FALSE | TRUE | FALSE | | | | FALSE |
| P13995 | Q6UB35 | 0.3896 | | | FALSE | TRUE | FALSE | | | | FALSE |
| O14936 | Q12959 | 0.3909 | HET,HOM | HOM | TRUE | FALSE | TRUE | 3207 | Human | LIN2-LIN7-SAP97-MINT1 complex | TRUE |
| Q96HC4 | Q9NR12 | 0.3909 | | | FALSE | TRUE | FALSE | | | | TRUE |
| P01892 | P17693 | 0.3916 | HET | HOM, HET | TRUE, FALSE | FALSE | FALSE | | | | TRUE |
| P49257 | Q12907 | 0.3939 | HOM | | TRUE | FALSE | FALSE | | | | FALSE |
| O43447 | P23284 | 0.3964 | HET | | FALSE | FALSE | FALSE | | | | FALSE |
| P50148 | P63096 | 0.3969 | | HOM | TRUE | FALSE | FALSE | | | | FALSE |
| Q93008 | Q93009 | 0.3983 | | HET | FALSE | FALSE | FALSE | | | | FALSE |
| P10253 | Q14697 | 0.3983 | | | FALSE | TRUE | FALSE | | | | FALSE |
| P12111 | Q02388 | 0.4004 | | | FALSE | TRUE | FALSE | | | | FALSE |
| Q13443 | Q9UKF2 | 0.4009 | | | FALSE | TRUE | FALSE | | | | FALSE |
| Q8IYD1 | Q9Y450 | 0.403 | | | FALSE | TRUE | FALSE | | | | FALSE |
| Q9C0C9 | Q9H832 | 0.4031 | | | FALSE | TRUE | FALSE | | | | FALSE |

Table S4 Human

| | | | | | | | | | | | |
|---|---|---|---|---|---|---|---|---|---|---|---|
| Q9P289 | Q9Y6E0 | 0.4075 | | | FALSE | TRUE | FALSE | | | | TRUE |
| P25815 | P26447 | 0.4088 | HOM | | TRUE | FALSE | FALSE | | | | FALSE |
| P08729 | Q5XKE5 | 0.409 | | | FALSE | TRUE | FALSE | | | | FALSE |
| P08729 | Q7Z794 | 0.4091 | | | FALSE | TRUE | FALSE | | | | FALSE |
| Q12905 | Q96KR1 | 0.4092 | HET | | FALSE | FALSE | FALSE | | | | TRUE |
| P05023 | P98194 | 0.4101 | | | FALSE | TRUE | FALSE | | | | FALSE |
| O60711 | P48059 | 0.4115 | | HET | FALSE | FALSE | FALSE | | | | TRUE |
| Q9UBF2 | Q9Y678 | 0.4115 | | | FALSE | TRUE | FALSE | | | | TRUE |
| Q07157 | Q9UDY2 | 0.4118 | HOM | HOM | TRUE | FALSE | FALSE | | | | TRUE |
| P57723 | Q15366 | 0.4125 | | HOM | TRUE | FALSE | FALSE | | | | FALSE |
| P10412 | P16401 | 0.413 | | | FALSE | TRUE | FALSE | | | | FALSE |
| P62873 | Q9HAV0 | 0.4133 | HET | | FALSE | FALSE | FALSE | | | | FALSE |
| P42224 | P42226 | 0.4144 | HOM | HOM | TRUE | FALSE | FALSE | | | | FALSE |
| Q13885 | Q9BUF5 | 0.4144 | HET | HET | FALSE | FALSE | FALSE | | | | FALSE |
| P35611 | Q9UEY8 | 0.415 | | | FALSE | TRUE | FALSE | | | | TRUE |
| O43252 | O95340 | 0.4155 | HOM | | TRUE | FALSE | FALSE | | | | TRUE |
| Q15785 | Q99614 | 0.4185 | HOM | HOM | TRUE | FALSE | FALSE | | | | FALSE |
| Q15286 | Q9H0U4 | 0.4217 | HET | HET | FALSE | FALSE | FALSE | | | | FALSE |
| O00303 | P51665 | 0.4225 | | HOM | TRUE | FALSE | FALSE | | | | FALSE |
| Q8NE71 | Q9NUQ8 | 0.4239 | | | FALSE | TRUE | FALSE | | | | FALSE |
| P00338 | P07195 | 0.4265 | | | FALSE | TRUE | FALSE | | | | TRUE |
| P08174 | P15529 | 0.4271 | HOM | | TRUE | FALSE | FALSE | | | | FALSE |
| Q92947 | Q9H845 | 0.4275 | HOM | | TRUE | FALSE | FALSE | | | | FALSE |
| P98179 | Q13595 | 0.4278 | | | FALSE | TRUE | FALSE | | | | FALSE |
| Q92888 | Q92974 | 0.4281 | | | FALSE | TRUE | TRUE | 6671 | Human | PI4K2A-WASH complex | FALSE |
| Q08752 | Q13427 | 0.4282 | HOM | | TRUE | FALSE | FALSE | | | | FALSE |

Table S4 Human

| | | | | | | | | | | | |
|---|---|---|---|---|---|---|---|---|---|---|---|
| P30084 | Q13011 | 0.4283 | HOM | HOM | TRUE | FALSE | FALSE | | | | FALSE |
| Q14192 | Q96HC4 | 0.4302 | | | FALSE | TRUE | FALSE | | | | FALSE |
| Q9H4A3 | Q9UHY1 | 0.4305 | | | FALSE | TRUE | FALSE | | | | FALSE |
| P45973 | Q13185 | 0.4323 | HOM | HOM | TRUE | FALSE | FALSE | | | | TRUE |
| O43823 | Q5BKZ1 | 0.4361 | | | FALSE | TRUE | FALSE | | | | FALSE |
| P06493 | Q00534 | 0.4366 | | HOM | TRUE | FALSE | FALSE | | | | FALSE |
| Q00839 | Q92499 | 0.4382 | | | FALSE | TRUE | TRUE | 1332 | Human | Large Drosha complex | TRUE |
| P43358 | P43363 | 0.4387 | | | FALSE | TRUE | FALSE | | | | TRUE |
| P23588 | Q15056 | 0.4397 | | | FALSE | TRUE | FALSE | | | | FALSE |
| O95302 | Q9NWM8 | 0.4408 | | | FALSE | TRUE | FALSE | | | | FALSE |
| P17844 | Q86XP3 | 0.4408 | | | FALSE | TRUE | FALSE | | | | FALSE |
| Q15365 | Q15366 | 0.4419 | | HOM | TRUE | FALSE | FALSE | | | | TRUE |
| P29992 | Q14344 | 0.4466 | | | FALSE | TRUE | FALSE | | | | FALSE |
| O15347 | P09429 | 0.4494 | | | FALSE | TRUE | FALSE | | | | FALSE |
| P06703 | P31949 | 0.45 | HOM | | TRUE | FALSE | FALSE | | | | FALSE |
| Q13509 | Q9BUF5 | 0.4516 | HET | HET | FALSE | FALSE | FALSE | | | | FALSE |
| Q08170 | Q13242 | 0.457 | HOM | HOM | TRUE | FALSE | TRUE | 351 | Human | Spliceosome | FALSE |
| P11279 | P13473 | 0.4576 | | HOM | TRUE | FALSE | FALSE | | | | FALSE |
| Q99614 | Q9H3U1 | 0.4586 | HOM | HOM | TRUE | FALSE | FALSE | | | | FALSE |
| O15347 | P26583 | 0.459 | | | FALSE | TRUE | FALSE | | | | FALSE |
| Q15942 | Q93052 | 0.46 | | | FALSE | TRUE | FALSE | | | | FALSE |
| Q15417 | Q99439 | 0.4659 | | | FALSE | TRUE | FALSE | | | | TRUE |
| O94804 | Q9P289 | 0.4664 | HOM | | TRUE | FALSE | FALSE | | | | FALSE |
| Q13509 | Q13885 | 0.4673 | HET | HET | FALSE | FALSE | FALSE | | | | TRUE |
| P07237 | Q8NBS9 | 0.4689 | HOM | | TRUE | FALSE | FALSE | | | | FALSE |
| P39687 | Q92688 | 0.4692 | | | FALSE | TRUE | FALSE | | | | TRUE |

Table S4 Human

| | | | | | | | | | | | |
|---|---|---|---|---|---|---|---|---|---|---|---|
| P62995 | Q13595 | 0.4714 | | | FALSE | TRUE | FALSE | | | | TRUE |
| P07942 | P11047 | 0.4722 | | | FALSE | TRUE | TRUE | 2318 | Human | ITGA6-ITGB4-Laminin10/12 complex | TRUE |
| O00469 | Q02809 | 0.4725 | | | FALSE | TRUE | FALSE | | | | TRUE |
| O43707 | P12814 | 0.4803 | | | FALSE | TRUE | FALSE | | | | TRUE |
| P13645 | P35527 | 0.4812 | HET | | FALSE | FALSE | FALSE | | | | FALSE |
| P07237 | P30101 | 0.4845 | HOM | HET | TRUE | FALSE | FALSE | | | | TRUE |
| P31949 | P60903 | 0.4852 | | | FALSE | TRUE | FALSE | | | | FALSE |
| P16219 | P49748 | 0.4853 | HOM | HOM | TRUE | FALSE | FALSE | | | | FALSE |
| O43852 | Q15293 | 0.4857 | | | FALSE | TRUE | FALSE | | | | FALSE |
| Q13619 | Q13620 | 0.4909 | HOM, HET | HET | TRUE, FALSE | FALSE | TRUE | 1162 | Human | Ubiquitin E3 ligase | TRUE |
| P68431 | P84243 | 0.494 | HET | HET | FALSE | FALSE | FALSE | | | | FALSE |
| O43172 | Q9UMS4 | 0.494 | HET | HOM | TRUE | FALSE | TRUE | 351 | Human | Spliceosome | TRUE |
| P16615 | P98194 | 0.4947 | | | FALSE | TRUE | FALSE | | | | FALSE |
| Q8N1G4 | Q9NSD9 | 0.4951 | | HOM | TRUE | FALSE | FALSE | | | | FALSE |
| Q96IJ6 | Q9Y5P6 | 0.4953 | | | FALSE | TRUE | FALSE | | | | TRUE |
| O14979 | Q96EP5 | 0.4966 | | | FALSE | TRUE | FALSE | | | | FALSE |
| P35637 | Q01844 | 0.4997 | | | FALSE | TRUE | TRUE | 1332 | Human | Large Drosha complex | TRUE |
| Q92688 | Q9BTT0 | 0.5003 | | | FALSE | TRUE | FALSE | | | | FALSE |
| P39687 | Q9BTT0 | 0.5053 | | | FALSE | TRUE | FALSE | | | | TRUE |
| Q15785 | Q9H3U1 | 0.5068 | HOM | HOM | TRUE | FALSE | FALSE | | | | FALSE |
| P32119 | Q06830 | 0.5069 | HOM | HOM | TRUE | FALSE | FALSE | | | | TRUE |
| Q14192 | Q9NR12 | 0.5102 | | | FALSE | TRUE | FALSE | | | | FALSE |
| P57721 | Q15366 | 0.5103 | | HOM | TRUE | FALSE | FALSE | | | | FALSE |
| P68402 | Q15102 | 0.5137 | HOM | | TRUE | FALSE | FALSE | | | | TRUE |
| P22695 | Q10713 | 0.5142 | HET | HET | FALSE | FALSE | FALSE | | | | FALSE |

Table S4 Human

| | | | | | | | | | | | |
|---|---|---|---|---|---|---|---|---|---|---|---|
| P15924 | P58107 | 0.5175 | HOM | HET | TRUE | FALSE | FALSE | | | | FALSE |
| Q32P28 | Q92791 | 0.518 | | | FALSE | TRUE | FALSE | | | | FALSE |
| Q99959 | Q9Y446 | 0.5191 | | | FALSE | TRUE | FALSE | | | | FALSE |
| P98179 | Q14011 | 0.5217 | | | FALSE | TRUE | FALSE | | | | FALSE |
| Q07157 | Q12959 | 0.5225 | HOM | HOM | TRUE | FALSE | FALSE | | | | FALSE |
| P45973 | P83916 | 0.5229 | HOM | HOM, HET | TRUE | FALSE | FALSE | | | | TRUE |
| P08754 | P29992 | 0.5232 | HET | | FALSE | FALSE | FALSE | | | | FALSE |
| P51149 | P57729 | 0.5245 | HOM | | TRUE | FALSE | FALSE | | | | FALSE |
| P57721 | P57723 | 0.5266 | | | FALSE | TRUE | FALSE | | | | FALSE |
| P51149 | Q13637 | 0.5267 | HOM | | TRUE | FALSE | FALSE | | | | FALSE |
| P27348 | P61981 | 0.5297 | | | FALSE | TRUE | TRUE | 5199 | Human | Kinase maturation complex 1 | TRUE |
| Q15654 | Q15942 | 0.5303 | | | FALSE | TRUE | FALSE | | | | FALSE |
| P07237 | Q15084 | 0.5397 | HOM | | TRUE | FALSE | FALSE | | | | TRUE |
| O75439 | P31930 | 0.5398 | HET | HET | FALSE | FALSE | FALSE | | | | FALSE |
| Q15717 | Q9UHX1 | 0.5405 | HOM | HOM | TRUE | FALSE | FALSE | | | | FALSE |
| O94842 | Q08945 | 0.5448 | HOM | HET,HOM | TRUE | FALSE | FALSE | | | | FALSE |
| O75718 | Q92791 | 0.5451 | | | FALSE | TRUE | FALSE | | | | FALSE |
| Q9NR31 | Q9Y6B6 | 0.5476 | HOM | HOM | TRUE | FALSE | FALSE | | | | TRUE |
| O95810 | Q6NZI2 | 0.5483 | | | FALSE | TRUE | FALSE | | | | FALSE |
| P31150 | P50395 | 0.5498 | HET | HET | FALSE | FALSE | FALSE | | | | TRUE |
| O43399 | P55327 | 0.5518 | HOM | HOM | TRUE | FALSE | FALSE | | | | TRUE |
| Q96AY3 | Q9NWM8 | 0.5533 | | | FALSE | TRUE | FALSE | | | | FALSE |
| P14866 | Q9UKA9 | 0.5546 | | | FALSE | TRUE | FALSE | | | | TRUE |
| P57721 | Q15365 | 0.5553 | | | FALSE | TRUE | FALSE | | | | TRUE |
| Q13263 | Q9UPN9 | 0.5553 | HOM | HET | TRUE | FALSE | FALSE | | | | FALSE |
| O60220 | Q9Y5J9 | 0.5562 | HET | HET | FALSE | FALSE | FALSE | | | | FALSE |

Table S4 Human

| | | | | | | | | | | | |
|---|---|---|---|---|---|---|---|---|---|---|---|
| P37802 | Q99439 | 0.5609 | | | FALSE | TRUE | FALSE | | | | FALSE |
| O00148 | Q13838 | 0.5621 | | HOM | TRUE | FALSE | FALSE | | | | TRUE |
| Q15019 | Q9UH03 | 0.564 | HOM | HET | TRUE | FALSE | FALSE | | | | FALSE |
| P62328 | P63313 | 0.5664 | | | FALSE | TRUE | FALSE | | | | FALSE |
| O60684 | P52292 | 0.5665 | HOM | HET,HOM | TRUE | FALSE | FALSE | | | | TRUE |
| P05141 | P12236 | 0.5674 | | | FALSE | TRUE | FALSE | | | | FALSE |
| P17844 | Q92841 | 0.5773 | | | FALSE | TRUE | TRUE | 351,1332,3082 | Human | Spliceosome,Large Drosha complex,DGCR8 multiprotein complex | TRUE |
| Q1KMD3 | Q92499 | 0.5791 | | | FALSE | TRUE | FALSE | | | | FALSE |
| Q14257 | Q15293 | 0.5802 | | | FALSE | TRUE | FALSE | | | | FALSE |
| O14979 | Q14103 | 0.5825 | | | FALSE | TRUE | FALSE | | | | TRUE |
| Q13148 | Q14103 | 0.5833 | | | FALSE | TRUE | FALSE | | | | TRUE |
| P02545 | Q16352 | 0.5841 | HOM | | TRUE | FALSE | FALSE | | | | FALSE |
| Q9H4M9 | Q9NZN4 | 0.5842 | HET | HOM, HET | TRUE, FALSE | FALSE | FALSE | | | | FALSE |
| P32322 | Q96C36 | 0.5866 | HOM | HOM | TRUE | FALSE | FALSE | | | | TRUE |
| P36915 | Q9H089 | 0.5874 | | | FALSE | TRUE | FALSE | | | | FALSE |
| O60884 | P31689 | 0.5916 | HOM | HOM | TRUE | FALSE | FALSE | | | | FALSE |
| O43837 | P51553 | 0.5927 | HET,HOM | HET,HOM | FALSE,TRUE | FALSE | FALSE | | | | TRUE |
| P57723 | Q15365 | 0.5956 | | | FALSE | TRUE | FALSE | | | | TRUE |
| O43684 | P78406 | 0.5975 | | HET | FALSE | FALSE | FALSE | | | | FALSE |
| O95045 | Q16831 | 0.6056 | HOM | HOM | TRUE | FALSE | FALSE | | | | FALSE |
| P05787 | Q7Z794 | 0.6086 | | | FALSE | TRUE | FALSE | | | | FALSE |
| Q00688 | Q02790 | 0.612 | | HOM | TRUE | FALSE | FALSE | | | | FALSE |
| Q96KQ7 | Q9H9B1 | 0.6139 | HET | HOM | TRUE | FALSE | TRUE | 642,1194 | Human | CtBP complex,E2F-6 complex | TRUE |
| Q13148 | Q96EP5 | 0.6147 | | | FALSE | TRUE | FALSE | | | | FALSE |

Table S4 Human

| | | | | | | | | | | | |
|---|---|---|---|---|---|---|---|---|---|---|---|
| P46926 | Q8TDQ7 | 0.6226 | HOM | | TRUE | FALSE | FALSE | | | | TRUE |
| P07237 | P13667 | 0.6232 | HOM | | TRUE | FALSE | FALSE | | | | TRUE |
| O60568 | Q02809 | 0.6248 | | | FALSE | TRUE | FALSE | | | | FALSE |
| P62306 | P62312 | 0.6271 | HOM | HOM | TRUE | FALSE | TRUE | 351 | Human | Spliceosome | TRUE |
| P78344 | Q04637 | 0.6272 | HET | HOM, HET | TRUE, FALSE | FALSE | FALSE | | | | TRUE |
| P11142 | P34931 | 0.631 | HOM | | TRUE | FALSE | FALSE | | | | FALSE |
| P09429 | P26583 | 0.6386 | | | FALSE | TRUE | TRUE | 280 | Human | HMGB1-HMGB2-HSC70-ERP60-GAPDH complex | TRUE |
| P26599 | Q9UKA9 | 0.6421 | | | FALSE | TRUE | FALSE | | | | FALSE |
| A5A3E0 | P0CG39 | 0.6545 | | | FALSE | TRUE | FALSE | | | | FALSE |
| P83916 | Q13185 | 0.6552 | HOM, HET | HOM | TRUE | FALSE | FALSE | | | | TRUE |
| Q15181 | Q9H2U2 | 0.6593 | | | FALSE | TRUE | FALSE | | | | FALSE |
| O95232 | Q9Y383 | 0.6715 | HOM | | TRUE | FALSE | FALSE | | | | TRUE |
| Q02809 | Q8NBJ5 | 0.6736 | | | FALSE | TRUE | FALSE | | | | TRUE |
| P05787 | Q5XKE5 | 0.6746 | | | FALSE | TRUE | FALSE | | | | FALSE |
| O95302 | Q96AY3 | 0.6777 | | | FALSE | TRUE | FALSE | | | | FALSE |
| P35579 | Q7Z406 | 0.6782 | | | FALSE | TRUE | FALSE | | | | TRUE |
| Q13242 | Q13243 | 0.6847 | HOM | HOM | TRUE | FALSE | TRUE | 351 | Human | Spliceosome | FALSE |
| Q13243 | Q13247 | 0.6878 | HOM | HOM | TRUE | FALSE | TRUE | 351,2589 | Human | Spliceosome,PGC-1-SRp40-SRp55-SRp75 complex | TRUE |
| P02545 | P08670 | 0.689 | HOM | HOM | TRUE | FALSE | FALSE | | | | FALSE |
| P51991 | Q13151 | 0.6904 | | | FALSE | TRUE | FALSE | | | | FALSE |
| Q14103 | Q96EP5 | 0.6944 | | | FALSE | TRUE | FALSE | | | | FALSE |
| P07900 | P14625 | 0.699 | HOM | HOM | TRUE | FALSE | FALSE | | | | FALSE |
| O14979 | Q13148 | 0.7022 | | | FALSE | TRUE | TRUE | 1332 | Human | Large Drosha complex | TRUE |
| Q9UH03 | Q9UHD8 | 0.7033 | HET | HET | FALSE | FALSE | FALSE | | | | FALSE |

Table S4 Human

| | | | | | | | | | | | | |
|---|---|---|---|---|---|---|---|---|---|---|---|---|
| O75431 | Q13505 | 0.7066 | | | FALSE | TRUE | TRUE | 6249 | Human | MIB complex | | FALSE |
| Q96PK6 | Q9BWF3 | 0.7084 | | | FALSE | TRUE | FALSE | | | | | FALSE |
| P04350 | P68371 | 0.7142 | HET | HET | FALSE | FALSE | FALSE | | | | | FALSE |
| O60568 | Q8NBJ5 | 0.7261 | | | FALSE | TRUE | FALSE | | | | | TRUE |
| P17844 | Q7L014 | 0.7278 | | | FALSE | TRUE | TRUE | 351 | Human | Spliceosome | | FALSE |
| Q00839 | Q1KMD3 | 0.7344 | | | FALSE | TRUE | FALSE | | | | | FALSE |
| Q7L014 | Q92841 | 0.7404 | | | FALSE | TRUE | TRUE | 351 | Human | Spliceosome | | FALSE |
| O43143 | O60231 | 0.7442 | | | FALSE | TRUE | TRUE | 351 | Human | Spliceosome | | FALSE |
| Q15019 | Q9UHD8 | 0.7504 | HOM | HET | TRUE | FALSE | TRUE | 1345 | Human | Septin complex | | FALSE |
| O15460 | P13674 | 0.7561 | | HOM | TRUE | FALSE | FALSE | | | | | FALSE |
| P07437 | P68371 | 0.7592 | HET | HET | FALSE | FALSE | FALSE | | | | | TRUE |
| Q5RKV6 | Q9NPD3 | 0.7735 | HOM, HET | HOM, HET | TRUE, FALSE | FALSE | TRUE | 789 | Human | Exosome | | TRUE |
| P15531 | P22392 | 0.7833 | HOM | HOM | TRUE | FALSE | FALSE | | | | | TRUE |
| P14866 | P26599 | 0.7876 | | | FALSE | TRUE | FALSE | | | | | TRUE |
| Q5XKE5 | Q7Z794 | 0.7879 | | | FALSE | TRUE | FALSE | | | | | FALSE |
| P42025 | P61163 | 0.7929 | | | FALSE | TRUE | FALSE | | | | | TRUE |
| Q7Z5L9 | Q9H1B7 | 0.7936 | | | FALSE | TRUE | TRUE | 6848 | Human | IRF2BP2-IRF2BP1-IRF2BPL complex | | TRUE |
| Q13242 | Q13247 | 0.8076 | HOM | HOM | TRUE | FALSE | TRUE | 351 | Human | Spliceosome | | TRUE |
| P08754 | P63096 | 0.8099 | HET | HOM | TRUE | FALSE | FALSE | | | | | TRUE |
| P14174 | P30046 | 0.8124 | HOM | HOM | TRUE | FALSE | FALSE | | | | | FALSE |
| P04439 | P17693 | 0.8201 | | HOM, HET | TRUE, FALSE | FALSE | FALSE | | | | | TRUE |
| P16989 | P67809 | 0.8205 | | HOM | TRUE | FALSE | FALSE | | | | | FALSE |
| O75955 | Q14254 | 0.8472 | | | FALSE | TRUE | FALSE | | | | | TRUE |
| P04350 | P07437 | 0.8501 | HET | HET | FALSE | FALSE | FALSE | | | | | TRUE |
| O43390 | O60506 | 0.8552 | | | FALSE | TRUE | TRUE | 1181 | Human | C complex spliceosome | | TRUE |

Table S4 Human

| | | | | | | | | | | | |
|---|---|---|---|---|---|---|---|---|---|---|---|
| P30101 | Q15084 | 0.8628 | HET | | FALSE | FALSE | FALSE | | | | FALSE |
| O75718 | Q32P28 | 0.8705 | | | FALSE | TRUE | FALSE | | | | TRUE |
| P62258 | P63104 | 0.8746 | HOM | | TRUE | FALSE | TRUE | 5199 | Human | Kinase maturation complex 1 | TRUE |
| P23246 | Q15233 | 0.8924 | HOM | HET | TRUE | FALSE | TRUE | 330,335,1148 | Human | PSF-p54 complex,p54-PSF-matrin3 complex,snRNP-free U1A complex | TRUE |
| P26232 | P35221 | 0.9101 | | HOM | TRUE | FALSE | FALSE | | | | FALSE |
| P11216 | P11217 | 0.9222 | HOM | HOM | TRUE | FALSE | FALSE | | | | FALSE |
| P10412 | P16403 | 0.9386 | | | FALSE | TRUE | FALSE | | | | FALSE |
| P25685 | Q9UDY4 | 0.9449 | HOM | HOM | TRUE | FALSE | FALSE | | | | TRUE |
| P04075 | P09972 | 0.9473 | | | FALSE | TRUE | FALSE | | | | TRUE |
| Q12905 | Q12906 | 0.9513 | HET | HET | FALSE | FALSE | TRUE | 1332,3055,5183 | Human | Large Drosha complex,Nop56p-associated pre-rRNA complex,DNA-PK-Ku-eIF2-NF90-NF45 complex | TRUE |
| O15397 | O95373 | 0.986 | HET | HET | FALSE | FALSE | FALSE | | | | FALSE |
| P40227 | Q92526 | 0.9939 | HET | HET | FALSE | FALSE | FALSE | | | | TRUE |

Table S4 Mouse

| Prot_1 | Prot_2 | Corr | Prot_1_Hom | Prot_2_Hom | Hom | Mon | CORUM | cluster_id | CORUM_organism | CORUM_name | Biogrid |
|---|---|---|---|---|---|---|---|---|---|---|---|
| Q8BYM5 | Q8VCT4 | -0.87 | HET | | FALSE | FALSE | FALSE | | | | FALSE |
| P04117 | P51880 | -0.85 | HOM | | TRUE | FALSE | FALSE | | | | FALSE |
| Q8CGY6 | Q9CZW5 | -0.84 | HOM | HOM | TRUE | FALSE | FALSE | | | | FALSE |
| O70250 | Q9DBJ1 | -0.83 | | | FALSE | TRUE | FALSE | | | | FALSE |
| P45376 | Q9JII6 | -0.8 | HOM | | TRUE | FALSE | FALSE | | | | FALSE |
| P35762 | Q8R3G9 | -0.8 | HOM | | TRUE | FALSE | FALSE | | | | FALSE |
| Q9QXY6 | Q9Z0R4 | -0.8 | HET | HET | FALSE | FALSE | FALSE | | | | FALSE |
| P45377 | Q9JII6 | -0.79 | | | FALSE | TRUE | FALSE | | | | FALSE |
| P20029 | P63017 | -0.76 | HOM | HOM | TRUE | FALSE | TRUE | 929,2721 | Human | CEN complex, HCF-1 complex | FALSE |
| Q9EQP2 | Q9QXY6 | -0.76 | HET | HET | FALSE | FALSE | FALSE | | | | FALSE |
| O09158 | Q9WVK8 | -0.74 | | | FALSE | TRUE | FALSE | | | | FALSE |
| Q8BG73 | Q9WUZ7 | -0.74 | | | FALSE | TRUE | FALSE | | | | FALSE |
| Q5SQM0 | Q7TNG5 | -0.73 | | | FALSE | TRUE | FALSE | | | | FALSE |
| P20060 | P29416 | -0.72 | | HOM | TRUE | FALSE | FALSE | | | | FALSE |
| Q64176 | Q8BYM5 | -0.72 | | HET | FALSE | FALSE | FALSE | | | | FALSE |
| Q8BYM5 | Q91WG0 | -0.71 | HET | | FALSE | FALSE | FALSE | | | | FALSE |
| P23953 | Q8BYM5 | -0.7 | | HET | FALSE | FALSE | FALSE | | | | FALSE |
| P19246 | P31001 | -0.7 | | | FALSE | TRUE | FALSE | | | | FALSE |
| Q8CI94 | Q9ET01 | -0.7 | HOM | HOM | TRUE | FALSE | FALSE | | | | FALSE |
| Q9R0Q6 | Q9WV32 | -0.69 | | | FALSE | TRUE | FALSE | | | | FALSE |
| Q64459 | Q9WVK8 | -0.69 | | | FALSE | TRUE | FALSE | | | | FALSE |
| Q8BFW7 | Q9WV69 | -0.68 | | | FALSE | TRUE | FALSE | | | | FALSE |
| O35215 | P34884 | -0.68 | HOM | | TRUE | FALSE | FALSE | | | | FALSE |
| Q8R429 | Q8VDN2 | -0.68 | | | FALSE | TRUE | FALSE | | | | FALSE |

Table S4 Mouse

| | | | | | | | | |
|---|---|---|---|---|---|---|---|---|
| Q80TY0 | Q8CJ53 | -0.68 | HOM | HOM | TRUE | FALSE | FALSE | FALSE |
| P97427 | Q9EQF5 | -0.67 | HET | HOM | TRUE | FALSE | FALSE | FALSE |
| P57780 | Q9JI91 | -0.67 | | HOM | TRUE | FALSE | FALSE | FALSE |
| A2AJK6 | A3KFM7 | -0.67 | HOM | HOM | TRUE | FALSE | FALSE | FALSE |
| O09110 | P31938 | -0.67 | | | FALSE | TRUE | FALSE | FALSE |
| P10126 | P62631 | -0.67 | HET | HET | FALSE | FALSE | FALSE | FALSE |
| Q8BLF1 | Q99PG0 | -0.66 | | | FALSE | TRUE | FALSE | FALSE |
| P11404 | P51880 | -0.66 | | | FALSE | TRUE | FALSE | FALSE |
| Q8BK48 | Q8BYM5 | -0.65 | | HET | FALSE | FALSE | FALSE | FALSE |
| Q3THS6 | Q91X83 | -0.64 | | | FALSE | TRUE | FALSE | FALSE |
| O35660 | P48774 | -0.64 | | | FALSE | TRUE | FALSE | FALSE |
| P14142 | P32037 | -0.64 | | | FALSE | TRUE | FALSE | FALSE |
| Q7TQ48 | Q9QXY6 | -0.64 | HET | HET | FALSE | FALSE | FALSE | FALSE |
| Q8JZR0 | Q99PU5 | -0.64 | | | FALSE | TRUE | FALSE | FALSE |
| P97449 | Q8C129 | -0.63 | | | FALSE | TRUE | FALSE | FALSE |
| Q61301 | Q64727 | -0.63 | | HET | FALSE | FALSE | FALSE | FALSE |
| Q8BHD7 | Q91Z31 | -0.61 | | | FALSE | TRUE | FALSE | FALSE |
| Q0II04 | Q61792 | -0.61 | | | FALSE | TRUE | FALSE | FALSE |
| P17742 | Q99KR7 | -0.61 | | | FALSE | TRUE | FALSE | FALSE |
| P53810 | P53811 | -0.6 | HOM | | TRUE | FALSE | FALSE | FALSE |
| Q69ZK9 | Q8VCU1 | -0.6 | | | FALSE | TRUE | FALSE | FALSE |
| P46660 | P48678 | -0.6 | | HOM | TRUE | FALSE | FALSE | FALSE |
| Q9QXY6 | Q9WVK4 | -0.59 | HET | HET | FALSE | FALSE | FALSE | FALSE |
| P04919 | Q5DTL9 | -0.59 | HOM | | TRUE | FALSE | FALSE | FALSE |
| P00920 | P23589 | -0.59 | HOM | | TRUE | FALSE | FALSE | FALSE |
| P18760 | P45591 | -0.59 | | | FALSE | TRUE | FALSE | FALSE |

Table S4 Mouse

| | | | | | | | | |
|---|---|---|---|---|---|---|---|---|
| Q8R326 | Q99K48 | -0.58 | HET | | FALSE | FALSE | FALSE | FALSE |
| P29351 | P35235 | -0.58 | | | FALSE | TRUE | FALSE | FALSE |
| P18654 | Q9WUA6 | -0.58 | | | FALSE | TRUE | FALSE | FALSE |
| Q8R3G9 | Q922J6 | -0.58 | | | FALSE | TRUE | FALSE | FALSE |
| P48318 | Q9DBE0 | -0.58 | HOM | HOM | TRUE | FALSE | FALSE | FALSE |
| Q8JZS0 | Q99L88 | -0.58 | | HOM | TRUE | FALSE | FALSE | FALSE |
| Q69ZK9 | Q91WG0 | -0.58 | | | FALSE | TRUE | FALSE | FALSE |
| Q8BH04 | Q9Z2V4 | -0.57 | | | FALSE | TRUE | FALSE | FALSE |
| P56501 | Q9CR58 | -0.57 | | | FALSE | TRUE | FALSE | FALSE |
| P97449 | Q11011 | -0.56 | | | FALSE | TRUE | FALSE | FALSE |
| Q8CHS7 | Q9EQ06 | -0.56 | | | FALSE | TRUE | FALSE | FALSE |
| P10107 | P97429 | -0.56 | | | FALSE | TRUE | FALSE | FALSE |
| P09542 | Q60605 | -0.56 | | | FALSE | TRUE | FALSE | FALSE |
| P10630 | P60843 | -0.56 | | HET | FALSE | FALSE | FALSE | FALSE |
| Q925N1 | Q925N2 | -0.56 | | | FALSE | TRUE | FALSE | FALSE |
| P12242 | Q9CR58 | -0.55 | HOM | | TRUE | FALSE | FALSE | FALSE |
| Q8BH66 | Q91YH5 | -0.55 | HOM | | TRUE | FALSE | FALSE | FALSE |
| P06801 | Q8BMF3 | -0.55 | | | FALSE | TRUE | FALSE | FALSE |
| B0V2N1 | P06800 | -0.54 | HOM | | TRUE | FALSE | FALSE | FALSE |
| Q9DBG1 | Q9WVK8 | -0.54 | | | FALSE | TRUE | FALSE | FALSE |
| Q9CRD0 | Q9D8W7 | -0.53 | | | FALSE | TRUE | FALSE | FALSE |
| P23953 | Q69ZK9 | -0.53 | | | FALSE | TRUE | FALSE | FALSE |
| P15327 | Q9DBJ1 | -0.53 | HOM | | TRUE | FALSE | FALSE | FALSE |
| Q8BGS1 | Q9WV92 | -0.52 | | | FALSE | TRUE | FALSE | FALSE |
| Q3UQ84 | Q9D0R2 | -0.52 | | | FALSE | TRUE | FALSE | FALSE |
| Q91VK1 | Q9CQC6 | -0.52 | | | FALSE | TRUE | FALSE | FALSE |

Table S4 Mouse

| | | | | | | | | |
|---|---|---|---|---|---|---|---|---|
| P37804 | Q9DAW9 | -0.52 | | | FALSE | TRUE | FALSE | FALSE |
| P63005 | Q8BG40 | -0.52 | HOM | | TRUE | FALSE | FALSE | FALSE |
| Q9QWW1 | Q9Z2Y3 | -0.51 | | | FALSE | TRUE | FALSE | TRUE |
| P08103 | P25911 | -0.51 | HOM | | TRUE | FALSE | FALSE | FALSE |
| Q3TTY5 | Q6IFZ6 | -0.51 | | | FALSE | TRUE | FALSE | FALSE |
| Q6ZQ38 | Q6ZQ73 | -0.51 | HET | | FALSE | FALSE | FALSE | FALSE |
| P35700 | Q61171 | -0.5 | HOM | HOM | TRUE | FALSE | FALSE | FALSE |
| P17182 | P21550 | -0.5 | HOM | HOM | TRUE | FALSE | FALSE | FALSE |
| Q64444 | Q9WVT6 | -0.5 | HOM | | TRUE | FALSE | FALSE | FALSE |
| P30999 | Q68FH0 | -0.5 | | | FALSE | TRUE | FALSE | FALSE |
| Q8BH64 | Q9QXY6 | -0.5 | HOM, HET | HET | TRUE, FALSE | FALSE | FALSE | FALSE |
| P70704 | Q9QZW0 | -0.5 | | | FALSE | TRUE | FALSE | FALSE |
| P16406 | Q11011 | -0.49 | | | FALSE | TRUE | FALSE | FALSE |
| Q8BU33 | Q9QXE0 | -0.49 | | | FALSE | TRUE | FALSE | FALSE |
| Q63880 | Q69ZK9 | -0.49 | | | FALSE | TRUE | FALSE | FALSE |
| P00920 | P16015 | -0.49 | HOM | | TRUE | FALSE | FALSE | FALSE |
| Q9DAW9 | Q9WVA4 | -0.49 | | | FALSE | TRUE | FALSE | FALSE |
| O70622 | Q8K0T0 | -0.49 | | | FALSE | TRUE | FALSE | FALSE |
| Q9DB41 | Q9QXX4 | -0.49 | | | FALSE | TRUE | FALSE | FALSE |
| F8VPU2 | P26041 | -0.48 | | | FALSE | TRUE | FALSE | FALSE |
| Q69ZK9 | Q8QZR3 | -0.48 | | | FALSE | TRUE | FALSE | FALSE |
| P48962 | P51881 | -0.48 | | | FALSE | TRUE | FALSE | FALSE |
| Q64176 | Q69ZK9 | -0.48 | | | FALSE | TRUE | FALSE | FALSE |
| P48320 | Q9DBE0 | -0.48 | | HOM | TRUE | FALSE | FALSE | FALSE |
| P70290 | Q9WV34 | -0.47 | HOM | HOM | TRUE | FALSE | FALSE | FALSE |

Table S4 Mouse

| | | | | | | | | |
|---|---|---|---|---|---|---|---|---|
| P60764 | Q05144 | -0.47 | | HET | FALSE | FALSE | FALSE | FALSE |
| O55143 | Q8R429 | -0.47 | | | FALSE | TRUE | FALSE | FALSE |
| P21836 | Q8QZR3 | -0.47 | HOM | | TRUE | FALSE | FALSE | FALSE |
| P28843 | Q9Z218 | -0.47 | HOM | HOM | TRUE | FALSE | FALSE | FALSE |
| P21836 | Q8VCT4 | -0.46 | HOM | | TRUE | FALSE | FALSE | FALSE |
| O88952 | Q61234 | -0.46 | | HOM | TRUE | FALSE | FALSE | FALSE |
| P06800 | P18052 | -0.46 | | | FALSE | TRUE | FALSE | FALSE |
| P21836 | Q8BK48 | -0.46 | HOM | | TRUE | FALSE | FALSE | FALSE |
| Q8BYM5 | Q8QZR3 | -0.46 | HET | | FALSE | FALSE | FALSE | FALSE |
| P51881 | Q3V132 | -0.46 | | | FALSE | TRUE | FALSE | FALSE |
| Q69ZK9 | Q8VCT4 | -0.45 | | | FALSE | TRUE | FALSE | FALSE |
| Q9EPL9 | Q9R0H0 | -0.45 | | | FALSE | TRUE | FALSE | FALSE |
| P48410 | P55096 | -0.45 | HOM | HET | TRUE | FALSE | FALSE | TRUE |
| P70290 | Q811D0 | -0.45 | HOM | HOM | TRUE | FALSE | FALSE | FALSE |
| A2AKK5 | Q9QYR9 | -0.45 | | | FALSE | TRUE | FALSE | FALSE |
| Q8VHQ9 | Q91V12 | -0.44 | HOM | HOM | TRUE | FALSE | FALSE | FALSE |
| P40237 | Q8R3G9 | -0.44 | | | FALSE | TRUE | FALSE | FALSE |
| Q8BSL7 | Q9D0J4 | -0.44 | | HET | FALSE | FALSE | FALSE | FALSE |
| Q64152 | Q9CQH7 | -0.44 | HET | | FALSE | FALSE | FALSE | FALSE |
| O09110 | Q63932 | -0.44 | | | FALSE | TRUE | FALSE | FALSE |
| O35381 | Q9EST5 | -0.44 | | | FALSE | TRUE | FALSE | FALSE |
| O08528 | Q91W97 | -0.44 | | | FALSE | TRUE | FALSE | FALSE |
| Q3TCH7 | Q9JLV5 | -0.43 | HOM, HET | HOM, HET | TRUE, FALSE | FALSE | FALSE | FALSE |
| P97823 | Q9WTL7 | -0.43 | | | FALSE | TRUE | FALSE | FALSE |
| Q8BTW9 | Q8CIN4 | -0.43 | HET | | FALSE | FALSE | FALSE | FALSE |

Table S4 Mouse

| | | | | | | | | |
|---|---|---|---|---|---|---|---|---|
| P05063 | Q91Y97 | -0.43 | | | FALSE | TRUE | FALSE | FALSE |
| O08807 | Q61171 | -0.42 | HOM | HOM | TRUE | FALSE | FALSE | FALSE |
| Q91VM9 | Q9D819 | -0.42 | | | FALSE | TRUE | FALSE | FALSE |
| Q9CR62 | Q9QZD8 | -0.42 | | | FALSE | TRUE | FALSE | FALSE |
| Q61490 | Q9R069 | -0.42 | | | FALSE | TRUE | FALSE | FALSE |
| P31649 | P31650 | -0.42 | | | FALSE | TRUE | FALSE | FALSE |
| P13597 | Q60625 | -0.42 | HOM | HET | TRUE | FALSE | FALSE | FALSE |
| Q9D0R2 | Q9JKF7 | -0.42 | | | FALSE | TRUE | FALSE | FALSE |
| P13541 | Q61879 | -0.42 | | | FALSE | TRUE | FALSE | FALSE |
| P28650 | P46664 | -0.42 | | | FALSE | TRUE | FALSE | FALSE |
| Q8BGA8 | Q99NB1 | -0.42 | | HOM | TRUE | FALSE | FALSE | FALSE |
| P84084 | Q9CQW2 | -0.42 | | | FALSE | TRUE | FALSE | FALSE |
| Q8BG95 | Q9DBR7 | -0.42 | | | FALSE | TRUE | FALSE | FALSE |
| O09110 | P47809 | -0.42 | | | FALSE | TRUE | FALSE | FALSE |
| Q811D0 | Q91XM9 | -0.41 | HOM | HOM | TRUE | FALSE | FALSE | FALSE |
| P84078 | Q9CQW2 | -0.41 | | | FALSE | TRUE | FALSE | FALSE |
| O35465 | Q62446 | -0.41 | HET | | FALSE | FALSE | FALSE | FALSE |
| P41216 | Q99PU5 | -0.41 | | | FALSE | TRUE | FALSE | FALSE |
| P07310 | Q04447 | -0.41 | | | FALSE | TRUE | FALSE | FALSE |
| O08600 | Q8C163 | -0.41 | | | FALSE | TRUE | FALSE | FALSE |
| P21836 | P23953 | -0.41 | HOM | | TRUE | FALSE | FALSE | FALSE |
| O54865 | P52785 | -0.41 | HOM | | TRUE | FALSE | FALSE | FALSE |
| P45591 | Q9R0P5 | -0.41 | | | FALSE | TRUE | FALSE | FALSE |
| P13020 | Q9JJ28 | -0.4 | | | FALSE | TRUE | FALSE | FALSE |
| Q80TH2 | Q80U72 | -0.4 | | | FALSE | TRUE | FALSE | FALSE |
| Q91V61 | Q99JR1 | -0.4 | | | FALSE | TRUE | FALSE | FALSE |

Table S4 Mouse

| | | | | | | | | | | | |
|---|---|---|---|---|---|---|---|---|---|---|---|
| P05480 | Q04736 | -0.4 | HOM | | TRUE | FALSE | FALSE | | | | FALSE |
| O89104 | Q62277 | -0.39 | | HOM | TRUE | FALSE | FALSE | | | | FALSE |
| Q9D4H8 | Q9WTX6 | -0.39 | HET | HOM, HET | TRUE, FALSE | FALSE | TRUE | 2715 | Human | Ubiquitin E3 ligase | FALSE |
| P62482 | Q8CG76 | -0.39 | HET | | FALSE | FALSE | FALSE | | | | FALSE |
| P14142 | P17809 | -0.39 | | | FALSE | TRUE | FALSE | | | | FALSE |
| Q8CGY6 | Q91Z38 | -0.39 | HOM | HOM | TRUE | FALSE | FALSE | | | | FALSE |
| Q9QYF9 | Q9QYG0 | -0.39 | | HOM | TRUE | FALSE | FALSE | | | | FALSE |
| P21836 | Q63880 | -0.39 | HOM | | TRUE | FALSE | FALSE | | | | FALSE |
| P62245 | Q9JKW0 | -0.39 | | | FALSE | TRUE | FALSE | | | | FALSE |
| Q60714 | Q91VE0 | -0.39 | | | FALSE | TRUE | FALSE | | | | FALSE |
| O88990 | P57780 | -0.39 | HOM | | TRUE | FALSE | FALSE | | | | FALSE |
| Q8C7K6 | Q9CQF9 | -0.39 | | | FALSE | TRUE | FALSE | | | | FALSE |
| P21836 | Q64176 | -0.38 | HOM | | TRUE | FALSE | FALSE | | | | FALSE |
| P03995 | P20152 | -0.38 | | | FALSE | TRUE | FALSE | | | | FALSE |
| P15532 | Q01768 | -0.38 | | | FALSE | TRUE | FALSE | | | | FALSE |
| P31324 | Q9DBC7 | -0.38 | | | FALSE | TRUE | FALSE | | | | FALSE |
| Q8BFR5 | Q9JHW4 | -0.38 | | | FALSE | TRUE | FALSE | | | | FALSE |
| P48036 | Q07076 | -0.38 | | | FALSE | TRUE | FALSE | | | | FALSE |
| Q6PE01 | Q8BG40 | -0.38 | | | FALSE | TRUE | FALSE | | | | FALSE |
| Q6GV12 | Q8VCR2 | -0.38 | | | FALSE | TRUE | FALSE | | | | FALSE |
| Q8BGS7 | Q8C025 | -0.38 | | | FALSE | TRUE | FALSE | | | | FALSE |
| P50462 | P97315 | -0.38 | | | FALSE | TRUE | FALSE | | | | FALSE |
| P70290 | Q62108 | -0.38 | HOM | HOM | TRUE | FALSE | FALSE | | | | FALSE |
| Q61490 | Q8R2Y2 | -0.38 | | | FALSE | TRUE | FALSE | | | | FALSE |
| P26039 | Q9JKY5 | -0.38 | HOM | | TRUE | FALSE | FALSE | | | | FALSE |

Table S4 Mouse

| | | | | | | | | |
|---|---|---|---|---|---|---|---|---|
| A2APT9 | Q8BNW9 | -0.37 | | | FALSE | TRUE | FALSE | FALSE |
| Q8VCC2 | Q8VCT4 | -0.37 | | | FALSE | TRUE | FALSE | FALSE |
| P10107 | P97384 | -0.37 | | | FALSE | TRUE | FALSE | FALSE |
| P16054 | P63318 | -0.37 | | | FALSE | TRUE | FALSE | FALSE |
| P46735 | Q5SYD0 | -0.37 | | HOM | TRUE | FALSE | FALSE | FALSE |
| P39447 | Q9WV34 | -0.37 | | HOM | TRUE | FALSE | FALSE | FALSE |
| A2ATU0 | Q60597 | -0.37 | | | FALSE | TRUE | FALSE | FALSE |
| O88951 | Q99L88 | -0.37 | HOM | HOM | TRUE | FALSE | FALSE | FALSE |
| Q7TMK9 | Q91WT8 | -0.36 | | | FALSE | TRUE | FALSE | FALSE |
| P48722 | Q9JKR6 | -0.36 | HOM | HOM | TRUE | FALSE | FALSE | FALSE |
| Q9CYH2 | Q9DB60 | -0.36 | | | FALSE | TRUE | FALSE | FALSE |
| O35488 | Q60714 | -0.36 | | | FALSE | TRUE | FALSE | FALSE |
| P16546 | Q9JI91 | -0.36 | HOM | HOM | TRUE | FALSE | FALSE | FALSE |
| Q9CQW2 | Q9D0J4 | -0.36 | | HET | FALSE | FALSE | FALSE | FALSE |
| O35955 | P70195 | -0.36 | HOM | HOM | TRUE | FALSE | FALSE | FALSE |
| Q04447 | Q6P8J7 | -0.36 | | HOM | TRUE | FALSE | FALSE | FALSE |
| P05480 | P08103 | -0.36 | HOM | HOM | TRUE | FALSE | FALSE | FALSE |
| Q63880 | Q8BYM5 | -0.36 | | HET | FALSE | FALSE | FALSE | FALSE |
| P05977 | Q60605 | -0.35 | | | FALSE | TRUE | FALSE | FALSE |
| P11679 | Q6IFZ6 | -0.35 | | | FALSE | TRUE | FALSE | FALSE |
| P20152 | P21619 | -0.35 | | | FALSE | TRUE | FALSE | FALSE |
| P31001 | P46660 | -0.35 | | | FALSE | TRUE | FALSE | FALSE |

Table S4 Mouse

| | | | | | | | | | | | |
|---|---|---|---|---|---|---|---|---|---|---|---|
| Q60972 | Q60973 | -0.35 | | | FALSE | TRUE | TRUE | 2750,61,100,282,587,632,646,649,650,685,696,732,738,739,743,886,888,889,996,1257,2721,2851 | Mouse,Human | Gata1-Fog1-MeCP1 complex,Mi2/NuRD complex,hNURF complex,SNF2h-cohesin-NuRD complex,NuRD.1 complex,Anti-HDAC2 complex,HDAC1-associated protein complex,HDAC1-associated core complex cII,HDAC2-asscociated core | FALSE |
| O08553 | Q9EQF5 | -0.35 | HOM | HOM | TRUE | FALSE | FALSE | | | | FALSE |
| P26040 | P26043 | -0.35 | | | FALSE | TRUE | FALSE | | | | FALSE |
| Q9QZB1 | Q9Z2H2 | -0.35 | | | FALSE | TRUE | FALSE | | | | FALSE |
| P15626 | P48774 | -0.35 | | | FALSE | TRUE | FALSE | | | | FALSE |
| Q91V61 | Q925N2 | -0.35 | | | FALSE | TRUE | FALSE | | | | FALSE |
| Q6GQS1 | Q8BMD8 | -0.35 | | | FALSE | TRUE | FALSE | | | | FALSE |
| P55258 | P62823 | -0.34 | | | FALSE | TRUE | FALSE | | | | FALSE |
| Q9EQF5 | Q9EQF6 | -0.34 | HOM | HET | TRUE | FALSE | FALSE | | | | TRUE |
| P32883 | P62071 | -0.34 | | | FALSE | TRUE | FALSE | | | | FALSE |
| Q8BYM5 | Q8VCU1 | -0.34 | HET | | FALSE | FALSE | FALSE | | | | FALSE |
| Q5YD48 | Q7TMK9 | -0.34 | | | FALSE | TRUE | FALSE | | | | FALSE |
| Q80TB8 | Q9DCS3 | -0.34 | HOM | HOM | TRUE | FALSE | FALSE | | | | FALSE |

Table S4 Mouse

| | | | | | | | | | | | |
|---|---|---|---|---|---|---|---|---|---|---|---|
| Q02248 | Q02257 | -0.34 | HET,HOM | | FALSE,TRUE | FALSE | TRUE | 5177 | Human | Polycystin-1 multiprotein complex | FALSE |
| Q920P5 | Q9R0Y5 | -0.34 | HOM | HOM | TRUE | FALSE | FALSE | | | | FALSE |
| Q64133 | Q8BW75 | -0.34 | HET | HOM | TRUE | FALSE | FALSE | | | | FALSE |
| Q6GQS1 | Q8C0K5 | -0.33 | | | FALSE | TRUE | FALSE | | | | FALSE |
| Q6GQS1 | Q8R0Y8 | -0.33 | | | FALSE | TRUE | FALSE | | | | FALSE |
| P23589 | Q64444 | -0.33 | | HOM | TRUE | FALSE | FALSE | | | | FALSE |
| P31649 | Q61327 | -0.33 | | | FALSE | TRUE | FALSE | | | | FALSE |
| P21836 | Q8VCU1 | -0.33 | HOM | | TRUE | FALSE | FALSE | | | | FALSE |
| Q99104 | Q9QZZ4 | -0.33 | HOM | HOM | TRUE | FALSE | FALSE | | | | FALSE |
| P70349 | Q9D0S9 | -0.33 | | | FALSE | TRUE | FALSE | | | | FALSE |
| Q64518 | Q8VDN2 | -0.32 | | | FALSE | TRUE | FALSE | | | | FALSE |
| P61202 | Q8BG32 | -0.32 | | | FALSE | TRUE | FALSE | | | | FALSE |
| P97742 | Q924X2 | -0.32 | HOM | | TRUE | FALSE | FALSE | | | | FALSE |
| P39447 | P70175 | -0.32 | | HOM | TRUE | FALSE | FALSE | | | | FALSE |
| P60764 | P84096 | -0.32 | | | FALSE | TRUE | FALSE | | | | FALSE |
| Q6IFZ6 | Q8VED5 | -0.32 | | | FALSE | TRUE | FALSE | | | | FALSE |
| Q6URW6 | Q8VDD5 | -0.32 | | | FALSE | TRUE | FALSE | | | | FALSE |
| P49813 | Q9JKK7 | -0.32 | | | FALSE | TRUE | FALSE | | | | FALSE |
| P14733 | P31001 | -0.32 | | | FALSE | TRUE | FALSE | | | | FALSE |
| P35293 | Q9D1G1 | -0.32 | | | FALSE | TRUE | FALSE | | | | FALSE |
| Q9WTP7 | Q9WUR9 | -0.31 | | | FALSE | TRUE | FALSE | | | | FALSE |
| Q8BH00 | Q9EQ20 | -0.31 | | | FALSE | TRUE | FALSE | | | | FALSE |
| O88792 | Q9D8B7 | -0.31 | HOM | | TRUE | FALSE | FALSE | | | | FALSE |
| Q9DB73 | Q9DCN2 | -0.31 | | | FALSE | TRUE | FALSE | | | | FALSE |

Table S4 Mouse

| | | | | | | | | | | | |
|---|---|---|---|---|---|---|---|---|---|---|---|
| P97457 | Q3THE2 | -0.31 | | | FALSE | TRUE | FALSE | | | | FALSE |
| P48787 | Q9WUZ5 | -0.31 | | | FALSE | TRUE | FALSE | | | | FALSE |
| P21300 | P45377 | -0.31 | | | FALSE | TRUE | FALSE | | | | FALSE |
| Q3V3R1 | Q922D8 | -0.31 | | | FALSE | TRUE | FALSE | | | | FALSE |
| P19096 | Q9D404 | -0.31 | HOM | HOM | TRUE | FALSE | FALSE | | | | FALSE |
| Q8JZR0 | Q91WC3 | -0.3 | | | FALSE | TRUE | FALSE | | | | FALSE |
| O35737 | Q9Z2X1 | -0.3 | | | FALSE | TRUE | TRUE | 5142,1181 | Mouse,Human | DCS complex, C complex spliceosome | FALSE |
| O89104 | Q8BGN8 | -0.3 | | | FALSE | TRUE | FALSE | | | | FALSE |
| O35488 | Q91VE0 | -0.3 | | | FALSE | TRUE | FALSE | | | | FALSE |
| Q91VW3 | Q9WUZ7 | -0.3 | HOM | | TRUE | FALSE | FALSE | | | | FALSE |
| Q9D6M3 | Q9QXX4 | -0.3 | | | FALSE | TRUE | FALSE | | | | FALSE |
| Q9JKK7 | Q9JLH8 | -0.3 | | | FALSE | TRUE | FALSE | | | | FALSE |
| P13541 | Q9JMH9 | -0.3 | | | FALSE | TRUE | FALSE | | | | FALSE |
| Q922J3 | Q9D1E6 | -0.3 | | HET | FALSE | FALSE | FALSE | | | | FALSE |
| Q925N0 | Q925N2 | -0.3 | | | FALSE | TRUE | FALSE | | | | FALSE |
| O08989 | P62071 | -0.3 | | | FALSE | TRUE | FALSE | | | | FALSE |
| P21300 | P45376 | -0.3 | | HOM | TRUE | FALSE | FALSE | | | | FALSE |
| A2ASZ8 | Q6GQS1 | -0.29 | | | FALSE | TRUE | FALSE | | | | FALSE |
| O88492 | P43883 | -0.29 | | | FALSE | TRUE | FALSE | | | | FALSE |
| P55258 | P61027 | -0.29 | | | FALSE | TRUE | FALSE | | | | FALSE |
| P63001 | Q05144 | -0.29 | | HET | FALSE | FALSE | FALSE | | | | FALSE |
| Q3UMU9 | Q9JMG7 | -0.29 | | | FALSE | TRUE | FALSE | | | | FALSE |
| P13634 | Q9WVT6 | -0.29 | | | FALSE | TRUE | FALSE | | | | FALSE |
| P17156 | P20029 | -0.29 | | HOM | TRUE | FALSE | FALSE | | | | FALSE |

Table S4 Mouse

| | | | | | | | | |
|---|---|---|---|---|---|---|---|---|
| O08989 | Q9JIW9 | -0.29 | | HET | FALSE | FALSE | FALSE | FALSE |
| B1AZP2 | Q6PFD5 | -0.29 | | | FALSE | TRUE | FALSE | FALSE |
| Q6P9R2 | Q8CIN4 | -0.29 | HOM | | TRUE | FALSE | FALSE | FALSE |
| Q61151 | Q6PD03 | -0.28 | HET | HET | FALSE | FALSE | FALSE | FALSE |
| Q60902 | Q7TQ48 | -0.28 | HET | HET | FALSE | FALSE | FALSE | FALSE |
| P21619 | P31001 | -0.28 | | | FALSE | TRUE | FALSE | FALSE |
| Q8BK48 | Q8VCC2 | -0.28 | | | FALSE | TRUE | FALSE | FALSE |
| Q02566 | Q8VDD5 | -0.28 | | | FALSE | TRUE | FALSE | FALSE |
| Q69ZK9 | Q8VCC2 | -0.28 | | | FALSE | TRUE | FALSE | FALSE |
| P45376 | Q8K023 | -0.28 | HOM | HOM | TRUE | FALSE | FALSE | FALSE |
| P61750 | Q9CQW2 | -0.28 | | | FALSE | TRUE | FALSE | FALSE |
| O54916 | Q60902 | -0.28 | HET | HET | FALSE | FALSE | FALSE | FALSE |
| P61205 | Q8BSL7 | -0.28 | | | FALSE | TRUE | FALSE | FALSE |
| Q8BGF9 | Q9QXX4 | -0.28 | | | FALSE | TRUE | FALSE | FALSE |
| P05480 | P25911 | -0.28 | HOM | | TRUE | FALSE | FALSE | FALSE |
| Q4LDG0 | Q60714 | -0.28 | | | FALSE | TRUE | FALSE | FALSE |
| P21836 | Q91WG0 | -0.28 | HOM | | TRUE | FALSE | FALSE | FALSE |
| P07901 | Q9CQN1 | -0.27 | HOM | HOM | TRUE | FALSE | FALSE | FALSE |
| O88343 | P04919 | -0.27 | | HOM | TRUE | FALSE | FALSE | FALSE |
| P45377 | Q8K023 | -0.27 | | HOM | TRUE | FALSE | FALSE | FALSE |
| P61750 | Q8BSL7 | -0.27 | | | FALSE | TRUE | FALSE | FALSE |
| Q9CR58 | Q9CR62 | -0.27 | | | FALSE | TRUE | FALSE | FALSE |
| Q3UV17 | Q6IFZ6 | -0.27 | | | FALSE | TRUE | FALSE | FALSE |
| A2AQ07 | Q7TMM9 | -0.27 | HET | HET | FALSE | FALSE | FALSE | FALSE |
| Q6GV12 | Q8CHS7 | -0.27 | | | FALSE | TRUE | FALSE | FALSE |
| Q9EPL9 | Q9QXD1 | -0.27 | | | FALSE | TRUE | FALSE | FALSE |

Table S4 Mouse

| | | | | | | | | |
|---|---|---|---|---|---|---|---|---|
| Q8CH18 | Q8VDP4 | -0.27 | | | FALSE | TRUE | FALSE | FALSE |
| P14106 | Q60994 | -0.27 | HET | HET,HOM | FALSE,TRUE | FALSE | FALSE | FALSE |
| Q4KUS2 | Q8K0T7 | -0.27 | HOM | | TRUE | FALSE | FALSE | FALSE |
| P45376 | P70694 | -0.26 | HOM | HOM | TRUE | FALSE | FALSE | FALSE |
| Q8K0T0 | Q99P72 | -0.26 | | | FALSE | TRUE | FALSE | FALSE |
| Q91V14 | Q9WVL3 | -0.26 | | | FALSE | TRUE | FALSE | FALSE |
| P19157 | P48774 | -0.26 | HOM | | TRUE | FALSE | FALSE | FALSE |
| P97429 | Q07076 | -0.26 | | | FALSE | TRUE | FALSE | FALSE |
| Q8BT60 | Q9Z140 | -0.26 | | | FALSE | TRUE | FALSE | FALSE |
| P28271 | Q99KI0 | -0.26 | | | FALSE | TRUE | FALSE | FALSE |
| P61205 | Q9CQW2 | -0.26 | | | FALSE | TRUE | FALSE | FALSE |
| Q61035 | Q99KK9 | -0.25 | | | FALSE | TRUE | FALSE | FALSE |
| Q3UGR5 | Q9D7I5 | -0.25 | | HOM | TRUE | FALSE | FALSE | FALSE |
| Q7SIG6 | Q9QWY8 | -0.25 | | HOM | TRUE | FALSE | FALSE | FALSE |
| A2AJK6 | Q6PDQ2 | -0.25 | HOM | HOM | TRUE | FALSE | FALSE | FALSE |
| P48193 | Q9WV92 | -0.25 | | | FALSE | TRUE | FALSE | FALSE |
| Q5SYD0 | Q9WTI7 | -0.25 | HOM | HOM | TRUE | FALSE | FALSE | FALSE |
| Q60996 | Q6PD03 | -0.25 | HET | HET | FALSE | FALSE | FALSE | FALSE |
| P18572 | P21995 | -0.25 | HOM | | TRUE | FALSE | FALSE | FALSE |
| Q11011 | Q9EQH2 | -0.25 | | HET | FALSE | FALSE | FALSE | FALSE |
| B1AZP2 | Q9D415 | -0.25 | | | FALSE | TRUE | FALSE | TRUE |
| P17183 | P21550 | -0.24 | HOM | HOM | TRUE | FALSE | FALSE | FALSE |
| Q08857 | Q61009 | -0.24 | HET | | FALSE | FALSE | FALSE | FALSE |
| O35685 | Q8R1N4 | -0.24 | | | FALSE | TRUE | FALSE | FALSE |
| O09117 | Q62277 | -0.24 | | HOM | TRUE | FALSE | FALSE | FALSE |

Table S4 Mouse

| | | | | | | | | | | | |
|---|---|---|---|---|---|---|---|---|---|---|---|
| P84084 | Q8BSL7 | -0.24 | | | FALSE | TRUE | FALSE | | | | FALSE |
| A2AQ07 | Q9CWF2 | -0.24 | HET | | FALSE | FALSE | FALSE | | | | FALSE |
| P29351 | P54830 | -0.23 | | | FALSE | TRUE | FALSE | | | | FALSE |
| P19639 | P48774 | -0.23 | | | FALSE | TRUE | FALSE | | | | FALSE |
| P55258 | P63011 | -0.23 | | | FALSE | TRUE | FALSE | | | | FALSE |
| P32883 | Q9JIW9 | -0.23 | | HET | FALSE | FALSE | FALSE | | | | FALSE |
| O88990 | Q7TPR4 | -0.23 | HOM | | TRUE | FALSE | FALSE | | | | FALSE |
| P28651 | Q64444 | -0.23 | | HOM | TRUE | FALSE | FALSE | | | | FALSE |
| P12382 | P47857 | -0.23 | HET | HET | FALSE | FALSE | FALSE | | | | FALSE |
| P97379 | P97855 | -0.23 | | HOM | TRUE | FALSE | FALSE | | | | FALSE |
| Q8JZU0 | Q9DCN1 | -0.22 | | | FALSE | TRUE | FALSE | | | | FALSE |
| F8VPU2 | P26043 | -0.22 | | | FALSE | TRUE | FALSE | | | | FALSE |
| Q8BG73 | Q9JJU8 | -0.22 | | | FALSE | TRUE | FALSE | | | | FALSE |
| P57780 | Q7TPR4 | -0.22 | | | FALSE | TRUE | FALSE | | | | FALSE |
| Q8VHE6 | Q91XQ0 | -0.22 | | | FALSE | TRUE | FALSE | | | | FALSE |
| P12849 | P31324 | -0.22 | | | FALSE | TRUE | FALSE | | | | FALSE |
| O55143 | Q6PIE5 | -0.22 | | | FALSE | TRUE | FALSE | | | | FALSE |
| P56371 | Q8BHC1 | -0.22 | HET | | FALSE | FALSE | FALSE | | | | FALSE |
| O09164 | Q9WU84 | -0.22 | HOM | HOM | TRUE | FALSE | FALSE | | | | FALSE |
| B2RSH2 | P21278 | -0.21 | HOM | | TRUE | FALSE | FALSE | | | | FALSE |
| D3YZU1 | Q4ACU6 | -0.21 | | | FALSE | TRUE | FALSE | | | | TRUE |
| Q80W22 | Q8BH55 | -0.21 | | | FALSE | TRUE | FALSE | | | | FALSE |
| P16015 | Q9WVT6 | -0.21 | | | FALSE | TRUE | FALSE | | | | FALSE |
| Q8R317 | Q9QZM0 | -0.21 | HOM, HET | HET | TRUE, FALSE | FALSE | TRUE | 5209 | Human | Ubiquilin-proteasome complex | TRUE |
| Q9DAW9 | Q9R1Q8 | -0.21 | | | FALSE | TRUE | FALSE | | | | FALSE |

Table S4 Mouse

| | | | | | | | | |
|---|---|---|---|---|---|---|---|---|
| Q3THE2 | Q9QVP4 | -0.21 | | | FALSE | TRUE | FALSE | FALSE |
| Q80UG2 | Q8CJH3 | -0.21 | | HOM | TRUE | FALSE | FALSE | FALSE |
| Q3URE1 | Q8VCW8 | -0.21 | | | FALSE | TRUE | FALSE | FALSE |
| P08551 | P48678 | -0.21 | | HOM | TRUE | FALSE | FALSE | FALSE |
| P35293 | P62821 | -0.21 | | | FALSE | TRUE | FALSE | FALSE |
| O70318 | Q9WV92 | -0.2 | | | FALSE | TRUE | FALSE | FALSE |
| P23953 | Q8VCC2 | -0.2 | | | FALSE | TRUE | FALSE | FALSE |
| B2RXS4 | Q80UG2 | -0.2 | HET | | FALSE | FALSE | FALSE | FALSE |
| Q8BGF9 | Q9D6M3 | -0.2 | | | FALSE | TRUE | FALSE | FALSE |
| Q8BH24 | Q9ET30 | -0.2 | | | FALSE | TRUE | FALSE | FALSE |
| P62331 | Q8BSL7 | -0.2 | HOM | | TRUE | FALSE | FALSE | FALSE |
| P61164 | Q8BFZ3 | -0.2 | | | FALSE | TRUE | FALSE | FALSE |
| Q00PI9 | Q8VEK3 | -0.2 | | | FALSE | TRUE | FALSE | FALSE |
| E9Q401 | P70227 | -0.2 | | | FALSE | TRUE | FALSE | FALSE |
| O70318 | Q8BGS1 | -0.2 | | | FALSE | TRUE | FALSE | FALSE |
| Q62261 | Q9JI91 | -0.2 | HOM | HOM | TRUE | FALSE | FALSE | FALSE |
| Q8VCA8 | Q9CZC8 | -0.19 | | | FALSE | TRUE | FALSE | FALSE |
| P45377 | P70694 | -0.19 | | HOM | TRUE | FALSE | FALSE | FALSE |
| P47934 | P97742 | -0.19 | | HOM | TRUE | FALSE | FALSE | FALSE |
| Q60598 | Q62418 | -0.19 | | | FALSE | TRUE | FALSE | FALSE |
| Q91XQ0 | Q9JHU4 | -0.19 | | HOM | TRUE | FALSE | FALSE | FALSE |
| Q8R3P0 | Q91XE4 | -0.19 | | HOM | TRUE | FALSE | FALSE | FALSE |
| Q8CC35 | Q91YE8 | -0.19 | | HOM | TRUE | FALSE | FALSE | FALSE |
| P04104 | P11679 | -0.19 | | | FALSE | TRUE | FALSE | FALSE |
| P48774 | Q80W21 | -0.19 | | HOM | TRUE | FALSE | FALSE | FALSE |
| P03995 | P48678 | -0.19 | | HOM | TRUE | FALSE | FALSE | FALSE |

Table S4 Mouse

| | | | | | | | | | | | |
|---|---|---|---|---|---|---|---|---|---|---|---|
| P70333 | Q9Z2X1 | -0.19 | | | FALSE | TRUE | FALSE | | | | FALSE |
| O55143 | Q64518 | -0.19 | | | FALSE | TRUE | FALSE | | | | FALSE |
| P43883 | Q8CGN5 | -0.19 | | | FALSE | TRUE | FALSE | | | | FALSE |
| P11276 | Q8BYI9 | -0.19 | HOM | | TRUE | FALSE | FALSE | | | | FALSE |
| Q61411 | Q9JIW9 | -0.18 | | HET | FALSE | FALSE | FALSE | | | | FALSE |
| O70209 | O70400 | -0.18 | | | FALSE | TRUE | FALSE | | | | FALSE |
| O55234 | P28063 | -0.18 | HOM | HOM | TRUE | FALSE | FALSE | | | | FALSE |
| Q62523 | Q9WV69 | -0.18 | | | FALSE | TRUE | FALSE | | | | FALSE |
| O70325 | P11352 | -0.18 | | HOM | TRUE | FALSE | FALSE | | | | FALSE |
| Q64464 | Q9WVK8 | -0.18 | | | FALSE | TRUE | FALSE | | | | FALSE |
| Q60605 | Q8CI43 | -0.18 | | | FALSE | TRUE | FALSE | | | | FALSE |
| A2AQP0 | Q8VDD5 | -0.18 | | | FALSE | TRUE | FALSE | | | | FALSE |
| P58021 | Q9ET30 | -0.18 | | | FALSE | TRUE | FALSE | | | | FALSE |
| P22723 | P26048 | -0.18 | | | FALSE | TRUE | FALSE | | | | FALSE |
| Q01279 | Q9QVP9 | -0.18 | HOM | HOM | TRUE | FALSE | FALSE | | | | FALSE |
| Q7TSQ8 | Q99LB7 | -0.18 | | | FALSE | TRUE | FALSE | | | | FALSE |
| P61750 | P62331 | -0.18 | | HOM | TRUE | FALSE | FALSE | | | | FALSE |
| Q9R0N5 | Q9R0N7 | -0.18 | | HET | FALSE | FALSE | FALSE | | | | FALSE |
| P51880 | Q05816 | -0.18 | | | FALSE | TRUE | FALSE | | | | FALSE |
| Q8VCR2 | Q99J47 | -0.17 | | | FALSE | TRUE | FALSE | | | | FALSE |
| Q60902 | Q9EQP2 | -0.17 | HET | HET | FALSE | FALSE | FALSE | | | | FALSE |
| Q9ERI6 | Q9QYF1 | -0.17 | | | FALSE | TRUE | FALSE | | | | FALSE |
| P45376 | Q8VCX1 | -0.17 | HOM | | TRUE | FALSE | FALSE | | | | FALSE |
| Q60875 | Q61210 | -0.17 | | | FALSE | TRUE | TRUE | 6671 | Human | PI4K2A-WASH complex | FALSE |
| P39447 | Q62108 | -0.17 | | HOM | TRUE | FALSE | FALSE | | | | FALSE |

Table S4 Mouse

| | | | | | | | | | | | |
|---|---|---|---|---|---|---|---|---|---|---|---|
| O55028 | Q922H2 | -0.16 | | | FALSE | TRUE | FALSE | | | | FALSE |
| O35381 | P97822 | -0.16 | | | FALSE | TRUE | FALSE | | | | FALSE |
| P62331 | Q9D0J4 | -0.16 | HOM | HET | TRUE | FALSE | FALSE | | | | FALSE |
| Q62108 | Q91XM9 | -0.16 | HOM | HOM | TRUE | FALSE | FALSE | | | | TRUE |
| P45377 | Q8VCX1 | -0.16 | | | FALSE | TRUE | FALSE | | | | FALSE |
| Q6PIC6 | Q8R429 | -0.16 | | | FALSE | TRUE | FALSE | | | | FALSE |
| O88533 | Q9DBE0 | -0.16 | HOM | HOM | TRUE | FALSE | FALSE | | | | FALSE |
| Q9JLV5 | Q9WTX6 | -0.16 | HOM, HET | HOM, HET | TRUE, FALSE | FALSE | TRUE | 2715 | Human | Ubiquitin E3 ligase | FALSE |
| Q3UE37 | Q6ZPJ3 | -0.16 | | | FALSE | TRUE | FALSE | | | | FALSE |
| P06801 | Q99KE1 | -0.16 | | HOM | TRUE | FALSE | FALSE | | | | FALSE |
| O35737 | Q8C5Q4 | -0.15 | | | FALSE | TRUE | FALSE | | | | FALSE |
| Q3UNZ8 | Q80TB8 | -0.15 | | HOM | TRUE | FALSE | FALSE | | | | FALSE |
| Q3UMU9 | Q99JF8 | -0.15 | | | FALSE | TRUE | FALSE | | | | FALSE |
| Q69ZK9 | Q8BK48 | -0.15 | | | FALSE | TRUE | FALSE | | | | FALSE |
| Q8CHS7 | Q99J47 | -0.15 | | | FALSE | TRUE | FALSE | | | | FALSE |
| P19246 | P48678 | -0.15 | | HOM | TRUE | FALSE | FALSE | | | | FALSE |
| P08752 | Q9DC51 | -0.15 | | HET | FALSE | FALSE | FALSE | | | | FALSE |
| Q9D0J4 | Q9WUL7 | -0.15 | HET | | FALSE | FALSE | FALSE | | | | FALSE |
| O35465 | P26883 | -0.15 | HET | HOM | TRUE | FALSE | FALSE | | | | FALSE |
| Q9CQE7 | Q9DC16 | -0.15 | | | FALSE | TRUE | FALSE | | | | FALSE |
| P51880 | Q00915 | -0.15 | | | FALSE | TRUE | FALSE | | | | FALSE |
| Q8BWN8 | Q9QYR9 | -0.15 | | | FALSE | TRUE | FALSE | | | | FALSE |
| P11499 | Q9CQN1 | -0.15 | HOM | HOM | TRUE | FALSE | FALSE | | | | FALSE |
| P62962 | Q9JJV2 | -0.15 | | HOM | TRUE | FALSE | FALSE | | | | TRUE |
| A2AKK5 | Q91X34 | -0.15 | | | FALSE | TRUE | FALSE | | | | FALSE |

Table S4 Mouse

| | | | | | | | | |
|---|---|---|---|---|---|---|---|---|
| E9PZQ0 | P70227 | -0.15 | HET | | FALSE | FALSE | FALSE | FALSE |
| P57780 | Q62261 | -0.15 | | HOM | TRUE | FALSE | FALSE | FALSE |
| P12242 | Q9CR62 | -0.15 | HOM | | TRUE | FALSE | FALSE | FALSE |
| Q62419 | Q9JK48 | -0.14 | | HET | FALSE | FALSE | FALSE | FALSE |
| P30275 | Q6P8J7 | -0.14 | | HOM | TRUE | FALSE | FALSE | FALSE |
| O88952 | Q99L88 | -0.14 | | HOM | TRUE | FALSE | FALSE | FALSE |
| P97772 | Q3UVX5 | -0.14 | HOM | | TRUE | FALSE | FALSE | FALSE |
| P23589 | Q9QZA0 | -0.14 | | | FALSE | TRUE | FALSE | FALSE |
| Q62261 | Q7TPR4 | -0.14 | HOM | | TRUE | FALSE | FALSE | FALSE |
| P28665 | P28666 | -0.14 | | | FALSE | TRUE | FALSE | FALSE |
| P70206 | Q80UG2 | -0.14 | | | FALSE | TRUE | FALSE | FALSE |
| P61750 | Q9WUL7 | -0.14 | | | FALSE | TRUE | FALSE | FALSE |
| Q80UW2 | Q9QZN4 | -0.14 | HET | | FALSE | FALSE | FALSE | FALSE |
| P45376 | Q9DCT1 | -0.13 | HOM | | TRUE | FALSE | FALSE | FALSE |
| P19123 | P20801 | -0.13 | HOM | | TRUE | FALSE | FALSE | FALSE |
| Q99KR7 | Q9ERU9 | -0.13 | | HET | FALSE | FALSE | FALSE | FALSE |
| Q2TPA8 | Q99L04 | -0.13 | | HOM | TRUE | FALSE | FALSE | FALSE |
| Q3UN02 | Q9D1E8 | -0.13 | | | FALSE | TRUE | FALSE | FALSE |
| B2RSH2 | Q9DC51 | -0.13 | HOM | HET | TRUE | FALSE | FALSE | FALSE |
| P37040 | Q9Z0J4 | -0.13 | | | FALSE | TRUE | FALSE | FALSE |
| P39447 | Q9Z0U1 | -0.13 | | HOM | TRUE | FALSE | FALSE | FALSE |
| P70158 | Q04519 | -0.13 | | | FALSE | TRUE | FALSE | FALSE |
| Q00612 | Q8CFX1 | -0.13 | HOM | | TRUE | FALSE | FALSE | FALSE |
| Q62189 | Q9CQI7 | -0.13 | HOM | | TRUE | FALSE | FALSE | FALSE |
| P35293 | Q6PHN9 | -0.12 | | | FALSE | TRUE | FALSE | FALSE |
| P45376 | Q8VC28 | -0.12 | HOM | HOM | TRUE | FALSE | FALSE | FALSE |

Table S4 Mouse

| | | | | | | | | | | | |
|---|---|---|---|---|---|---|---|---|---|---|---|
| P08207 | P50114 | -0.12 | | | FALSE | TRUE | FALSE | | | | FALSE |
| P28661 | P42208 | -0.12 | HET | HOM | TRUE | FALSE | FALSE | | | | FALSE |
| Q3UN02 | Q9D517 | -0.12 | | | FALSE | TRUE | FALSE | | | | FALSE |
| O54916 | Q7TQ48 | -0.12 | HET | HET | FALSE | FALSE | FALSE | | | | FALSE |
| P45377 | Q8VC28 | -0.12 | | HOM | TRUE | FALSE | FALSE | | | | FALSE |
| P11404 | Q00915 | -0.12 | | | FALSE | TRUE | FALSE | | | | FALSE |
| P26048 | P62812 | -0.12 | | | FALSE | TRUE | FALSE | | | | FALSE |
| Q9JME5 | Q9Z1T1 | -0.12 | HET | HET | FALSE | FALSE | TRUE | 59,652 | Human | AP3 adaptor complex, AP3-BLOC1 complex | FALSE |
| P50247 | Q80SW1 | -0.12 | | | FALSE | TRUE | FALSE | | | | FALSE |
| O54829 | Q9QZB1 | -0.12 | | | FALSE | TRUE | FALSE | | | | FALSE |
| Q60823 | Q9WUA6 | -0.12 | | | FALSE | TRUE | FALSE | | | | FALSE |
| P29387 | P62880 | -0.12 | | | FALSE | TRUE | FALSE | | | | FALSE |
| Q8BH59 | Q9QXX4 | -0.12 | | | FALSE | TRUE | FALSE | | | | FALSE |
| P07356 | Q07076 | -0.11 | HOM | | TRUE | FALSE | FALSE | | | | FALSE |
| Q9D2V7 | Q9WUM3 | -0.11 | | HOM | TRUE | FALSE | FALSE | | | | FALSE |
| Q8BHD7 | Q8R081 | -0.11 | | | FALSE | TRUE | FALSE | | | | FALSE |
| P10833 | P32883 | -0.11 | | | FALSE | TRUE | FALSE | | | | FALSE |
| P12382 | Q9WUA3 | -0.11 | HET | HET | FALSE | FALSE | FALSE | | | | FALSE |
| P26040 | P26041 | -0.11 | | | FALSE | TRUE | FALSE | | | | FALSE |
| P70333 | Q8C5Q4 | -0.11 | | | FALSE | TRUE | FALSE | | | | FALSE |
| P20152 | P31001 | -0.11 | | | FALSE | TRUE | FALSE | | | | FALSE |
| P58404 | Q8CIE6 | -0.11 | | HOM | TRUE | FALSE | FALSE | | | | FALSE |
| P04104 | Q6IFZ6 | -0.11 | | | FALSE | TRUE | FALSE | | | | FALSE |
| P61290 | P97372 | -0.11 | | | FALSE | TRUE | FALSE | | | | FALSE |

Table S4 Mouse

| | | | | | | | | | |
|---|---|---|---|---|---|---|---|---|---|
| P00920 | P13634 | -0.11 | HOM | | TRUE | FALSE | FALSE | | FALSE |
| P24547 | Q99L27 | -0.1 | | HOM | TRUE | FALSE | FALSE | | FALSE |
| P36993 | P49443 | -0.1 | | | FALSE | TRUE | FALSE | | FALSE |
| P28656 | Q78ZA7 | -0.1 | | | FALSE | TRUE | FALSE | | FALSE |
| P84078 | Q8BSL7 | -0.1 | | | FALSE | TRUE | FALSE | | FALSE |
| Q8C167 | Q9QUR6 | -0.1 | | | FALSE | TRUE | FALSE | | FALSE |
| O89053 | Q9D2V7 | -0.1 | | | FALSE | TRUE | FALSE | | FALSE |
| P35762 | Q8BJU2 | -0.1 | HOM | | TRUE | FALSE | FALSE | | FALSE |
| Q02053 | Q8C7R4 | -0.1 | | | FALSE | TRUE | FALSE | | FALSE |
| P08551 | P20152 | -0.09 | | | FALSE | TRUE | FALSE | | FALSE |
| Q64518 | Q6PIE5 | -0.09 | | | FALSE | TRUE | FALSE | | FALSE |
| A2ASZ8 | Q8C0K5 | -0.09 | | | FALSE | TRUE | FALSE | | FALSE |
| O08528 | P52792 | -0.09 | | | FALSE | TRUE | FALSE | | FALSE |
| P10107 | P14824 | -0.09 | | | FALSE | TRUE | FALSE | | FALSE |
| Q91V12 | Q9DBK0 | -0.09 | HOM | HOM | TRUE | FALSE | FALSE | | FALSE |
| Q4KWH5 | Q9Z1B3 | -0.09 | | | FALSE | TRUE | FALSE | | FALSE |
| P39447 | Q9JLB0 | -0.09 | | | FALSE | TRUE | FALSE | | FALSE |
| P45377 | Q9DCT1 | -0.09 | | | FALSE | TRUE | FALSE | | FALSE |
| Q31125 | Q5FWH7 | -0.09 | | | FALSE | TRUE | FALSE | | FALSE |
| Q91VT4 | Q91X52 | -0.09 | HOM | HOM | TRUE | FALSE | FALSE | | FALSE |
| P02463 | Q9QZS0 | -0.08 | | | FALSE | TRUE | FALSE | | FALSE |
| P06800 | Q64487 | -0.08 | | | FALSE | TRUE | FALSE | | FALSE |
| Q62433 | Q9QYG0 | -0.08 | | HOM | TRUE | FALSE | FALSE | | FALSE |
| Q3TCH7 | Q9D4H8 | -0.08 | HOM, HET | HET | TRUE, FALSE | FALSE | FALSE | | FALSE |
| Q64176 | Q8VCC2 | -0.08 | | | FALSE | TRUE | FALSE | | FALSE |

Table S4 Mouse

| | | | | | | | | |
|---|---|---|---|---|---|---|---|---|
| Q80X90 | Q8VHX6 | -0.08 | | HOM | TRUE | FALSE | FALSE | FALSE |
| Q61234 | Q8JZS0 | -0.08 | HOM | | TRUE | FALSE | FALSE | FALSE |
| P14246 | P17809 | -0.08 | | | FALSE | TRUE | FALSE | FALSE |
| O70433 | P97447 | -0.08 | | | FALSE | TRUE | FALSE | FALSE |
| O70433 | Q3TJD7 | -0.08 | | | FALSE | TRUE | FALSE | FALSE |
| P14246 | P32037 | -0.08 | | | FALSE | TRUE | FALSE | FALSE |
| Q925N2 | Q99JR1 | -0.08 | | | FALSE | TRUE | FALSE | FALSE |
| P25911 | Q04736 | -0.08 | | | FALSE | TRUE | FALSE | FALSE |
| O35114 | Q08857 | -0.08 | | HET | FALSE | FALSE | FALSE | FALSE |
| P06151 | P16125 | -0.08 | | | FALSE | TRUE | FALSE | FALSE |
| P70290 | Q9Z0U1 | -0.08 | HOM | HOM | TRUE | FALSE | FALSE | FALSE |
| A2ASZ8 | Q8R0Y8 | -0.07 | | | FALSE | TRUE | FALSE | FALSE |
| Q8BHD7 | Q921F4 | -0.07 | | | FALSE | TRUE | FALSE | FALSE |
| O88876 | Q8CHS7 | -0.07 | | | FALSE | TRUE | FALSE | FALSE |
| P39447 | Q91XM9 | -0.07 | | HOM | TRUE | FALSE | FALSE | FALSE |
| O35954 | P53811 | -0.07 | | | FALSE | TRUE | FALSE | FALSE |
| P11881 | P70227 | -0.07 | | | FALSE | TRUE | FALSE | FALSE |
| P51667 | Q3THE2 | -0.07 | | | FALSE | TRUE | FALSE | FALSE |
| Q31125 | Q6P5F6 | -0.07 | | | FALSE | TRUE | FALSE | FALSE |
| P21619 | P46660 | -0.07 | | | FALSE | TRUE | FALSE | FALSE |
| P56501 | Q9CR62 | -0.07 | | | FALSE | TRUE | FALSE | FALSE |
| P48036 | P97384 | -0.07 | | | FALSE | TRUE | FALSE | FALSE |
| P40237 | P40240 | -0.07 | | HOM | TRUE | FALSE | FALSE | FALSE |
| P08551 | P21619 | -0.07 | | | FALSE | TRUE | FALSE | FALSE |
| Q3TCH7 | Q9WTX6 | -0.07 | HOM, HET | HOM, HET | TRUE, FALSE | FALSE | FALSE | FALSE |

Table S4 Mouse

| | | | | | | | | | | | |
|---|---|---|---|---|---|---|---|---|---|---|---|
| P54285 | Q8R0S4 | -0.07 | | | FALSE | TRUE | FALSE | | | | FALSE |
| P13634 | P28651 | -0.06 | | | FALSE | TRUE | FALSE | | | | FALSE |
| Q3UNX5 | Q99NB1 | -0.06 | | HOM | TRUE | FALSE | FALSE | | | | FALSE |
| Q62393 | Q9CYZ2 | -0.06 | | | FALSE | TRUE | FALSE | | | | FALSE |
| O55028 | Q8BFP9 | -0.06 | | HOM | TRUE | FALSE | FALSE | | | | FALSE |
| P63318 | P70268 | -0.06 | | HET | FALSE | FALSE | FALSE | | | | FALSE |
| P18572 | P97300 | -0.06 | HOM | | TRUE | FALSE | FALSE | | | | FALSE |
| A2AQ07 | Q9D6F9 | -0.06 | HET | HET | FALSE | FALSE | FALSE | | | | FALSE |
| O70571 | Q922H2 | -0.06 | | | FALSE | TRUE | FALSE | | | | FALSE |
| P12960 | Q61330 | -0.05 | HET | | FALSE | FALSE | FALSE | | | | FALSE |
| P50171 | Q91X52 | -0.05 | HOM | HOM | TRUE | FALSE | FALSE | | | | FALSE |
| O08917 | Q60634 | -0.05 | | | FALSE | TRUE | FALSE | | | | FALSE |
| E9Q557 | Q9QXS1 | -0.05 | HOM | HOM | TRUE | FALSE | FALSE | | | | FALSE |
| O08599 | Q60770 | -0.05 | | | FALSE | TRUE | FALSE | | | | FALSE |
| Q60902 | Q9Z0R4 | -0.05 | HET | HET | FALSE | FALSE | FALSE | | | | FALSE |
| P50114 | P50543 | -0.05 | | | FALSE | TRUE | FALSE | | | | FALSE |
| O54916 | Q9QXY6 | -0.05 | HET | HET | FALSE | FALSE | FALSE | | | | FALSE |
| Q61234 | Q99L88 | -0.05 | HOM | HOM | TRUE | FALSE | TRUE | 349 | Mouse | Sarcoglycan-sarcospan-syntrophin-dystrobrevin complex | FALSE |
| Q7TSQ8 | Q9DBT9 | -0.05 | | | FALSE | TRUE | FALSE | | | | FALSE |
| O55106 | Q8CIE6 | -0.05 | | HOM | TRUE | FALSE | FALSE | | | | FALSE |
| P19157 | Q80W21 | -0.05 | HOM | HOM | TRUE | FALSE | FALSE | | | | FALSE |
| Q62452 | Q64435 | -0.05 | | | FALSE | TRUE | FALSE | | | | FALSE |
| P00920 | Q9QZA0 | -0.04 | HOM | | TRUE | FALSE | FALSE | | | | FALSE |

Table S4 Mouse

| | | | | | | | | |
|---|---|---|---|---|---|---|---|---|
| P21440 | Q9QY30 | -0.04 | | | FALSE | TRUE | FALSE | FALSE |
| P05064 | Q91Y97 | -0.04 | | | FALSE | TRUE | FALSE | FALSE |
| P68404 | P70268 | -0.04 | | HET | FALSE | FALSE | FALSE | FALSE |
| O08529 | O35350 | -0.04 | HOM | HET | TRUE | FALSE | FALSE | FALSE |
| P20108 | P35700 | -0.04 | HOM | HOM | TRUE | FALSE | FALSE | FALSE |
| O88545 | P26516 | -0.04 | | | FALSE | TRUE | FALSE | FALSE |
| O35639 | Q07076 | -0.04 | | | FALSE | TRUE | FALSE | FALSE |
| Q8K0T0 | Q9ES97 | -0.04 | | | FALSE | TRUE | FALSE | FALSE |
| Q61699 | Q9JKR6 | -0.04 | HOM | HOM | TRUE | FALSE | FALSE | FALSE |
| P24547 | Q9DCZ1 | -0.03 | | | FALSE | TRUE | FALSE | FALSE |
| Q3UNZ8 | Q62465 | -0.03 | | | FALSE | TRUE | FALSE | FALSE |
| P07310 | P30275 | -0.03 | | | FALSE | TRUE | FALSE | FALSE |
| P03995 | P14733 | -0.03 | | | FALSE | TRUE | FALSE | FALSE |
| P10649 | P48774 | -0.03 | | | FALSE | TRUE | FALSE | FALSE |
| Q8BTG7 | Q9QYG0 | -0.03 | | HOM | TRUE | FALSE | FALSE | FALSE |
| A2AKK5 | Q8BGG9 | -0.03 | | | FALSE | TRUE | FALSE | FALSE |
| Q62465 | Q9DCS3 | -0.03 | | HOM | TRUE | FALSE | FALSE | FALSE |
| O35367 | P51885 | -0.03 | | | FALSE | TRUE | FALSE | FALSE |
| Q6P8X1 | Q9D8U8 | -0.03 | | HOM | TRUE | FALSE | FALSE | FALSE |
| P47199 | Q80TB8 | -0.03 | HOM | HOM | TRUE | FALSE | FALSE | FALSE |
| P53395 | Q8BMF4 | -0.03 | | HOM | TRUE | FALSE | FALSE | FALSE |
| P46735 | Q9WTI7 | -0.03 | | HOM | TRUE | FALSE | FALSE | FALSE |
| P41216 | Q91WC3 | -0.03 | | | FALSE | TRUE | FALSE | FALSE |
| Q4LDG0 | Q91VE0 | -0.02 | | | FALSE | TRUE | FALSE | FALSE |
| Q99NB1 | Q9QXG4 | -0.02 | HOM | | TRUE | FALSE | FALSE | FALSE |
| Q9Z210 | Q9Z211 | -0.02 | | | FALSE | TRUE | FALSE | FALSE |

Table S4 Mouse

| | | | | | | | | |
|---|---|---|---|---|---|---|---|---|
| Q8BH44 | Q9WUM3 | -0.02 | | HOM | TRUE | FALSE | FALSE | FALSE |
| Q91X34 | Q9QYR9 | -0.02 | | | FALSE | TRUE | FALSE | FALSE |
| P84078 | Q9D0J4 | -0.02 | | HET | FALSE | FALSE | FALSE | FALSE |
| Q8BGF9 | Q8BH59 | -0.02 | | | FALSE | TRUE | FALSE | FALSE |
| Q9D1E8 | Q9D517 | -0.02 | | | FALSE | TRUE | FALSE | FALSE |
| O35367 | Q9JK53 | -0.02 | | | FALSE | TRUE | FALSE | FALSE |
| Q8QZR3 | Q8VCC2 | -0.02 | | | FALSE | TRUE | FALSE | FALSE |
| O70443 | P08752 | -0.01 | | | FALSE | TRUE | FALSE | FALSE |
| O08989 | P32883 | -0.01 | | | FALSE | TRUE | FALSE | FALSE |
| Q8BMD8 | Q8R0Y8 | -0.01 | | | FALSE | TRUE | FALSE | FALSE |
| P97371 | P97372 | -0.01 | | | FALSE | TRUE | FALSE | TRUE |
| Q9WUM3 | Q9WUM4 | -0.01 | HOM | HOM | TRUE | FALSE | FALSE | FALSE |
| Q91VA0 | Q99NB1 | -0.01 | | HOM | TRUE | FALSE | FALSE | FALSE |
| O35367 | Q99MQ4 | -0.01 | | | FALSE | TRUE | FALSE | FALSE |
| O88951 | Q61234 | -0.01 | HOM | HOM | TRUE | FALSE | FALSE | FALSE |
| Q91YW3 | Q9QYI3 | -0.01 | | | FALSE | TRUE | FALSE | FALSE |
| Q5SQM0 | Q8BQM8 | -0.01 | | | FALSE | TRUE | FALSE | FALSE |
| O08788 | Q922J3 | -0.01 | HOM | | TRUE | FALSE | FALSE | FALSE |
| Q8R5J9 | Q9JIG8 | -0 | | | FALSE | TRUE | FALSE | FALSE |
| P08551 | P14733 | -0 | | | FALSE | TRUE | FALSE | FALSE |
| P43274 | P43276 | -0 | | | FALSE | TRUE | FALSE | FALSE |
| P23589 | Q9WVT6 | 0.001 | | | FALSE | TRUE | FALSE | FALSE |
| P61750 | P84078 | 0.001 | | | FALSE | TRUE | FALSE | FALSE |
| P56501 | Q9QZD8 | 0.002 | | | FALSE | TRUE | FALSE | FALSE |
| P00920 | P28651 | 0.002 | HOM | | TRUE | FALSE | FALSE | FALSE |
| Q8VIJ6 | Q99K48 | 0.004 | HOM | | TRUE | FALSE | FALSE | FALSE |

Table S4 Mouse

| | | | | | | | | |
|---|---|---|---|---|---|---|---|---|
| Q60902 | Q9WVK4 | 0.008 | HET | HET | FALSE | FALSE | FALSE | FALSE |
| O35855 | P24288 | 0.008 | HOM | HOM | TRUE | FALSE | FALSE | FALSE |
| P47199 | Q62465 | 0.009 | HOM | | TRUE | FALSE | FALSE | FALSE |
| Q3B7Z2 | Q91XL9 | 0.011 | HOM | | TRUE | FALSE | FALSE | FALSE |
| P54869 | Q8JZK9 | 0.011 | HOM | HOM | TRUE | FALSE | FALSE | FALSE |
| P62746 | Q62159 | 0.012 | | | FALSE | TRUE | FALSE | FALSE |
| P14231 | P97370 | 0.012 | | | FALSE | TRUE | FALSE | FALSE |
| Q6A065 | Q80U49 | 0.013 | | | FALSE | TRUE | FALSE | FALSE |
| O54991 | Q6P9K9 | 0.014 | | | FALSE | TRUE | FALSE | FALSE |
| P14824 | P48036 | 0.014 | | | FALSE | TRUE | FALSE | FALSE |
| Q6GYP7 | Q8C0T5 | 0.015 | HET | | FALSE | FALSE | FALSE | FALSE |
| P12242 | Q9QZD8 | 0.016 | HOM | | TRUE | FALSE | FALSE | FALSE |
| E9PUL5 | Q8C838 | 0.017 | | | FALSE | TRUE | FALSE | FALSE |
| P51667 | P97457 | 0.017 | | | FALSE | TRUE | FALSE | FALSE |
| P40240 | Q8R3G9 | 0.017 | HOM | | TRUE | FALSE | FALSE | FALSE |
| Q8BGF9 | Q9DB41 | 0.017 | | | FALSE | TRUE | FALSE | FALSE |
| P62331 | Q9CQW2 | 0.017 | HOM | | TRUE | FALSE | FALSE | FALSE |
| O08808 | Q8BPM0 | 0.017 | HOM | HOM | TRUE | FALSE | FALSE | FALSE |
| P16054 | P70268 | 0.017 | | HET | FALSE | FALSE | FALSE | FALSE |
| Q60766 | Q9QZ85 | 0.018 | | | FALSE | TRUE | FALSE | FALSE |
| P55258 | P61028 | 0.018 | | | FALSE | TRUE | FALSE | FALSE |
| Q62188 | Q9EQF5 | 0.019 | HET | HOM | TRUE | FALSE | FALSE | FALSE |
| P50171 | Q8JZV9 | 0.019 | HOM | HOM | TRUE | FALSE | FALSE | FALSE |
| P03995 | P31001 | 0.02 | | | FALSE | TRUE | FALSE | FALSE |
| Q8C129 | Q9EQH2 | 0.021 | | HET | FALSE | FALSE | FALSE | FALSE |
| Q6KAR6 | Q8BI71 | 0.021 | | | FALSE | TRUE | FALSE | FALSE |

Table S4 Mouse

| | | | | | | | | | | | |
|---|---|---|---|---|---|---|---|---|---|---|---|
| Q7TNG5 | Q8BQM8 | 0.021 | | | FALSE | TRUE | FALSE | | | | FALSE |
| Q8CFA2 | Q99LB7 | 0.022 | HOM | | TRUE | FALSE | FALSE | | | | FALSE |
| O35114 | Q61009 | 0.023 | | | FALSE | TRUE | FALSE | | | | FALSE |
| P04104 | Q3TTY5 | 0.023 | | | FALSE | TRUE | FALSE | | | | FALSE |
| P63321 | Q9JIW9 | 0.024 | | HET | FALSE | FALSE | FALSE | | | | FALSE |
| O55143 | Q6PIC6 | 0.025 | | | FALSE | TRUE | FALSE | | | | FALSE |
| O70433 | Q9R059 | 0.025 | | | FALSE | TRUE | TRUE | 3187 | Human | FHL2-FHL3 complex | FALSE |
| P16406 | Q8C129 | 0.026 | | | FALSE | TRUE | FALSE | | | | FALSE |
| P53395 | Q8BKZ9 | 0.026 | | HET | FALSE | FALSE | FALSE | | | | FALSE |
| P17182 | P17183 | 0.027 | HOM | HOM | TRUE | FALSE | FALSE | | | | FALSE |
| P28651 | Q9QZA0 | 0.027 | | | FALSE | TRUE | FALSE | | | | FALSE |
| P29341 | P70372 | 0.028 | | HOM | TRUE | FALSE | FALSE | | | | FALSE |
| P13634 | Q64444 | 0.029 | | HOM | TRUE | FALSE | FALSE | | | | FALSE |
| P68373 | Q9JJZ2 | 0.029 | | HET | FALSE | FALSE | FALSE | | | | FALSE |
| Q8BJU2 | Q8R3G9 | 0.031 | | | FALSE | TRUE | FALSE | | | | FALSE |
| P19246 | P20152 | 0.031 | | | FALSE | TRUE | FALSE | | | | FALSE |
| O08532 | Q6PHS9 | 0.031 | | | FALSE | TRUE | FALSE | | | | FALSE |
| P08551 | P31001 | 0.032 | | | FALSE | TRUE | FALSE | | | | FALSE |
| Q7TME0 | Q99JY8 | 0.032 | | | FALSE | TRUE | FALSE | | | | FALSE |
| Q5FWH7 | Q6P5F6 | 0.032 | | | FALSE | TRUE | FALSE | | | | FALSE |
| P97822 | Q9EST5 | 0.033 | | | FALSE | TRUE | FALSE | | | | FALSE |
| Q9R1Q8 | Q9WVA4 | 0.034 | | | FALSE | TRUE | FALSE | | | | FALSE |
| P60843 | Q91VC3 | 0.035 | HET | HET | FALSE | FALSE | FALSE | | | | FALSE |
| O09164 | P08228 | 0.036 | HOM | HOM | TRUE | FALSE | FALSE | | | | FALSE |
| F6SEU4 | Q9Z268 | 0.036 | | | FALSE | TRUE | FALSE | | | | TRUE |

Table S4 Mouse

| | | | | | | | | | | | |
|---|---|---|---|---|---|---|---|---|---|---|---|
| O08663 | P50580 | 0.037 | | | FALSE | TRUE | FALSE | | | | FALSE |
| P70206 | Q8CJH3 | 0.037 | | HOM | TRUE | FALSE | TRUE | 5791 | Mouse | PlexinA1-PlexinB1 complex | FALSE |
| Q64444 | Q9QZA0 | 0.037 | HOM | | TRUE | FALSE | FALSE | | | | FALSE |
| P26049 | P62812 | 0.038 | | | FALSE | TRUE | FALSE | | | | TRUE |
| P12960 | Q810U4 | 0.042 | HET | | FALSE | FALSE | FALSE | | | | FALSE |
| Q7TPR4 | Q9JI91 | 0.043 | | HOM | TRUE | FALSE | FALSE | | | | FALSE |
| O35136 | Q80Z24 | 0.044 | HOM | | TRUE | FALSE | FALSE | | | | FALSE |
| P70175 | Q811D0 | 0.044 | HOM | HOM | TRUE | FALSE | FALSE | | | | FALSE |
| Q8BFP9 | Q922H2 | 0.045 | HOM | | TRUE | FALSE | FALSE | | | | FALSE |
| O89053 | Q8BH44 | 0.045 | | | FALSE | TRUE | FALSE | | | | FALSE |
| P58021 | Q8BH24 | 0.046 | | | FALSE | TRUE | FALSE | | | | FALSE |
| Q924X2 | Q9DC50 | 0.046 | | | FALSE | TRUE | FALSE | | | | FALSE |
| O88533 | P48320 | 0.046 | HOM | | TRUE | FALSE | FALSE | | | | FALSE |
| Q8CIE6 | Q9ERG2 | 0.048 | HOM | | TRUE | FALSE | FALSE | | | | FALSE |
| P70695 | Q9QXD6 | 0.048 | | | FALSE | TRUE | FALSE | | | | FALSE |
| Q3UQ44 | Q9JKF1 | 0.05 | | HOM | TRUE | FALSE | FALSE | | | | FALSE |
| Q811D0 | Q9Z0U1 | 0.051 | HOM | HOM | TRUE | FALSE | FALSE | | | | FALSE |
| P39447 | P70290 | 0.052 | | HOM | TRUE | FALSE | FALSE | | | | FALSE |
| Q3V009 | Q9CXE7 | 0.053 | | | FALSE | TRUE | FALSE | | | | FALSE |
| P62748 | Q8BNY6 | 0.053 | HET | HET | FALSE | FALSE | FALSE | | | | FALSE |
| Q02566 | Q5SX40 | 0.054 | | | FALSE | TRUE | FALSE | | | | FALSE |
| O54916 | Q8BH64 | 0.054 | HET | HOM, HET | TRUE, FALSE | FALSE | FALSE | | | | FALSE |
| Q9CQ62 | Q9WV68 | 0.056 | HOM | | TRUE | FALSE | FALSE | | | | FALSE |
| Q3UJU9 | Q8BSE0 | 0.056 | | | FALSE | TRUE | FALSE | | | | FALSE |
| Q8VCB3 | Q9Z1E4 | 0.056 | HOM | HOM | TRUE | FALSE | FALSE | | | | FALSE |

Table S4 Mouse

| | | | | | | | | | | | |
|---|---|---|---|---|---|---|---|---|---|---|---|
| O70318 | P48193 | 0.056 | | | FALSE | TRUE | FALSE | | | | FALSE |
| O35639 | P10107 | 0.056 | | | FALSE | TRUE | FALSE | | | | FALSE |
| P97449 | Q9EQH2 | 0.057 | | HET | FALSE | FALSE | FALSE | | | | FALSE |
| P20152 | P46660 | 0.057 | | | FALSE | TRUE | FALSE | | | | FALSE |
| A2AQ07 | P68372 | 0.058 | HET | HET | FALSE | FALSE | FALSE | | | | FALSE |
| P31786 | Q5XG73 | 0.058 | | | FALSE | TRUE | FALSE | | | | FALSE |
| O54916 | Q9Z0R4 | 0.059 | HET | HET | FALSE | FALSE | FALSE | | | | FALSE |
| P36371 | Q9JI39 | 0.059 | | HOM | TRUE | FALSE | FALSE | | | | FALSE |
| O35367 | P28654 | 0.059 | | | FALSE | TRUE | FALSE | | | | FALSE |
| P60766 | Q05144 | 0.06 | HET | HET | FALSE | FALSE | FALSE | | | | FALSE |
| P20357 | P27546 | 0.06 | | | FALSE | TRUE | FALSE | | | | FALSE |
| P16546 | P57780 | 0.061 | HOM | | TRUE | FALSE | FALSE | | | | FALSE |
| P63213 | Q80SZ7 | 0.064 | HET | | FALSE | FALSE | FALSE | | | | FALSE |
| P68368 | Q9JJZ2 | 0.064 | HET | HET | FALSE | FALSE | FALSE | | | | FALSE |
| P51859 | Q3UMU 9 | 0.065 | HOM | | TRUE | FALSE | FALSE | | | | FALSE |
| P61205 | Q9D0J4 | 0.065 | | HET | FALSE | FALSE | FALSE | | | | FALSE |
| D3Z7P3 | Q571F8 | 0.067 | HOM | | TRUE | FALSE | FALSE | | | | FALSE |
| P16406 | P97449 | 0.068 | | | FALSE | TRUE | FALSE | | | | FALSE |
| P21278 | P21279 | 0.069 | | | FALSE | TRUE | TRUE | 117 | Human | GPR56-CD81-Galpha-Gbeta complex | FALSE |
| P07356 | P10107 | 0.07 | HOM | | TRUE | FALSE | FALSE | | | | FALSE |
| Q8BMD 8 | Q8C0K5 | 0.071 | | | FALSE | TRUE | FALSE | | | | FALSE |
| O35367 | P28653 | 0.071 | | | FALSE | TRUE | FALSE | | | | FALSE |
| Q7TQ48 | Q9WVK 4 | 0.071 | HET | HET | FALSE | FALSE | FALSE | | | | FALSE |
| P62880 | Q8CGF6 | 0.072 | | | FALSE | TRUE | FALSE | | | | FALSE |

Table S4 Mouse

| | | | | | | | | |
|---|---|---|---|---|---|---|---|---|
| P62331 | P84084 | 0.072 | HOM | | TRUE | FALSE | FALSE | FALSE |
| Q4KWH5 | Q8K394 | 0.072 | | | FALSE | TRUE | FALSE | FALSE |
| Q5SUC9 | Q8VCL2 | 0.073 | | | FALSE | TRUE | FALSE | FALSE |
| P97772 | Q9QYS2 | 0.074 | HOM | | TRUE | FALSE | FALSE | FALSE |
| P00920 | Q64444 | 0.074 | HOM | HOM | TRUE | FALSE | FALSE | FALSE |
| Q8VHQ9 | Q9DBK0 | 0.074 | HOM | HOM | TRUE | FALSE | FALSE | FALSE |
| Q8BSL7 | Q9WUL7 | 0.076 | | | FALSE | TRUE | FALSE | FALSE |
| B2RXS4 | P70206 | 0.079 | HET | | FALSE | FALSE | FALSE | FALSE |
| O88876 | Q6GV12 | 0.079 | | | FALSE | TRUE | FALSE | FALSE |
| Q5PR73 | Q80ZJ1 | 0.079 | | HOM | TRUE | FALSE | FALSE | FALSE |
| Q8BGD9 | Q9WUK2 | 0.079 | | | FALSE | TRUE | FALSE | FALSE |
| P61164 | P68134 | 0.08 | | | FALSE | TRUE | FALSE | FALSE |
| P26048 | P26049 | 0.081 | | | FALSE | TRUE | FALSE | FALSE |
| P08752 | P21278 | 0.081 | | | FALSE | TRUE | FALSE | FALSE |
| Q8C0L0 | Q8VBT0 | 0.081 | | | FALSE | TRUE | FALSE | FALSE |
| O35639 | P97429 | 0.082 | | | FALSE | TRUE | FALSE | FALSE |
| Q8K021 | Q9JKD3 | 0.082 | | | FALSE | TRUE | FALSE | FALSE |
| Q6P9K9 | Q9CS84 | 0.083 | | HET | FALSE | FALSE | FALSE | FALSE |
| P56371 | Q91ZR1 | 0.083 | HET | | FALSE | FALSE | FALSE | FALSE |
| P14069 | P50114 | 0.084 | | | FALSE | TRUE | FALSE | FALSE |
| Q80SZ7 | Q9CXP8 | 0.084 | | | FALSE | TRUE | FALSE | FALSE |
| Q60854 | Q9D154 | 0.084 | | | FALSE | TRUE | FALSE | FALSE |
| F8VPU2 | P26040 | 0.084 | | | FALSE | TRUE | FALSE | FALSE |
| P63216 | Q80SZ7 | 0.084 | | | FALSE | TRUE | FALSE | FALSE |
| P62071 | Q61411 | 0.085 | | | FALSE | TRUE | FALSE | FALSE |
| P50247 | Q68FL4 | 0.085 | | | FALSE | TRUE | FALSE | FALSE |

Table S4 Mouse

| | | | | | | | | | | | |
|---|---|---|---|---|---|---|---|---|---|---|---|
| P62881 | Q8CGF6 | 0.086 | HET | | FALSE | FALSE | FALSE | | | | FALSE |
| P13542 | Q8VDD5 | 0.086 | | | FALSE | TRUE | FALSE | | | | FALSE |
| P46638 | P56371 | 0.086 | | HET | FALSE | FALSE | FALSE | | | | FALSE |
| P10107 | P48036 | 0.086 | | | FALSE | TRUE | FALSE | | | | FALSE |
| Q80ZJ1 | Q99JI6 | 0.087 | HOM | | TRUE | FALSE | FALSE | | | | FALSE |
| P22723 | P26049 | 0.087 | | | FALSE | TRUE | FALSE | | | | FALSE |
| P47934 | Q9DC50 | 0.087 | | | FALSE | TRUE | FALSE | | | | FALSE |
| Q8VCH0 | Q921H8 | 0.089 | | HOM | TRUE | FALSE | FALSE | | | | FALSE |
| P12960 | Q810U3 | 0.089 | HET | | FALSE | FALSE | FALSE | | | | FALSE |
| P62259 | Q9CQV8 | 0.089 | HOM | | TRUE | FALSE | TRUE | 5199,5613 | Human | Kinase maturation complex 1,Emerin complex 25 | TRUE |
| Q8R2Y2 | Q9R069 | 0.09 | | | FALSE | TRUE | FALSE | | | | FALSE |
| P08752 | P27601 | 0.091 | | | FALSE | TRUE | FALSE | | | | FALSE |
| Q8BH44 | Q9D2V7 | 0.093 | | | FALSE | TRUE | FALSE | | | | FALSE |
| Q63880 | Q8VCC2 | 0.095 | | | FALSE | TRUE | FALSE | | | | FALSE |
| P50153 | Q80SZ7 | 0.095 | | | FALSE | TRUE | FALSE | | | | FALSE |
| P26231 | Q64727 | 0.095 | HOM | HET | TRUE | FALSE | FALSE | | | | FALSE |
| Q91V61 | Q925N0 | 0.095 | | | FALSE | TRUE | FALSE | | | | FALSE |
| O70503 | Q8BTX9 | 0.097 | | | FALSE | TRUE | FALSE | | | | FALSE |
| Q8K023 | Q9JII6 | 0.097 | HOM | | TRUE | FALSE | FALSE | | | | FALSE |
| Q6GV12 | Q9EQ06 | 0.098 | | | FALSE | TRUE | FALSE | | | | FALSE |
| P07758 | P22599 | 0.099 | HOM | HOM | TRUE | FALSE | FALSE | | | | FALSE |
| Q8VDD5 | Q91Z83 | 0.099 | | | FALSE | TRUE | FALSE | | | | FALSE |
| Q5SSM3 | Q91Z67 | 0.101 | | HET | FALSE | FALSE | FALSE | | | | FALSE |

Table S4 Mouse

| | | | | | | | | | | | |
|---|---|---|---|---|---|---|---|---|---|---|---|
| P48771 | Q61387 | 0.102 | | | FALSE | TRUE | FALSE | | | | FALSE |
| P24527 | Q8VCT3 | 0.103 | | | FALSE | TRUE | FALSE | | | | FALSE |
| Q91XD7 | Q9CYA0 | 0.103 | | | FALSE | TRUE | FALSE | | | | FALSE |
| P47791 | Q9JMH6 | 0.104 | HOM | HOM | TRUE | FALSE | FALSE | | | | FALSE |
| P62880 | P62881 | 0.105 | | HET | FALSE | FALSE | FALSE | | | | FALSE |
| P61205 | P61750 | 0.106 | | | FALSE | TRUE | FALSE | | | | FALSE |
| Q64518 | Q6PIC6 | 0.107 | | | FALSE | TRUE | FALSE | | | | FALSE |
| P08752 | P21279 | 0.107 | | | FALSE | TRUE | FALSE | | | | FALSE |
| Q9Z0V7 | Q9Z0V8 | 0.108 | | | FALSE | TRUE | FALSE | | | | FALSE |
| P20152 | P48678 | 0.108 | | HOM | TRUE | FALSE | FALSE | | | | FALSE |
| P29387 | P62874 | 0.109 | | HET | FALSE | FALSE | FALSE | | | | FALSE |
| O08539 | Q7TQF7 | 0.109 | | | FALSE | TRUE | FALSE | | | | FALSE |
| P07934 | Q6PGN3 | 0.11 | HET | | FALSE | FALSE | FALSE | | | | FALSE |
| Q7TQ48 | Q8BH64 | 0.11 | HET | HOM, HET | TRUE, FALSE | FALSE | FALSE | | | | FALSE |
| P63037 | Q99KV1 | 0.111 | HOM | HOM | TRUE | FALSE | FALSE | | | | FALSE |
| P14733 | P48678 | 0.114 | | HOM | TRUE | FALSE | TRUE | 5608,5611 | Human | Emerin architectural complex, Emerin complex 24 | TRUE |
| Q8BTZ7 | Q922H4 | 0.114 | | | FALSE | TRUE | FALSE | | | | FALSE |
| P08551 | P19246 | 0.115 | | | FALSE | TRUE | FALSE | | | | TRUE |
| O88951 | O88952 | 0.117 | HOM | | TRUE | FALSE | FALSE | | | | FALSE |
| A2AQ07 | P99024 | 0.118 | HET | HET | FALSE | FALSE | FALSE | | | | FALSE |
| P35762 | P40240 | 0.12 | HOM | HOM | TRUE | FALSE | FALSE | | | | FALSE |
| O88990 | Q9JI91 | 0.12 | HOM | HOM | TRUE | FALSE | FALSE | | | | FALSE |
| Q6PA06 | Q8BH66 | 0.12 | | HOM | TRUE | FALSE | FALSE | | | | FALSE |

| | | | | | | | | | | | |
|---|---|---|---|---|---|---|---|---|---|---|---|
| P17742 | Q9ERU9 | 0.121 | | HET | FALSE | FALSE | FALSE | | | | FALSE |
| Q00915 | Q05816 | 0.122 | | | FALSE | TRUE | FALSE | | | | FALSE |
| P16388 | P63141 | 0.122 | HET | HET | FALSE | FALSE | FALSE | | | | TRUE |
| O08807 | P20108 | 0.122 | HOM | HOM | TRUE | FALSE | FALSE | | | | FALSE |
| Q8BR92 | Q9Z0P4 | 0.123 | | | FALSE | TRUE | FALSE | | | | FALSE |
| Q501J6 | Q810A7 | 0.123 | | | FALSE | TRUE | FALSE | | | | FALSE |
| Q8BJU2 | Q922J6 | 0.123 | | | FALSE | TRUE | FALSE | | | | FALSE |
| P10107 | Q07076 | 0.124 | | | FALSE | TRUE | FALSE | | | | FALSE |
| P40240 | Q8BJU2 | 0.125 | HOM | | TRUE | FALSE | FALSE | | | | FALSE |
| P21278 | P27601 | 0.126 | | | FALSE | TRUE | FALSE | | | | FALSE |
| O88844 | P54071 | 0.126 | HOM | HOM | TRUE | FALSE | FALSE | | | | FALSE |
| O35728 | Q9EP75 | 0.126 | | | FALSE | TRUE | FALSE | | | | FALSE |
| Q61644 | Q99JB8 | 0.127 | HET | HOM | TRUE | FALSE | FALSE | | | | FALSE |
| Q8R464 | Q8R5M8 | 0.129 | HOM | | TRUE | FALSE | FALSE | | | | FALSE |
| O89053 | Q9WUM4 | 0.13 | | HOM | TRUE | FALSE | FALSE | | | | FALSE |
| Q14BI2 | Q3UVX5 | 0.131 | | | FALSE | TRUE | FALSE | | | | FALSE |
| P53994 | P56371 | 0.131 | | HET | FALSE | FALSE | FALSE | | | | FALSE |
| P11679 | Q8VED5 | 0.131 | | | FALSE | TRUE | FALSE | | | | FALSE |
| P62874 | Q8CGF6 | 0.132 | HET | | FALSE | FALSE | FALSE | | | | FALSE |
| P62331 | P84078 | 0.132 | HOM | | TRUE | FALSE | FALSE | | | | FALSE |
| Q6PHS9 | Q9Z1L5 | 0.132 | | | FALSE | TRUE | FALSE | | | | FALSE |
| Q80TL0 | Q8CGA0 | 0.134 | | | FALSE | TRUE | FALSE | | | | FALSE |
| Q8R0F9 | Q99J08 | 0.134 | HOM | HOM | TRUE | FALSE | FALSE | | | | FALSE |
| P70175 | P70290 | 0.134 | HOM | HOM | TRUE | FALSE | FALSE | | | | FALSE |
| Q91VN4 | Q9CRB9 | 0.134 | | | FALSE | TRUE | TRUE | 6249,6255 | Human | MIB complex, MICOS complex | FALSE |

Table S4 Mouse

| | | | | | | | | |
|---|---|---|---|---|---|---|---|---|
| P42208 | Q9Z2Q6 | 0.135 | HOM | HET | TRUE | FALSE | FALSE | FALSE |
| Q8VC28 | Q9JII6 | 0.135 | HOM | | TRUE | FALSE | FALSE | FALSE |
| Q8BXK9 | Q9Z1Q5 | 0.136 | | HOM | TRUE | FALSE | FALSE | FALSE |
| Q8JZV9 | Q91X52 | 0.137 | HOM | HOM | TRUE | FALSE | FALSE | FALSE |
| O55106 | Q9ERG2 | 0.137 | | | FALSE | TRUE | FALSE | FALSE |
| Q8CFJ7 | Q9Z2Z6 | 0.138 | | | FALSE | TRUE | FALSE | FALSE |
| Q8CEE7 | Q9QYF1 | 0.138 | | | FALSE | TRUE | FALSE | FALSE |
| Q8VCC2 | Q91WG0 | 0.139 | | | FALSE | TRUE | FALSE | FALSE |
| P39053 | P39054 | 0.139 | HOM | | TRUE | FALSE | FALSE | FALSE |
| Q8BSE0 | Q9DCV4 | 0.141 | | | FALSE | TRUE | FALSE | FALSE |
| P13412 | Q9WUZ5 | 0.141 | | | FALSE | TRUE | FALSE | FALSE |
| Q8BG51 | Q8JZN7 | 0.142 | | | FALSE | TRUE | FALSE | FALSE |
| Q3UIZ8 | Q8VCR8 | 0.143 | | | FALSE | TRUE | FALSE | FALSE |
| P01029 | P28666 | 0.145 | | | FALSE | TRUE | FALSE | FALSE |
| Q9QZA0 | Q9WVT6 | 0.145 | | | FALSE | TRUE | FALSE | FALSE |
| Q91XM9 | Q9WV34 | 0.148 | HOM | HOM | TRUE | FALSE | FALSE | FALSE |
| P15209 | Q6VNS1 | 0.149 | HOM | | TRUE | FALSE | FALSE | FALSE |
| P16015 | Q64444 | 0.15 | | HOM | TRUE | FALSE | FALSE | FALSE |
| Q8VCR2 | Q9EQ06 | 0.152 | | | FALSE | TRUE | FALSE | FALSE |
| Q9D2V7 | Q9WUM 4 | 0.153 | | HOM | TRUE | FALSE | FALSE | FALSE |
| Q61361 | Q9QUP5 | 0.153 | | | FALSE | TRUE | FALSE | FALSE |
| P08122 | Q9QZS0 | 0.154 | | | FALSE | TRUE | FALSE | FALSE |
| P11352 | P46412 | 0.154 | HOM | HOM | TRUE | FALSE | FALSE | FALSE |
| Q99KV1 | Q9QYJ0 | 0.155 | HOM | HOM | TRUE | FALSE | FALSE | FALSE |
| Q6PIE5 | Q8R429 | 0.155 | | | FALSE | TRUE | FALSE | FALSE |
| Q8JZV9 | Q91VT4 | 0.156 | HOM | HOM | TRUE | FALSE | FALSE | FALSE |

Table S4 Mouse

| | | | | | | | | |
|---|---|---|---|---|---|---|---|---|
| O70161 | Q91XU3 | 0.157 | | HOM | TRUE | FALSE | FALSE | FALSE |
| P08103 | Q04736 | 0.157 | HOM | | TRUE | FALSE | FALSE | FALSE |
| P84075 | Q8BNY6 | 0.158 | HET | HET | FALSE | FALSE | FALSE | FALSE |
| P51859 | Q99JF8 | 0.159 | HOM | | TRUE | FALSE | FALSE | FALSE |
| P55258 | Q99P58 | 0.159 | | HET | FALSE | FALSE | FALSE | FALSE |
| P63213 | Q9DAS9 | 0.16 | HET | | FALSE | FALSE | FALSE | FALSE |
| Q8C3Q5 | Q9CZN4 | 0.161 | | | FALSE | TRUE | FALSE | FALSE |
| O09174 | Q7TNE1 | 0.161 | | | FALSE | TRUE | FALSE | FALSE |
| Q811D0 | Q9JLB0 | 0.161 | HOM | | TRUE | FALSE | FALSE | FALSE |
| P31648 | P31649 | 0.162 | | | FALSE | TRUE | FALSE | FALSE |
| P63080 | P63137 | 0.162 | | | FALSE | TRUE | FALSE | FALSE |
| Q60902 | Q9QXY6 | 0.162 | HET | HET | FALSE | FALSE | FALSE | FALSE |
| P70290 | Q9JLB0 | 0.164 | HOM | | TRUE | FALSE | FALSE | FALSE |
| Q91W90 | Q921X9 | 0.164 | | | FALSE | TRUE | FALSE | FALSE |
| P21619 | P48678 | 0.164 | | HOM | TRUE | FALSE | FALSE | TRUE |
| O08989 | Q61411 | 0.165 | | | FALSE | TRUE | FALSE | FALSE |
| Q62523 | Q8BFW7 | 0.165 | | | FALSE | TRUE | FALSE | FALSE |
| Q9CWS4 | Q9QXK7 | 0.169 | | | FALSE | TRUE | FALSE | FALSE |
| O89017 | Q9CXY9 | 0.17 | | | FALSE | TRUE | FALSE | FALSE |
| Q922H2 | Q9JK42 | 0.17 | | HOM | TRUE | FALSE | FALSE | FALSE |
| P40240 | Q922J6 | 0.171 | HOM | | TRUE | FALSE | FALSE | FALSE |
| O09117 | Q8BGN8 | 0.171 | | | FALSE | TRUE | FALSE | FALSE |
| O70433 | Q9JKS4 | 0.171 | | | FALSE | TRUE | FALSE | FALSE |
| Q00PI9 | Q91VR5 | 0.171 | | | FALSE | TRUE | FALSE | FALSE |
| P63001 | P84096 | 0.172 | | | FALSE | TRUE | FALSE | FALSE |
| Q5FWK3 | Q5SSM3 | 0.173 | HET | | FALSE | FALSE | FALSE | FALSE |

Table S4 Mouse

| | | | | | | | | |
|---|---|---|---|---|---|---|---|---|
| Q5FW57 | Q9DCY0 | 0.173 | | | FALSE | TRUE | FALSE | FALSE |
| Q9CQW2 | Q9WUL7 | 0.174 | | | FALSE | TRUE | FALSE | FALSE |
| P26231 | Q61301 | 0.174 | HOM | | TRUE | FALSE | FALSE | FALSE |
| P10605 | P97821 | 0.174 | HET | HET | FALSE | FALSE | FALSE | FALSE |
| Q6PA06 | Q91YH5 | 0.175 | | | FALSE | TRUE | FALSE | FALSE |
| O08989 | P63321 | 0.176 | | | FALSE | TRUE | FALSE | FALSE |
| Q9DCT1 | Q9JII6 | 0.176 | | | FALSE | TRUE | FALSE | FALSE |
| P14733 | P46660 | 0.177 | | | FALSE | TRUE | FALSE | FALSE |
| P61022 | Q63810 | 0.177 | HET | HET | FALSE | FALSE | FALSE | FALSE |
| O09117 | O89104 | 0.177 | | | FALSE | TRUE | FALSE | FALSE |
| Q8BTG7 | Q9QYF9 | 0.177 | | | FALSE | TRUE | FALSE | FALSE |
| P49935 | Q9R013 | 0.179 | | HOM | TRUE | FALSE | FALSE | FALSE |
| P67984 | Q9D7S7 | 0.18 | | | FALSE | TRUE | FALSE | FALSE |
| Q5SX39 | Q9JMH9 | 0.18 | | | FALSE | TRUE | FALSE | FALSE |
| P11404 | Q05816 | 0.18 | | | FALSE | TRUE | FALSE | FALSE |
| P53986 | P57787 | 0.182 | | | FALSE | TRUE | FALSE | FALSE |
| Q6GV12 | Q9CXR1 | 0.183 | | | FALSE | TRUE | FALSE | FALSE |
| Q6P3A8 | Q9D051 | 0.184 | HOM | HOM | TRUE | FALSE | FALSE | FALSE |
| Q8JZV9 | Q99LB2 | 0.184 | HOM | | TRUE | FALSE | FALSE | FALSE |
| P63254 | P97315 | 0.184 | | | FALSE | TRUE | FALSE | FALSE |
| P50544 | Q60759 | 0.184 | HOM | HOM | TRUE | FALSE | FALSE | FALSE |
| O09159 | P27046 | 0.185 | | | FALSE | TRUE | FALSE | FALSE |
| P16015 | P28651 | 0.185 | | | FALSE | TRUE | FALSE | FALSE |
| Q01768 | Q9WV85 | 0.185 | | HOM | TRUE | FALSE | FALSE | FALSE |
| Q9CXR1 | Q9EQ06 | 0.187 | | | FALSE | TRUE | FALSE | FALSE |
| P62748 | P62761 | 0.188 | HET | | FALSE | FALSE | FALSE | FALSE |

Table S4 Mouse

| | | | | | | | | |
|---|---|---|---|---|---|---|---|---|
| Q99MZ7 | Q9CQ62 | 0.189 | | HOM | TRUE | FALSE | FALSE | FALSE |
| P60904 | Q9QYJ3 | 0.189 | | HOM | TRUE | FALSE | FALSE | FALSE |
| Q8BFZ3 | Q8R5C5 | 0.19 | | | FALSE | TRUE | FALSE | FALSE |
| E9PZQ0 | E9Q401 | 0.191 | HET | | FALSE | FALSE | FALSE | FALSE |
| P04104 | Q8VED5 | 0.191 | | | FALSE | TRUE | FALSE | FALSE |
| P04117 | Q00915 | 0.192 | HOM | | TRUE | FALSE | FALSE | FALSE |
| P98192 | Q61586 | 0.193 | HET | | FALSE | FALSE | FALSE | FALSE |
| P26883 | P30416 | 0.194 | HOM | HOM | TRUE | FALSE | FALSE | FALSE |
| B2RSH2 | O70443 | 0.195 | HOM | | TRUE | FALSE | FALSE | FALSE |
| Q91WS0 | Q9CQB5 | 0.197 | HOM | HOM | TRUE | FALSE | FALSE | FALSE |
| P62835 | Q80ZJ1 | 0.197 | | HOM | TRUE | FALSE | FALSE | FALSE |
| P10833 | P62071 | 0.198 | | | FALSE | TRUE | FALSE | FALSE |
| Q91X52 | Q99LB2 | 0.198 | HOM | | TRUE | FALSE | FALSE | FALSE |
| Q8VHE6 | Q9JHU4 | 0.2 | | HOM | TRUE | FALSE | FALSE | FALSE |
| Q5SX40 | Q6URW6 | 0.201 | | | FALSE | TRUE | FALSE | FALSE |
| Q3V1D3 | Q9DBT5 | 0.202 | | | FALSE | TRUE | FALSE | FALSE |
| Q64459 | Q64464 | 0.202 | | | FALSE | TRUE | FALSE | FALSE |
| Q9CZR3 | Q9QYA2 | 0.203 | | | FALSE | TRUE | FALSE | FALSE |
| P07309 | Q9CRB3 | 0.203 | | | FALSE | TRUE | FALSE | FALSE |
| P70245 | Q9D0P0 | 0.203 | | | FALSE | TRUE | FALSE | FALSE |
| P68040 | Q3UKJ7 | 0.203 | HOM | | TRUE | FALSE | FALSE | FALSE |
| O35459 | Q3TLP5 | 0.204 | HOM | HOM | TRUE | FALSE | FALSE | FALSE |
| Q61792 | Q9DC07 | 0.204 | | | FALSE | TRUE | FALSE | FALSE |
| Q62419 | Q8R3V5 | 0.204 | | | FALSE | TRUE | FALSE | FALSE |
| P16546 | Q7TPR4 | 0.205 | HOM | | TRUE | FALSE | FALSE | FALSE |
| Q6PIE5 | Q8VDN2 | 0.205 | | | FALSE | TRUE | FALSE | FALSE |

Table S4 Mouse

| | | | | | | | | |
|---|---|---|---|---|---|---|---|---|
| Q6TEK5 | Q9CRC0 | 0.205 | | | FALSE | TRUE | FALSE | FALSE |
| Q3TLP5 | Q8BH95 | 0.207 | HOM | HOM | TRUE | FALSE | FALSE | FALSE |
| P50153 | Q61016 | 0.207 | | | FALSE | TRUE | FALSE | FALSE |
| P37804 | Q9R1Q8 | 0.208 | | | FALSE | TRUE | FALSE | FALSE |
| P70372 | Q61701 | 0.208 | HOM | | TRUE | FALSE | FALSE | FALSE |
| Q8CAY6 | Q99JY0 | 0.209 | | HOM | TRUE | FALSE | FALSE | FALSE |
| P31648 | P31650 | 0.21 | | | FALSE | TRUE | FALSE | FALSE |
| P62259 | P68254 | 0.21 | HOM | | TRUE | FALSE | FALSE | FALSE |
| P63044 | Q62442 | 0.21 | | | FALSE | TRUE | FALSE | FALSE |
| Q8C0K5 | Q8R0Y8 | 0.21 | | | FALSE | TRUE | FALSE | FALSE |
| Q8BZF8 | Q9D0F9 | 0.21 | | | FALSE | TRUE | FALSE | FALSE |
| P61027 | Q99P58 | 0.21 | | HET | FALSE | FALSE | FALSE | FALSE |
| P28659 | Q9Z0H4 | 0.211 | HOM | | TRUE | FALSE | FALSE | FALSE |
| Q52KR3 | Q8BHE3 | 0.211 | | | FALSE | TRUE | FALSE | FALSE |
| P08074 | Q91VT4 | 0.211 | HOM | HOM | TRUE | FALSE | FALSE | FALSE |
| P01029 | Q61838 | 0.212 | | | FALSE | TRUE | FALSE | FALSE |
| P18052 | Q64487 | 0.212 | | | FALSE | TRUE | FALSE | FALSE |
| Q61102 | Q9DC29 | 0.212 | HET | | FALSE | FALSE | FALSE | FALSE |
| O88990 | P16546 | 0.213 | HOM | HOM | TRUE | FALSE | FALSE | FALSE |
| O70622 | Q99P72 | 0.213 | | | FALSE | TRUE | FALSE | FALSE |
| P68254 | P68510 | 0.213 | | | FALSE | TRUE | FALSE | FALSE |
| P54116 | Q99JB2 | 0.215 | HOM | HOM | TRUE | FALSE | FALSE | FALSE |
| P62746 | Q9QUI0 | 0.216 | | | FALSE | TRUE | FALSE | FALSE |
| P49813 | Q9JLH8 | 0.216 | | | FALSE | TRUE | FALSE | FALSE |
| P63011 | Q99P58 | 0.216 | | HET | FALSE | FALSE | FALSE | FALSE |
| O35490 | Q91WS4 | 0.217 | HOM | | TRUE | FALSE | FALSE | FALSE |

Table S4 Mouse

| | | | | | | | | |
|---|---|---|---|---|---|---|---|---|
| P11679 | Q3UV17 | 0.219 | | | FALSE | TRUE | FALSE | FALSE |
| P14873 | Q8C052 | 0.22 | | | FALSE | TRUE | FALSE | FALSE |
| P14094 | P97370 | 0.221 | | | FALSE | TRUE | FALSE | FALSE |
| O35136 | Q8BLK3 | 0.222 | HOM | | TRUE | FALSE | FALSE | FALSE |
| P00920 | Q9WVT6 | 0.222 | HOM | | TRUE | FALSE | FALSE | FALSE |
| Q8K406 | Q9JIA1 | 0.223 | | | FALSE | TRUE | FALSE | FALSE |
| O54916 | Q9EQP2 | 0.224 | HET | HET | FALSE | FALSE | FALSE | FALSE |
| Q9Z2I8 | Q9Z2I9 | 0.224 | | | FALSE | TRUE | FALSE | FALSE |
| Q91XM9 | Q9Z0U1 | 0.224 | HOM | HOM | TRUE | FALSE | FALSE | FALSE |
| P60904 | Q9QYI5 | 0.225 | | HOM | TRUE | FALSE | FALSE | FALSE |
| Q61330 | Q810U4 | 0.225 | | | FALSE | TRUE | FALSE | FALSE |
| Q8JZV9 | Q8VCC1 | 0.226 | HOM | HOM | TRUE | FALSE | FALSE | FALSE |
| B9EKR1 | O35239 | 0.227 | | | FALSE | TRUE | FALSE | FALSE |
| Q80XN0 | Q9R092 | 0.227 | | | FALSE | TRUE | FALSE | FALSE |
| Q8CHS7 | Q9CXR1 | 0.228 | | | FALSE | TRUE | FALSE | FALSE |
| O35621 | Q9Z2M7 | 0.229 | | | FALSE | TRUE | FALSE | FALSE |
| P97457 | Q9QVP4 | 0.23 | | | FALSE | TRUE | FALSE | FALSE |
| O88833 | Q9EP75 | 0.23 | | | FALSE | TRUE | FALSE | FALSE |
| P49312 | Q8BG05 | 0.23 | | | FALSE | TRUE | FALSE | FALSE |
| P15116 | Q9WTR5 | 0.23 | HET,HOM | HOM | TRUE | FALSE | FALSE | FALSE |
| Q62418 | Q9QXS6 | 0.23 | | | FALSE | TRUE | FALSE | FALSE |
| P62071 | P63321 | 0.231 | | | FALSE | TRUE | FALSE | FALSE |
| B2RSH2 | P27601 | 0.231 | HOM | | TRUE | FALSE | FALSE | FALSE |
| P04247 | Q9CX80 | 0.231 | | HOM | TRUE | FALSE | FALSE | FALSE |
| O55042 | Q9Z0F7 | 0.232 | | | FALSE | TRUE | FALSE | FALSE |
| Q60759 | Q8JZN5 | 0.232 | HOM | | TRUE | FALSE | FALSE | FALSE |

Table S4 Mouse

| | | | | | | | | | | | |
|---|---|---|---|---|---|---|---|---|---|---|---|
| Q11011 | Q8C129 | 0.232 | | | FALSE | TRUE | FALSE | | | | FALSE |
| O89020 | P07724 | 0.233 | | HOM | TRUE | FALSE | FALSE | | | | FALSE |
| P62259 | P63101 | 0.234 | HOM | | TRUE | FALSE | FALSE | | | | TRUE |
| Q80TZ3 | Q8CGB6 | 0.234 | | | FALSE | TRUE | FALSE | | | | FALSE |
| O54916 | Q9WVK4 | 0.235 | HET | HET | FALSE | FALSE | FALSE | | | | FALSE |
| Q8CHQ9 | Q9JIY7 | 0.235 | | | FALSE | TRUE | FALSE | | | | FALSE |
| O88908 | Q9Z2A7 | 0.236 | HET | HOM | TRUE | FALSE | FALSE | | | | FALSE |
| P97772 | Q14BI2 | 0.236 | HOM | | TRUE | FALSE | FALSE | | | | FALSE |
| O88951 | Q8JZS0 | 0.236 | HOM | | TRUE | FALSE | FALSE | | | | FALSE |
| Q8CGK3 | Q9DBN5 | 0.237 | HOM | | TRUE | FALSE | FALSE | | | | FALSE |
| Q64669 | Q9JI75 | 0.237 | | | FALSE | TRUE | FALSE | | | | FALSE |
| P35802 | P60202 | 0.237 | | | FALSE | TRUE | FALSE | | | | FALSE |
| P15626 | Q80W21 | 0.237 | | HOM | TRUE | FALSE | FALSE | | | | FALSE |
| P48036 | P97429 | 0.238 | | | FALSE | TRUE | FALSE | | | | FALSE |
| Q8R081 | Q91Z31 | 0.238 | | | FALSE | TRUE | FALSE | | | | FALSE |
| Q3UEB3 | Q61701 | 0.239 | HOM | | TRUE | FALSE | FALSE | | | | FALSE |
| P42208 | Q80UG5 | 0.239 | HOM | HET | TRUE | FALSE | TRUE | 1345 | Human | Septin complex | FALSE |
| Q8BGA9 | Q8VC74 | 0.239 | | | FALSE | TRUE | FALSE | | | | FALSE |
| Q8VCX1 | Q9DCT1 | 0.24 | | | FALSE | TRUE | FALSE | | | | FALSE |
| O88545 | Q9DCH4 | 0.24 | | | FALSE | TRUE | FALSE | | | | FALSE |
| P39447 | Q811D0 | 0.241 | | HOM | TRUE | FALSE | FALSE | | | | FALSE |
| P08113 | Q9CQN1 | 0.241 | HOM | HOM | TRUE | FALSE | FALSE | | | | FALSE |
| Q3UHU5 | Q6NZL0 | 0.243 | | | FALSE | TRUE | FALSE | | | | FALSE |
| Q60749 | Q9QYS9 | 0.243 | HOM | HOM | TRUE | FALSE | FALSE | | | | FALSE |
| Q91VW3 | Q9JJU8 | 0.243 | HOM | | TRUE | FALSE | FALSE | | | | FALSE |
| Q8VCR2 | Q9CXR1 | 0.243 | | | FALSE | TRUE | FALSE | | | | FALSE |

Table S4 Mouse

| | | | | | | | | |
|---|---|---|---|---|---|---|---|---|
| P43883 | Q9DBG5 | 0.244 | | HOM | TRUE | FALSE | FALSE | FALSE |
| P14211 | P35564 | 0.245 | | | FALSE | TRUE | FALSE | FALSE |
| Q3TLP5 | Q9D7J9 | 0.245 | HOM | HOM | TRUE | FALSE | FALSE | FALSE |
| Q62433 | Q9QYF9 | 0.247 | | | FALSE | TRUE | FALSE | FALSE |
| P29341 | Q61701 | 0.248 | | | FALSE | TRUE | FALSE | FALSE |
| Q64518 | Q8R429 | 0.249 | | | FALSE | TRUE | FALSE | FALSE |
| O70166 | P55821 | 0.249 | | | FALSE | TRUE | FALSE | FALSE |
| Q7TN29 | Q91VZ6 | 0.25 | | | FALSE | TRUE | FALSE | FALSE |
| Q91V92 | Q9WUM5 | 0.251 | | | FALSE | TRUE | FALSE | FALSE |
| Q60902 | Q8BH64 | 0.251 | HET | HOM, HET | TRUE, FALSE | FALSE | FALSE | FALSE |
| Q8CI94 | Q9WUB3 | 0.251 | HOM | HOM | TRUE | FALSE | FALSE | FALSE |
| P52792 | Q91W97 | 0.251 | | | FALSE | TRUE | FALSE | FALSE |
| P14824 | P97429 | 0.252 | | | FALSE | TRUE | FALSE | FALSE |
| P02469 | P97927 | 0.253 | | | FALSE | TRUE | FALSE | FALSE |
| Q01063 | Q01065 | 0.253 | | | FALSE | TRUE | FALSE | FALSE |
| Q80Z24 | Q8BLK3 | 0.253 | | | FALSE | TRUE | FALSE | FALSE |
| Q922J3 | Q9Z0H8 | 0.254 | | | FALSE | TRUE | FALSE | FALSE |
| P47941 | Q64010 | 0.255 | HOM | HET | TRUE | FALSE | FALSE | FALSE |
| P17156 | P63017 | 0.255 | | HOM | TRUE | FALSE | FALSE | FALSE |
| P26883 | Q62446 | 0.256 | HOM | | TRUE | FALSE | FALSE | FALSE |
| Q8BK67 | Q8VE37 | 0.256 | | | FALSE | TRUE | FALSE | FALSE |
| Q9ESW4 | Q9JIA7 | 0.257 | | | FALSE | TRUE | FALSE | FALSE |
| P63213 | Q61016 | 0.257 | HET | | FALSE | FALSE | FALSE | FALSE |
| P13634 | Q9QZA0 | 0.257 | | | FALSE | TRUE | FALSE | FALSE |
| P56375 | P56376 | 0.257 | | | FALSE | TRUE | FALSE | FALSE |

Table S4 Mouse

| | | | | | | | | |
|---|---|---|---|---|---|---|---|---|
| Q61016 | Q80SZ7 | 0.257 | | | FALSE | TRUE | FALSE | FALSE |
| P60766 | P84096 | 0.257 | HET | | FALSE | FALSE | FALSE | FALSE |
| Q9ET01 | Q9WUB3 | 0.258 | HOM | HOM | TRUE | FALSE | FALSE | FALSE |
| P40336 | Q8C0E2 | 0.259 | HET | | FALSE | FALSE | FALSE | TRUE |
| Q8BHC1 | Q91V41 | 0.26 | | | FALSE | TRUE | FALSE | FALSE |
| P14733 | P21619 | 0.26 | | | FALSE | TRUE | FALSE | FALSE |
| O35465 | P30416 | 0.261 | HET | HOM | TRUE | FALSE | FALSE | FALSE |
| Q91Z61 | Q99JI6 | 0.261 | | | FALSE | TRUE | FALSE | FALSE |
| Q3UHD9 | Q9QWY8 | 0.262 | | HOM | TRUE | FALSE | FALSE | TRUE |
| P62071 | Q9JIW9 | 0.262 | | HET | FALSE | FALSE | FALSE | FALSE |
| Q3TTY5 | Q3UV17 | 0.263 | | | FALSE | TRUE | FALSE | FALSE |
| P97427 | Q62188 | 0.263 | HET | HET | FALSE | FALSE | FALSE | TRUE |
| Q3U1J4 | Q921M3 | 0.263 | HOM | | TRUE | FALSE | FALSE | FALSE |
| O35660 | Q80W21 | 0.263 | | HOM | TRUE | FALSE | FALSE | FALSE |
| E9Q401 | P11881 | 0.264 | | | FALSE | TRUE | FALSE | FALSE |
| P61226 | P62835 | 0.264 | | | FALSE | TRUE | FALSE | FALSE |
| A2AMM0 | O54724 | 0.264 | | | FALSE | TRUE | FALSE | FALSE |
| Q9DBW0 | Q9EP75 | 0.265 | | | FALSE | TRUE | FALSE | FALSE |
| O08989 | P10833 | 0.265 | | | FALSE | TRUE | FALSE | FALSE |
| Q63886 | Q64435 | 0.265 | | | FALSE | TRUE | FALSE | FALSE |
| Q8R0Y6 | Q9CZS1 | 0.265 | | | FALSE | TRUE | FALSE | FALSE |
| O35609 | Q8K021 | 0.266 | | | FALSE | TRUE | FALSE | FALSE |
| Q920L1 | Q9Z0R9 | 0.268 | | | FALSE | TRUE | FALSE | FALSE |
| Q9D164 | Q9Z239 | 0.268 | | | FALSE | TRUE | FALSE | FALSE |
| Q3UZZ6 | Q9QWG7 | 0.269 | | HOM | TRUE | FALSE | FALSE | FALSE |
| Q8BLQ9 | Q8R5M8 | 0.27 | HOM | | TRUE | FALSE | FALSE | FALSE |

Table S4 Mouse

| | | | | | | | | | | | |
|---|---|---|---|---|---|---|---|---|---|---|---|
| P03995 | P19246 | 0.271 | | | FALSE | TRUE | FALSE | | | | FALSE |
| P01027 | Q61838 | 0.271 | HOM | | TRUE | FALSE | FALSE | | | | FALSE |
| P05977 | P09542 | 0.271 | | | FALSE | TRUE | FALSE | | | | FALSE |
| P62835 | Q91Z61 | 0.271 | | | FALSE | TRUE | FALSE | | | | FALSE |
| Q9EQC1 | Q9R1J0 | 0.271 | | | FALSE | TRUE | FALSE | | | | FALSE |
| Q8C5Q4 | Q9Z2X1 | 0.272 | | | FALSE | TRUE | FALSE | | | | FALSE |
| Q91ZZ3 | Q9Z0F7 | 0.272 | | | FALSE | TRUE | FALSE | | | | FALSE |
| Q7TSQ8 | Q8CFA2 | 0.272 | | HOM | TRUE | FALSE | FALSE | | | | FALSE |
| Q8VC52 | Q9WVB0 | 0.275 | | HOM | TRUE | FALSE | FALSE | | | | FALSE |
| Q569Z6 | Q8K019 | 0.276 | | | FALSE | TRUE | FALSE | | | | FALSE |
| P61028 | Q99P58 | 0.276 | | HET | FALSE | FALSE | FALSE | | | | FALSE |
| Q8R326 | Q8VIJ6 | 0.278 | HET | HOM | TRUE | FALSE | FALSE | | | | FALSE |
| Q641P0 | Q99JY9 | 0.278 | | | FALSE | TRUE | FALSE | | | | FALSE |
| Q91WC3 | Q99PU5 | 0.279 | | | FALSE | TRUE | FALSE | | | | FALSE |
| Q9CWF2 | Q9ERD7 | 0.28 | | HET | FALSE | FALSE | FALSE | | | | FALSE |
| P68372 | P99024 | 0.281 | HET | HET | FALSE | FALSE | FALSE | | | | FALSE |
| Q8BXK9 | Q9QYB1 | 0.281 | | | FALSE | TRUE | FALSE | | | | FALSE |
| P51150 | Q9R0M6 | 0.282 | | | FALSE | TRUE | FALSE | | | | FALSE |
| Q9CXY6 | Q9Z1X4 | 0.282 | HET | HET | FALSE | FALSE | TRUE | 1332,3055,5183 | Human | Large Drosha complex, Nop56p-associated pre-rRNA complex, DNA-PK-Ku-eIF2-NF90-NF45 complex | FALSE |
| Q64464 | Q9DBG1 | 0.283 | | | FALSE | TRUE | FALSE | | | | FALSE |
| P55258 | Q9CZT8 | 0.284 | | | FALSE | TRUE | FALSE | | | | FALSE |

Table S4 Mouse

| | | | | | | | | |
|---|---|---|---|---|---|---|---|---|
| P32883 | P63321 | 0.285 | | | FALSE | TRUE | FALSE | FALSE |
| P01027 | P28666 | 0.286 | HOM | | TRUE | FALSE | FALSE | FALSE |
| Q6P9R2 | Q8BTW9 | 0.286 | HOM | HET | TRUE | FALSE | FALSE | FALSE |
| Q62093 | Q9R0U0 | 0.286 | HOM | | TRUE | FALSE | FALSE | FALSE |
| Q62277 | Q8BGN8 | 0.286 | HOM | | TRUE | FALSE | FALSE | FALSE |
| P04117 | Q05816 | 0.287 | HOM | | TRUE | FALSE | FALSE | FALSE |
| Q8BLR2 | Q8BT60 | 0.287 | | | FALSE | TRUE | FALSE | FALSE |
| P17426 | P17427 | 0.287 | HET | HET | FALSE | FALSE | FALSE | FALSE |
| O70433 | Q8CI51 | 0.288 | | | FALSE | TRUE | FALSE | FALSE |
| O89020 | P21614 | 0.289 | | | FALSE | TRUE | FALSE | FALSE |
| P68372 | Q9ERD7 | 0.289 | HET | HET | FALSE | FALSE | FALSE | FALSE |
| Q3UVL4 | Q71KT5 | 0.289 | | | FALSE | TRUE | FALSE | FALSE |
| Q8VCT4 | Q8VCU1 | 0.289 | | | FALSE | TRUE | FALSE | FALSE |
| Q8CGN5 | Q9DBG5 | 0.29 | | HOM | TRUE | FALSE | FALSE | FALSE |
| P26041 | P26043 | 0.29 | | | FALSE | TRUE | FALSE | FALSE |
| O70325 | P46412 | 0.29 | | HOM | TRUE | FALSE | FALSE | FALSE |
| Q8CAY6 | Q8QZT1 | 0.29 | | HOM | TRUE | FALSE | FALSE | FALSE |
| P62484 | Q8CBW3 | 0.291 | | HET | FALSE | FALSE | FALSE | FALSE |
| Q8K023 | Q8VCX1 | 0.291 | HOM | | TRUE | FALSE | FALSE | FALSE |
| Q62108 | Q9JLB0 | 0.292 | HOM | | TRUE | FALSE | FALSE | FALSE |
| P29387 | P62881 | 0.293 | | HET | FALSE | FALSE | FALSE | FALSE |
| Q64435 | Q8BWQ1 | 0.294 | | | FALSE | TRUE | FALSE | FALSE |
| P05977 | Q8CI43 | 0.294 | | | FALSE | TRUE | FALSE | FALSE |
| P85094 | Q9DCC7 | 0.295 | | | FALSE | TRUE | FALSE | FALSE |
| Q810U3 | Q810U4 | 0.295 | | | FALSE | TRUE | FALSE | FALSE |
| P20108 | Q61171 | 0.295 | HOM | HOM | TRUE | FALSE | FALSE | FALSE |

Table S4 Mouse

| | | | | | | | | |
|---|---|---|---|---|---|---|---|---|
| P11679 | Q3TTY5 | 0.295 | | | FALSE | TRUE | FALSE | FALSE |
| Q3TTY5 | Q8VED5 | 0.295 | | | FALSE | TRUE | FALSE | FALSE |
| P62307 | P62313 | 0.296 | | | FALSE | TRUE | FALSE | FALSE |
| P15532 | Q9WV85 | 0.296 | | HOM | TRUE | FALSE | FALSE | FALSE |
| P09103 | Q91W90 | 0.297 | HOM | | TRUE | FALSE | FALSE | FALSE |
| P50428 | Q571E4 | 0.297 | | | FALSE | TRUE | FALSE | FALSE |
| A2AMM0 | Q91VJ2 | 0.298 | | | FALSE | TRUE | FALSE | FALSE |
| P27773 | Q921X9 | 0.298 | HET | | FALSE | FALSE | FALSE | FALSE |
| P58404 | Q9ERG2 | 0.299 | | | FALSE | TRUE | FALSE | FALSE |
| P68372 | Q9D6F9 | 0.299 | HET | HET | FALSE | FALSE | FALSE | FALSE |
| Q7TMM9 | Q922F4 | 0.299 | HET | | FALSE | FALSE | FALSE | FALSE |
| Q5SX40 | Q8VDD5 | 0.3 | | | FALSE | TRUE | FALSE | FALSE |
| Q8BH43 | Q8R5H6 | 0.3 | | HET | FALSE | FALSE | FALSE | FALSE |
| P62748 | Q8BGZ1 | 0.301 | HET | | FALSE | FALSE | FALSE | FALSE |
| Q8VCC2 | Q8VCU1 | 0.301 | | | FALSE | TRUE | FALSE | FALSE |
| Q9D0F3 | Q9DBH5 | 0.302 | HOM | | TRUE | FALSE | FALSE | FALSE |
| O88876 | Q99J47 | 0.302 | | | FALSE | TRUE | FALSE | FALSE |
| P62141 | P63087 | 0.302 | | | FALSE | TRUE | FALSE | FALSE |
| O88833 | Q9DBW0 | 0.302 | | | FALSE | TRUE | FALSE | FALSE |
| Q8VI47 | Q9R1S7 | 0.302 | | | FALSE | TRUE | FALSE | FALSE |
| P10630 | Q91VC3 | 0.302 | | HET | FALSE | FALSE | FALSE | FALSE |
| Q8BGG9 | Q9QYR9 | 0.303 | | | FALSE | TRUE | FALSE | FALSE |
| P63037 | Q9QYJ0 | 0.303 | HOM | HOM | TRUE | FALSE | FALSE | FALSE |
| P63318 | P68404 | 0.303 | | | FALSE | TRUE | FALSE | FALSE |
| P21836 | Q69ZK9 | 0.304 | HOM | | TRUE | FALSE | FALSE | FALSE |

Table S4 Mouse

| | | | | | | | | | | | |
|---|---|---|---|---|---|---|---|---|---|---|---|
| P68510 | Q9CQV8 | 0.304 | | | FALSE | TRUE | TRUE | 5199 | Human | Kinase maturatio n complex 1 | FALSE |
| P48318 | P48320 | 0.305 | HOM | | TRUE | FALSE | FALSE | | | | FALSE |
| Q80UG5 | Q9Z2Q6 | 0.305 | HET | HET | FALSE | FALSE | FALSE | | | | FALSE |
| Q8K394 | Q9Z1B3 | 0.306 | | | FALSE | TRUE | FALSE | | | | FALSE |
| B0V2N1 | Q64487 | 0.306 | HOM | | TRUE | FALSE | FALSE | | | | FALSE |
| P41216 | Q8JZR0 | 0.307 | | | FALSE | TRUE | FALSE | | | | FALSE |
| P62761 | P84075 | 0.308 | | HET | FALSE | FALSE | FALSE | | | | FALSE |
| Q3UV17 | Q8VED5 | 0.308 | | | FALSE | TRUE | FALSE | | | | FALSE |
| O70166 | P54227 | 0.308 | | | FALSE | TRUE | FALSE | | | | FALSE |
| P36371 | Q9CXJ4 | 0.309 | | | FALSE | TRUE | FALSE | | | | FALSE |
| O70443 | P21278 | 0.31 | | | FALSE | TRUE | FALSE | | | | FALSE |
| G3X982 | Q00519 | 0.31 | | | FALSE | TRUE | FALSE | | | | FALSE |
| P46096 | Q9R0N5 | 0.31 | | | FALSE | TRUE | FALSE | | | | FALSE |
| Q922F4 | Q9CWF2 | 0.31 | | | FALSE | TRUE | FALSE | | | | FALSE |
| O09043 | P18242 | 0.311 | HET | HET | FALSE | FALSE | FALSE | | | | FALSE |
| P04104 | Q3UV17 | 0.311 | | | FALSE | TRUE | FALSE | | | | FALSE |
| Q3TLP5 | Q9WUR 2 | 0.311 | HOM | HOM | TRUE | FALSE | FALSE | | | | FALSE |
| Q9QYB8 | Q9QYC0 | 0.312 | | | FALSE | TRUE | FALSE | | | | FALSE |
| Q62318 | Q9ESN6 | 0.312 | | | FALSE | TRUE | FALSE | | | | FALSE |
| P70372 | Q3UEB3 | 0.313 | HOM | HOM | TRUE | FALSE | FALSE | | | | FALSE |
| O88343 | Q5DTL9 | 0.313 | | | FALSE | TRUE | FALSE | | | | FALSE |
| Q99LD8 | Q9CWS0 | 0.314 | | | FALSE | TRUE | FALSE | | | | FALSE |
| B2RSH2 | P21279 | 0.315 | HOM | | TRUE | FALSE | FALSE | | | | FALSE |
| Q62420 | Q9JK48 | 0.316 | HOM | HET | TRUE | FALSE | FALSE | | | | FALSE |

Table S4 Mouse

| | | | | | | | | |
|---|---|---|---|---|---|---|---|---|
| P10833 | Q9JIW9 | 0.317 | | HET | FALSE | FALSE | FALSE | FALSE |
| Q922Q1 | Q9CW42 | 0.318 | | | FALSE | TRUE | FALSE | FALSE |
| P03995 | P21619 | 0.32 | | | FALSE | TRUE | FALSE | FALSE |
| P68372 | Q922F4 | 0.32 | HET | | FALSE | FALSE | FALSE | FALSE |
| Q921I1 | Q9DBD0 | 0.32 | | | FALSE | TRUE | FALSE | FALSE |
| P08207 | P14069 | 0.321 | | | FALSE | TRUE | FALSE | FALSE |
| P40124 | Q9CYT6 | 0.322 | HOM | HOM | TRUE | FALSE | FALSE | FALSE |
| P18872 | Q9DC51 | 0.322 | | HET | FALSE | FALSE | FALSE | FALSE |
| Q3TWW8 | Q6PDM2 | 0.322 | HOM | HOM | TRUE | FALSE | FALSE | FALSE |
| P19246 | P46660 | 0.323 | | | FALSE | TRUE | FALSE | FALSE |
| P21836 | Q8VCC2 | 0.323 | HOM | | TRUE | FALSE | FALSE | FALSE |
| B2RXS4 | Q8CJH3 | 0.323 | HET | HOM | TRUE | FALSE | FALSE | FALSE |
| O55022 | Q80UU9 | 0.324 | | | FALSE | TRUE | FALSE | FALSE |
| P63216 | Q9DAS9 | 0.324 | | | FALSE | TRUE | FALSE | FALSE |
| P70677 | P97864 | 0.325 | HOM | HOM | TRUE | FALSE | FALSE | FALSE |
| A2AMM0 | Q63918 | 0.325 | | | FALSE | TRUE | FALSE | FALSE |
| P08074 | Q8JZV9 | 0.325 | HOM | HOM | TRUE | FALSE | FALSE | FALSE |
| P13634 | P23589 | 0.325 | | | FALSE | TRUE | FALSE | FALSE |
| P07356 | P97429 | 0.326 | HOM | | TRUE | FALSE | FALSE | FALSE |
| P61226 | Q91Z61 | 0.326 | | | FALSE | TRUE | FALSE | FALSE |
| Q9DB77 | Q9DC61 | 0.326 | HET | | FALSE | FALSE | FALSE | FALSE |
| P14142 | P14246 | 0.327 | | | FALSE | TRUE | FALSE | FALSE |
| P35278 | P35282 | 0.328 | | | FALSE | TRUE | FALSE | FALSE |
| P21278 | Q9DC51 | 0.328 | | HET | FALSE | FALSE | FALSE | FALSE |
| P53994 | Q91V41 | 0.328 | | | FALSE | TRUE | FALSE | FALSE |
| Q91Z31 | Q921F4 | 0.328 | | | FALSE | TRUE | FALSE | FALSE |

Table S4 Mouse

| | | | | | | | | | |
|---|---|---|---|---|---|---|---|---|---|
| Q8BYM5 | Q8VCC2 | 0.329 | HET | | FALSE | FALSE | FALSE | | FALSE |
| Q925N1 | Q99JR1 | 0.329 | | | FALSE | TRUE | FALSE | | FALSE |
| P16015 | Q9QZA0 | 0.33 | | | FALSE | TRUE | FALSE | | FALSE |
| B2RSH2 | P18872 | 0.331 | HOM | | TRUE | FALSE | FALSE | | FALSE |
| Q8VCX1 | Q9JII6 | 0.332 | | | FALSE | TRUE | FALSE | | FALSE |
| Q9CZY3 | Q9D2M8 | 0.332 | | | FALSE | TRUE | FALSE | | FALSE |
| Q9JJU8 | Q9WUZ7 | 0.333 | | | FALSE | TRUE | FALSE | | FALSE |
| Q63880 | Q8VCT4 | 0.333 | | | FALSE | TRUE | FALSE | | FALSE |
| O54749 | P24457 | 0.333 | | HOM | TRUE | FALSE | FALSE | | FALSE |
| Q6P5F7 | Q9D3A9 | 0.335 | | | FALSE | TRUE | FALSE | | FALSE |
| Q61425 | Q9DBM2 | 0.337 | HOM | | TRUE | FALSE | FALSE | | FALSE |
| P21958 | Q9JI39 | 0.338 | | HOM | TRUE | FALSE | FALSE | | FALSE |
| Q6PGN3 | Q9JLM8 | 0.338 | | | FALSE | TRUE | FALSE | | FALSE |
| P09103 | P27773 | 0.338 | HOM | HET | TRUE | FALSE | FALSE | | FALSE |
| P28666 | Q61838 | 0.338 | | | FALSE | TRUE | FALSE | | FALSE |
| Q3UHL1 | Q6PHZ2 | 0.338 | | HET | FALSE | FALSE | FALSE | | FALSE |
| P68372 | Q9CWF2 | 0.339 | HET | | FALSE | FALSE | FALSE | | FALSE |
| P84084 | Q9D0J4 | 0.339 | | HET | FALSE | FALSE | FALSE | | FALSE |
| O35367 | P50608 | 0.34 | | | FALSE | TRUE | FALSE | | FALSE |
| P53994 | Q91ZR1 | 0.34 | | | FALSE | TRUE | FALSE | | FALSE |
| Q8C1B7 | Q9R1T4 | 0.341 | | HOM | TRUE | FALSE | FALSE | | FALSE |
| Q8BMS1 | Q9DBM2 | 0.342 | | | FALSE | TRUE | FALSE | | FALSE |
| Q5SX39 | Q61879 | 0.342 | | | FALSE | TRUE | FALSE | | FALSE |
| P18872 | P21278 | 0.342 | | | FALSE | TRUE | FALSE | | FALSE |
| O70172 | Q91XU3 | 0.342 | | HOM | TRUE | FALSE | FALSE | | FALSE |
| P28665 | Q61838 | 0.344 | | | FALSE | TRUE | FALSE | | FALSE |

Table S4 Mouse

| | | | | | | | | | | | |
|---|---|---|---|---|---|---|---|---|---|---|---|
| P50153 | P63216 | 0.346 | | | FALSE | TRUE | FALSE | | | | FALSE |
| Q8R3V5 | Q9JK48 | 0.346 | | HET | FALSE | FALSE | FALSE | | | | FALSE |
| O35728 | Q9DBW0 | 0.346 | | | FALSE | TRUE | FALSE | | | | FALSE |
| P60764 | P60766 | 0.347 | | HET | FALSE | FALSE | FALSE | | | | FALSE |
| Q5SQX6 | Q7TMB8 | 0.347 | | HET | FALSE | FALSE | FALSE | | | | TRUE |
| Q8VEK3 | Q91VR5 | 0.348 | | | FALSE | TRUE | TRUE | 1332 | Human | Large Drosha complex | FALSE |
| P24549 | P47738 | 0.348 | | HOM | TRUE | FALSE | FALSE | | | | FALSE |
| O08810 | P58252 | 0.349 | | | FALSE | TRUE | FALSE | | | | FALSE |
| P84084 | Q9WUL7 | 0.35 | | | FALSE | TRUE | FALSE | | | | FALSE |
| Q3TMH2 | Q9CZC8 | 0.35 | | | FALSE | TRUE | FALSE | | | | FALSE |
| Q8VED9 | Q9JL15 | 0.35 | HOM | | TRUE | FALSE | FALSE | | | | FALSE |
| P28738 | Q61768 | 0.35 | HOM | HOM | TRUE | FALSE | FALSE | | | | TRUE |
| Q8VCC1 | Q91VT4 | 0.351 | HOM | HOM | TRUE | FALSE | FALSE | | | | FALSE |
| P08074 | P50171 | 0.351 | HOM | HOM | TRUE | FALSE | FALSE | | | | FALSE |
| A2AQ07 | Q9ERD7 | 0.352 | HET | HET | FALSE | FALSE | FALSE | | | | FALSE |
| Q8R238 | Q8VBT2 | 0.352 | HOM | | TRUE | FALSE | FALSE | | | | FALSE |
| P70694 | Q9DCT1 | 0.353 | HOM | | TRUE | FALSE | FALSE | | | | FALSE |
| Q61233 | Q99K51 | 0.354 | | | FALSE | TRUE | FALSE | | | | FALSE |
| P70694 | Q9JII6 | 0.355 | HOM | | TRUE | FALSE | FALSE | | | | FALSE |
| Q62419 | Q62420 | 0.355 | | HOM | TRUE | FALSE | FALSE | | | | FALSE |
| Q9CR58 | Q9QZD8 | 0.355 | | | FALSE | TRUE | FALSE | | | | FALSE |
| Q5SX40 | Q91Z83 | 0.355 | | | FALSE | TRUE | FALSE | | | | FALSE |
| P70175 | Q91XM9 | 0.355 | HOM | HOM | TRUE | FALSE | FALSE | | | | TRUE |
| P21981 | Q8BH61 | 0.357 | HOM | | TRUE | FALSE | FALSE | | | | FALSE |
| Q9JLB0 | Q9WV34 | 0.358 | | HOM | TRUE | FALSE | FALSE | | | | FALSE |

Table S4 Mouse

| | | | | | | | | | | | |
|---|---|---|---|---|---|---|---|---|---|---|---|
| P99024 | Q7TMM9 | 0.358 | HET | HET | FALSE | FALSE | FALSE | | | | FALSE |
| Q8BGZ1 | Q8BNY6 | 0.359 | | HET | FALSE | FALSE | FALSE | | | | FALSE |
| Q9CXP8 | Q9DAS9 | 0.359 | | | FALSE | TRUE | FALSE | | | | FALSE |
| P21300 | Q9JII6 | 0.36 | | | FALSE | TRUE | FALSE | | | | FALSE |
| Q7TMM9 | Q9ERD7 | 0.36 | HET | HET | FALSE | FALSE | FALSE | | | | FALSE |
| P16054 | P68404 | 0.36 | | | FALSE | TRUE | FALSE | | | | FALSE |
| P50153 | Q9DAS9 | 0.36 | | | FALSE | TRUE | FALSE | | | | FALSE |
| P61982 | P68510 | 0.362 | | | FALSE | TRUE | TRUE | 5199 | Human | Kinase maturation complex 1 | FALSE |
| P53994 | Q8BHC1 | 0.365 | | | FALSE | TRUE | FALSE | | | | FALSE |
| P28661 | Q80UG5 | 0.365 | HET | HET | FALSE | FALSE | FALSE | | | | FALSE |
| Q8BNY6 | Q91X97 | 0.365 | HET | HET | FALSE | FALSE | FALSE | | | | FALSE |
| P28661 | Q9Z2Q6 | 0.366 | HET | HET | FALSE | FALSE | FALSE | | | | FALSE |
| Q791T5 | Q791V5 | 0.367 | | | FALSE | TRUE | FALSE | | | | FALSE |
| P56392 | Q61387 | 0.367 | | | FALSE | TRUE | FALSE | | | | FALSE |
| Q8K023 | Q9DCT1 | 0.368 | HOM | | TRUE | FALSE | FALSE | | | | FALSE |
| Q4V9Z5 | Q923L3 | 0.368 | | | FALSE | TRUE | FALSE | | | | FALSE |
| P70290 | Q91XM9 | 0.368 | HOM | HOM | TRUE | FALSE | FALSE | | | | FALSE |
| P39039 | P41317 | 0.369 | | | FALSE | TRUE | FALSE | | | | FALSE |
| O89112 | Q9JJK2 | 0.37 | | | FALSE | TRUE | FALSE | | | | FALSE |
| A3KFM7 | Q6PDQ2 | 0.371 | HOM | HOM | TRUE | FALSE | FALSE | | | | FALSE |
| P16406 | Q9EQH2 | 0.371 | | HET | FALSE | FALSE | FALSE | | | | FALSE |
| Q9JJW5 | Q9JK37 | 0.373 | | | FALSE | TRUE | FALSE | | | | FALSE |
| P21279 | Q9DC51 | 0.373 | | HET | FALSE | FALSE | FALSE | | | | FALSE |
| P23589 | P28651 | 0.374 | | | FALSE | TRUE | FALSE | | | | FALSE |

Table S4 Mouse

| | | | | | | | | |
|---|---|---|---|---|---|---|---|---|
| P31648 | Q61327 | 0.375 | | | FALSE | TRUE | FALSE | FALSE |
| P46638 | Q91V41 | 0.376 | | | FALSE | TRUE | FALSE | FALSE |
| Q8VCC1 | Q91X52 | 0.376 | HOM | HOM | TRUE | FALSE | FALSE | FALSE |
| Q8BTM8 | Q8VHX6 | 0.377 | HOM | HOM | TRUE | FALSE | FALSE | FALSE |
| Q3V009 | Q9R0Q3 | 0.378 | | | FALSE | TRUE | FALSE | FALSE |
| P24472 | P30115 | 0.378 | HOM | HOM | TRUE | FALSE | FALSE | FALSE |
| O88741 | Q8VE33 | 0.378 | | | FALSE | TRUE | FALSE | FALSE |
| P62137 | P62141 | 0.38 | HOM | | TRUE | FALSE | FALSE | FALSE |
| P50462 | Q9DCT8 | 0.38 | | | FALSE | TRUE | FALSE | FALSE |
| P50285 | P97872 | 0.381 | | | FALSE | TRUE | FALSE | FALSE |
| P35235 | P54830 | 0.382 | | | FALSE | TRUE | FALSE | FALSE |
| Q91V61 | Q925N1 | 0.383 | | | FALSE | TRUE | FALSE | FALSE |
| O35526 | P61264 | 0.384 | | | FALSE | TRUE | FALSE | FALSE |
| Q69ZN7 | Q9ESD7 | 0.386 | | | FALSE | TRUE | FALSE | FALSE |
| Q60759 | Q9DBL1 | 0.386 | HOM | HOM | TRUE | FALSE | FALSE | FALSE |
| Q8C166 | Q9Z140 | 0.386 | HOM | | TRUE | FALSE | FALSE | FALSE |
| P21300 | Q8K023 | 0.386 | | HOM | TRUE | FALSE | FALSE | FALSE |
| P58771 | Q6IRU2 | 0.387 | | | FALSE | TRUE | FALSE | FALSE |
| A2BDX3 | Q8VE47 | 0.387 | | HOM | TRUE | FALSE | FALSE | FALSE |
| P84096 | Q05144 | 0.388 | | HET | FALSE | FALSE | FALSE | FALSE |
| P56654 | Q6XVG2 | 0.389 | | | FALSE | TRUE | FALSE | FALSE |
| O35954 | P53810 | 0.389 | | HOM | TRUE | FALSE | FALSE | FALSE |
| P10833 | Q61411 | 0.39 | | | FALSE | TRUE | FALSE | FALSE |
| P45952 | Q60759 | 0.391 | HOM | HOM | TRUE | FALSE | FALSE | FALSE |
| Q80Y17 | Q8K400 | 0.391 | | | FALSE | TRUE | FALSE | FALSE |
| P97384 | P97429 | 0.392 | | | FALSE | TRUE | FALSE | FALSE |

Table S4 Mouse

| | | | | | | | | | | | |
|---|---|---|---|---|---|---|---|---|---|---|---|
| Q9ESN6 | Q9R1R2 | 0.393 | | | FALSE | TRUE | FALSE | | | | FALSE |
| P50462 | P63254 | 0.394 | | | FALSE | TRUE | FALSE | | | | FALSE |
| P43006 | P56564 | 0.394 | | | FALSE | TRUE | FALSE | | | | FALSE |
| Q99J47 | Q9EQ06 | 0.394 | | | FALSE | TRUE | FALSE | | | | FALSE |
| Q6NSR8 | Q9CPY7 | 0.395 | | | FALSE | TRUE | FALSE | | | | FALSE |
| Q8BSL7 | Q9CQW2 | 0.395 | | | FALSE | TRUE | FALSE | | | | FALSE |
| Q60930 | Q60931 | 0.395 | | | FALSE | TRUE | FALSE | | | | FALSE |
| Q91V41 | Q91ZR1 | 0.395 | | | FALSE | TRUE | FALSE | | | | FALSE |
| P63101 | P68510 | 0.396 | | | FALSE | TRUE | FALSE | | | | TRUE |
| P40237 | Q8BJU2 | 0.396 | | | FALSE | TRUE | FALSE | | | | FALSE |
| Q61694 | Q9EQC1 | 0.396 | | | FALSE | TRUE | FALSE | | | | FALSE |
| P14733 | P19246 | 0.396 | | | FALSE | TRUE | FALSE | | | | FALSE |
| P97384 | Q07076 | 0.397 | | | FALSE | TRUE | FALSE | | | | FALSE |
| Q6X893 | Q8BY89 | 0.397 | | | FALSE | TRUE | FALSE | | | | FALSE |
| Q9WV34 | Q9Z0U1 | 0.397 | HOM | HOM | TRUE | FALSE | FALSE | | | | FALSE |
| P30416 | Q62446 | 0.398 | HOM | | TRUE | FALSE | FALSE | | | | FALSE |
| P63011 | Q9CZT8 | 0.399 | | | FALSE | TRUE | FALSE | | | | FALSE |
| P46638 | Q91ZR1 | 0.4 | | | FALSE | TRUE | FALSE | | | | FALSE |
| P61226 | Q99JI6 | 0.4 | | | FALSE | TRUE | FALSE | | | | FALSE |
| O88602 | Q9JJV5 | 0.4 | | | FALSE | TRUE | FALSE | | | | FALSE |
| P10649 | P19157 | 0.4 | | HOM | TRUE | FALSE | FALSE | | | | FALSE |
| Q8BLK3 | Q99PJ0 | 0.403 | | | FALSE | TRUE | FALSE | | | | FALSE |
| P84091 | Q9JKC8 | 0.403 | HET | HET | FALSE | FALSE | FALSE | | | | FALSE |
| Q9D4H8 | Q9JLV5 | 0.403 | HET | HOM, HET | TRUE, FALSE | FALSE | TRUE | 2715 | Human | Ubiquitin E3 ligase | FALSE |
| Q64458 | Q6XVG2 | 0.403 | | | FALSE | TRUE | FALSE | | | | FALSE |

Table S4 Mouse

| | | | | | | | | |
|---|---|---|---|---|---|---|---|---|
| P02469 | Q60675 | 0.404 | | HOM | TRUE | FALSE | FALSE | FALSE |
| Q922F4 | Q9D6F9 | 0.405 | | HET | FALSE | FALSE | FALSE | FALSE |
| Q91VR7 | Q9CQV6 | 0.406 | HOM | HOM | TRUE | FALSE | FALSE | FALSE |
| Q07417 | Q60759 | 0.406 | HOM | HOM | TRUE | FALSE | FALSE | FALSE |
| Q8CIB5 | Q8K1B8 | 0.406 | | | FALSE | TRUE | FALSE | FALSE |
| P21995 | P97300 | 0.406 | | | FALSE | TRUE | FALSE | FALSE |
| O88569 | P49312 | 0.407 | | | FALSE | TRUE | FALSE | FALSE |
| Q5FWK3 | Q91Z67 | 0.408 | HET | HET | FALSE | FALSE | FALSE | FALSE |
| P62874 | P62881 | 0.409 | HET | HET | FALSE | FALSE | FALSE | FALSE |
| B0V2N1 | P18052 | 0.41 | HOM | | TRUE | FALSE | FALSE | FALSE |
| P19639 | Q80W21 | 0.41 | | HOM | TRUE | FALSE | FALSE | FALSE |
| Q91XM9 | Q9JLB0 | 0.41 | HOM | | TRUE | FALSE | FALSE | FALSE |
| O89086 | Q91VM5 | 0.411 | | | FALSE | TRUE | FALSE | FALSE |
| Q61361 | Q9ESM3 | 0.412 | | | FALSE | TRUE | FALSE | FALSE |
| P12849 | Q9DBC7 | 0.413 | | | FALSE | TRUE | FALSE | FALSE |
| Q3ULF4 | Q920A7 | 0.413 | HET | | FALSE | FALSE | FALSE | FALSE |
| Q8QZR3 | Q8VCT4 | 0.414 | | | FALSE | TRUE | FALSE | FALSE |
| P63216 | Q9CXP8 | 0.415 | | | FALSE | TRUE | FALSE | FALSE |
| Q505F5 | Q9WUA2 | 0.415 | | HOM | TRUE | FALSE | FALSE | FALSE |
| P49312 | Q9CX86 | 0.415 | | | FALSE | TRUE | FALSE | FALSE |
| O54991 | Q9CS84 | 0.416 | | HET | FALSE | FALSE | FALSE | FALSE |
| Q8BWN8 | Q91X34 | 0.416 | | | FALSE | TRUE | FALSE | FALSE |
| P63213 | P63216 | 0.417 | HET | | FALSE | FALSE | FALSE | FALSE |
| Q8JZN5 | Q9DBL1 | 0.417 | | HOM | TRUE | FALSE | FALSE | FALSE |
| P23953 | Q8QZR3 | 0.418 | | | FALSE | TRUE | FALSE | FALSE |
| O88346 | P50752 | 0.419 | | | FALSE | TRUE | FALSE | FALSE |

Table S4 Mouse

| | | | | | | | | |
|---|---|---|---|---|---|---|---|---|
| Q9QZD9 | Q9Z1Z2 | 0.42 | HET | | FALSE | FALSE | FALSE | FALSE |
| Q91V64 | Q9DCC7 | 0.42 | | | FALSE | TRUE | FALSE | FALSE |
| Q8BHC1 | Q91ZR1 | 0.42 | | | FALSE | TRUE | FALSE | FALSE |
| P21300 | Q9DCT1 | 0.421 | | | FALSE | TRUE | FALSE | FALSE |
| P56371 | Q91V41 | 0.422 | HET | | FALSE | FALSE | FALSE | FALSE |
| P60766 | P63001 | 0.422 | HET | | FALSE | FALSE | FALSE | FALSE |
| Q68FL4 | Q80SW1 | 0.422 | | | FALSE | TRUE | FALSE | FALSE |
| Q8BG73 | Q91VW3 | 0.426 | | HOM | TRUE | FALSE | FALSE | FALSE |
| Q8CFA2 | Q9DBT9 | 0.428 | HOM | | TRUE | FALSE | FALSE | FALSE |
| P13542 | Q02566 | 0.428 | | | FALSE | TRUE | FALSE | FALSE |
| P47738 | Q8R0Y6 | 0.429 | HOM | | TRUE | FALSE | FALSE | FALSE |
| O88712 | Q61753 | 0.429 | HET | HOM | TRUE | FALSE | FALSE | FALSE |
| P97315 | Q9DCT8 | 0.43 | | | FALSE | TRUE | FALSE | FALSE |
| P05063 | P05064 | 0.43 | | | FALSE | TRUE | FALSE | FALSE |
| P13634 | P16015 | 0.431 | | | FALSE | TRUE | FALSE | FALSE |
| P63005 | Q6PE01 | 0.431 | HOM | | TRUE | FALSE | FALSE | FALSE |
| Q80XI4 | Q91XU3 | 0.431 | | HOM | TRUE | FALSE | FALSE | FALSE |
| Q6PFD5 | Q9D415 | 0.431 | | | FALSE | TRUE | FALSE | TRUE |
| P07934 | Q9JLM8 | 0.431 | HET | | FALSE | FALSE | FALSE | FALSE |
| Q3UVX5 | Q9QYS2 | 0.432 | | | FALSE | TRUE | FALSE | FALSE |
| Q80ZJ1 | Q91Z61 | 0.432 | HOM | | TRUE | FALSE | FALSE | FALSE |
| Q8VC28 | Q9DCT1 | 0.433 | HOM | | TRUE | FALSE | FALSE | FALSE |
| O35609 | Q9JKD3 | 0.433 | | | FALSE | TRUE | FALSE | FALSE |
| P07758 | Q00898 | 0.433 | HOM | HOM | TRUE | FALSE | FALSE | FALSE |
| P46638 | Q8BHC1 | 0.433 | | | FALSE | TRUE | FALSE | FALSE |
| P61021 | Q9CQD1 | 0.435 | | | FALSE | TRUE | FALSE | FALSE |

Table S4 Mouse

| | | | | | | | | |
|---|---|---|---|---|---|---|---|---|
| P19157 | P19639 | 0.436 | HOM | | TRUE | FALSE | FALSE | FALSE |
| Q99P72 | Q9ES97 | 0.436 | | | FALSE | TRUE | FALSE | FALSE |
| P97821 | Q99JR5 | 0.436 | HET | | FALSE | FALSE | FALSE | FALSE |
| O54829 | Q9Z2H2 | 0.436 | | | FALSE | TRUE | FALSE | FALSE |
| Q5PR73 | Q91Z61 | 0.436 | | | FALSE | TRUE | FALSE | FALSE |
| Q6PIC6 | Q8VDN2 | 0.437 | | | FALSE | TRUE | FALSE | FALSE |
| O35737 | P70333 | 0.437 | | | FALSE | TRUE | FALSE | FALSE |
| Q9D1E6 | Q9Z0H8 | 0.438 | HET | | FALSE | FALSE | FALSE | FALSE |
| O09158 | Q64464 | 0.438 | | | FALSE | TRUE | FALSE | FALSE |
| Q8BGG9 | Q91X34 | 0.438 | | | FALSE | TRUE | FALSE | FALSE |
| P30275 | Q04447 | 0.439 | | | FALSE | TRUE | FALSE | FALSE |
| P32883 | Q61411 | 0.439 | | | FALSE | TRUE | FALSE | FALSE |
| Q69ZK9 | Q8BYM5 | 0.44 | | HET | FALSE | FALSE | FALSE | FALSE |
| P63254 | Q9DCT8 | 0.44 | | | FALSE | TRUE | FALSE | FALSE |
| Q8R1V4 | Q99KF1 | 0.441 | | | FALSE | TRUE | FALSE | FALSE |
| O88990 | Q62261 | 0.441 | HOM | HOM | TRUE | FALSE | FALSE | FALSE |
| Q60759 | Q9JHI5 | 0.442 | HOM | | TRUE | FALSE | FALSE | FALSE |
| O35945 | P24549 | 0.442 | | | FALSE | TRUE | FALSE | FALSE |
| P61750 | P84084 | 0.442 | | | FALSE | TRUE | FALSE | FALSE |
| P52430 | Q62086 | 0.442 | | | FALSE | TRUE | FALSE | FALSE |
| Q80X90 | Q8BTM8 | 0.444 | | HOM | TRUE | FALSE | FALSE | FALSE |
| P10639 | Q8CDN6 | 0.444 | HOM, HET | HET | TRUE, FALSE | FALSE | FALSE | FALSE |
| P99024 | Q9CWF2 | 0.445 | HET | | FALSE | FALSE | FALSE | FALSE |
| P50396 | Q61598 | 0.445 | HET | HET | FALSE | FALSE | FALSE | FALSE |
| P07356 | P97384 | 0.445 | HOM | | TRUE | FALSE | FALSE | FALSE |

Table S4 Mouse

| | | | | | | | | |
|---|---|---|---|---|---|---|---|---|
| P28651 | Q9WVT6 | 0.447 | | | FALSE | TRUE | FALSE | FALSE |
| P46096 | Q9R0N7 | 0.448 | | HET | FALSE | FALSE | FALSE | FALSE |
| P62823 | Q99P58 | 0.448 | | HET | FALSE | FALSE | FALSE | FALSE |
| Q62159 | Q9QUI0 | 0.448 | | | FALSE | TRUE | FALSE | FALSE |
| P48193 | Q8BGS1 | 0.449 | | | FALSE | TRUE | FALSE | FALSE |
| P70175 | Q62108 | 0.449 | HOM | HOM | TRUE | FALSE | FALSE | TRUE |
| O35639 | P97384 | 0.45 | | | FALSE | TRUE | FALSE | FALSE |
| P10649 | Q80W21 | 0.45 | | HOM | TRUE | FALSE | FALSE | FALSE |
| P84075 | Q91X97 | 0.45 | HET | HET | FALSE | FALSE | FALSE | FALSE |
| P14824 | Q07076 | 0.45 | | | FALSE | TRUE | FALSE | FALSE |
| Q8CEE7 | Q9ERI6 | 0.45 | | | FALSE | TRUE | FALSE | FALSE |
| P27601 | Q9DC51 | 0.451 | | HET | FALSE | FALSE | FALSE | FALSE |
| O88492 | Q9DBG5 | 0.452 | | HOM | TRUE | FALSE | FALSE | FALSE |
| P50153 | Q9CXP8 | 0.452 | | | FALSE | TRUE | FALSE | FALSE |
| P10605 | Q99JR5 | 0.452 | HET | | FALSE | FALSE | FALSE | FALSE |
| P84078 | Q9WUL7 | 0.453 | | | FALSE | TRUE | FALSE | FALSE |
| P26516 | Q9DCH4 | 0.454 | | | FALSE | TRUE | FALSE | FALSE |
| P21958 | Q9CXJ4 | 0.454 | | | FALSE | TRUE | FALSE | FALSE |
| Q8JZQ2 | Q920A7 | 0.454 | HET | | FALSE | FALSE | FALSE | FALSE |
| Q61644 | Q9WVE8 | 0.454 | HET | | FALSE | FALSE | FALSE | FALSE |
| Q8BK48 | Q8VCU1 | 0.455 | | | FALSE | TRUE | FALSE | FALSE |
| P61226 | Q5PR73 | 0.455 | | | FALSE | TRUE | FALSE | FALSE |
| P68372 | Q7TMM9 | 0.455 | HET | HET | FALSE | FALSE | FALSE | FALSE |
| P23953 | Q8VCU1 | 0.455 | | | FALSE | TRUE | FALSE | FALSE |
| Q8BGG9 | Q8BWN8 | 0.456 | | | FALSE | TRUE | FALSE | FALSE |
| P07901 | P08113 | 0.456 | HOM | HOM | TRUE | FALSE | FALSE | FALSE |

Table S4 Mouse

| | | | | | | | | |
|---|---|---|---|---|---|---|---|---|
| P70175 | Q9WV34 | 0.456 | HOM | HOM | TRUE | FALSE | FALSE | FALSE |
| P03995 | P46660 | 0.456 | | | FALSE | TRUE | FALSE | FALSE |
| Q8BGY7 | Q9D8B6 | 0.456 | | | FALSE | TRUE | FALSE | FALSE |
| O09158 | Q9DBG1 | 0.457 | | | FALSE | TRUE | FALSE | FALSE |
| P61458 | Q9CZL5 | 0.458 | HOM | HOM | TRUE | FALSE | FALSE | TRUE |
| P08752 | P18872 | 0.458 | | | FALSE | TRUE | FALSE | FALSE |
| Q9CR51 | Q9WTT4 | 0.459 | | | FALSE | TRUE | FALSE | FALSE |
| Q3THG9 | Q9R0Q7 | 0.462 | | | FALSE | TRUE | FALSE | FALSE |
| Q61016 | Q9CXP8 | 0.462 | | | FALSE | TRUE | FALSE | FALSE |
| Q8R574 | Q9D7G0 | 0.463 | | HOM | TRUE | FALSE | FALSE | FALSE |
| Q91Z38 | Q9CZW5 | 0.463 | HOM | HOM | TRUE | FALSE | FALSE | FALSE |
| Q8BWT1 | Q8CAY6 | 0.463 | | | FALSE | TRUE | FALSE | FALSE |
| P68134 | Q8BFZ3 | 0.464 | | | FALSE | TRUE | FALSE | FALSE |
| Q61330 | Q810U3 | 0.465 | | | FALSE | TRUE | FALSE | FALSE |
| Q9D6F9 | Q9ERD7 | 0.465 | HET | HET | FALSE | FALSE | FALSE | FALSE |
| P62823 | Q9CZT8 | 0.466 | | | FALSE | TRUE | FALSE | FALSE |
| P23953 | Q8BK48 | 0.466 | | | FALSE | TRUE | FALSE | FALSE |
| Q7TQ48 | Q9Z0R4 | 0.467 | HET | HET | FALSE | FALSE | FALSE | FALSE |
| P50608 | Q9JK53 | 0.468 | | | FALSE | TRUE | FALSE | FALSE |
| Q6PHN9 | Q9D1G1 | 0.469 | | | FALSE | TRUE | FALSE | FALSE |
| P57722 | Q61990 | 0.47 | | | FALSE | TRUE | FALSE | FALSE |
| Q71RI9 | Q8BTY1 | 0.47 | HOM | | TRUE | FALSE | FALSE | FALSE |
| P62331 | Q9WUL7 | 0.471 | HOM | | TRUE | FALSE | FALSE | FALSE |
| A2AQ07 | Q922F4 | 0.471 | HET | | FALSE | FALSE | FALSE | FALSE |
| O54749 | P11714 | 0.472 | | HOM | TRUE | FALSE | FALSE | FALSE |
| Q8BGA8 | Q9QXG4 | 0.474 | | | FALSE | TRUE | FALSE | FALSE |

Table S4 Mouse

| | | | | | | | | |
|---|---|---|---|---|---|---|---|---|
| Q8CHS7 | Q8VCR2 | 0.474 | | | FALSE | TRUE | FALSE | FALSE |
| Q3TES0 | Q8R0S2 | 0.474 | | HOM | TRUE | FALSE | FALSE | FALSE |
| O88451 | Q80XN0 | 0.475 | | | FALSE | TRUE | FALSE | FALSE |
| P57722 | P61979 | 0.476 | | HOM | TRUE | FALSE | FALSE | FALSE |
| Q9QYB5 | Q9QYB8 | 0.476 | | | FALSE | TRUE | FALSE | FALSE |
| P31001 | P48678 | 0.478 | | HOM | TRUE | FALSE | FALSE | FALSE |
| P14873 | Q9QYR6 | 0.478 | | | FALSE | TRUE | FALSE | FALSE |
| Q6PDL0 | Q8R1Q8 | 0.479 | HOM | HOM | TRUE | FALSE | FALSE | FALSE |
| P33267 | Q64458 | 0.479 | | | FALSE | TRUE | FALSE | FALSE |
| P61750 | Q9D0J4 | 0.479 | | HET | FALSE | FALSE | FALSE | FALSE |
| Q8BLR2 | Q8C166 | 0.479 | | HOM | TRUE | FALSE | FALSE | FALSE |
| P21300 | P70694 | 0.48 | | HOM | TRUE | FALSE | FALSE | FALSE |
| Q62108 | Q9Z0U1 | 0.48 | HOM | HOM | TRUE | FALSE | FALSE | FALSE |
| P24456 | P24457 | 0.481 | HOM | HOM | TRUE | FALSE | FALSE | FALSE |
| P56656 | Q6XVG2 | 0.481 | | | FALSE | TRUE | FALSE | FALSE |
| A2ASZ8 | Q8BMD8 | 0.482 | | | FALSE | TRUE | FALSE | FALSE |
| Q61879 | Q9JMH9 | 0.482 | | | FALSE | TRUE | FALSE | FALSE |
| Q6GV12 | Q99J47 | 0.483 | | | FALSE | TRUE | FALSE | FALSE |
| Q811D0 | Q9WV34 | 0.484 | HOM | HOM | TRUE | FALSE | FALSE | FALSE |
| Q8C854 | Q9D0E1 | 0.484 | | | FALSE | TRUE | FALSE | FALSE |
| P00329 | Q9QYY9 | 0.484 | | HOM | TRUE | FALSE | FALSE | FALSE |
| Q8BGA8 | Q91VA0 | 0.484 | | | FALSE | TRUE | FALSE | FALSE |
| Q8BLR2 | Q9Z140 | 0.485 | | | FALSE | TRUE | FALSE | FALSE |
| Q62318 | Q9R1R2 | 0.486 | | | FALSE | TRUE | FALSE | FALSE |
| Q3U0V1 | Q91WJ8 | 0.486 | | HET | FALSE | FALSE | FALSE | FALSE |
| P21836 | Q8BYM5 | 0.486 | HOM | HET | TRUE | FALSE | FALSE | FALSE |

Table S4 Mouse

| | | | | | | | | | | | |
|---|---|---|---|---|---|---|---|---|---|---|---|
| P63085 | Q63844 | 0.486 | HOM | HOM | TRUE | FALSE | TRUE | 5227,6799,6800,6801,6802 | Mouse,Human | p14-Mp1-Erk1/2 complex, VEGFR2-S1PR1-ERK1/2-PKC-alpha complex, VEGFR2-S1PR2-ERK1/2-PKC-alpha complex, VEGFR2-S1PR3-ERK1/2-PKC-alpha complex, VEGFR2-S1PR5-ERK1/2-PKC-alpha complex | FALSE |
| O35488 | Q4LDG0 | 0.489 | | | FALSE | TRUE | FALSE | | | | FALSE |
| Q5DQR4 | Q80Y17 | 0.489 | | | FALSE | TRUE | FALSE | | | | FALSE |
| Q60931 | Q60932 | 0.49 | | | FALSE | TRUE | FALSE | | | | FALSE |
| Q8C3X2 | Q9CXD6 | 0.491 | | | FALSE | TRUE | FALSE | | | | FALSE |
| P21958 | P36371 | 0.491 | | | FALSE | TRUE | FALSE | | | | FALSE |
| P97742 | Q9DC50 | 0.491 | HOM | | TRUE | FALSE | FALSE | | | | FALSE |
| P21300 | Q8VCX1 | 0.492 | | | FALSE | TRUE | FALSE | | | | FALSE |
| P61028 | P62823 | 0.493 | | | FALSE | TRUE | FALSE | | | | FALSE |
| O35945 | Q9CZS1 | 0.493 | | | FALSE | TRUE | FALSE | | | | FALSE |
| P09103 | Q921X9 | 0.493 | HOM | | TRUE | FALSE | FALSE | | | | FALSE |
| Q9QYB1 | Q9Z1Q5 | 0.495 | | HOM | TRUE | FALSE | FALSE | | | | FALSE |
| Q7TMM9 | Q9D6F9 | 0.495 | HET | HET | FALSE | FALSE | FALSE | | | | FALSE |

Table S4 Mouse

| | | | | | | | | |
|---|---|---|---|---|---|---|---|---|
| P48758 | Q8K354 | 0.495 | | | FALSE | TRUE | FALSE | FALSE |
| P18654 | Q60823 | 0.496 | | | FALSE | TRUE | FALSE | FALSE |
| Q3UHD9 | Q7SIG6 | 0.497 | | | FALSE | TRUE | FALSE | FALSE |
| P10605 | Q9WUU7 | 0.497 | HET | | FALSE | FALSE | FALSE | FALSE |
| O88428 | Q60967 | 0.498 | | | FALSE | TRUE | FALSE | FALSE |
| P10649 | P15626 | 0.498 | | | FALSE | TRUE | FALSE | FALSE |
| P29699 | Q9QXC1 | 0.5 | | | FALSE | TRUE | FALSE | FALSE |
| O70161 | Q80XI4 | 0.5 | | | FALSE | TRUE | FALSE | FALSE |
| Q8QZR3 | Q91WG0 | 0.501 | | | FALSE | TRUE | FALSE | FALSE |
| P60764 | P63001 | 0.502 | | | FALSE | TRUE | FALSE | FALSE |
| P61027 | Q9CZT8 | 0.502 | | | FALSE | TRUE | FALSE | FALSE |
| Q62465 | Q80TB8 | 0.502 | | HOM | TRUE | FALSE | FALSE | FALSE |
| Q6PER3 | Q8R001 | 0.502 | | | FALSE | TRUE | FALSE | FALSE |
| O70571 | Q8BFP9 | 0.503 | | HOM | TRUE | FALSE | FALSE | FALSE |
| Q3UEG6 | Q8BWU8 | 0.503 | | | FALSE | TRUE | FALSE | FALSE |
| Q61166 | Q8R001 | 0.505 | | | FALSE | TRUE | FALSE | FALSE |
| Q99JF8 | Q9JMG7 | 0.505 | | | FALSE | TRUE | FALSE | FALSE |
| P50171 | Q8VCC1 | 0.506 | HOM | HOM | TRUE | FALSE | FALSE | FALSE |
| P13542 | Q6URW6 | 0.507 | | | FALSE | TRUE | FALSE | FALSE |
| P12815 | Q6P069 | 0.507 | HOM | | TRUE | FALSE | FALSE | FALSE |
| Q5H8C4 | Q8BX70 | 0.507 | | | FALSE | TRUE | FALSE | FALSE |
| O35660 | P10649 | 0.508 | | | FALSE | TRUE | FALSE | FALSE |
| P24549 | Q8R0Y6 | 0.508 | | | FALSE | TRUE | FALSE | FALSE |
| Q8BT60 | Q8C166 | 0.508 | | HOM | TRUE | FALSE | FALSE | FALSE |
| P13541 | Q5SX39 | 0.508 | | | FALSE | TRUE | FALSE | FALSE |
| O09161 | O09165 | 0.509 | HOM | HOM | TRUE | FALSE | FALSE | FALSE |

Table S4 Mouse

| | | | | | | | | | | | |
|---|---|---|---|---|---|---|---|---|---|---|---|
| Q99J47 | Q9CXR1 | 0.51 | | | FALSE | TRUE | FALSE | | | | FALSE |
| P47857 | Q9WUA3 | 0.51 | HET | HET | FALSE | FALSE | FALSE | | | | FALSE |
| P15626 | P19639 | 0.511 | | | FALSE | TRUE | FALSE | | | | FALSE |
| P07724 | P21614 | 0.511 | HOM | | TRUE | FALSE | FALSE | | | | FALSE |
| P62748 | Q91X97 | 0.513 | HET | HET | FALSE | FALSE | FALSE | | | | FALSE |
| P01027 | P28665 | 0.514 | HOM | | TRUE | FALSE | FALSE | | | | FALSE |
| Q9QXD1 | Q9R0H0 | 0.514 | | | FALSE | TRUE | FALSE | | | | FALSE |
| Q8K009 | Q8R0Y6 | 0.514 | | | FALSE | TRUE | FALSE | | | | FALSE |
| P19221 | P20918 | 0.515 | | HOM | TRUE | FALSE | FALSE | | | | FALSE |
| Q6P1F6 | Q925E7 | 0.516 | HET | HET | FALSE | FALSE | FALSE | | | | FALSE |
| Q61166 | Q6PER3 | 0.516 | | | FALSE | TRUE | FALSE | | | | FALSE |
| O08553 | Q62188 | 0.516 | HOM | HET | TRUE | FALSE | FALSE | | | | FALSE |
| Q2NL51 | Q9WV60 | 0.516 | | | FALSE | TRUE | FALSE | | | | FALSE |
| P11714 | P24457 | 0.517 | HOM | HOM | TRUE | FALSE | FALSE | | | | FALSE |
| Q64458 | Q91W64 | 0.517 | | | FALSE | TRUE | FALSE | | | | FALSE |
| O70443 | Q9DC51 | 0.518 | | HET | FALSE | FALSE | FALSE | | | | FALSE |
| P63168 | Q9D0M5 | 0.518 | | HOM | TRUE | FALSE | FALSE | | | | FALSE |
| P50608 | P51885 | 0.518 | | | FALSE | TRUE | FALSE | | | | FALSE |
| Q5FW57 | Q91XE0 | 0.518 | | | FALSE | TRUE | FALSE | | | | FALSE |
| Q91ZE0 | Q924Y0 | 0.518 | | HOM | TRUE | FALSE | FALSE | | | | FALSE |
| P23953 | Q63880 | 0.519 | | | FALSE | TRUE | FALSE | | | | FALSE |
| P60521 | Q9CQV6 | 0.519 | HOM | HOM | TRUE | FALSE | FALSE | | | | FALSE |
| Q3ULF4 | Q8JZQ2 | 0.519 | HET | HET | FALSE | FALSE | FALSE | | | | FALSE |
| P12815 | Q8BFY6 | 0.519 | HOM | HET | TRUE | FALSE | TRUE | 6375 | Human | PEF1-ALG2 complex | FALSE |
| O88876 | Q9CXR1 | 0.519 | | | FALSE | TRUE | FALSE | | | | FALSE |

Table S4 Mouse

| | | | | | | | | |
|---|---|---|---|---|---|---|---|---|
| P70694 | Q8K023 | 0.52 | HOM | HOM | TRUE | FALSE | FALSE | FALSE |
| P63040 | P84086 | 0.52 | | | FALSE | TRUE | FALSE | FALSE |
| Q8BGT8 | Q8K0S0 | 0.52 | | | FALSE | TRUE | FALSE | FALSE |
| P19246 | P21619 | 0.521 | | | FALSE | TRUE | FALSE | FALSE |
| P97447 | Q8CI51 | 0.521 | | | FALSE | TRUE | FALSE | FALSE |
| P14733 | P20152 | 0.521 | | | FALSE | TRUE | FALSE | FALSE |
| P70297 | Q5SRX1 | 0.521 | | | FALSE | TRUE | FALSE | FALSE |
| Q80Z24 | Q99PJ0 | 0.522 | | | FALSE | TRUE | FALSE | FALSE |
| O70443 | P18872 | 0.523 | | | FALSE | TRUE | FALSE | FALSE |
| O55028 | Q9JK42 | 0.523 | | HOM | TRUE | FALSE | FALSE | FALSE |
| Q8CI51 | Q9R059 | 0.526 | | | FALSE | TRUE | FALSE | FALSE |
| Q3TMH2 | Q8VCA8 | 0.526 | | | FALSE | TRUE | FALSE | FALSE |
| P84075 | Q8BGZ1 | 0.527 | HET | | FALSE | FALSE | FALSE | FALSE |
| Q9ESM3 | Q9QUP5 | 0.527 | | | FALSE | TRUE | FALSE | FALSE |
| P62761 | Q8BGZ1 | 0.529 | | | FALSE | TRUE | FALSE | FALSE |
| O35660 | P19639 | 0.53 | | | FALSE | TRUE | FALSE | FALSE |
| P97927 | Q60675 | 0.531 | | HOM | TRUE | FALSE | FALSE | FALSE |
| Q60930 | Q60932 | 0.531 | | | FALSE | TRUE | FALSE | FALSE |
| P52430 | Q62087 | 0.531 | | | FALSE | TRUE | FALSE | FALSE |
| O08807 | P35700 | 0.532 | HOM | HOM | TRUE | FALSE | FALSE | FALSE |
| Q6ZQ82 | Q91YM2 | 0.532 | | | FALSE | TRUE | FALSE | FALSE |
| P23818 | Q9Z2W9 | 0.533 | HET | HET | FALSE | FALSE | FALSE | FALSE |
| P61226 | Q80ZJ1 | 0.533 | | HOM | TRUE | FALSE | FALSE | FALSE |
| Q8C052 | Q9QYR6 | 0.533 | | | FALSE | TRUE | FALSE | FALSE |
| P62761 | Q91X97 | 0.534 | | HET | FALSE | FALSE | FALSE | FALSE |
| Q8R5M8 | Q99N28 | 0.534 | | HOM | TRUE | FALSE | FALSE | FALSE |

Table S4 Mouse

| | | | | | | | | | | | |
|---|---|---|---|---|---|---|---|---|---|---|---|
| O35945 | Q8R0Y6 | 0.535 | | | FALSE | TRUE | FALSE | | | | FALSE |
| P22599 | Q00897 | 0.535 | HOM | HOM | TRUE | FALSE | FALSE | | | | FALSE |
| P35585 | P84091 | 0.535 | HET | HET | FALSE | FALSE | FALSE | | | | FALSE |
| P35282 | Q9CQD1 | 0.536 | | | FALSE | TRUE | FALSE | | | | FALSE |
| Q02566 | Q6URW6 | 0.536 | | | FALSE | TRUE | FALSE | | | | FALSE |
| O35639 | P48036 | 0.537 | | | FALSE | TRUE | FALSE | | | | FALSE |
| P20065 | Q6ZWY8 | 0.537 | | | FALSE | TRUE | FALSE | | | | FALSE |
| Q9JKB1 | Q9R0P9 | 0.538 | HET | HET | FALSE | FALSE | FALSE | | | | FALSE |
| Q91VA0 | Q9QXG4 | 0.539 | | | FALSE | TRUE | FALSE | | | | FALSE |
| P99024 | Q9D6F9 | 0.54 | HET | HET | FALSE | FALSE | FALSE | | | | FALSE |
| P08228 | Q9WU84 | 0.541 | HOM | HOM | TRUE | FALSE | FALSE | | | | FALSE |
| Q62188 | Q9EQF6 | 0.541 | HET | HET | FALSE | FALSE | FALSE | | | | TRUE |
| Q9JLB0 | Q9Z0U1 | 0.542 | | HOM | TRUE | FALSE | FALSE | | | | FALSE |
| P61290 | P97371 | 0.543 | | | FALSE | TRUE | FALSE | | | | FALSE |
| Q64459 | Q9DBG1 | 0.543 | | | FALSE | TRUE | FALSE | | | | FALSE |
| O88876 | Q9EQ06 | 0.544 | | | FALSE | TRUE | FALSE | | | | FALSE |
| P35438 | Q9Z2W9 | 0.546 | HET | HET | FALSE | FALSE | FALSE | | | | FALSE |
| P35278 | Q9CQD1 | 0.547 | | | FALSE | TRUE | FALSE | | | | FALSE |
| P51658 | Q80XN0 | 0.548 | | | FALSE | TRUE | FALSE | | | | FALSE |
| Q80U63 | Q811U4 | 0.548 | HET | HOM | TRUE | FALSE | FALSE | | | | TRUE |
| P07758 | Q00897 | 0.548 | HOM | HOM | TRUE | FALSE | FALSE | | | | FALSE |
| P22723 | P62812 | 0.548 | | | FALSE | TRUE | TRUE | 5809 | Human | GABAA receptor | TRUE |
| P10649 | P19639 | 0.548 | | | FALSE | TRUE | FALSE | | | | FALSE |
| P35585 | Q9JKC8 | 0.55 | HET | HET | FALSE | FALSE | FALSE | | | | FALSE |
| P12658 | Q08331 | 0.553 | | | FALSE | TRUE | FALSE | | | | FALSE |
| P33587 | Q61646 | 0.553 | | | FALSE | TRUE | FALSE | | | | FALSE |

Table S4 Mouse

| | | | | | | | | |
|---|---|---|---|---|---|---|---|---|
| P35762 | P40237 | 0.553 | HOM | | TRUE | FALSE | FALSE | FALSE |
| P15626 | P19157 | 0.554 | | HOM | TRUE | FALSE | FALSE | FALSE |
| Q91XE0 | Q9DCY0 | 0.556 | | | FALSE | TRUE | FALSE | FALSE |
| O88451 | Q9R092 | 0.556 | | | FALSE | TRUE | FALSE | FALSE |
| P35282 | P61021 | 0.557 | | | FALSE | TRUE | FALSE | FALSE |
| P21300 | Q8VC28 | 0.557 | | HOM | TRUE | FALSE | FALSE | FALSE |
| P03995 | P08551 | 0.557 | | | FALSE | TRUE | FALSE | FALSE |
| Q9EQP2 | Q9WVK4 | 0.558 | HET | HET | FALSE | FALSE | FALSE | FALSE |
| P45878 | Q9D1M7 | 0.558 | | | FALSE | TRUE | FALSE | FALSE |
| P06909 | Q01339 | 0.559 | | HET | FALSE | FALSE | FALSE | FALSE |
| P07356 | P14824 | 0.561 | HOM | | TRUE | FALSE | FALSE | FALSE |
| P28653 | P50608 | 0.564 | | | FALSE | TRUE | FALSE | FALSE |
| P70402 | Q3KNY0 | 0.564 | | | FALSE | TRUE | FALSE | FALSE |
| Q8BH64 | Q9EQP2 | 0.565 | HOM, HET | HET | TRUE, FALSE | FALSE | FALSE | FALSE |
| P50608 | Q99MQ4 | 0.566 | | | FALSE | TRUE | FALSE | FALSE |
| P09542 | Q8CI43 | 0.566 | | | FALSE | TRUE | FALSE | FALSE |
| P23818 | P35438 | 0.566 | HET | HET | FALSE | FALSE | FALSE | TRUE |
| P28652 | Q3UHL1 | 0.567 | | | FALSE | TRUE | FALSE | FALSE |
| P47809 | Q63932 | 0.567 | | | FALSE | TRUE | FALSE | FALSE |
| P51830 | P84309 | 0.567 | | | FALSE | TRUE | FALSE | FALSE |
| P08074 | Q91X52 | 0.568 | HOM | HOM | TRUE | FALSE | FALSE | FALSE |
| Q6URW6 | Q91Z83 | 0.568 | | | FALSE | TRUE | FALSE | FALSE |
| P29387 | Q8CGF6 | 0.57 | | | FALSE | TRUE | FALSE | FALSE |
| Q8C1B7 | Q8CHH9 | 0.57 | | HET | FALSE | FALSE | FALSE | FALSE |
| P61164 | Q8R5C5 | 0.57 | | | FALSE | TRUE | FALSE | FALSE |

Table S4 Mouse

| | | | | | | | | |
|---|---|---|---|---|---|---|---|---|
| Q3ULD5 | Q99MN9 | 0.57 | | | FALSE | TRUE | FALSE | FALSE |
| P56593 | Q6XVG2 | 0.571 | | | FALSE | TRUE | FALSE | FALSE |
| O70250 | P15327 | 0.572 | | HOM | TRUE | FALSE | FALSE | FALSE |
| P61982 | P63101 | 0.572 | | | FALSE | TRUE | FALSE | TRUE |
| P24721 | P34927 | 0.572 | | | FALSE | TRUE | FALSE | FALSE |
| A2AQP0 | Q5SX40 | 0.573 | | | FALSE | TRUE | FALSE | FALSE |
| Q8VCZ9 | Q9WU79 | 0.573 | | | FALSE | TRUE | FALSE | FALSE |
| P24549 | Q8K009 | 0.573 | | | FALSE | TRUE | FALSE | FALSE |
| P16015 | P23589 | 0.573 | | | FALSE | TRUE | FALSE | FALSE |
| Q8R464 | Q99N28 | 0.574 | HOM | HOM | TRUE | FALSE | FALSE | FALSE |
| Q925N0 | Q925N1 | 0.575 | | | FALSE | TRUE | FALSE | FALSE |
| Q8BK48 | Q91WG0 | 0.575 | | | FALSE | TRUE | FALSE | FALSE |
| A2AQP0 | Q02566 | 0.576 | | | FALSE | TRUE | FALSE | FALSE |
| Q3UHL1 | Q923T9 | 0.576 | | | FALSE | TRUE | FALSE | FALSE |
| P61205 | P62331 | 0.577 | | HOM | TRUE | FALSE | FALSE | FALSE |
| Q8BG39 | Q9JIS5 | 0.577 | | | FALSE | TRUE | FALSE | FALSE |
| Q3TJD7 | Q9R059 | 0.58 | | | FALSE | TRUE | FALSE | FALSE |
| O35728 | Q91WL5 | 0.58 | | | FALSE | TRUE | FALSE | FALSE |
| P99024 | Q922F4 | 0.58 | HET | | FALSE | FALSE | FALSE | FALSE |
| P26150 | Q9R1J0 | 0.58 | | | FALSE | TRUE | FALSE | FALSE |
| P23927 | Q99PR8 | 0.583 | HOM | | TRUE | FALSE | FALSE | FALSE |
| Q91WL5 | Q9EP75 | 0.584 | | | FALSE | TRUE | FALSE | FALSE |
| P14094 | P14231 | 0.585 | | | FALSE | TRUE | FALSE | FALSE |
| Q6PIC6 | Q6PIE5 | 0.585 | | | FALSE | TRUE | FALSE | FALSE |
| Q8BGT5 | Q8QZR5 | 0.585 | | | FALSE | TRUE | FALSE | FALSE |
| Q9CXE7 | Q9R0Q3 | 0.585 | | | FALSE | TRUE | FALSE | FALSE |

Table S4 Mouse

| | | | | | | | | | | | |
|---|---|---|---|---|---|---|---|---|---|---|---|
| A2AKK5 | Q8BWN8 | 0.586 | | | FALSE | TRUE | FALSE | | | | FALSE |
| Q99JR5 | Q9WUU7 | 0.586 | | | FALSE | TRUE | FALSE | | | | FALSE |
| P48453 | P63328 | 0.586 | | | FALSE | TRUE | FALSE | | | | FALSE |
| P63213 | Q9CXP8 | 0.586 | HET | | FALSE | FALSE | FALSE | | | | FALSE |
| P49442 | Q9Z0S1 | 0.586 | | | FALSE | TRUE | FALSE | | | | FALSE |
| Q62452 | Q63886 | 0.587 | | | FALSE | TRUE | FALSE | | | | FALSE |
| Q8BH59 | Q9D6M3 | 0.587 | | | FALSE | TRUE | FALSE | | | | FALSE |
| P08074 | Q99LB2 | 0.587 | HOM | | TRUE | FALSE | FALSE | | | | FALSE |
| P61982 | Q9CQV8 | 0.588 | | | FALSE | TRUE | TRUE | 5199 | Human | Kinase maturation complex 1 | FALSE |
| Q9JJL3 | Q9QXZ6 | 0.588 | | | FALSE | TRUE | FALSE | | | | FALSE |
| P70402 | Q5XKE0 | 0.589 | | | FALSE | TRUE | FALSE | | | | FALSE |
| Q99JB8 | Q9WVE8 | 0.589 | HOM | | TRUE | FALSE | FALSE | | | | FALSE |
| Q61016 | Q9DAS9 | 0.589 | | | FALSE | TRUE | FALSE | | | | FALSE |
| P08551 | P46660 | 0.59 | | | FALSE | TRUE | FALSE | | | | FALSE |
| P97447 | Q3TJD7 | 0.59 | | | FALSE | TRUE | FALSE | | | | FALSE |
| P11588 | Q5FW60 | 0.592 | | | FALSE | TRUE | FALSE | | | | FALSE |
| O88833 | Q91WL5 | 0.593 | | | FALSE | TRUE | FALSE | | | | FALSE |
| B2RQC6 | Q8C196 | 0.594 | | HOM | TRUE | FALSE | FALSE | | | | FALSE |
| P70694 | Q8VCX1 | 0.594 | HOM | | TRUE | FALSE | FALSE | | | | FALSE |
| P48771 | P56392 | 0.594 | | | FALSE | TRUE | FALSE | | | | FALSE |
| Q3UNX5 | Q8BGA8 | 0.594 | | | FALSE | TRUE | FALSE | | | | FALSE |
| P28654 | P50608 | 0.595 | | | FALSE | TRUE | FALSE | | | | FALSE |
| P62835 | Q5PR73 | 0.595 | | | FALSE | TRUE | FALSE | | | | FALSE |
| P97447 | Q9R059 | 0.596 | | | FALSE | TRUE | FALSE | | | | FALSE |

Table S4 Mouse

| | | | | | | | | |
|---|---|---|---|---|---|---|---|---|
| Q8JZN5 | Q9JHI5 | 0.597 | | | FALSE | TRUE | FALSE | FALSE |
| P28474 | Q9QYY9 | 0.597 | HOM | HOM | TRUE | FALSE | FALSE | FALSE |
| P11798 | Q3UHL1 | 0.598 | | | FALSE | TRUE | FALSE | FALSE |
| P62748 | P84075 | 0.598 | HET | HET | FALSE | FALSE | FALSE | FALSE |
| P18760 | Q9R0P5 | 0.599 | | | FALSE | TRUE | FALSE | FALSE |
| P63216 | Q61016 | 0.599 | | | FALSE | TRUE | FALSE | FALSE |
| Q3UP75 | Q8JZZ0 | 0.599 | | | FALSE | TRUE | FALSE | FALSE |
| Q8CHH9 | Q9R1T4 | 0.599 | HET | HOM | TRUE | FALSE | FALSE | FALSE |
| P26150 | Q9EQC1 | 0.599 | | | FALSE | TRUE | FALSE | FALSE |
| Q63880 | Q91WG0 | 0.599 | | | FALSE | TRUE | FALSE | FALSE |
| P07356 | P48036 | 0.602 | HOM | | TRUE | FALSE | FALSE | FALSE |
| Q05421 | Q6XVG2 | 0.602 | | | FALSE | TRUE | FALSE | FALSE |
| O70172 | Q80XI4 | 0.604 | | | FALSE | TRUE | FALSE | FALSE |
| O35639 | P14824 | 0.607 | | | FALSE | TRUE | FALSE | FALSE |
| O88876 | Q8VCR2 | 0.607 | | | FALSE | TRUE | FALSE | FALSE |
| P22599 | Q00898 | 0.608 | HOM | HOM | TRUE | FALSE | FALSE | FALSE |
| Q62420 | Q8R3V5 | 0.61 | HOM | | TRUE | FALSE | FALSE | FALSE |
| P33267 | P56654 | 0.611 | | | FALSE | TRUE | FALSE | FALSE |
| P97447 | Q9JKS4 | 0.611 | | | FALSE | TRUE | FALSE | FALSE |
| Q9ET78 | Q9ET80 | 0.611 | | | FALSE | TRUE | FALSE | FALSE |
| O88533 | P48318 | 0.611 | HOM | HOM | TRUE | FALSE | FALSE | FALSE |
| Q9ET54 | Q9JIF9 | 0.612 | | | FALSE | TRUE | FALSE | FALSE |
| O70622 | Q9ES97 | 0.612 | | | FALSE | TRUE | FALSE | FALSE |
| O55143 | Q8VDN2 | 0.614 | | | FALSE | TRUE | FALSE | FALSE |
| P01029 | P28665 | 0.615 | | | FALSE | TRUE | FALSE | FALSE |
| Q8BXB6 | Q9JJL3 | 0.615 | | | FALSE | TRUE | FALSE | FALSE |

Table S4 Mouse

| | | | | | | | | | | | |
|---|---|---|---|---|---|---|---|---|---|---|---|
| P97927 | Q61292 | 0.615 | | | FALSE | TRUE | FALSE | | | | FALSE |
| P24549 | Q9CZS1 | 0.615 | | | FALSE | TRUE | FALSE | | | | FALSE |
| O88952 | Q8JZS0 | 0.616 | | | FALSE | TRUE | FALSE | | | | FALSE |
| P18872 | P21279 | 0.616 | | | FALSE | TRUE | FALSE | | | | FALSE |
| O55028 | O70571 | 0.617 | | | FALSE | TRUE | FALSE | | | | FALSE |
| P47199 | Q9DCS3 | 0.617 | HOM | HOM | TRUE | FALSE | FALSE | | | | FALSE |
| P49817 | Q9WVC3 | 0.617 | | HOM | TRUE | FALSE | FALSE | | | | FALSE |
| A2AQP0 | Q91Z83 | 0.618 | | | FALSE | TRUE | FALSE | | | | FALSE |
| P21956 | Q61147 | 0.619 | | | FALSE | TRUE | FALSE | | | | FALSE |
| Q64176 | Q8QZR3 | 0.619 | | | FALSE | TRUE | FALSE | | | | FALSE |
| Q8BXA5 | Q8VBZ3 | 0.621 | | | FALSE | TRUE | FALSE | | | | FALSE |
| P56654 | Q91W64 | 0.622 | | | FALSE | TRUE | FALSE | | | | FALSE |
| P14602 | Q9JK92 | 0.623 | | | FALSE | TRUE | FALSE | | | | FALSE |
| P61205 | Q9WUL7 | 0.625 | | | FALSE | TRUE | FALSE | | | | FALSE |
| P21279 | P27601 | 0.625 | | | FALSE | TRUE | FALSE | | | | FALSE |
| P01898 | P01899 | 0.626 | HET,HOM | | FALSE,TRUE | FALSE | FALSE | | | | FALSE |
| Q62433 | Q8BTG7 | 0.626 | | | FALSE | TRUE | FALSE | | | | FALSE |
| Q63880 | Q8BK48 | 0.627 | | | FALSE | TRUE | FALSE | | | | FALSE |
| P31650 | Q61327 | 0.628 | | | FALSE | TRUE | FALSE | | | | FALSE |
| O35660 | P15626 | 0.631 | | | FALSE | TRUE | FALSE | | | | FALSE |
| P68134 | Q8R5C5 | 0.631 | | | FALSE | TRUE | FALSE | | | | FALSE |
| P62259 | P68510 | 0.632 | HOM | | TRUE | FALSE | TRUE | 5199 | Human | Kinase maturation complex 1 | TRUE |
| Q60598 | Q9QXS6 | 0.634 | | | FALSE | TRUE | FALSE | | | | FALSE |
| O70443 | P27601 | 0.634 | | | FALSE | TRUE | FALSE | | | | FALSE |

Table S4 Mouse

| | | | | | | | | |
|---|---|---|---|---|---|---|---|---|
| Q9R1V6 | Q9R1V7 | 0.634 | | | FALSE | TRUE | FALSE | FALSE |
| P32020 | P51660 | 0.634 | HET | HOM | TRUE | FALSE | FALSE | FALSE |
| P13542 | Q91Z83 | 0.635 | | | FALSE | TRUE | FALSE | FALSE |
| P61967 | P62743 | 0.635 | HET | HET | FALSE | FALSE | FALSE | FALSE |
| Q7TQ48 | Q9EQP2 | 0.636 | HET | HET | FALSE | FALSE | FALSE | FALSE |
| Q14DH7 | Q9D2R0 | 0.636 | | | FALSE | TRUE | FALSE | FALSE |
| P40237 | Q922J6 | 0.636 | | | FALSE | TRUE | FALSE | FALSE |
| Q9JKS4 | Q9R059 | 0.637 | | | FALSE | TRUE | FALSE | FALSE |
| P01898 | P01901 | 0.637 | HET,HOM | HOM | TRUE | FALSE | FALSE | FALSE |
| Q9QYI5 | Q9QYJ3 | 0.638 | HOM | HOM | TRUE | FALSE | FALSE | FALSE |
| Q3UQ84 | Q9JKF7 | 0.639 | | | FALSE | TRUE | FALSE | FALSE |
| P23927 | Q9JK92 | 0.64 | HOM | | TRUE | FALSE | FALSE | FALSE |
| Q8VCU1 | Q91WG0 | 0.642 | | | FALSE | TRUE | FALSE | FALSE |
| P18872 | P27601 | 0.642 | | | FALSE | TRUE | FALSE | FALSE |
| Q8BK48 | Q8VCT4 | 0.644 | | | FALSE | TRUE | FALSE | FALSE |
| P13412 | P48787 | 0.644 | | | FALSE | TRUE | FALSE | FALSE |
| P08074 | Q8VCC1 | 0.645 | HOM | HOM | TRUE | FALSE | FALSE | FALSE |
| P84078 | P84084 | 0.648 | | | FALSE | TRUE | FALSE | FALSE |
| P51658 | Q9R092 | 0.648 | | | FALSE | TRUE | FALSE | FALSE |
| Q9D6M3 | Q9DB41 | 0.649 | | | FALSE | TRUE | FALSE | FALSE |
| Q6P069 | Q8BFY6 | 0.649 | | HET | FALSE | FALSE | FALSE | FALSE |
| Q3UNX5 | Q91VA0 | 0.65 | | | FALSE | TRUE | FALSE | FALSE |
| Q05421 | Q64458 | 0.65 | | | FALSE | TRUE | FALSE | FALSE |
| P70318 | Q8VH51 | 0.651 | | | FALSE | TRUE | FALSE | FALSE |
| O35660 | P19157 | 0.652 | | HOM | TRUE | FALSE | FALSE | FALSE |
| P08113 | P11499 | 0.652 | HOM | HOM | TRUE | FALSE | FALSE | FALSE |

Table S4 Mouse

| | | | | | | | | | | | |
|---|---|---|---|---|---|---|---|---|---|---|---|
| O35945 | P47738 | 0.653 | | HOM | TRUE | FALSE | FALSE | | | | FALSE |
| P61028 | Q9CZT8 | 0.653 | | | FALSE | TRUE | FALSE | | | | FALSE |
| O35136 | Q99PJ0 | 0.653 | HOM | | TRUE | FALSE | FALSE | | | | FALSE |
| P61982 | P62259 | 0.653 | | HOM | TRUE | FALSE | TRUE | 5199 | Human | Kinase maturation complex 1 | TRUE |
| P47738 | Q9CZS1 | 0.653 | HOM | | TRUE | FALSE | FALSE | | | | FALSE |
| P35278 | P61021 | 0.654 | | | FALSE | TRUE | FALSE | | | | FALSE |
| Q05421 | Q91W64 | 0.655 | | | FALSE | TRUE | FALSE | | | | FALSE |
| P16546 | Q62261 | 0.655 | HOM | HOM | TRUE | FALSE | FALSE | | | | FALSE |
| Q78IK4 | Q9DCZ4 | 0.655 | | | FALSE | TRUE | TRUE | 6249,6255 | Human | MIB complex, MICOS complex | FALSE |
| Q07417 | Q8JZN5 | 0.656 | HOM | | TRUE | FALSE | FALSE | | | | FALSE |
| P09041 | P09411 | 0.656 | | | FALSE | TRUE | FALSE | | | | FALSE |
| P10833 | P63321 | 0.656 | | | FALSE | TRUE | FALSE | | | | FALSE |
| P62821 | Q6PHN9 | 0.658 | | | FALSE | TRUE | FALSE | | | | FALSE |
| P61205 | P84084 | 0.658 | | | FALSE | TRUE | FALSE | | | | FALSE |
| P61028 | P63011 | 0.658 | | | FALSE | TRUE | FALSE | | | | FALSE |
| B2RSH2 | P08752 | 0.659 | HOM | | TRUE | FALSE | FALSE | | | | FALSE |
| P70175 | Q9Z0U1 | 0.659 | HOM | HOM | TRUE | FALSE | FALSE | | | | FALSE |
| P35279 | P61294 | 0.66 | | | FALSE | TRUE | FALSE | | | | FALSE |
| O08788 | Q9Z0H8 | 0.66 | HOM | | TRUE | FALSE | FALSE | | | | FALSE |
| P31938 | P47809 | 0.661 | | | FALSE | TRUE | FALSE | | | | FALSE |
| P17809 | P32037 | 0.661 | | | FALSE | TRUE | FALSE | | | | FALSE |
| Q8BG05 | Q9CX86 | 0.661 | | | FALSE | TRUE | FALSE | | | | FALSE |
| P62137 | P63087 | 0.661 | HOM | | TRUE | FALSE | FALSE | | | | FALSE |

Table S4 Mouse

| | | | | | | | | | |
|---|---|---|---|---|---|---|---|---|---|
| Q6XVG2 | Q91W64 | 0.662 | | | FALSE | TRUE | FALSE | | FALSE |
| P97821 | Q9WUU7 | 0.662 | HET | | FALSE | FALSE | FALSE | | FALSE |
| Q99P58 | Q9CZT8 | 0.662 | HET | | FALSE | FALSE | FALSE | | FALSE |
| P28663 | Q9DB05 | 0.663 | | | FALSE | TRUE | FALSE | | FALSE |
| Q8BMF3 | Q99KE1 | 0.663 | | HOM | TRUE | FALSE | FALSE | | FALSE |
| O55042 | Q91ZZ3 | 0.663 | | | FALSE | TRUE | FALSE | | FALSE |
| O88569 | Q9CX86 | 0.663 | | | FALSE | TRUE | FALSE | | FALSE |
| Q9CXJ4 | Q9JI39 | 0.665 | | HOM | TRUE | FALSE | FALSE | | FALSE |
| Q8BLQ9 | Q99N28 | 0.665 | HOM | HOM | TRUE | FALSE | FALSE | | FALSE |
| P08207 | P50543 | 0.667 | | | FALSE | TRUE | FALSE | | FALSE |
| Q99LB7 | Q9DBT9 | 0.667 | | | FALSE | TRUE | FALSE | | FALSE |
| Q8BFP9 | Q9JK42 | 0.667 | HOM | HOM | TRUE | FALSE | FALSE | | FALSE |
| Q64176 | Q8VCT4 | 0.668 | | | FALSE | TRUE | FALSE | | FALSE |
| P58771 | P58774 | 0.668 | | | FALSE | TRUE | FALSE | | FALSE |
| P97364 | Q8BH69 | 0.668 | | HOM | TRUE | FALSE | FALSE | | FALSE |
| Q61694 | Q9R1J0 | 0.668 | | | FALSE | TRUE | FALSE | | FALSE |
| Q8VC28 | Q8VCX1 | 0.669 | HOM | | TRUE | FALSE | FALSE | | FALSE |
| Q5PR73 | Q99JI6 | 0.67 | | | FALSE | TRUE | FALSE | | FALSE |
| Q60675 | Q61292 | 0.67 | HOM | | TRUE | FALSE | FALSE | | FALSE |
| P11438 | P17047 | 0.67 | | | FALSE | TRUE | FALSE | | FALSE |
| P57722 | P60335 | 0.67 | | | FALSE | TRUE | FALSE | | FALSE |
| Q02566 | Q91Z83 | 0.671 | | | FALSE | TRUE | FALSE | | FALSE |
| P62821 | Q9D1G1 | 0.672 | | | FALSE | TRUE | FALSE | | FALSE |
| P29341 | Q3UEB3 | 0.673 | | HOM | TRUE | FALSE | FALSE | | FALSE |
| P58774 | Q6IRU2 | 0.673 | | | FALSE | TRUE | FALSE | | FALSE |
| P13707 | Q3ULJ0 | 0.676 | HOM | | TRUE | FALSE | FALSE | | FALSE |

Table S4 Mouse

| | | | | | | | | | | | |
|---|---|---|---|---|---|---|---|---|---|---|---|
| Q3KNY0 | Q5XKE0 | 0.676 | | | FALSE | TRUE | FALSE | | | | FALSE |
| P50153 | P63213 | 0.676 | | HET | FALSE | FALSE | FALSE | | | | FALSE |
| P56654 | Q05421 | 0.677 | | | FALSE | TRUE | FALSE | | | | FALSE |
| P97427 | Q9EQF6 | 0.677 | HET | HET | FALSE | FALSE | FALSE | | | | FALSE |
| Q14BI2 | Q9QYS2 | 0.678 | | | FALSE | TRUE | FALSE | | | | FALSE |
| P56593 | Q64458 | 0.678 | | | FALSE | TRUE | FALSE | | | | FALSE |
| Q7TMM9 | Q9CWF2 | 0.679 | HET | | FALSE | FALSE | FALSE | | | | FALSE |
| Q9CPW4 | Q9D898 | 0.68 | | | FALSE | TRUE | FALSE | | | | FALSE |
| O54749 | P24456 | 0.681 | | HOM | TRUE | FALSE | FALSE | | | | FALSE |
| P61982 | P68254 | 0.681 | | | FALSE | TRUE | FALSE | | | | FALSE |
| P18826 | Q7TSH2 | 0.682 | HET | HET | FALSE | FALSE | TRUE | 6640 | Human | Phosphorylase kinase complex | FALSE |
| Q60996 | Q61151 | 0.683 | HET | HET | FALSE | FALSE | FALSE | | | | FALSE |
| P70402 | Q62234 | 0.684 | | HOM | TRUE | FALSE | FALSE | | | | FALSE |
| O08553 | Q9EQF6 | 0.685 | HOM | HET | TRUE | FALSE | FALSE | | | | TRUE |
| P23953 | Q64176 | 0.687 | | | FALSE | TRUE | FALSE | | | | FALSE |
| P61027 | P62823 | 0.689 | | | FALSE | TRUE | FALSE | | | | FALSE |
| O09158 | Q64459 | 0.689 | | | FALSE | TRUE | FALSE | | | | FALSE |
| Q62108 | Q811D0 | 0.689 | HOM | HOM | TRUE | FALSE | FALSE | | | | TRUE |
| P01899 | P01901 | 0.689 | | HOM | TRUE | FALSE | FALSE | | | | FALSE |
| P56656 | Q05421 | 0.689 | | | FALSE | TRUE | FALSE | | | | FALSE |
| P33267 | P56656 | 0.689 | | | FALSE | TRUE | FALSE | | | | FALSE |
| Q91WL5 | Q9DBW0 | 0.69 | | | FALSE | TRUE | FALSE | | | | FALSE |
| Q64176 | Q8VCU1 | 0.691 | | | FALSE | TRUE | FALSE | | | | FALSE |
| Q99L27 | Q9DCZ1 | 0.691 | HOM | | TRUE | FALSE | FALSE | | | | FALSE |
| Q99P30 | Q9CR24 | 0.691 | | | FALSE | TRUE | FALSE | | | | FALSE |

Table S4 Mouse

| | | | | | | | | | |
|---|---|---|---|---|---|---|---|---|---|
| Q9QYB5 | Q9QYC0 | 0.692 | | | FALSE | TRUE | FALSE | | FALSE |
| Q99PR8 | Q9JK92 | 0.692 | | | FALSE | TRUE | FALSE | | FALSE |
| Q925N0 | Q99JR1 | 0.693 | | | FALSE | TRUE | FALSE | | FALSE |
| Q91VT4 | Q99LB2 | 0.693 | HOM | | TRUE | FALSE | FALSE | | FALSE |
| P33267 | Q05421 | 0.694 | | | FALSE | TRUE | FALSE | | FALSE |
| A2AQP0 | Q6URW 6 | 0.695 | | | FALSE | TRUE | FALSE | | FALSE |
| P60521 | Q91VR7 | 0.696 | HOM | HOM | TRUE | FALSE | FALSE | | FALSE |
| P14602 | Q99PR8 | 0.697 | | | FALSE | TRUE | FALSE | | FALSE |
| Q61133 | Q64471 | 0.697 | HOM | | TRUE | FALSE | FALSE | | FALSE |
| P70175 | Q9JLB0 | 0.697 | HOM | | TRUE | FALSE | FALSE | | FALSE |
| Q8BK48 | Q8QZR3 | 0.698 | | | FALSE | TRUE | FALSE | | FALSE |
| O70443 | P21279 | 0.699 | | | FALSE | TRUE | FALSE | | FALSE |
| O70161 | O70172 | 0.699 | | | FALSE | TRUE | FALSE | | FALSE |
| P56656 | Q91W64 | 0.699 | | | FALSE | TRUE | FALSE | | FALSE |
| O35945 | Q8K009 | 0.7 | | | FALSE | TRUE | FALSE | | FALSE |
| P61205 | P84078 | 0.7 | | | FALSE | TRUE | FALSE | | FALSE |
| P60487 | Q8CHP8 | 0.701 | HOM | | TRUE | FALSE | FALSE | | FALSE |
| Q8BXB6 | Q9QXZ6 | 0.701 | | | FALSE | TRUE | FALSE | | FALSE |
| Q8BH59 | Q9DB41 | 0.702 | | | FALSE | TRUE | FALSE | | FALSE |
| P33267 | Q6XVG2 | 0.702 | | | FALSE | TRUE | FALSE | | FALSE |
| O08532 | Q9Z1L5 | 0.705 | | | FALSE | TRUE | FALSE | | FALSE |
| Q3UNZ8 | Q9DCS3 | 0.705 | | HOM | TRUE | FALSE | FALSE | | FALSE |
| P45952 | Q8JZN5 | 0.707 | HOM | | TRUE | FALSE | FALSE | | FALSE |
| Q64176 | Q8BK48 | 0.707 | | | FALSE | TRUE | FALSE | | FALSE |
| Q63880 | Q64176 | 0.707 | | | FALSE | TRUE | FALSE | | FALSE |
| Q62108 | Q9WV34 | 0.708 | HOM | HOM | TRUE | FALSE | FALSE | | TRUE |

Table S4 Mouse

| | | | | | | | | | | | |
|---|---|---|---|---|---|---|---|---|---|---|---|
| Q8VCT4 | Q91WG0 | 0.709 | | | FALSE | TRUE | FALSE | | | | FALSE |
| P02469 | Q61292 | 0.711 | | | FALSE | TRUE | FALSE | | | | FALSE |
| P51859 | Q9JMG7 | 0.712 | HOM | | TRUE | FALSE | FALSE | | | | FALSE |
| P07901 | P11499 | 0.712 | HOM | HOM | TRUE | FALSE | TRUE | 25,5199,5212,5234,5266,5268, 5269,5286 | Mouse,Human | 9S-cytosolic aryl hydrocarbon receptor non-ligand activated complex, Kinase maturation complex 1,Kinase maturation complex 2,IKBKB-CDC37-KIAA1967-HSP90AB1-HSP90AA1 complex, TNF-alpha/NF-kappa B signaling complex | FALSE |
| Q8K009 | Q9CZS1 | 0.712 | | | FALSE | TRUE | FALSE | | | | FALSE |
| P85094 | Q91V64 | 0.713 | | | FALSE | TRUE | FALSE | | | | FALSE |
| Q921F2 | Q99020 | 0.713 | HOM | | TRUE | FALSE | FALSE | | | | FALSE |
| P51667 | Q9QVP4 | 0.714 | | | FALSE | TRUE | FALSE | | | | FALSE |
| Q8BH64 | Q9Z0R4 | 0.715 | HOM, HET | HET | TRUE, FALSE | FALSE | FALSE | | | | FALSE |
| P19324 | P32261 | 0.715 | | HOM | TRUE | FALSE | FALSE | | | | FALSE |
| P62761 | Q8BNY6 | 0.716 | | HET | FALSE | FALSE | FALSE | | | | FALSE |
| Q922F4 | Q9ERD7 | 0.718 | | HET | FALSE | FALSE | FALSE | | | | FALSE |

Table S4 Mouse

| | | | | | | | | | | | |
|---|---|---|---|---|---|---|---|---|---|---|---|
| O89053 | Q9WUM3 | 0.719 | | HOM | TRUE | FALSE | FALSE | | | | FALSE |
| O08553 | P97427 | 0.72 | HOM | HET | TRUE | FALSE | FALSE | | | | TRUE |
| P68368 | P68373 | 0.72 | HET | | FALSE | FALSE | FALSE | | | | FALSE |
| Q3UJU9 | Q9DCV4 | 0.721 | | | FALSE | TRUE | FALSE | | | | FALSE |
| Q8QZR3 | Q8VCU1 | 0.722 | | | FALSE | TRUE | FALSE | | | | FALSE |
| P56593 | P56656 | 0.723 | | | FALSE | TRUE | FALSE | | | | FALSE |
| O55100 | Q8R191 | 0.723 | | | FALSE | TRUE | FALSE | | | | FALSE |
| Q9CWF2 | Q9D6F9 | 0.724 | | HET | FALSE | FALSE | FALSE | | | | FALSE |
| P63321 | Q61411 | 0.724 | | | FALSE | TRUE | FALSE | | | | FALSE |
| Q80SZ7 | Q9DAS9 | 0.724 | | | FALSE | TRUE | FALSE | | | | FALSE |
| P35762 | Q922J6 | 0.726 | HOM | | TRUE | FALSE | FALSE | | | | FALSE |
| P05132 | P68181 | 0.726 | HET,HOM | HET | FALSE,TRUE | FALSE | FALSE | | | | FALSE |
| Q80T41 | Q9WV18 | 0.727 | | | FALSE | TRUE | TRUE | 6437 | Human | GABBR1-GABBR2 complex | FALSE |
| P02463 | P08122 | 0.729 | | | FALSE | TRUE | FALSE | | | | FALSE |
| P48722 | Q61699 | 0.729 | HOM | HOM | TRUE | FALSE | FALSE | | | | FALSE |
| P97441 | Q60738 | 0.73 | | | FALSE | TRUE | FALSE | | | | FALSE |
| P62077 | Q9WVA2 | 0.732 | HET | HET | FALSE | FALSE | FALSE | | | | FALSE |
| E9PZQ0 | P11881 | 0.733 | HET | | FALSE | FALSE | FALSE | | | | FALSE |
| Q62452 | Q8BWQ1 | 0.735 | | | FALSE | TRUE | FALSE | | | | FALSE |
| P23953 | Q91WG0 | 0.736 | | | FALSE | TRUE | FALSE | | | | FALSE |
| Q99MZ7 | Q9WV68 | 0.737 | | | FALSE | TRUE | FALSE | | | | FALSE |
| Q8VCC1 | Q99LB2 | 0.737 | HOM | | TRUE | FALSE | FALSE | | | | FALSE |
| Q8R081 | Q921F4 | 0.738 | | | FALSE | TRUE | FALSE | | | | FALSE |
| Q8K023 | Q8VC28 | 0.738 | HOM | HOM | TRUE | FALSE | FALSE | | | | FALSE |
| P70694 | Q8VC28 | 0.738 | HOM | HOM | TRUE | FALSE | FALSE | | | | FALSE |

Table S4 Mouse

| | | | | | | | | |
|---|---|---|---|---|---|---|---|---|
| P52196 | Q99J99 | 0.74 | | | FALSE | TRUE | FALSE | FALSE |
| P56656 | Q64458 | 0.74 | | | FALSE | TRUE | FALSE | FALSE |
| P46638 | P53994 | 0.74 | | | FALSE | TRUE | FALSE | FALSE |
| A2AQP0 | P13542 | 0.74 | | | FALSE | TRUE | FALSE | FALSE |
| P00329 | P28474 | 0.742 | | HOM | TRUE | FALSE | FALSE | FALSE |
| Q5YD48 | Q91WT8 | 0.744 | | | FALSE | TRUE | FALSE | FALSE |
| P31938 | Q63932 | 0.744 | | | FALSE | TRUE | FALSE | FALSE |
| Q62086 | Q62087 | 0.745 | | | FALSE | TRUE | FALSE | FALSE |
| P04117 | P11404 | 0.745 | HOM | | TRUE | FALSE | FALSE | FALSE |
| Q9EQP2 | Q9Z0R4 | 0.746 | HET | HET | FALSE | FALSE | FALSE | FALSE |
| O35643 | Q9DBG3 | 0.747 | | HOM | TRUE | FALSE | FALSE | FALSE |
| P50544 | Q9JHI5 | 0.748 | HOM | | TRUE | FALSE | FALSE | FALSE |
| P37804 | Q9WVA4 | 0.75 | | | FALSE | TRUE | FALSE | FALSE |
| Q8BGZ1 | Q91X97 | 0.75 | | HET | FALSE | FALSE | FALSE | FALSE |
| O55106 | P58404 | 0.751 | | | FALSE | TRUE | FALSE | FALSE |
| Q61205 | Q61206 | 0.751 | | HOM | TRUE | FALSE | FALSE | FALSE |
| P50544 | Q9DBL1 | 0.752 | HOM | HOM | TRUE | FALSE | FALSE | FALSE |
| Q3TJD7 | Q8CI51 | 0.752 | | | FALSE | TRUE | FALSE | FALSE |
| P11714 | P24456 | 0.753 | HOM | HOM | TRUE | FALSE | FALSE | FALSE |
| Q9WVK4 | Q9Z0R4 | 0.756 | HET | HET | FALSE | FALSE | FALSE | FALSE |
| P33267 | Q91W64 | 0.757 | | | FALSE | TRUE | FALSE | FALSE |
| Q9CXT8 | Q9CZ13 | 0.758 | HET | HET | FALSE | FALSE | FALSE | FALSE |
| P11798 | Q6PHZ2 | 0.758 | | HET | FALSE | FALSE | FALSE | FALSE |
| P56593 | Q05421 | 0.759 | | | FALSE | TRUE | FALSE | FALSE |
| O35295 | P42669 | 0.761 | HET | HET | FALSE | FALSE | FALSE | FALSE |
| P56593 | Q91W64 | 0.761 | | | FALSE | TRUE | FALSE | FALSE |

Table S4 Mouse

| | | | | | | | | | | | |
|---|---|---|---|---|---|---|---|---|---|---|---|
| A2AS89 | Q61176 | 0.762 | | HOM | TRUE | FALSE | FALSE | | | | FALSE |
| P62835 | Q99JI6 | 0.763 | | | FALSE | TRUE | FALSE | | | | FALSE |
| P60335 | P61979 | 0.764 | | HOM | TRUE | FALSE | FALSE | | | | TRUE |
| P23953 | Q8VCT4 | 0.766 | | | FALSE | TRUE | FALSE | | | | FALSE |
| P62823 | P63011 | 0.767 | | | FALSE | TRUE | FALSE | | | | FALSE |
| Q921F2 | Q9JII5 | 0.769 | HOM | | TRUE | FALSE | FALSE | | | | FALSE |
| O88935 | Q64332 | 0.77 | | | FALSE | TRUE | TRUE | 2835 | Mouse | Profilin 2 complex | FALSE |
| P0C192 | Q9D1T0 | 0.771 | | | FALSE | TRUE | FALSE | | | | FALSE |
| Q6PHZ2 | Q923T9 | 0.771 | HET | | FALSE | FALSE | FALSE | | | | FALSE |
| P56593 | P56654 | 0.774 | | | FALSE | TRUE | FALSE | | | | FALSE |
| P01027 | P01029 | 0.775 | HOM | | TRUE | FALSE | FALSE | | | | FALSE |
| P61027 | P61028 | 0.777 | | | FALSE | TRUE | FALSE | | | | FALSE |
| P47199 | Q3UNZ8 | 0.778 | HOM | | TRUE | FALSE | FALSE | | | | FALSE |
| P11087 | Q01149 | 0.781 | | | FALSE | TRUE | FALSE | | | | FALSE |
| Q3KNY0 | Q62234 | 0.782 | | HOM | TRUE | FALSE | FALSE | | | | FALSE |
| O35639 | P07356 | 0.784 | | HOM | TRUE | FALSE | FALSE | | | | FALSE |
| P13542 | Q5SX40 | 0.784 | | | FALSE | TRUE | FALSE | | | | FALSE |
| O35728 | O88833 | 0.784 | | | FALSE | TRUE | FALSE | | | | FALSE |
| Q8BH44 | Q9WUM4 | 0.786 | | HOM | TRUE | FALSE | FALSE | | | | FALSE |
| P28652 | Q6PHZ2 | 0.787 | | HET | FALSE | FALSE | FALSE | | | | FALSE |
| O08788 | Q9D1E6 | 0.789 | HOM | HET | TRUE | FALSE | FALSE | | | | FALSE |
| Q91ZA3 | Q99MR8 | 0.789 | | | FALSE | TRUE | FALSE | | | | FALSE |
| P23927 | Q5EBG6 | 0.79 | HOM | HOM | TRUE | FALSE | FALSE | | | | FALSE |
| Q8BH64 | Q9WVK4 | 0.792 | HOM, HET | HET | TRUE, FALSE | FALSE | FALSE | | | | FALSE |
| P61027 | P63011 | 0.793 | | | FALSE | TRUE | FALSE | | | | FALSE |

Table S4 Mouse

| | | | | | | | | | | | |
|---|---|---|---|---|---|---|---|---|---|---|---|
| Q5XKE0 | Q62234 | 0.795 | | HOM | TRUE | FALSE | FALSE | | | | FALSE |
| P56395 | Q9CQX2 | 0.795 | | | FALSE | TRUE | FALSE | | | | FALSE |
| Q9CXN7 | Q9DCG6 | 0.797 | HOM | | TRUE | FALSE | FALSE | | | | FALSE |
| Q3UNX5 | Q9QXG4 | 0.798 | | | FALSE | TRUE | FALSE | | | | FALSE |
| Q3TJD7 | Q9JKS4 | 0.8 | | | FALSE | TRUE | FALSE | | | | FALSE |
| Q8K0E8 | Q8VCM7 | 0.8 | HOM | HOM | TRUE | FALSE | TRUE | 6417 | Human | Fibrinogen complex | FALSE |
| P56654 | Q64458 | 0.802 | | | FALSE | TRUE | FALSE | | | | FALSE |
| P26150 | Q61694 | 0.802 | | | FALSE | TRUE | FALSE | | | | FALSE |
| P56654 | P56656 | 0.803 | | | FALSE | TRUE | FALSE | | | | FALSE |
| Q9DBL1 | Q9JHI5 | 0.804 | HOM | | TRUE | FALSE | FALSE | | | | FALSE |
| Q5DQR4 | Q8K400 | 0.804 | | | FALSE | TRUE | FALSE | | | | FALSE |
| P50544 | Q8JZN5 | 0.81 | HOM | | TRUE | FALSE | FALSE | | | | FALSE |
| P60335 | Q61990 | 0.81 | | | FALSE | TRUE | FALSE | | | | FALSE |
| P12242 | P56501 | 0.811 | HOM | | TRUE | FALSE | FALSE | | | | FALSE |
| Q60668 | Q99020 | 0.812 | | | FALSE | TRUE | FALSE | | | | FALSE |
| P99024 | Q9ERD7 | 0.813 | HET | HET | FALSE | FALSE | FALSE | | | | FALSE |
| P27773 | Q91W90 | 0.814 | HET | | FALSE | FALSE | FALSE | | | | FALSE |
| P50171 | Q99LB2 | 0.815 | HOM | | TRUE | FALSE | FALSE | | | | FALSE |
| P33267 | P56593 | 0.817 | | | FALSE | TRUE | FALSE | | | | FALSE |
| Q5EBG6 | Q99PR8 | 0.818 | HOM | | TRUE | FALSE | FALSE | | | | FALSE |
| O35459 | Q9D7J9 | 0.819 | HOM | HOM | TRUE | FALSE | FALSE | | | | FALSE |
| Q61425 | Q8BMS1 | 0.82 | HOM | | TRUE | FALSE | FALSE | | | | FALSE |
| P14602 | Q5EBG6 | 0.821 | | HOM | TRUE | FALSE | FALSE | | | | FALSE |
| P47738 | Q8K009 | 0.824 | HOM | | TRUE | FALSE | FALSE | | | | FALSE |
| P48962 | Q3V132 | 0.826 | | | FALSE | TRUE | FALSE | | | | FALSE |

| | | | | | | | | | | | |
|---|---|---|---|---|---|---|---|---|---|---|---|
| P14824 | P97384 | 0.828 | | | FALSE | TRUE | FALSE | | | | FALSE |
| Q6R891 | Q9QZQ1 | 0.829 | | | FALSE | TRUE | FALSE | | | | FALSE |
| P50171 | Q91VT4 | 0.83 | HOM | HOM | TRUE | FALSE | FALSE | | | | FALSE |
| Q00897 | Q00898 | 0.831 | HOM | HOM | TRUE | FALSE | FALSE | | | | FALSE |
| P63101 | P68254 | 0.833 | | | FALSE | TRUE | FALSE | | | | FALSE |
| Q64176 | Q91WG0 | 0.834 | | | FALSE | TRUE | FALSE | | | | FALSE |
| Q63880 | Q8VCU1 | 0.836 | | | FALSE | TRUE | FALSE | | | | FALSE |
| Q8QZT1 | Q99JY0 | 0.838 | HOM | HOM | TRUE | FALSE | FALSE | | | | FALSE |
| P62874 | P62880 | 0.84 | HET | | FALSE | FALSE | FALSE | | | | FALSE |
| Q8BLQ9 | Q8R464 | 0.84 | HOM | HOM | TRUE | FALSE | FALSE | | | | FALSE |
| Q5EBG6 | Q9JK92 | 0.84 | HOM | | TRUE | FALSE | FALSE | | | | FALSE |
| Q07417 | Q9JHI5 | 0.842 | HOM | | TRUE | FALSE | FALSE | | | | FALSE |
| Q60668 | Q9JII5 | 0.843 | | | FALSE | TRUE | FALSE | | | | FALSE |
| Q8BWT1 | Q8QZT1 | 0.845 | | HOM | TRUE | FALSE | FALSE | | | | FALSE |
| P56959 | Q61545 | 0.846 | | | FALSE | TRUE | TRUE | 1332 | Human | Large Drosha complex | FALSE |
| Q63880 | Q8QZR3 | 0.847 | | | FALSE | TRUE | FALSE | | | | FALSE |
| P68254 | Q9CQV8 | 0.847 | | | FALSE | TRUE | FALSE | | | | FALSE |
| O88569 | Q8BG05 | 0.848 | | | FALSE | TRUE | FALSE | | | | FALSE |
| O88492 | Q8CGN5 | 0.851 | | | FALSE | TRUE | FALSE | | | | FALSE |
| Q8BKZ9 | Q8BMF4 | 0.853 | HET | HOM | TRUE | FALSE | FALSE | | | | FALSE |
| Q8BH95 | Q9WUR2 | 0.856 | HOM | HOM | TRUE | FALSE | FALSE | | | | FALSE |
| P45952 | Q9DBL1 | 0.857 | HOM | HOM | TRUE | FALSE | FALSE | | | | FALSE |
| Q8CI51 | Q9JKS4 | 0.86 | | | FALSE | TRUE | FALSE | | | | FALSE |
| O54724 | Q91VJ2 | 0.86 | | | FALSE | TRUE | FALSE | | | | FALSE |
| Q8BHZ0 | Q921M7 | 0.861 | | | FALSE | TRUE | FALSE | | | | FALSE |

Table S4 Mouse

| | | | | | | | | |
|---|---|---|---|---|---|---|---|---|
| P14069 | P50543 | 0.862 | | | FALSE | TRUE | FALSE | FALSE |
| P07310 | Q6P8J7 | 0.862 | | HOM | TRUE | FALSE | FALSE | FALSE |
| P28653 | Q9JK53 | 0.863 | | | FALSE | TRUE | FALSE | FALSE |
| Q63886 | Q8BWQ1 | 0.863 | | | FALSE | TRUE | FALSE | FALSE |
| Q9D7J9 | Q9WUR2 | 0.863 | HOM | HOM | TRUE | FALSE | FALSE | FALSE |
| P63101 | Q9CQV8 | 0.864 | | | FALSE | TRUE | FALSE | TRUE |
| O88451 | P51658 | 0.865 | | | FALSE | TRUE | FALSE | FALSE |
| O70571 | Q9JK42 | 0.869 | | HOM | TRUE | FALSE | FALSE | FALSE |
| P28652 | Q923T9 | 0.869 | | | FALSE | TRUE | FALSE | FALSE |
| Q8BH95 | Q9D7J9 | 0.869 | HOM | HOM | TRUE | FALSE | FALSE | FALSE |
| P51885 | Q9JK53 | 0.874 | | | FALSE | TRUE | FALSE | FALSE |
| P06728 | Q00623 | 0.877 | | HOM | TRUE | FALSE | FALSE | FALSE |
| Q99MQ4 | Q9JK53 | 0.878 | | | FALSE | TRUE | FALSE | FALSE |
| P28654 | Q9JK53 | 0.882 | | | FALSE | TRUE | FALSE | FALSE |
| P50544 | Q07417 | 0.882 | HOM | HOM | TRUE | FALSE | FALSE | FALSE |
| P45952 | Q9JHI5 | 0.883 | HOM | | TRUE | FALSE | FALSE | FALSE |
| Q60668 | Q921F2 | 0.883 | | HOM | TRUE | FALSE | FALSE | FALSE |
| Q07417 | Q9DBL1 | 0.888 | HOM | HOM | TRUE | FALSE | FALSE | FALSE |
| Q99020 | Q9JII5 | 0.891 | | | FALSE | TRUE | FALSE | FALSE |
| P45952 | P50544 | 0.892 | HOM | HOM | TRUE | FALSE | FALSE | FALSE |
| P28653 | P51885 | 0.897 | | | FALSE | TRUE | FALSE | FALSE |
| P28653 | P28654 | 0.898 | | | FALSE | TRUE | FALSE | FALSE |
| P54227 | P55821 | 0.898 | | | FALSE | TRUE | FALSE | FALSE |
| P05201 | P05202 | 0.898 | HOM | HOM | TRUE | FALSE | FALSE | FALSE |
| Q8BWT1 | Q99JY0 | 0.9 | | HOM | TRUE | FALSE | FALSE | FALSE |
| P14602 | P23927 | 0.903 | | HOM | TRUE | FALSE | FALSE | FALSE |

Table S4 Mouse

| | | | | | | | | |
|---|---|---|---|---|---|---|---|---|
| Q02788 | Q04857 | 0.904 | | | FALSE | TRUE | FALSE | FALSE |
| O35459 | Q8BH95 | 0.905 | HOM | HOM | TRUE | FALSE | FALSE | FALSE |
| P61979 | Q61990 | 0.91 | HOM | | TRUE | FALSE | FALSE | TRUE |
| P28653 | Q99MQ4 | 0.91 | | | FALSE | TRUE | FALSE | FALSE |
| P11798 | Q923T9 | 0.911 | | | FALSE | TRUE | FALSE | FALSE |
| Q63918 | Q91VJ2 | 0.917 | | | FALSE | TRUE | FALSE | FALSE |
| P11798 | P28652 | 0.926 | | | FALSE | TRUE | FALSE | FALSE |
| P51885 | Q99MQ4 | 0.932 | | | FALSE | TRUE | FALSE | FALSE |
| P47934 | Q924X2 | 0.936 | | | FALSE | TRUE | FALSE | FALSE |
| O35459 | Q9WUR2 | 0.937 | HOM | HOM | TRUE | FALSE | FALSE | FALSE |
| P45376 | P45377 | 0.938 | HOM | | TRUE | FALSE | FALSE | FALSE |
| P28654 | P51885 | 0.942 | | | FALSE | TRUE | FALSE | FALSE |
| O54724 | Q63918 | 0.947 | | | FALSE | TRUE | FALSE | FALSE |
| P45952 | Q07417 | 0.951 | HOM | HOM | TRUE | FALSE | FALSE | FALSE |
| P28654 | Q99MQ4 | 0.959 | | | FALSE | TRUE | FALSE | FALSE |
